# Virtual dynamics of spatiotemporal single-cell interactions

**DOI:** 10.64898/2026.11.04.764438

**Authors:** Zhenyi Zhang, Zihan Wang, Qinxue Tian, Han Hou, Jiaqi Han, Jiantao Shen, Yuhao Sun, Xueya Zhou, Qiangwei Peng, Qing Nie, Tiejun Li, Peijie Zhou

**Author notes:** These authors contributed equally (Zhenyi Zhang, Zihan Wang, Qinxue Tian, Han Hou, Jiaqi Han). Corresponding authors. (Q.N.), (T.L.), (P.Z.).

## Abstract

Building virtual cells and tissues requires reconstructing how interacting cell populations evolve in space and time, yet spatial omics experiments provide only sparse snapshots of this process. Here, we introduce **CytoBridge**, a computational method that reconstructs continuous virtual cell dynamics from discrete spatial transcriptomic data. Starting from a joint transcriptomic–spatial space, CytoBridge learns stochastic trajectories of cell states, positions, and population sizes through an unbalanced mean-field Schrödinger bridge formulation that explicitly incorporates cell–cell interaction as a dynamical driver. A time-varying interaction graph built from spatial proximity and ligand–receptor priors allows the velocity to be decomposed into an interaction component estimated from observed neighborhoods and ligand–receptor features, and an intrinsic-context component that captures intrinsic regulatory processes and unmeasured contextual influences. Applied to five datasets across three spatial transcriptomics platforms and spanning development, regeneration, and neurodegeneration, CytoBridge accurately generates tissue dynamics at unmeasured time points, follows ligand–receptor signaling through measured and unmeasured stages, and supports hypothesis-generating in silico perturbation. CytoBridge provides a general method for turning heterogeneous spatiotemporal transcriptomic measurements into continuously evolving virtual tissues.

## Introduction

A long-standing aspiration in computational biology is to build virtual cells and virtual tissues that reproduce how living systems develop, regenerate, and degenerate^1–6^. Such virtual representations would allow researchers to simulate developmental processes, predict the effect of perturbations in silico, and support computational studies of multicellular dynamics^7^. Realizing this vision requires moving beyond static cellular atlases toward continuous dynamical models that capture how individual cells change their transcriptional states, migrate through space, proliferate or die, and coordinate with their neighbors through intercellular signaling^8^. Spatial transcriptomics now provides increasingly resolved snapshots of gene expression and tissue architecture at multiple time points^9^, yet each cell is observed only once, and the data do not directly reveal the trajectories connecting states across time and space^10,11^. A central challenge, therefore, is to infer continuous spatiotemporal dynamics from these discrete molecular snapshots.

To achieve this goal, for snapshot scRNA-seq datasets, pseudotime^12–14^ and RNA velocity^15–24^ methods have been developed to recover local progression from single-time-point data. For temporally resolved scRNA-seq and time-series omics^25–27^, optimal transport (OT)^8,10,28–33^, flow matching^34–38^, unbalanced transport^39,40^, and Schrödinger bridge^41–47^ formulations couple cell populations across time and provide continuous or stochastic population evolution. UNAGI further models disease progression and in silico drug responses from time-series single-cell transcriptomes^48^. In spatial and spatiotemporal transcriptomics, dynamics unfold in a joint transcriptomic–spatial space, where local neighborhood structure can shape both transcriptional change and physical movement. Recent methods such as MIOFlow 2.0^49^, MOSCOT^28^, Spateo^50^, DeST-OT^51^, HM-OT^52^, STORIES^53^, and stVCR^54^ show that incorporating spatial coordinates can substantially improve cross-time alignment, coupling, and reconstruction. These technologies and models have advanced the reconstruction of dynamic spatiotemporal biological processes and provide a useful basis for ongoing efforts toward virtual-cell dynamics.

Yet a key limitation persists in the pursuit of virtual cells and virtual tissues. For biomedical translation, the value of a virtual tissue lies not only in reconstructing cellular states across space and time, but also in predicting how a diseased tissue evolves, which communication programs drive this evolution, and how the system may respond to molecular or cellular perturbations. This is particularly important in progressive disorders, where early spatially localized changes in cell state and intercellular signaling may influence pathology, inflammatory remodeling, or loss of tissue function^55,56^. Although current spatiotemporal methods have brought us closer to dynamic reconstruction, they do not yet provide predictive models of how interacting cellular populations collectively shape future tissue states or perturbation responses. Indeed, during development or disease progression, multicellular tissues do not evolve as collections of isolated cells. Instead, they are shaped by coordinated signaling and local interactions that drive cell-state transitions, spatial patterning, and collective tissue dynamics^57–65^. Building predictive virtual tissues, therefore, requires treating cell–cell interaction as an explicit driver of continuous dynamics.

Here we introduce CytoBridge, a computational method designed to learn continuous spatiotemporal dynamics from discrete spatial transcriptomic snapshots while treating cell–cell interaction as an explicit component of the dynamics. CytoBridge is built on an unbalanced mean-field Schrödinger bridge generative model defined in a joint transcriptomic–spatial space, allowing stochasticity and changes in cell number to be modeled together with spatially structured dynamics. To address partial observability caused by sliced spatial transcriptomics measurement, CytoBridge decomposes the drift into an interaction program, estimated on a time-varying cell–cell graph that integrates spatial proximity with ligand–receptor priors, and an intrinsic-context program that absorbs intrinsic regulatory progression together with unmeasured contextual influences.

We evaluate CytoBridge in two settings. First, we construct an atlas-grounded in silico spatiotemporal benchmark with ground truth for velocities, growth rates, and interaction structure, enabling direct evaluation of recovery under incomplete observation. We also evaluate interaction recovery using an independent simulator and assess the inferred dynamics against lineage-tracing and metabolic-labeling measurements in two single-cell datasets. We then apply CytoBridge to five atlases spanning three spatial transcriptomics platforms and multiple biological and biomedical contexts, including development, regeneration, and neurodegeneration. Across these analyses, CytoBridge interpolates unmeasured stages, reconstructs interaction-resolved velocity and signaling patterns, and predicts responses to in silico perturbations.

## Results

### Overview of CytoBridge

Time-series spatial transcriptomics samples different tissue sections at discrete time points, across which population sizes vary and cell–cell interactions shape both transcriptional programs and spatial rearrangements. CytoBridge learns a continuous spatiotemporal mapping from a sequence of discrete transcriptomic snapshots 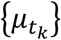 (Fig. 1a and Supplementary Fig. S1). Each snapshot is represented in a joint state space that couples a low-dimensional transcriptomic embedding **x** with spatial coordinates **r**, enabling CytoBridge to model transcriptional progression and spatial movement within a single framework (Methods).

**Figure 1:**
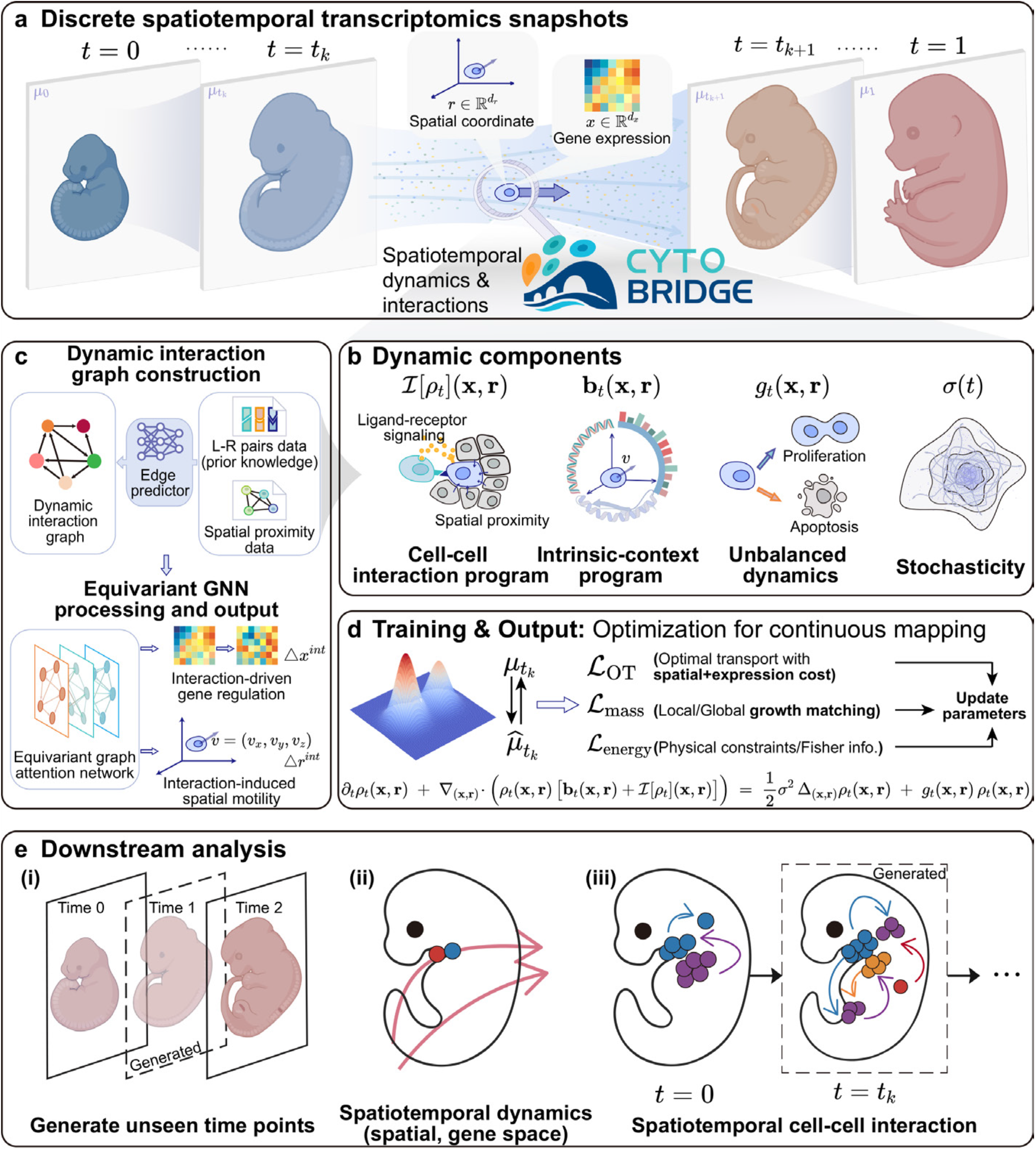
CytoBridge maps continuous spatiotemporal cell dynamics and interactions from discrete transcriptomic snapshots. **a**, Discrete spatiotemporal transcriptomics snapshots acquired at multiple sampling times {***t***_***k***_} are represented as empirical measures 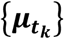 over a joint state space comprising transcriptomic states 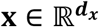 and spatial coordinates 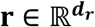. CytoBridge infers a continuous-time population evolution between snapshots. **b**, CytoBridge decomposes the dynamics into interpretable components: an interaction program ***ℐ***[***ρ***_***t***_](**x, r**) capturing cell–cell interaction effects, an intrinsic-context program **b**_***t***_(**x, r**) encoding intrinsic regulation together with unmeasured contextual influences, an unbalanced growth field ***g***_***t***_(**x, r**) modeling proliferation and loss, and stochasticity modeled through a time-dependent diffusion scale ***σ***(***t***). c, Construction of a time-varying interaction graph ***G***_***t***_ = (***V, ℰ***) by integrating prior knowledge (ligand–receptor pairs) with spatial proximity. A graph attention-style equivariant GNN predicts edge-dependent interactions and estimates how interactions contribute to transcriptomic and spatial velocity. **d**, Training and output. CytoBridge optimizes a continuous mapping that matches observed distributions at each ***t***_***k***_ while respecting the controlled mean-field Fokker–Planck dynamics of the unbalanced mean-field Schrödinger bridge (optimized in its Fisher-information probability-flow form). The objective combines a spatiotemporal transport loss (***ℒ***_**OT**_), local/global growth-matching (***ℒ***_**mass**_), and an action/regularization term (***ℒ***_**energy**_) derived from the underlying variational principle. **e**, Downstream analyses enabled by the learned continuous dynamics, including generating unmeasured stages, spatiotemporal dynamics, and continuous ligand–receptor interaction dynamics through space and time. In e(ii), red arrows denote inferred cell trajectories, whereas in e(iii), colored arrows indicate cell–cell interaction flows, with arrow colors matching the sender cell population.

At the core of CytoBridge is an unbalanced mean-field Schrödinger bridge (UMFSB) formulation expressed in a Fisher-information regularized probability-flow form (Methods). This formulation supports stochasticity in cellular dynamics, non-conservation of mass reflecting proliferation and loss, and mean-field interactions that couple cells through population-dependent forces (Fig. 1b). It yields a dynamics decomposition into four biologically interpretable components. The intrinsic-context program **b**_*t*_(**x, r**) represents intrinsic regulatory progression and unmeasured contextual influences, while the interaction term ℐ[*ρ*_*t*_](**x, r**) describes the contribution of neighboring cells to gene-expression and spatial velocity. The growth field *g*_*t*_(**x, r**) describes local population expansion or loss, and the diffusion scale *σ*(*t*) sets the magnitude of stochastic fluctuations (Methods, Supplementary Notes S1 and S2).

To model interactions without fixing an analytic kernel, CytoBridge parameterizes the interaction term with an attention-style equivariant graph neural network (GNN) on a time-varying cell–cell graph that combines spatial proximity with ligand–receptor (LR) pairing (Fig. 1c; Methods and Supplementary Note S3). Because the graph is rebuilt as cells move and change state, the interaction term evolves with the population (ℐ[*ρ*_*t*_]).

CytoBridge is trained in stages by optimizing a composite objective that aligns predicted intermediate distributions with observed snapshots while enforcing the underlying variational structure (Fig. 1d, Supplementary Note S4). The objective couples a spatiotemporal transport term that matches distributions across time, a mass-matching term that aligns local and global growth patterns, and an action/regularization term derived from the Fisher-information formulation (Methods, Supplementary Notes S2 and S4). The learned dynamics generate intermediate cell populations and describe changes in velocity, growth and ligand–receptor interactions over time (Fig. 1e; Methods). Supplementary Note S3.3 uses the UMFSB action to establish conditions for reconstructed population distributions to converge to the true distributions as observations become increasingly dense and accurate. Within a specified model class, it further gives conditions for the unique and stable recovery of intrinsic-context velocity, growth and interaction.

### Evaluation of CytoBridge in simulations and single-cell datasets

Real datasets do not measure continuous trajectories or interaction forces, so to evaluate recovery of continuous dynamics we constructed an atlas-grounded in silico spatiotemporal (AGIST) benchmark with known velocities, growth rates and interaction networks (Methods, Supplementary Note S5). A CytoBridge model pretrained on the MOSTA mouse organogenesis atlas simulated the continuous evolution of 5,000 Brain-annotated cells, and snapshots at four equally spaced time points were used to train a new CytoBridge model (Fig. 2a).

**Figure 2:**
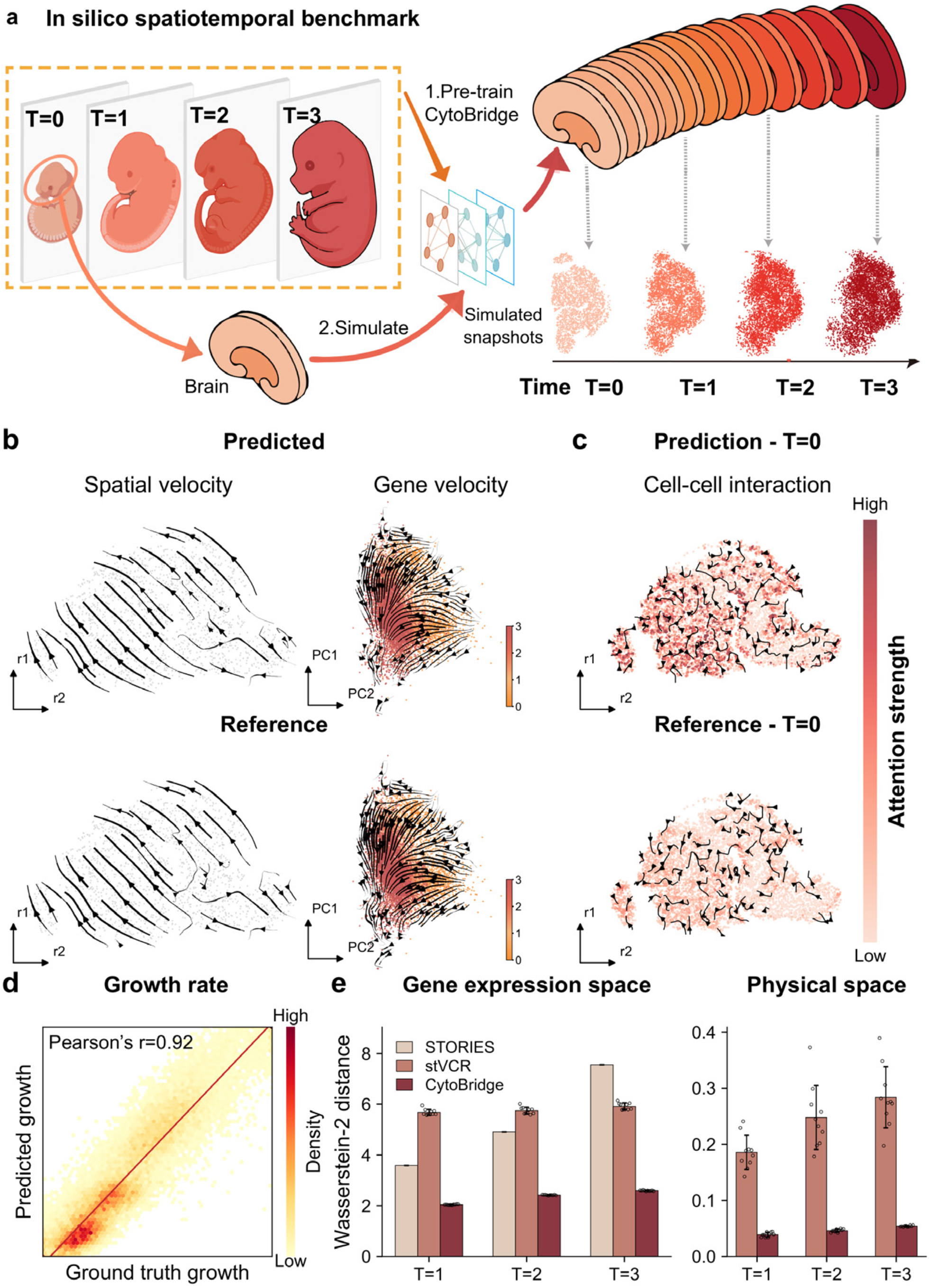
Evaluation of CytoBridge on an atlas-grounded in silico spatiotemporal (AGIST) benchmark. **a**, Overview of the AGIST benchmark. The CytoBridge model pre-trained on a mouse organogenesis atlas is used to simulate continuous brain development (upper right). Starting from 5,000 Brain-annotated cells, the simulation yields 5,000, 7,637, 9,733 and 10,667 cells at the four sampled time points. **b**, Comparison of inferred versus ground truth velocity fields. Streamlines depict the spatial velocity (left) and gene expression velocity (right). **c**, Recovery of cell–cell interaction patterns. Maps showing the spatial distribution of attention strengths in the predicted model versus the ground truth at *T* = 0. Black arrows indicate attention-weighted directions between cells. The spatial maps show the main tissue region. **d**, Evaluation of growth rate inference. Comparison of predicted and ground-truth growth rates for individual cells. Each distribution is clipped at its first and 99th percentiles and scaled to the interval from zero to one for display. **e**, Wasserstein-2 distance between inferred and ground-truth distributions in gene-expression space (left) and physical space (right). For CytoBridge and stVCR, each open circle shows the Wasserstein-2 distance for one of ten predictions with different random seeds from one trained model per method. Bars and error bars show the mean and standard deviation across these predictions. The stVCR predictions include stochastic cell division and death. STORIES supplies one deterministic gene-state prediction per target time. Each method was trained on all four snapshots, and all predictions started from the same 5,000 initial cells. Across the ten repeated predictions, CytoBridge had lower Wasserstein-2 distance than each comparison method at all evaluated time points (two-sided sign-flip tests, Holm-adjusted P = 0.0176 across nine comparisons).

We applied CytoBridge to the benchmark snapshots and reconstructed the underlying continuous dynamics. The inferred full velocity fields agreed with the generator fields, with aggregate cosine similarity of 0.977 in physical space and 0.952 in gene-expression space (Fig. 2b). The recovered attention strengths were correlated with the generator values, with a mean Spearman correlation of 0.80 across the four time points (Fig. 2c and Supplementary Fig. S2b). Inferred and generator growth values were also correlated (Pearson’s r = 0.92; Fig. 2d and Supplementary Fig. S2a). Median velocity agreement remained high across the evaluated time points and expression-state clusters (Supplementary Fig. S3).

We compared CytoBridge with two spatiotemporal trajectory inference methods, stVCR and STORIES, on the same snapshots. Across ten stochastic predictions from each trained CytoBridge and stVCR model, CytoBridge had a lower Wasserstein-2 distance than stVCR in physical and gene-expression space, and than STORIES in gene-expression space, at all three evaluated time points (Fig. 2e). It also tracked the relative population mass more closely than both methods (Supplementary Fig. S2c).

Because the AGIST generator is itself a CytoBridge model, we also tested recovery on an independent interacting-particle simulator with finite-range attraction, diffusion and population growth (Supplementary Fig. S4, Supplementary Note S6). CytoBridge recovered the radial attraction profile (Pearson r = 0.876) and the increase in population mass, and retaining the inferred interaction during propagation lowered the final-time W1 distance in both physical and gene-expression space compared with switching it off.

These simulations provide ground truth but not real biology. We therefore tested the inferred dynamics against measurements withheld from training in two non-spatial single-cell time courses, with interaction graphs built from ligand–receptor information in gene-expression space (Supplementary Figs. S5 and S6, Supplementary Note S7). In the LARRY mouse hematopoiesis dataset^66^, lineage barcodes were used only to evaluate the predicted fate proportions of each clone. The full model had lower mean distributional error at days 4 and 6 and higher agreement with the observed clone-fate proportions than the separately trained no-interaction model (Supplementary Fig. S5c,d). In the scNT-seq cortical activation dataset^67^, CytoBridge was trained on total RNA and evaluated against transcriptional directions estimated from metabolically labeled new RNA. Across three training runs per condition, the full model had lower mean distributional error at all four evaluated endpoints and a similar percentage of cells aligned with the new-RNA direction (Supplementary Fig. S6c, d).

### Benchmarking CytoBridge and evaluating its inferred cell–cell interactions

We benchmarked CytoBridge in five spatial datasets: chicken-heart development^68^, mouse organogenesis (MOSTA)^69^, axolotl brain regeneration (ARTISTA)^70^, Alzheimer’s disease (AD) progression in mice^71^, and zebrafish development^72^. For each dataset, we excluded one intermediate time point at a time from model fitting and compared the predicted cell population at that time with the withheld observations. This leave-one-time-point-out benchmark compared CytoBridge with stVCR^54^, MOSCOT^28^, PASTE2^73,74^, Spateo^50^, SpaTrack^75^, MIOFlow 2.0^49,76^, STORIES^53^, and Waddington-OT^29^, together with Linear OT displacement and random-interpolation controls. Prediction error was measured using Sliced-W2 in physical space, gene-expression space, and their joint space, depending on the outputs available for each method. Across the 33 combinations of held-out stage and evaluation space, CytoBridge had the lowest Sliced-W2 in 12, more than any other method or control, while providing interaction-resolved continuous dynamics not modeled by the other methods compared here (Supplementary Figs. S7 and S8, and Supplementary Note S4.8).

Removing the ligand–receptor prior during training or switching off the learned interaction term during prediction increased mean reconstruction error in all five datasets (Supplementary Fig. S9, Supplementary Note S4.8). The inferred interactions were also stable in zebrafish: cell-type interaction rankings from five independently trained models had a median pairwise Spearman correlation of 0.980, and changing the spatial-neighborhood radius or loss weights gave correlations of 0.963–0.997 (Supplementary Fig. S10, Supplementary Note S4.7).

To compare the inferred interactions with established communication tools, we summarized them by sender and receiver cell type and compared their rankings with COMMOT^59^ and CellAgentChat^77^ in chicken heart, MOSTA, ARTISTA and zebrafish (with CellAgentChat run using the same ligand–receptor database), using the full rankings and the overlap among the highest-ranked 20*k* of pairs. Rank correlations were positive in all four datasets, with median Spearman correlations of 0.84 for COMMOT and 0.45 for CellAgentChat (Supplementary Fig. S11a). We then combined the inferred interaction contributions with ligand expression in sender cells and receptor expression in receiver cells to identify candidate ligand–receptor axes. Representative axes selected by CytoBridge were also highly ranked by COMMOT within the corresponding sender–receiver pairs (Supplementary Fig. S11b). We further used NicheNet^57^ to examine candidate downstream targets of selected ligands in receiver cells (Supplementary Fig. S11c and Supplementary Note S8.9).

### CytoBridge uncovers fate transitions and interaction remodeling in the developing chicken heart

We applied CytoBridge to an integrated embryonic chicken heart dataset^68^ generated using 10x Visium spatial transcriptomics at embryonic days 4, 7, 10, and 14 (D4, D7, D10, and D14). We selected one tissue section per stage before alignment, retaining 3,550 Visium spots in total. Heart sections from the four stages were aligned into a common spatial coordinate system (Methods, Supplementary Fig. S12a), and the aligned geometry and velocity patterns changed little under small coordinate perturbations (Supplementary Figs. S12b and S13). Using these snapshots, CytoBridge reconstructs the continuous development of the chicken heart, predicting changes in cell states and spatial organization between the observed stages (Fig. 3a, Supplementary Fig. S14).

**Figure 3:**
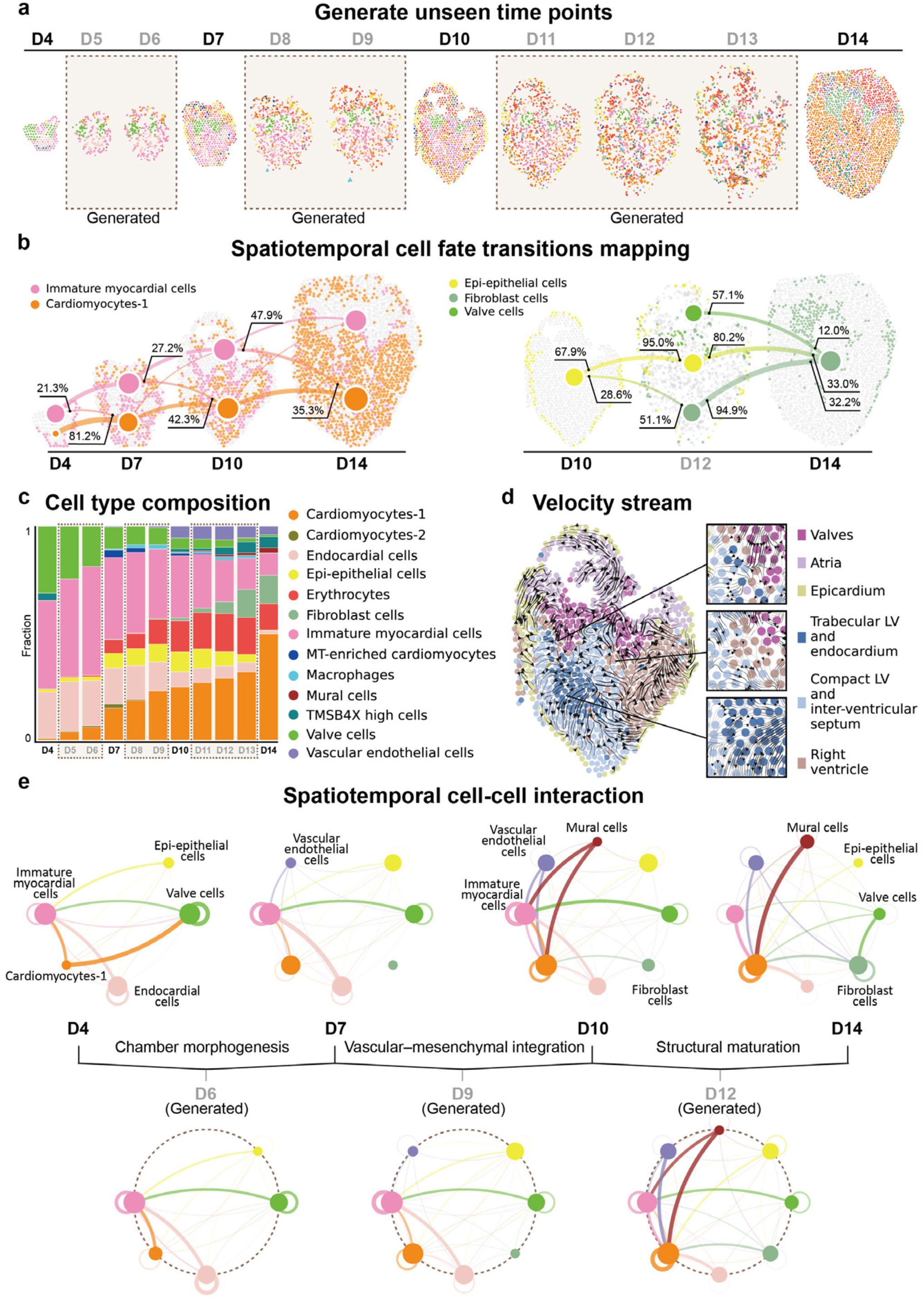
CytoBridge enables continuous spatiotemporal reconstruction of embryonic chicken heart development. **a**, Generation of unobserved developmental stages between experimentally sampled time points. Generated stages are marked by light gray time labels and dashed boxes. **b**, Spatiotemporal fate-transition map inferred from the learned continuous dynamics. Node size denotes the log-transformed relative abundance of each cell type at each stage. Directed edges indicate transition probabilities. Edge width scales with transition strength, and percentage labels report the outbound fraction from the source and the inbound contribution to the target. Generated stages are shown with light gray time labels. **c**, Cell-type composition dynamics across development. Stacked bars show relative cell-type fractions at each stage. Generated stages are highlighted with light gray time labels and dashed boxes. **d**, Full gene-expression velocity at D10 projected onto tissue coordinates. Streamlines show the inferred directions of transcriptional change, with inset panels enlarging local flow patterns. **e**, Time-resolved cell–cell interaction network inferred by CytoBridge. Node size denotes log-transformed relative abundance. Edges represent interaction strength (width and transparency), with edge color indicating the signaling source cell type. For clarity, only top-ranked interactions are emphasized, with weaker interactions shown as background. Dashed circles and light gray time labels mark generated stages (D6, D9, and D12).

CytoBridge generates intermediate stages (D5, D6, D8, D9, D11, D12, and D13) embedded in a common spatial space, with the resulting sections retaining the global heart geometry (Fig. 3a). Cell-type compositions vary smoothly along the reconstructed timeline (Fig. 3c), with generated stages occupying intermediate fractions between adjacent time points. Cell-type-specific growth rates were also inferred through the model (Supplementary Fig. S14). Using the learned continuous flow, we constructed a directed transition map across time points (Fig. 3b). Within the myocardial lineage, CytoBridge identifies a dominant transition from immature myocardial cells toward the more mature Cardiomyocytes-1 state. Introducing the generated intermediate stage D12 between D10 and D14 further resolves mesenchymal diversification, revealing a stepwise transition from epicardial epithelial (epi-epithelial) cells toward fibroblasts and an additional contribution from valve cells between D12 and D14, suggesting a valve-associated fibroblast-like mesenchymal contribution^78,79^.

To relate discrete fate transitions to continuous transcriptional dynamics, we projected the inferred full gene velocity field onto D10 tissue coordinates and identified streamline patterns that varied across anatomical compartments (Fig. 3d). Specifically, streamlines showed a local directional trend from trabecular ventricular myocardium toward the adjacent compact ventricular wall and septum. At valve-adjacent interfaces, streamlines tended to extend from valve-rich regions toward neighboring compartments, suggesting local boundary remodeling during valve-associated morphogenesis. In epicardial, atrial, and ventricular regions, flow patterns were largely local and spatially restricted, consistent with regional differences in gene expression during heart development^68^. The model further decomposes the inferred gene-level velocity into distinct components, capturing intrinsic drift dynamics and cell–cell interaction effects, together with their relative magnitudes and directional agreement (Supplementary Fig. S15a,b). In the observed D14 section, CytoBridge and veloAgent^24^ showed a similar flow from the trabecular left ventricle toward the compact left ventricle (Supplementary Fig. S15c). This qualitative agreement is consistent with coordinated maturation of ventricular myocardial states.

CytoBridge also infers a dynamic spatiotemporal cell–cell interaction network at each developmental stage (Fig. 3e). At D4, the network is relatively sparse and is dominated by interactions among endocardial, valvular, epi-epithelial, and immature myocardial compartments, with prominent valve and endocardial self-coupling together with strong bidirectional interactions between endocardial/valve cells and immature myocardium. By D7, the network becomes more integrated while remaining centered on immature myocardial cells, which receive strengthened inputs from endocardial, valvular, epi-epithelial, and vascular endothelial compartments, indicating increasing coordination between myocardial and neighboring non-myocardial cell states. At D10, mural-, fibroblast-, and vascular endothelial-associated interactions become more prominent, particularly in their coupling to myocardial cells, while valve-associated interactions remain evident. By D14, the network adopts a denser and more interconnected topology, with strong cardiomyocyte and fibroblast self-coupling, prominent valve-to-fibroblast interactions, and persistent endothelial-, epithelial-, and fibroblast-associated interactions with myocardial compartments. The generated intermediate stages D6, D9, and D12 display connectivity patterns between those of adjacent observed stages, consistent with gradual remodeling of cell–cell interaction networks during heart development^80,81^.

### CytoBridge reconstructs interaction-resolved spatiotemporal landscape in mouse organogenesis

We applied CytoBridge to the MOSTA mouse organogenesis Stereo-seq atlas^69^ comprising 344,603 cells across embryonic days E12.5 to E15.5 to test whether a continuous spatiotemporal dynamical system learned from discrete snapshots can yield coherent interpolation and interaction-resolved dynamics (Fig. 4). CytoBridge generated intermediate stages (E13.0, E14.0, and E15.0) that preserved the global embryonic layout and boundaries of major compartments while producing smooth transitions between adjacent observed time points (Fig. 4a). The interpolated maps preserve coherent tissue geometry across the developmental series, and growth maps of brain cells show the expected decline in proliferative activity as development proceeds (Supplementary Figs. S16 and S17).

**Figure 4:**
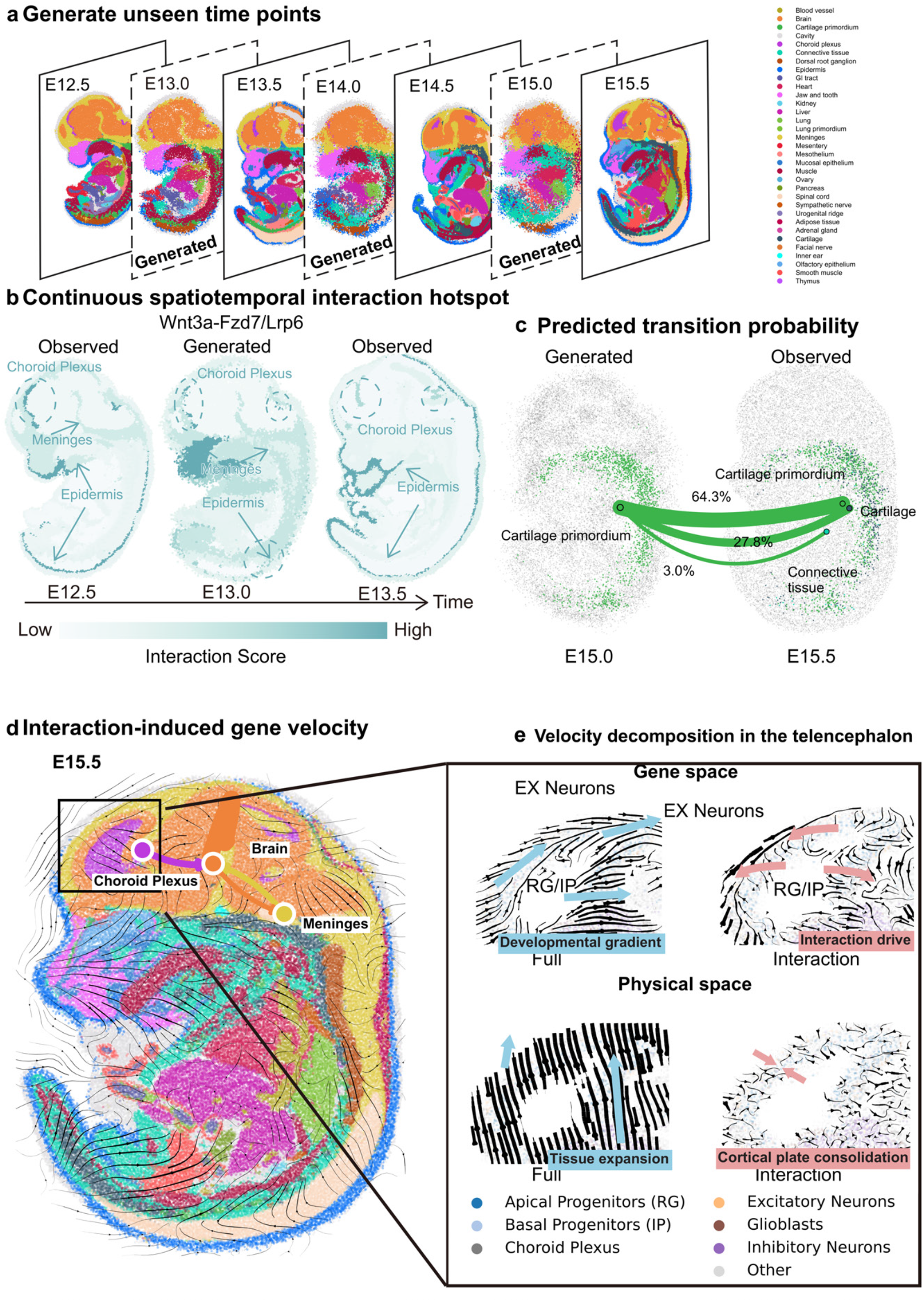
CytoBridge reconstructs continuous spatiotemporal dynamics, interactions, and fate flow during mouse organogenesis. **a**, Generation of unmeasured developmental stages from discrete spatial transcriptomic snapshots. CytoBridge interpolates E13.0, E14.0, and E15.0 (dashed) between observed stages E12.5–E15.5. **b**, Spatial patterns of the *Wnt3a*–*Fzd7*/*Lrp6* interaction score at E12.5, generated E13.0, and observed E13.5. Colors are normalized to the 1st–99th percentiles within each stage and show the spatial distribution of the score. **c**, Cell-state assignments after propagation from E15.0 to E15.5 for **1**,**464** generated cells originating from cartilage primordium. Cell types were assigned by the classifier. The three largest target groups were Cartilage primordium (64.3*k*), Cartilage (27.8*k*), and Connective tissue (3.0*k*). Arrow width indicates transition proportion. **d**, Interaction-associated gene-expression velocity at E15.5, shown as streamlines over the spatial map together with selected interaction links among choroid plexus, brain, and meninges. **e**, Velocity decomposition in the developing telencephalon. Gene-space views compare the full gene-expression velocity with the interaction component across radial glia/intermediate progenitors and excitatory neurons. Physical-space views compare full tissue expansion with the interaction-associated cortical-plate pattern. Colors denote the indicated major cell classes.

From the reconstructed dynamics, we mapped spatial interaction hotspots using the attention weights from the interaction network (Methods). For the *Wnt3a*–*Fzd7*/*Lrp6* axis, CytoBridge identified a dorsal interaction focus in the head region, with enrichment around the cortical hem–choroid plexus–meningeal vicinity and extensions along the dorsal epidermis (Fig. 4b). This localization agrees with the known role of the dorsal telencephalic midline as a Wnt-rich organizer of adjacent cortical and hippocampal territories^82^. The generated stages describe how this *Wnt3a*–*Fzd7*/*Lrp6* pattern changes between the observed sections (Fig. 4b).

CytoBridge also inferred a directed transition structure among mesenchymal-derived populations. Between E15.0 and E15.5, 1,464 generated cells classified as cartilage primordium were assigned mainly to Cartilage primordium (64.3%), Cartilage (27.8%), and Connective tissue (3.0%) (Fig. 4c), supporting persistence together with branching toward cartilage and neighboring connective-tissue states^83^. These transitions are accompanied by coordinated shifts in absolute cell abundance and composition, while major lineage connections remain continuous across interpolated stages (Supplementary Figs. S18 and S19).

We next examined how the learned interaction program contributes to local velocity patterns in the developing brain. At E15.5, the interaction component of velocity revealed coherent streams in the developing brain that spanned tissue boundaries and aligned with spatially adjacent brain–choroid plexus– meningeal structures (Fig. 4d). Within the boxed telencephalic region, where brain-annotated cells were further subclassified into telencephalic cell types (Supplementary Note S8.13), decomposing velocities separated full developmental gradients from interaction-linked effects. In gene space, the full velocity followed a canonical progression from radial glia/intermediate progenitors (RG/IP) toward excitatory neuronal states^84^, whereas the interaction component showed locally varying directions around the RG/IP region (Fig. 4e), suggesting that cell–cell interactions modulate local cell-state transitions during cortical development. In physical space, the full velocity emphasized global tissue expansion, while the interaction component of velocity was enriched along the cortical plate and pointed inward (Fig. 4e), suggesting interaction-associated consolidation of the cortical wall during mid-gestation development.

Along the reconstructed Brain trajectory, gene programs split into early-decreasing and late-increasing patterns (Supplementary Figs. S20 and S21). Ordering genes by peak expression time revealed three waves, from nuclear and metabolic functions through signaling to cell adhesion and extracellular-matrix organization (Supplementary Fig. S22), and ligand–receptor profiles grouped into three temporal patterns (Supplementary Fig. S23).

### CytoBridge maps coordinated spatiotemporal organization during axolotl brain regeneration

We applied CytoBridge to the ARTISTA Stereo-seq dataset of axolotl brain regeneration following surgical removal of a portion of the dorsal pallium in the left telencephalic hemisphere^70^. We analyzed one annotated section at each of 2, 5, 10, 15, and 20 days post injury (DPI), comprising 46,199 cells. CytoBridge embeds the annotated populations into a shared spatiotemporal coordinate system and reconstructs continuous trajectories across regeneration (Fig. 5a). Using the learned dynamics, CytoBridge generates intermediate, unmeasured stages (Fig. 5b and Supplementary Fig. S24).

**Figure 5:**
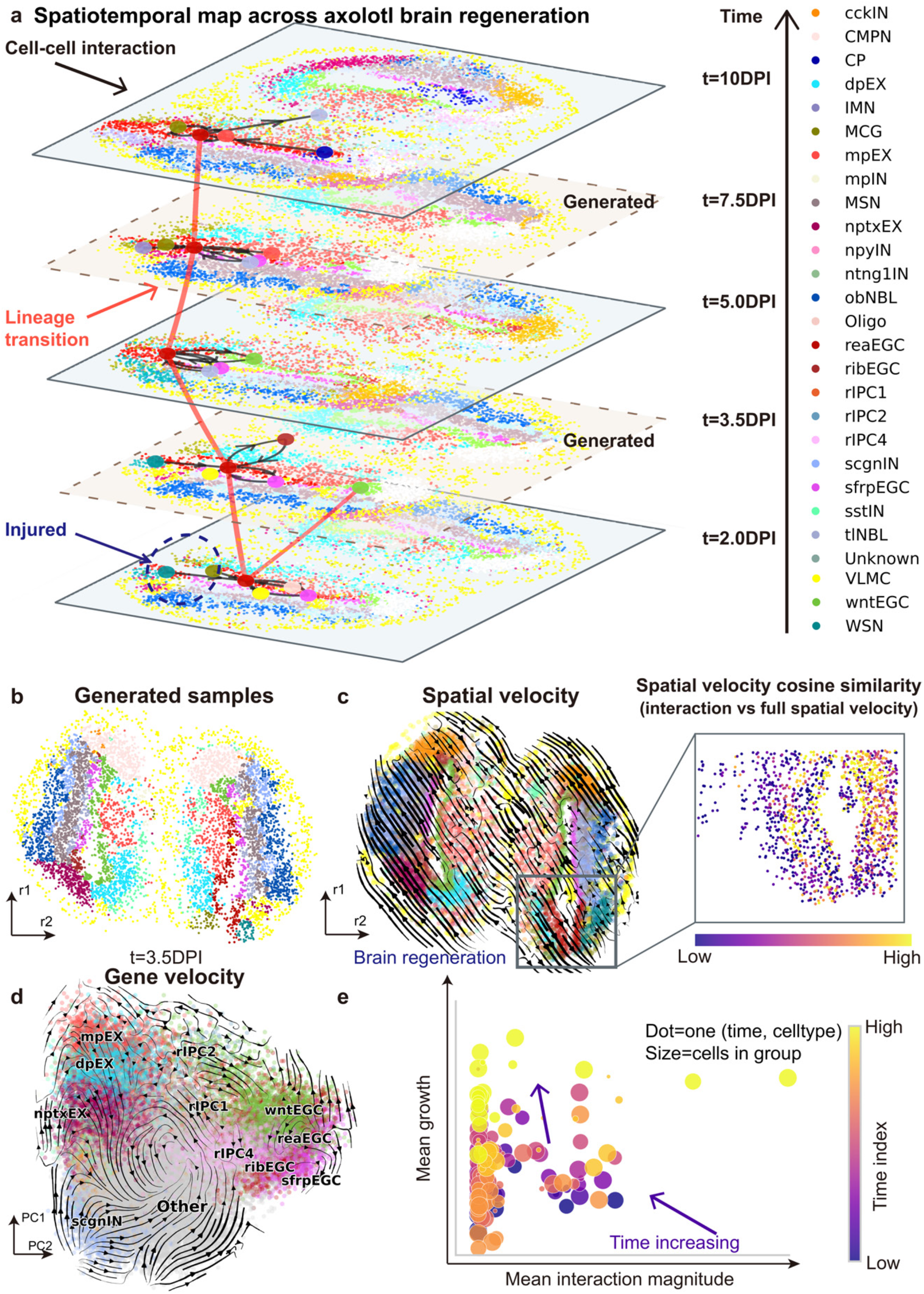
CytoBridge maps spatiotemporal dynamics, lineage transitions, and cell–cell interaction during axolotl brain regeneration. **a**, Joint spatiotemporal map of annotated populations at 2.0, generated 3.5, 5.0, generated 7.5, and 10 DPI. Red trajectories show directed state-transition summaries and black curves show the five strongest displayed reaEGC-associated interaction edges at each stage. The dashed outline indicates the approximate injury region. **b**, Representative generated cells at 3.5 DPI projected in spatial coordinates (r1, r2). **c**, Full spatial velocity visualized as streamlines over the tissue map. Right, cellwise cosine similarity between the full spatial velocity and the spatial component of interaction velocity within the boxed region. Higher values indicate closer directional agreement. **d**, Intrinsic-context gene-expression velocity streamlines in transcriptomic principal-component space. Local directions vary across annotated populations. **e**, Mean interaction magnitude and mean growth for each cell type at each time point (dot size indicates the number of cells and color indicates the time index). Later time points show increased growth while interaction magnitude remains heterogeneous.

Within this unified spatiotemporal map, CytoBridge inferred directed transition structures and cell–cell interaction edges. For the reactive ependymoglial cell (reaEGC) population, selected high-scoring interaction edges varied across time and space, with the approximate injury region marked by a dashed outline (Fig. 5a). This heterogeneity is compatible with the localized reactive and regenerative states reported for axolotl ependymoglial populations^70,85^. The transition summary showed substantial reaEGC self-retention together with time-dependent outgoing assignments to the regeneration intermediate progenitor cell (rIPC) subtypes rIPC1 and rIPC4 and to *Wnt8b*-expressing ependymoglial cells (wntEGCs; Fig. 5a, Supplementary Fig. S25). These predicted transitions are consistent with the reported contribution of ependymoglial populations to regenerated neuronal diversity^86^.

CytoBridge also estimates the full spatial velocity field across the regenerating telencephalon. The inferred streamlines suggest coordinated spatial rearrangement of cells during tissue repair (Fig. 5c). Cellwise cosine similarity between the full field and the interaction-program spatial component was spatially heterogeneous within the displayed region (Fig. 5c, right). To test whether the heterogeneous field contained organized local programs, we defined two spatially connected 5-DPI domains before evaluating their model edges (Supplementary Fig. S26). Attention in the two domains was 1.69-fold and 3.06-fold that of random cell sets with the same cell-type composition (both P = 0.0001), and their candidate ligand– receptor programs differed, with matrix and trophic signals in the sfrpEGC–VLMC-associated domain (sfrpEGC, *Sfrp1*-expressing ependymoglial cell; VLMC, vascular leptomeningeal cell) and reactive, adhesion and guidance signals in the reaEGC–wntEGC-associated domain.

In transcriptomic principal-component space, the intrinsic-context component of the CytoBridge-inferred gene-expression velocity field suggested local progression from reaEGCs toward rIPC states and from rIPCs toward excitatory neurons (Fig. 5d). These trends are consistent with the reported neurogenic role of ependymoglial cells during regeneration^70^. Reconstructed gene trajectories formed two temporal patterns. Genes in the first pattern were enriched for synaptic signaling, including glutamatergic transmission, and synapse organization (Supplementary Fig. S27). Ligand–receptor scores across the nine observed and generated stages grouped into two temporal programs (Supplementary Figs. S28 and S29).

To analyze population-level dynamics, we summarized each cell type at each time point by its mean interaction magnitude and mean growth, with point size reflecting the number of cells. At early stages, several cell populations showed strong interaction activity alongside relatively low growth. By 20 DPI, higher growth rates were observed across populations with a range of interaction magnitudes (Fig. 5e). Growth maps across observed and generated stages show higher inferred growth in the injured hemisphere, including the injury region (Supplementary Fig. S30).

### CytoBridge reveals continuous spatiotemporal microglial activation and signaling trajectories in AD

To resolve microglial state changes during Alzheimer’s disease (AD) progression, we applied CytoBridge to Xenium spatial transcriptomics data^71^ from TgCRND8 mice profiled at 2.5, 5.7, and 17.9 months (Fig. 6a). Across 172,092 cells, CytoBridge reconstructed a continuous trajectory spanning early, intermediate, and late disease stages (Fig. 6b, Supplementary Fig. S31a–c).

**Figure 6:**
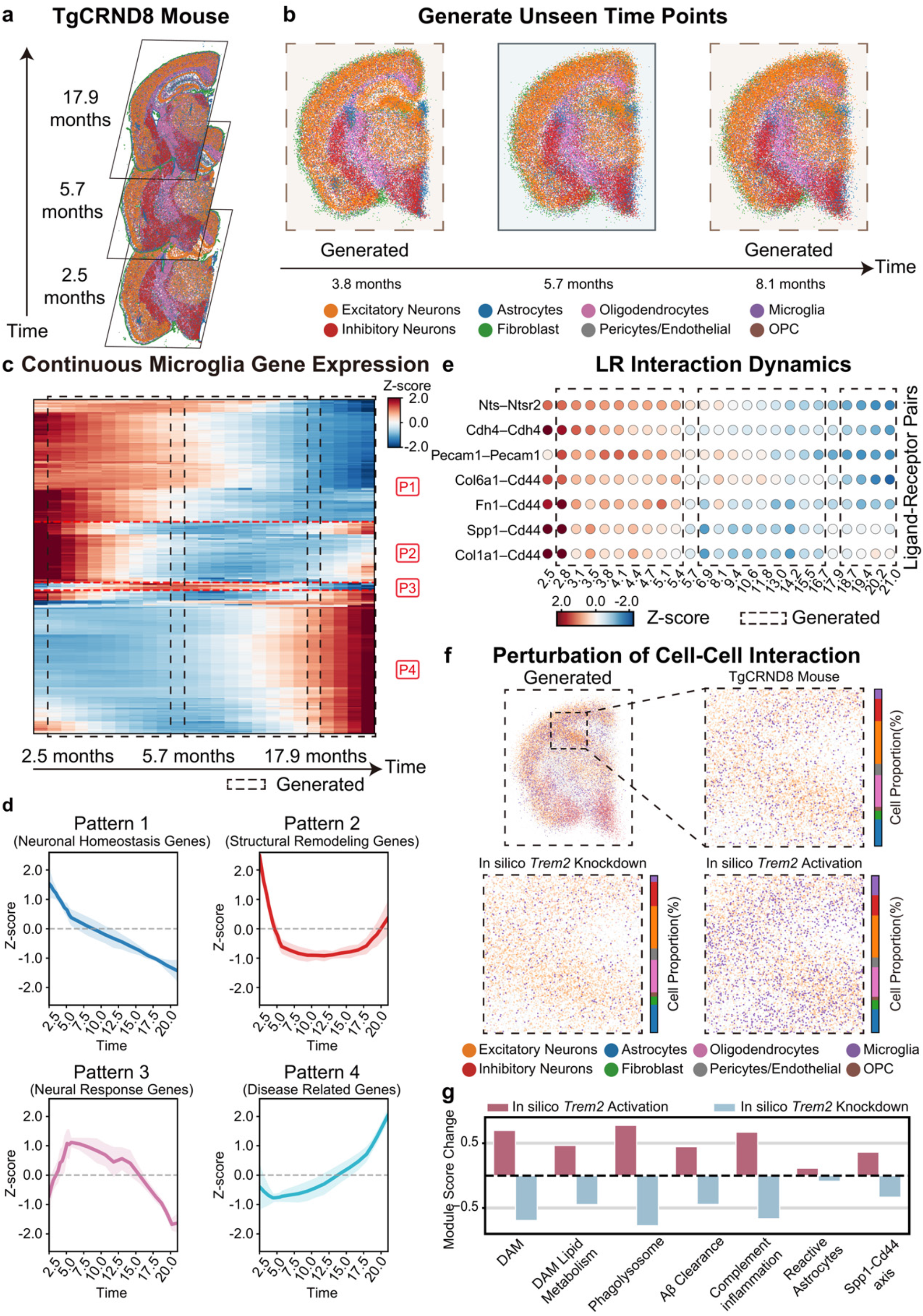
CytoBridge reconstructs spatiotemporal changes during Alzheimer’s disease progression in TgCRND8 mice. **a**, TgCRND8 brain sections at 2.5, 5.7, and 17.9 months. **b**, Generated intermediate states at 3.8 and 8.1 months between the observed sections. **c**, Reconstructed microglial gene-expression dynamics across the inferred disease trajectory. Dashed boxes mark generated intervals. **d**, Four temporal gene-expression patterns (P1–P4). Lines show the mean Z-scored expression and shading shows ± s.d. across genes in each pattern. **e**, Temporal profiles of selected model-derived ligand–receptor scores. In c– g, predictions after the last observed age of 17.9 months are extrapolated. **f**, Generated cell-type distributions and cell-type proportions for the reference, in silico *Trem2* knockdown, and in silico *Trem2* activation. **g**, Changes in functional-module scores after *Trem2* perturbation relative to the unperturbed TgCRND8 reference.

Continuous-time analysis of microglial gene expression identified four temporal patterns across the inferred disease trajectory (Fig. 6c,d). Pattern 1 declined over time and was associated in exploratory Gene Ontology analysis with synaptic and neuronal processes (Supplementary Fig. S32). Pattern 4 increased at later ages and was enriched for immune effector processes, TNF-associated cytokine production, and inflammatory responses. *Apoe, Tyrobp, Trem2*, and *Ctsd* contributed to this disease-associated microglial program (Supplementary Fig. S33)^87–89^. A complementary NicheNet analysis prioritized candidate ligands across adjacent intervals of the reconstructed trajectory and linked them to receptors expressed in microglia (Supplementary Fig. S34). Prioritized ligands shifted from *Dcn, Sema3a*, and *Clu* in earlier intervals toward *C1qb, C3, Apoe*, and *Spp1* in later intervals.

Ligand–receptor scores also changed across the inferred trajectory (Fig. 6e). Several extracellular-matrix and adhesion-associated pairs, including *Fn1–Cd44, Col1a1–Cd44, Col6a1–Cd44*, and *Pecam1–Pecam1*, declined over time, indicating stage-dependent remodeling of the model-derived communication program.

We next used the trained model to simulate perturbations of microglial regulators. In silico *Trem2* activation and knockdown altered the generated cell-type distribution across hippocampal and cortical regions (Fig. 6f). *Trem2* activation increased the disease-associated microglia (DAM), phagolysosome and Aβ-clearance module scores, whereas knockdown decreased them (Fig. 6g), consistent with increased microglial activation and amyloid clearance after TREM2 activation^90^. In a separate *Spp1* simulation, *Spp1* knockdown reduced the *Spp1–Cd44*, DAM, and antigen-presentation modules and increased the myelination and endothelial–blood–brain barrier modules (Supplementary Fig. S35)^88,91^.

### CytoBridge resolves zebrafish morphogenesis and ablation responses

To test whether CytoBridge could capture interaction-guided embryonic morphogenesis, we applied it to Stereo-seq data comprising nearly 12,000 cells spanning zebrafish embryogenesis^72^. The reconstructed trajectory generated intermediate stages that preserved the global body plan and local cell-type organization between neighboring observations, whereas inferred growth maps revealed a stage-dependent redistribution of proliferative activity over development (Supplementary Figs. S36 and S37).

We further evaluated the learned interaction attention against external cell–cell interaction rankings and the zebrafish junctional adhesion molecule (JAM) program (Supplementary Fig. S38). Across all 361 directed cell-type pairs, attention ranks agreed with COMMOT after adjustment for sender and receiver abundance, spatial distance, and whether the two cells had the same cell type (Spearman ρ = 0.886). Among the 677 graph edges between somite cells at 18 hpf, those joining a *jam2a*-expressing cell to a *jam3b*-expressing cell made up 25.9% of the highest-attention quartile after interaction training but 1.8% of the lowest. With attention from the model before interaction training, the two quartiles differed little (8.8% and 7.1%). Complementary *jam2a*+/*jam3b*+ spatial neighbors were also enriched relative to random permutations of gene-detection labels within somite cells, and *jam3b* was preferentially detected in *myog*-positive cells. These findings suggest that higher attention among neighboring somite cells is associated with molecular features of the myocyte-fusion program, consistent with the reported roles of Jam2a–Jam3b interactions and Myogenin-dependent *jam3b* expression in this process^92,93^.

We next examined whether the learned dynamics could support in silico ablation analysis. Virtual ablation of yolk syncytial layer (YSL) support slowed developmental progression while largely preserving coarse body-plan organization, consistent with the established role of the YSL as a transient extraembryonic tissue that contributes to early patterning and coordinated morphogenetic progression in zebrafish development^94^. In contrast, ablation of the enveloping layer (EVL) support produced greater morphological instability, in line with the role of EVL in maintaining epithelial integrity and mechanical continuity during epiboly and surface morphogenesis^95^ (Supplementary Figs. S39 and S40). Within the yolk syncytial layer, reconstructed variable-gene trajectories showed distinct temporal patterns across observed and interpolated stages (Supplementary Fig. S41), and inverse-PCA reconstruction matched the mean expression profiles of observed YSL cells at the measured stages (Pearson r = 0.890–0.996; Supplementary Fig. S42).

## Discussion

Time-series spatial transcriptomics measures each tissue section once, so cell movement, population expansion and neighborhood-dependent changes in cell state are observed only indirectly. CytoBridge infers them jointly with an unbalanced mean-field Schrödinger bridge in the joint transcriptomic–spatial space, splitting the drift into an interaction component, estimated from spatial neighborhoods and ligand– receptor expression, and an intrinsic-context component. Applied to development, regeneration and neurodegeneration, the model generated unmeasured intermediate stages and showed where and when interactions contribute to the inferred dynamics, and its in silico perturbations point to candidate regulators for experimental tests.

Several limitations of the current framework and data should be noted. First, because the AGIST generator is itself a CytoBridge model and shares its structural assumptions, we used AGIST as a controlled recovery test and complemented it with an independent simulator, lineage-tracing and metabolic-labeling data, and held-out stages in five spatial datasets. Second, each tissue section samples only one plane of a three-dimensional tissue, and sparse temporal sampling or alignment error limits how well intermediate populations can be recovered; in the AD dataset, for example, the interval of more than a year between 5.7 and 17.9 months limits the temporal resolution of the inferred changes. Third, separating interaction from intrinsic-context contributions requires ligand–receptor and neighborhood features that vary enough among comparable cells to change the population in distinguishable ways (Supplementary Note S3.3). Finally, because these data are observational, the inferred interactions and the simulated perturbation responses are model predictions, and their causal roles need to be tested in perturbation experiments.

Addressing these limitations points to several directions for future work. Extending CytoBridge from two-dimensional sections to three-dimensional tissue representations and integrating sections across individuals and time, together with multimodal measurements such as chromatin accessibility, protein localization, imaging-derived morphology and biomechanical context^96,97^, should further help constrain latent dynamics^98^. Incorporating lineage recording or tracing^99,100^ or, when spliced and unspliced counts are available, RNA velocity estimates from methods such as veloAgent^24^ and STT^17^ into the learning objective could likewise constrain fate transitions and transcriptional directions. Time-resolved spatial measurements collected under both control and perturbed conditions will also be valuable for directly testing the predicted intervention responses. Computationally, larger spatiotemporal datasets will require more scalable training schemes, including reduced-cost solvers and simulation-free or flow-matching approximations. In the longer term, such advances may bring virtual tissues closer to reproducing how living systems develop, regenerate and degenerate.

## Methods

### Data and problem formulation

We consider time-series spatial transcriptomic data measured at ordered experimental times *t*_0_ < *t*_1_ < ⋯ < *t*_*K*_. At each time point *t*_*k*_, we observe *N*_*k*_ spots or cells indexed by *i* = 1, …, *N*_*k*_, *k* = 0, …, *K*, each with a spatial coordinate 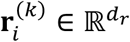 (e.g. physical tissue coordinates) and a transcriptomic state 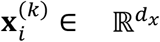 (e.g. the high-dimensional gene expression). The joint state space is

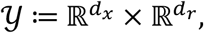

with generic element (**x, r**) ∈ *y*. The empirical distribution of cells at time *t*_*k*_ is

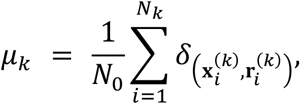

with total mass 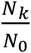 relative to the initial population. Dividing *µ*_*k*_ by its total mass gives the empirical probability distribution of cell states. Our goal is to infer a continuous-time evolution {*ρ*_*t*_}_*t*∈[0,1]_ and the underlying single-cell dynamics that interpolate the observed snapshots and jointly capture transcriptional drift, stochasticity, cell proliferation and death, and cell–cell interactions in the joint state space *y*. Unless otherwise specified, when considering a fixed snapshot at time *t*_*k*_, we drop the index *k* and write *N* and (**x, r**) in place of *N*_*k*_ and (**x**^(*k*)^, **r**^(*k*)^).

### Unbalanced mean-field Schrödinger bridge for spatiotemporal cell populations

A natural way to relate cell populations observed at different times is to treat them as snapshots of an underlying continuous population flow on the joint transcriptomic–spatial domain. Existing frameworks based on optimal transport^101–103^, unbalanced transport^104,105^, and Schrödinger bridge^106,107^ formulations provide a mathematical basis for coupling distributions across time, while accounting for cell movement, changes in population size, and stochasticity (Supplementary Note S1). However, these approaches primarily describe population evolution under prescribed transport or diffusion principles and do not explicitly model how local cell–cell interactions shape multicellular dynamics in space and time.

To model the coordinated evolution of interacting cells in space and expression, we adopt an unbalanced mean-field Schrödinger bridge (UMFSB) formulation^44^. In this framework, the drift experienced by a cell at state (**x, r**) depends not only on its own state but also on the current population distribution. We denote this interaction-induced component by ℐ[*ρ*_*t*_](**x, r**), which represents the net effect of neighboring cells on cells at state (**x, r**) under the population density *ρ*_*t*_. We learn this interaction term using a time-varying cell–cell interaction graph informed by spatial proximity and ligand–receptor connectivity. This yields a discrete approximation of ℐ[*ρ*_*t*_] that can be incorporated directly into the population dynamics model (Supplementary Note S1).

We describe the evolution of cells on *y* using a controlled mean-field diffusion model, where changes in cell state arise from intrinsic progression **b**_*t*_, population-dependent interactions ℐ[*ρ*_!_], stochastic fluctuations with scale *σ*(*t*), and local growth or death captured by *g*_*t*_. The resulting population density *ρ*_*t*_ then satisfies

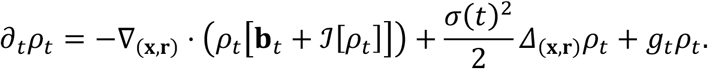

The formulation seeks an evolution (*ρ*, **b**, *g*) that is compatible with the observed snapshots while minimizing the intrinsic-control cost relative to the mass-conserving mean-field reference diffusion, together with a separate reaction cost for changes in population mass. This is expressed by minimizing an action functional

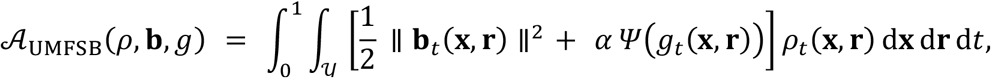

subject to the dynamics above and the marginal constraints 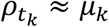at the observed time points. In this decomposition, **b**_*t*_ captures the intrinsic-context program in the joint transcriptomic–spatial space, *g*_*t*_ encodes local net expansion or loss of the population, and ℐ[*ρ*_*t*_] summarizes collective effects mediated by neighboring cells.

The quadratic action has different interpretations in optimal transport and Schrödinger bridge formulations. In Benamou–Brenier optimal transport^102^, the kinetic action follows from the quadratic transport cost, which is a modeling choice rather than a biophysical law. The Schrödinger bridge^106^ instead seeks a path distribution that satisfies the observed marginal constraints while minimizing relative entropy with respect to a reference stochastic process. Through its large-deviation interpretation, this describes the most likely collective evolution under the assumed reference dynamics. For a diffusion reference, Girsanov’s theorem expresses the drift-dependent relative entropy as a quadratic control cost.

In CytoBridge, the reference is a mass-conserving McKean–Vlasov diffusion driven by the interaction field ℐ[*ρ*_*t*_] and stochastic diffusion. Its dependence on the population density makes the reference law depend on the candidate population evolution. Conditional on the interaction operator, the intrinsic drift **b**_*t*_ controls departures from this interacting reference. In the mass-conserving case with constant isotropic diffusion, the quadratic drift action equals *σ*^2^ times the drift-dependent path-space relative entropy^108^. To accommodate population growth and loss, the unbalanced formulation additionally introduces *g*_*t*_*ρ*_*t*_ and a separate reaction penalty. With Ψ(*g*) = *g*^4^, this penalty integrates the Fisher–Rao cost^104^ of the pure reaction tangent 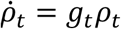. The objective thus combines a statistical interpretation of departures from interacting dynamics with a geometric cost for changes in population mass (Supplementary Notes S1.1 and S1.4.1 and Supplementary Table 1).

For model fitting, we assign model time k to observed stage k (k = 0, …, K), so that each experimental interval has unit length; the derivation above uses [0, 1] only as a generic time domain. The inferred drift and growth therefore describe rates per unit of model time and must be rescaled by the corresponding experimental interval when expressed in physical time units. Below, we parameterize **b**_*t*_, *g*_*t*_, and ℐ[*ρ*_*t*_] using neural networks defined on a data-driven cell–cell graph (including a graph neural network for the interaction term), and we describe how the resulting neural model is trained from spatiotemporal single-cell data. A fuller derivation of the joint transcriptomic–spatial formulation and its Fisher-information reformulation is provided in Supplementary Note S1.

### Data preprocessing and spatial alignment

We analyzed five spatial transcriptomic time series covering chicken heart development, mouse organogenesis (MOSTA), axolotl brain regeneration (ARTISTA), Alzheimer’s disease progression in TgCRND8 mice, and zebrafish embryogenesis, using one tissue section per sampled stage. Sampling times, cell or spot counts, and dataset-specific preprocessing details are provided in Supplementary Note S4.1.

Across all five datasets, experimental time labels were mapped to an ordered numerical timeline, expression profiles were library-size normalized and log-transformed, and highly variable genes were used to construct a low-dimensional representation for training.

Spatial alignment was then performed on preprocessed data to place snapshots from different time points into a coherent spatial reference frame while preserving local tissue geometry. Coordinates were centered and globally rescaled across time points, after which a transformation learned by a neural network was optimized with two complementary objectives: preserving within-timepoint spatial structure and minimizing cross-timepoint discrepancy in joint spatial-transcriptomic space via optimal transport.

The alignment objective was

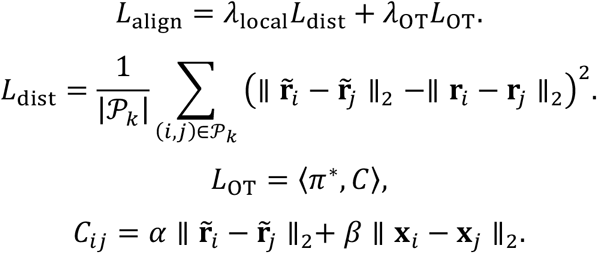

Here, *P*_*k*_ denotes the set of sampled cell pairs within time point *k*, and *π*^*∗*^ denotes the optimal transport coupling between adjacent time points. **r**_*i*_ denotes the original spatial coordinate of cell *i*, 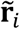 denotes its aligned coordinate after transformation, and **x**_*i*_ denotes its low-dimensional transcriptomic representation. In the definition of *C*_*ij*_, cell *i* is taken from time point *k* and cell *j* from time point *k* + 1. The resulting aligned coordinates were used for training. To assess sensitivity to the input spatial coordinates, we repeated alignment and model training after applying small translations and rotations to the chicken-heart sections. The resulting aligned geometries and velocity patterns were similar to those obtained without coordinate perturbation (Supplementary Figs. S12 and S13).

### Spatiotemporal interaction graph construction

To estimate interactions from observed neighborhoods and ligand–receptor features, we construct a dynamic graph *G* = (*V, E*), where nodes *V* represent cells and directed edges *E* represent interaction. At observed time points, candidate edges *e*_*ij*_ are established from a sender cell *i* to a receiver cell *j* if they are within a pre-defined spatial distance and cell *i* expresses a ligand for which cell *j* expresses the corresponding receptor, as determined by prior biological knowledge (e.g., COMMOT^59^ and CellChat^58^ database). As transcriptomic states (**x**) and spatial positions (**r**) are dynamic, these interaction patterns change over time. To capture this behavior, we introduce a neural network edge predictor, *f*_*θ*_, which takes the concatenated states of two cells, (**r**_*i*_, **x**_*i*_; **r**_*j*_, **x**_*j*_), as input and predicts whether the pair meets these criteria.

We train this predictor using data from observed time points, sampling ordered cell pairs from the observed data. Pairs that meet the spatial and ligand–receptor criteria using curated databases of known ligand– receptor pairs are treated as positive samples (*y*_*ij*_ = 1), and other sampled pairs are treated as negative samples (*y*_*ij*_ = 0). Let *Q*_edge_ denote the resulting set of *N* sampled ordered pairs (*i, j*). The parameters *θ* are optimized by minimizing the Binary Cross-Entropy (BCE) loss

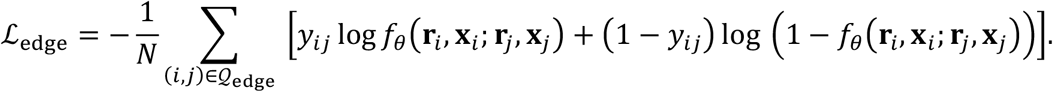

Once trained, the parameters of *f*_*θ*_ are held fixed, allowing the predictor to be applied to generated cell states at any unobserved time point to construct a continuous spatiotemporal interaction graph. Additional details of dynamic graph construction are provided in Supplementary Note S3.

### GNN-based interaction operator

We use an equivariant graph neural network (GNN) to parameterize the interaction operator and estimate the interaction component ℐ[*ρ*_*t*_]. This operator models the influence of neighboring cells on single-cell dynamics through two components: the interaction-driven gene regulation 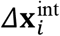 and spatial motility due to interactions 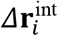. Given the discrete particle approximation of the density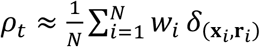, where *w*_*i*_ represents the weight of cell *i*, the GNN approximates the integral interaction field by aggregating messages over the dynamic graph constructed in the previous step.

Let 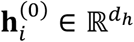 denote the initial embedding of the gene expression state **x**_*i*_, projected via a multi-layer perceptron (MLP). The network consists of an equivariant graph attention layer. For every edge (*j, i*) in the interaction graph, we compute the relative spatial position **r**_*ij*_ = **r**_*j*_ – **r**_*i*_, the Euclidean distance *d*_*ij*_ =∥ **r**_*ij*_ ∥, and the normalized direction 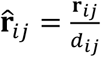. To capture high-frequency spatial dependencies, the scalar distance is expanded using a radial basis function (RBF) basis, **e**_*ij*_ = RBF(*d*_*ij*_), followed by a linear projection.

We employ a multi-head attention mechanism modified to respect the geometry of the joint space. For each head *k*, the attention coefficient 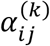 is computed based on the scalar features of the interacting cells and the edge attributes:

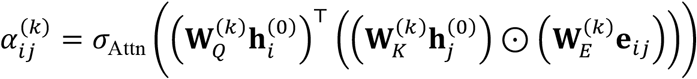

Here 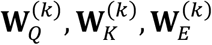 are learnable projection matrices for the *k*-th head, ⊙ denotes the element-wise product, and *σ*_Attn_ is the attention activation function. Thus, the attention weight for each cell pair depends on their gene-expression features and the distance between them.

The message propagation involves a coupled update of scalar and vector fields. For each head, we compute a scalar message 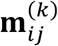 that combines the neighbor’s state features with the edge geometry and the center cell’s residual features:

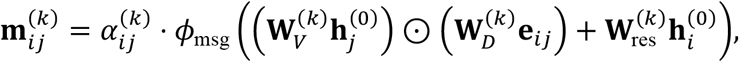

where 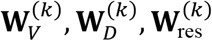 are projection matrices for the values, edge attributes, and residual connections, respectively, and *ϕ*_msg_ is a linear transformation.

Simultaneously, we update the spatial vector field. To maintain rotation equivariance, the vector update is driven by scalar coefficients derived from the cell-state message. We compute the vector message 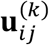 from the scalar message and relative spatial direction:

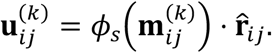

where *ϕ*_*s*_ is a projection layer that generates invariant scaling factors from the scalar message.

Finally, the network approximates the mean-field interaction integral by aggregating these messages. This aggregation is weighted by the cell density weights *w*_G_ from the unbalanced transport formulation:

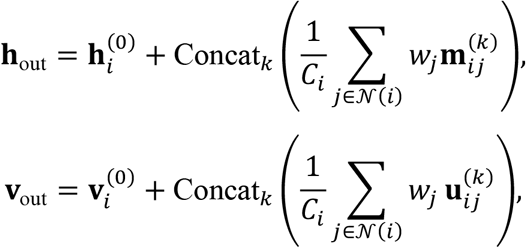

where *C*_*i*_ = ∑_*j*∈*N*(*i*)_ *w*_*j*_ is the normalization factor, and the vector features are initialized as 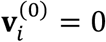. The final interaction drifts are obtained via readout functions: the transcriptomic interaction 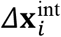 is projected from **h**_out_, and the spatial interaction force 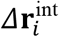 is derived by averaging **v**_out_ across feature channels. The corresponding equivariant GNN parameterization and the interpretation of attention weights are detailed in Supplementary Note S3.

### Particle-based approximation and numerical solver

The formulation introduces stochastic drift, diffusion, and growth terms at the population level. For numerical implementation, we adopt a Fisher-information regularized dynamic formulation that replaces the stochastic dynamics by an equivalent deterministic probability-flow representation. This leads to a first-order continuity equation driven by a velocity field **u**_*t*_ and a growth term, which can be efficiently approximated using interacting weighted particles (Supplementary Note S2).

The stochastic UMFSB problem can be reformulated in terms of a probability-flow drift 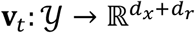 and a transport velocity **u**_*t*_ = **v**_*t*_ + ℐ[*ρ*_*t*_]. The Fisher-information action takes the form

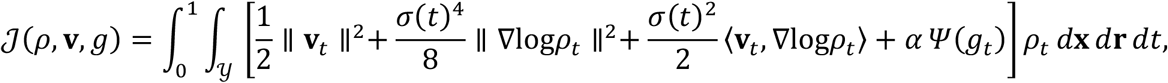

where 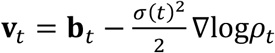 is the probability-flow drift. The corresponding density *ρ*_*t*_ satisfies the deterministic continuity equation

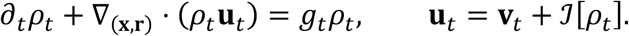

Thus probability flow refers to **v**_*t*_, whereas the continuity equation is driven by the transport velocity **u**_*t*_. This deterministic formulation is equivalent to the stochastic UMFSB problem under standard regularity assumptions (Supplementary Note S2).

To approximate the evolution of *ρ*_*t*_ in the joint transcriptomic–spatial space *y*, we represent *ρ*_*t*_ by a finite system of interacting weighted particles 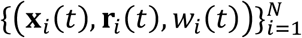. At any time *t*, the empirical measure

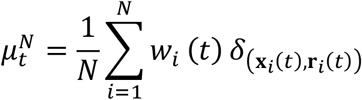

provides a discrete approximation of the population density *ρ*_*t*_. The particle states and weights evolve according to the coupled ordinary differential equation (ODE) system

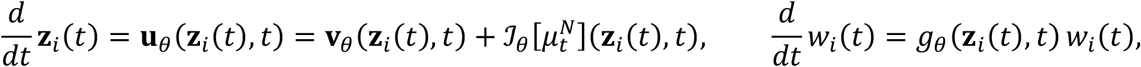

where **z**_*i*_(*t*) = (**x**_*i*_(*t*), **r**_*i*_(*t*)). We write *s*_θ_ for the estimated density score ∇log*ρ*_*t*_. A scalar potential *φ*_θ_ parameterizes the score drift 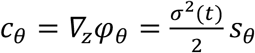, giving the intrinsic diffusive drift *b*_*θ*_ = *v* _*θ*_ + *c*_*θ*_. The transport velocity remains *u*_*θ*_ = *v*_*θ*_ + *I*_*θ*_. We call the drift of the simulated stochastic dynamics, the sum of the probability-flow, score and interaction contributions, the full velocity.

For each mini-batch of initial cells sampled from the earliest time point *t*_”_, we initialize particle states by the observed cell states and set *w*_*i*_(*t*_0_) = 1. We then integrate the interacting ODE system forward in time from *t*_0_ to *t*_*K*_ using a standard explicit ODE solver with a uniform time grid. At each solver step, we update the spatiotemporal cell–cell graph, evaluate **v**_*θ*_, *g*_*θ*_, **s**_*θ*_, and ℐ_*θ*_ for all particles, form 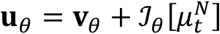, and advance both particle states and weights. The resulting particle ensemble at the observation times {*t*_*k*_} provides Monte Carlo estimates of 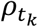, which are used to compute the training objectives described below.

The total loss is defined as

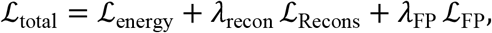

where ℒ_FP_ is an optional weak-form or physics-informed neural network penalty enforcing the transport continuity equation. Specifically, the variational energy loss ℒ_energy_ approximates the Fisher-information action in the deterministic formulation. Direct optimization of the inner product ⟨**v**_*θ*_, **s**_*θ*_ ⟩ can be numerically unstable. We therefore optimize the corresponding Cauchy–Schwarz upper bound along the simulated particle trajectories:

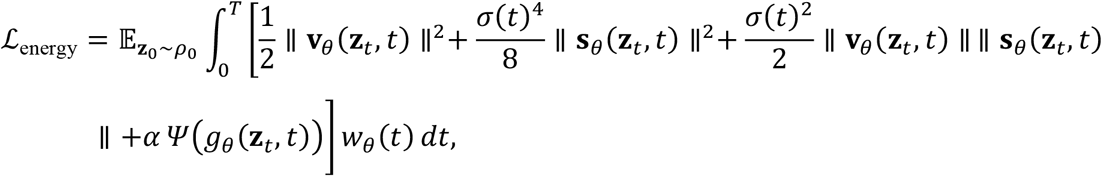

where **z**_*t*_ = (**x**_*t*_, **r**_*t*_) denotes the joint state of gene expression and spatial coordinates, and 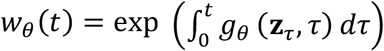 is the trajectory weight induced by growth (integrated up to terminal model time T = K). We used this upper bound as the energy regularization term during training.

The reconstruction loss ℒ_Recons_ aligns the simulated particle system with the observed data snapshots across time points *t*_*k*_. It is composed of an optimal transport loss ℒ_OT_ and a mass matching loss ℒ_mass_. The total reconstruction loss is given by:

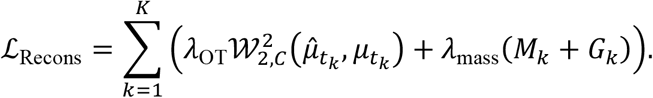

Here the optimal transport loss 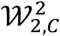 compares the simulated and observed distributions after normalizing each to unit total mass. It uses the joint-space ground cost matrix *C* that explicitly accounts for both spatial organization and transcriptomic similarity:

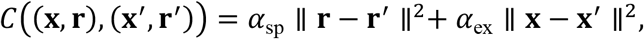

where *α*_sp_ and *α*_ex_ weight the contribution of spatial proximity and gene expression similarity, respectively. Simultaneously, ℒ_mass_ contains local and total-population terms. The local mass error *M*_*k*_ ensures that the weight distribution of simulated particles aligns with the local density of the real data:

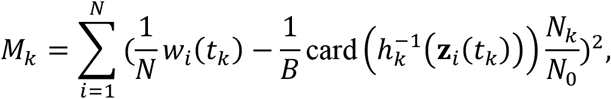

where *h*_*k*_ assigns the B observed points in the matched batch to their closest simulated particles. Here B is the observed batch size at this time point, which may differ from the number N of simulated particles. We also use the global mass term 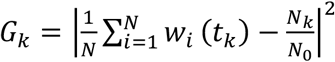, which constrains the total accumulated weight to match the observed relative expansion or contraction of the cell population size.

We train the probability-flow, growth, interaction, and score networks in stages. We first initialize **v**_*θ*_ and *g*_*θ*_ while freezing the interaction term ℐ_*θ*_ and score network **φ**_*θ*_. We then introduce ℐ_*θ*_ and refine the transport dynamics using the reconstruction loss. Before the final joint optimization, we initialize **φ**_*θ*_ using simulation-free score matching, and finally alternate updates of the dynamics and score networks (**v**_*θ*_, *g*_*θ*_, ℐ_*θ*_, **φ**_*θ*_) using their respective objectives. The score field is held fixed during the dynamics update, followed by a final score update (Supplementary Notes S2 and S4.4). The training objective was recorded at each epoch (six-stage curves for ARTISTA in Supplementary Fig. S43), and sensitivity to the expression-loss weight and the transport-to-mass loss ratio was evaluated in zebrafish: increasing the expression-loss weight improved expression reconstruction but worsened spatial reconstruction, whereas increasing the mass weight produced the opposite trade-off, so an expression-loss weight of 0.015 and a 1:1 transport-to-mass ratio were retained in the interaction stages (Supplementary Fig. S44).

The networks were implemented in PyTorch. We used neural ODE integration^109^ to simulate the trajectories and POT^110^ to calculate the optimal transport distance. To capture the distinct dynamic characteristics of physical migration and transcriptomic state transitions, we employ a decoupled multi-layer perceptron (MLP) architecture for the probability-flow network. This network consists of two parallel branches: a spatial branch outputting the 2-dimensional physical component and a gene branch outputting the high-dimensional transcriptomic component, both taking the concatenated time and state vector (**z**, *t*) as input. Specifically, we use 5 hidden layers with 256 units each and LeakyReLU activation functions. The growth rate *g*_*θ*_ is modeled by a 2-layer MLP mapping (**z**, *t*) to a scalar value with a hidden dimension of 256, while the scalar potential **φ**_*θ*_ uses a similar architecture with a hidden dimension of 400. The interaction operator is parameterized using a graph neural network with 8 attention heads and a hidden dimension of 256, together with a pre-trained link predictor to dynamically prune edges based on ligand– receptor interaction probability. Network architectures are described in Supplementary Note S4.3, with training schedules and parameter-selection rules in Supplementary Notes S4.4–S4.5 and Supplementary Table 2. We trained CytoBridge on each of the five datasets using an NVIDIA GeForce RTX 4090 D GPU, with one complete training run per dataset taking 11.6–33.3 min. Supplementary Table 3 lists the sampled stages, training times, and peak host and GPU memory.

We assessed the contribution of the ligand–receptor prior by retraining CytoBridge without the prior in all five spatial datasets, while retaining the interaction network and spatial-neighborhood cutoff. Separately, we disabled the interaction output during prediction without retraining and compared reconstruction errors from both analyses with those of the corresponding full models (Supplementary Note S4.8 and Supplementary Fig. S9). To evaluate reproducibility across training runs and model settings in zebrafish, five models trained with different random seeds were compared across all pairs, and models with altered neighborhood radius or loss weights were compared against the default model trained with the same seed. These comparisons evaluated reconstruction error, agreement between predicted dynamical components, and the rankings of directed cell-type interactions (Supplementary Note S4.7 and Supplementary Fig. S10).

### Trajectory simulation, interpolation, and downstream analysis

Next, we describe the procedures used to simulate cellular trajectories from the trained CytoBridge model, assign cell-type identities to simulated states, and quantify intercellular communication. Additional details of forward simulation, interpolation, label assignment, and communication analysis are provided in Supplementary Note S8.

### Forward simulation of developmental trajectories

Given the trained probability-flow network **v**_*θ*_, growth network *g*_*θ*_, scalar score potential **φ**_*θ*_, and interaction operator ℐ_*θ*_, we simulated developmental trajectories from an observed initial population using stochastic dynamics consistent with the UMFSB formulation. Here **s**_*θ*_(**x**, *t*) approximates the score ∇_**x**_log*ρ*_*t*_(**x**) (with x denoting the d-dimensional joint state in this subsection), and the intrinsic diffusive drift is recovered as

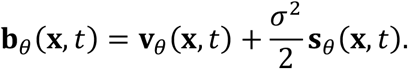

We considered two complementary simulation modes. In the weighted formulation, each particle *i* (*i* = 1, …, *N*) carries a joint state **x**_*i*_(*t*) ∈ ℝ^*d*)^ and a log-weight *l*_*i*_(*t*) = log*w*_*i*_(*t*). The coupled system evolves as

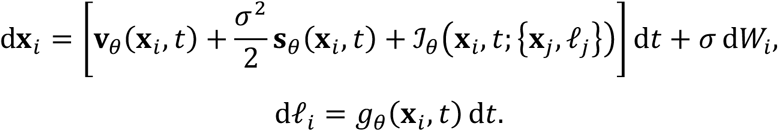

Here *W*_*i*_ is a standard *d*-dimensional Wiener process and *σ* ≥ 0 is the diffusion coefficient. When *σ* = 0, the state dynamics reduce to the deterministic transport ODE. The interaction term is computed by the attention-based GNN operating on random sub-groups of *m* particles: ℐ_*θ*_ (**x**_*i*_, *t*) = InteractionNet 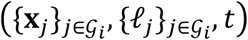, where *G*_*i*_ denotes the random sub-group of size *m* containing particle *i* (default *m* = 1024). Within each sub-group, the network selects edges using the spatial cutoff and trained link predictor, computes multi-head attention weights, and returns per-particle interaction forces. The log-weight *l*_*i*_ evolves under the growth network *g*_*θ*_, encoding changes in cellular abundance over time. In this formulation, the total particle count remains fixed at *N* throughout the simulation, and heterogeneous growth is represented through the weight distribution. The system is integrated numerically using the Euler–Maruyama scheme:

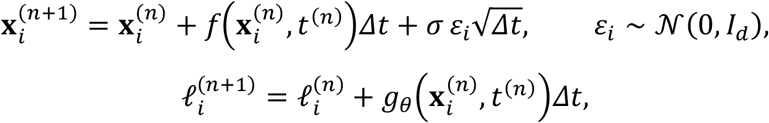

where

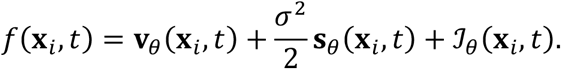

In the split formulation, the particle count varies dynamically over time to directly represent cellular proliferation and death. The state stochastic differential equation (SDE) is the same as in the weighted formulation:

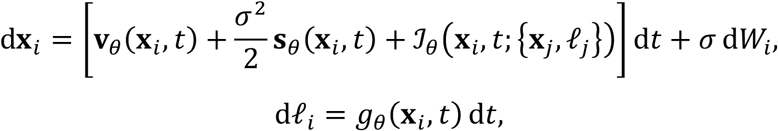

with interaction sub-group size *m*. The key distinction occurs at each output time point *t*_*k*_, where particles undergo stochastic branching and extinction based on the accumulated weight change. For each particle *i*, let

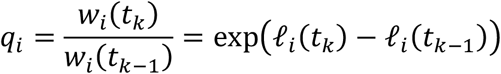

denote the weight ratio relative to the previous output time. **Proliferation** (*q*_*i*_ ≥ 1): particle *i* is replaced by *m*_*i*_ daughter particles, where

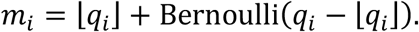

Each daughter inherits the parent state, with optional additive Gaussian noise:

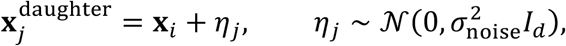

where *σ*_noise_ is a small perturbation scale. **Death** (*q*_*i*_ < 1): particle *i* survives with probability *q*_*i*_ and is removed otherwise. After branching, all surviving particles are re-assigned uniform log-weights

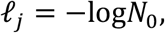

where *N*_0_ is the initial particle count. The updated population is then used as input for the next integration interval. This procedure ensures that the simulated particle count at each output time reflects the model-inferred growth and death dynamics, representing population changes through particle numbers. Intermediate states were generated by forward simulation of the learned split-SDE. We initialized daughter cells without additional perturbation (*σ*_noise_ = 0), while retaining the diffusion noise in the SDE. We also tested nonzero daughter-cell perturbations in zebrafish, which progressively altered cell-type composition and lineage-transition fractions as the perturbation scale increased (Supplementary Fig. S45).

### Cell-type label inference at simulated time points

To assign biological identities to cells generated by the split-SDE simulation at unobserved time points, we trained a multi-layer perceptron (MLP) classifier on the observed data and, for cell-type maps and proportions, applied spatial *k*-nearest neighbors (k-NN) smoothing. Lineage transitions were calculated from classifier cell-type predictions by following the same particles across time.

Let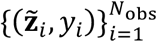denote the observed dataset, where

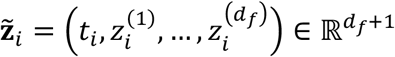

is the concatenation of the normalized time stamp and the *d*_*f*_ joint-state coordinates (52 dimensions), and

*y*_*i*_ ∈ {1, …, *C*} is the cell-type label. The classifier 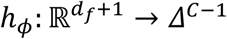 is a residual MLP trained by minimizing cross-entropy loss. We used a stratified 90:10 training–validation split within cell types and used the classifier from the epoch with the highest validation balanced accuracy (macro-averaged recall) for prediction. The classifier architecture and training schedule are described in Supplementary Note S8.7. To enforce local spatial coherence, the MLP predictions were refined by a *k*-nearest neighbors classifier in the 2D spatial coordinate space:

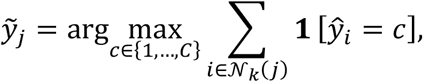

where 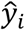 is the MLP-predicted label and *N*_*k*_(*j*) denotes the spatial *k*-nearest neighbors of cell *j*. We applied spatial voting with *k* = 10 to cell-type maps and proportions in zebrafish, MOSTA, and ARTISTA. For AD mouse and chicken heart, we retained the unsmoothed classifier labels (*k* = 1) because spatial voting substantially reduced held-out classification accuracy. We evaluated sensitivity to *k* = 1,5,10,20,50 using held-out classification accuracy and, in zebrafish, changes in cell-type composition and transition fractions (Supplementary Fig. S46 and Supplementary Note S8.7).

### Estimation of spatiotemporal cell–cell interactions

To summarize the learned interaction network, we extracted and aggregated attention weights from the GNN-based interaction operator. At each time point *t*, the interaction operator ℐ _*θ*_ constructs a spatiotemporal cell–cell graph with edge set *ℰ*_*t*_ Each edge (*i, j*) carries attention weights 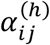 from *H* attention heads. We extracted the absolute multi-head mean:

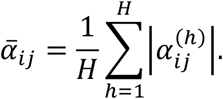

Given *C* cell types with *n*_*c*_ cells per type, we summarized attention between sender and receiver cell types in matrices at each time point. The raw summed matrix is

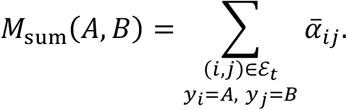

To account for differences in cell-type abundance, we divided the summed attention by the number of sender cells:

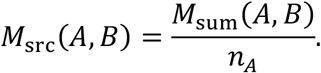

We also computed the row-normalized matrix

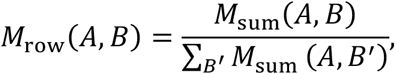

and the per-edge mean

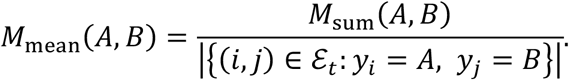

Directional asymmetry was measured as

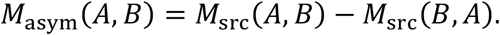

Cell-type interaction networks were embedded in three-dimensional space with horizontal axes for physical coordinates and the vertical axis for developmental time. Each time point was placed on a horizontal slice at height *z* = *t* ⋅ *Δz*. Edges were drawn as lines connecting cell-type centroids, with width proportional to *M*_src_(*A, B*). We added ribbons to the same 3D map to show cell-fate transitions from time *t*_*s*_ to *t*_*e*_. Each ribbon connects the centroid of source cell type *A* at *t*_*s*_ to that of target cell type *B* at *t*_*e*_, with its width proportional to the transition fraction *π*_*A*→*B*_.

### Time-resolved communication analysis and in silico perturbation

We followed ligand–receptor programs over time by aggregating directed interaction weights by sender and receiver cell type and combining these weights with ligand expression in sender cells and receptor expression in receiver cells. Multi-subunit complexes were represented by the expression of the least abundant required subunit. Across zebrafish, MOSTA, ARTISTA, and chicken heart, replacing this minimum with a geometric mean gave similar overall ligand–receptor rankings (Supplementary Fig. S47).

For the AD dataset, we used NicheNet to identify candidate ligands associated with changes in microglial gene expression along the reconstructed trajectory. For each pair of consecutive time points, the 50 microglial genes with the largest predicted increase in expression formed the target set for ligand-activity analysis. We then linked prioritized ligands to receptors expressed in microglia using the NicheNet prior and examined how these candidate signals changed over time (Supplementary Note S8.18).

In the same dataset, we used the trained CytoBridge model to predict the effects of *Trem2* and *Spp1* perturbations. Each perturbation was applied along the gene’s PCA loading direction without changing the initial spatial coordinates. Perturbed and unperturbed states were then propagated with the same split-SDE settings, assigned cell types with the trained classifier, and mapped back to gene space. We compared the resulting cell-type distributions and functional-module scores with those of the unperturbed TgCRND8 reference.

## Acknowledgments

This work was supported by the National Key R&D Program of China (No. 2021YFA1003301 to T.L.) and the National Natural Science Foundation of China (NSFC No. 12288101 to T.L. and P.Z.). We acknowledge the support from the High-performance Computing Platform of Peking University for computation.

## Author contributions

P.Z., T.L., Q.N., and Z.Z. conceived this research. Z.Z., Z.W., Y.S., T.L., and P.Z. designed the algorithm. Z.W. and Z.Z. implemented the algorithm. Z.Z., Z.W., Q.T., H.H., and J.H. analyzed the data. All authors interpreted the results. Z.Z., Z.W., Q.T., H.H., J.S., and X.Z. prepared the visualizations. Z.Z., Z.W., Q.T., and H.H. wrote the original draft. P.Z., T.L., and Q.N. reviewed and edited the manuscript. P.Z., T.L., and Q.N. supervised the work. All authors read and approved the manuscript.

## Data availability

All datasets used in this study are publicly available. The developing chicken heart dataset is available from the NCBI Gene Expression Omnibus (GEO) database under accession code GSE149457, and from https://github.com/madhavmantri/chicken_heart^68^. The mouse organogenesis dataset is available at https://db.cngb.org/stomics/mosta/download/^69^. The axolotl brain regeneration dataset is available at https://db.cngb.org/stomics/datasets/STDS0000056/data^70^. The TgCRND8 Alzheimer’s disease model mouse dataset is available at https://www.10xgenomics.com/datasets/xenium-in-situ-analysis-of-alzheimers-disease-mouse-model-brain-coronal-sections-from-one-hemisphere-over-a-time-course-1-standard^71^. The zebrafish development dataset is available at https://db.cngb.org/stomics/datasets/STDS0000057/data^72^. The LARRY hematopoiesis^66^ and scNT-seq cortical activation^67^ datasets are available from GEO under accession codes GSE140802 and GSE141851, respectively.

## References

1 Bunne, C. et al. How to build the virtual cell with artificial intelligence: priorities and opportunities. Cell 187, 7045–7063 (2024).

2 Qian, L., Dong, Z. & Guo, T. Grow AI virtual cells: three data pillars and closed-loop learning. Cell Res. 35, 319–321 (2025).

3 Roohani, Y. H. et al. Virtual Cell Challenge: Toward a Turing test for the virtual cell. Cell188, 3370–3374 (2025).

4 Tang, L. The virtual cell. Nat. Methods 22, 2493 (2025).

5 Kalafut, N. C. et al. Inferring virtual cell environments using multi-agent reinforcement learning. Preprint at 10.1101/2025.11.21.689815 (2025).

6 Wenckstern, J. et al. The Virtual Tissues foundation model resolves spatial proteomics across scales. Nature 657, 744–754 (2026).

7 Dimitrov, D., Schrod, S., Rohbeck, M. & Stegle, O. Interpretation, extrapolation and perturbation of single cells. Nat. Rev. Genet. 27, 349–370 (2026).

8 Zhang, Z. et al. Deciphering cell-fate trajectories using spatiotemporal single-cell transcriptomic data. npj Syst. Biol. Appl. 12, 2 (2026).

9 Liu, L. et al. Spatiotemporal omics for biology and medicine. Cell 187, 4488–4519 (2024).

10 Bunne, C., Schiebinger, G., Krause, A., Regev, A. & Cuturi, M. Optimal transport for single-cell and spatial omics. Nat. Rev. Methods Prim. 4, 58 (2024).

11 Zhang, Z., Sun, Y., Peng, Q., Li, T. & Zhou, P. Integrating dynamical systems modeling with spatiotemporal scRNA-seq data analysis. Entropy 27, 453 (2025).

12 Saelens, W., Cannoodt, R., Todorov, H. & Saeys, Y. A comparison of single-cell trajectory inference methods. Nat. Biotechnol. 37, 547–554 (2019).

13 Gulati, G. S. et al. Single-cell transcriptional diversity is a hallmark of developmental potential. Science 367, 405–411 (2020).

14 Kang, M. et al. Improved reconstruction of single-cell developmental potential with CytoTRACE 2. Nat. Methods 22, 2258–2263 (2025).

15 La Manno, G. et al. RNA velocity of single cells. Nature 560, 494–498 (2018).

16 Gu, Y. et al. Topological velocity inference from spatial transcriptomic data. Nat. Biotechnol. 44, 771–782 (2026).

17 Zhou, P., Bocci, F., Li, T. & Nie, Q. Spatial transition tensor of single cells. Nat. Methods 21, 1053–1062 (2024).

18 Abdelaal, T. et al. SIRV: spatial inference of RNA velocity at the single-cell resolution. NAR Genom. Bioinform. 6, qae100 (2024).

19 Li, S. et al. A relay velocity model infers cell-dependent RNA velocity.Nat. Biotechnol. 42, 99–108 (2024).

20 Gao, M., Qiao, C. & Huang, Y. UniTVelo: temporally unified RNA velocity reinforces single-cell trajectory inference. Nat. Commun. 13, 6586 (2022).

21 Qiu, X. et al. Mapping transcriptomic vector fields of single cells. Cell 185, 690–711.e45 (2022).

22 Bergen, V., Lange, M., Peidli, S., Wolf, F. A. & Theis, F. J. Generalizing RNA velocity to transient cell states through dynamical modeling. Nat. Biotechnol. 38, 1408–1414 (2020).

23 Li, C., Virgilio, M. C., Collins, K. L. & Welch, J. D. Multi-omic single-cell velocity models epigenome–transcriptome interactions and improves cell fate prediction. Nat. Biotechnol. 41, 387–398 (2023).

24 Raghavan, V., Yoon, B., Fonseca, G. J., Li, Y. & Ding, J. Dissecting and steering cell dynamics using spatially-informed RNA velocity with veloAgent. Mol. Syst. Biol. 22, 1180–1200 (2026).

25 Ding, J., Sharon, N. & Bar-Joseph, Z. Temporal modelling using single-cell transcriptomics. Nat. Rev. Genet. 23, 355–368 (2022).

26 Liu, Y. et al. TemporalVAE: atlas-assisted temporal mapping of time-series single-cell transcriptomes during embryogenesis. Nat. Cell Biol. 27, 1982–1992 (2025).

27 Moon, K. R. et al. Visualizing structure and transitions in high-dimensional biological data. Nat. Biotechnol. 37, 1482–1492 (2019).

28 Klein, D. et al. Mapping cells through time and space with moscot. Nature 638, 1065–1075 (2025).

29 Schiebinger, G. et al. Optimal-Transport Analysis of Single-Cell Gene Expression Identifies Developmental Trajectories in Reprogramming. Cell 176, 928–943.e22 (2019).

30 Tong, A., Huang, J., Wolf, G., Van Dijk, D. & Krishnaswamy, S. TrajectoryNet: A Dynamic Optimal Transport Network for Modeling Cellular Dynamics. In Proceedings of the 37th International Conference on Machine Learning Vol. 119 9526–9536 (PMLR, 2020).

31 Lavenant, H., Zhang, S.Kim, Y.-H. & Schiebinger, G. Toward a mathematical theory of trajectory inference. Ann. Appl. Probab. 34, 428–500 (2024).

32 Zhao, W., Larschan, E., Sandstede, B. & Singh, R. Optimal transport reveals dynamicgene regulatory networks via gene velocity estimation. PLoS Comput. Biol. 21, e1012476 (2025).

33 Baker, C. et al. SCOT+: a comprehensive software suite for single-cell alignment using optimal transport. Bioinform. Adv. 6, vbaf314 (2026).

34 Lipman, Y., Chen, R. T. Q., Ben-Hamu, H., Nickel, M. & Le, M. Flow Matching for Generative Modeling. In The Eleventh International Conference on Learning Representations (2023).

35 Wang, D. et al. Joint Velocity-Growth Flow Matching for Single-Cell Dynamics Modeling. In Advances in Neural Information Processing Systems Vol. 38 178180–178216 (Curran Associates, Inc., 2025).

36 Morehead, A. et al. Flow matching for generative modelling in bioinformatics and computational biology. Nat. Mach. Intell. 8, 517–534 (2026).

37 Peng, Q. et al. WFR-FM: Simulation-Free Dynamic Unbalanced Optimal Transport. In The Fourteenth International Conference on Learning Representations (2026).

38 Wang, X., Wang, R., Peng, Q., Zhou, P. & Li, T. WFR-MFM: One-Step Inference for Dynamic Unbalanced Optimal Transport. Preprint at https://arxiv.org/abs/2601.20606 (2026).

39 Sha, Y., Qiu, Y., Zhou, P. & Nie, Q. Reconstructing growth and dynamic trajectories from single-cell transcriptomics data. Nat. Mach. Intell. 6, 25–39 (2024).

40 Zhang, Z., Li, T. & Zhou, P. Learning Stochastic Dynamics from Snapshots through Regularized Unbalanced Optimal Transport. In The Thirteenth International Conference on Learning Representations (2025).

41 Pariset, M., Hsieh, Y.-P., Bunne, C., Krause, A. & De Bortoli, V. Unbalanced Diffusion Schrödinger Bridge. In ICML Workshop on New Frontiers in Learning, Control, and Dynamical Systems (2023).

42 Jiang, Q. & Wan, L. A physics-informed neural SDE network for learning cellular dynamics from time-series scRNA-seq data. Bioinformatics 40, ii120–ii127 (2024).

43 Tong, A. Y. et al. Simulation-Free Schrödinger Bridges via Score and Flow Matching. In Proceedings of the 27th International Conference on Artificial Intelligence and Statistics Vol. 238 1279–1287 (PMLR, 2024).

44 Zhang, Z., Wang, Z., Sun, Y., Li, T. & Zhou, P. Modeling Cell Dynamics and Interactions with Unbalanced Mean Field Schrödinger Bridge. In Advances in Neural Information Processing Systems Vol. 38 154733–154773 (Curran Associates, Inc., 2025).

45 Jiang, Q., Li, L., Zhang, L. & Wan, L. Learning collective multicellular dynamics with an interacting mean field neural SDE model. PLoS Comput. Biol. 22, e1013916 (2026).

46 Alatkar, S. A. & Wang, D. ARTEMIS integrates autoencoders and Schrödinger Bridges to predict continuous dynamics of gene expression, cell population, and perturbation from time-series single-cell data. Bioinformatics 41, i189–i197 (2025).

47 Vinyard, M. E. et al. Learning cell dynamics with neural differential equations. Nat. Mach. Intell. 7, 1969–1984 (2025).

48 Zheng, Y. et al. A deep generative model for deciphering cellular dynamics and in silico drug discovery in complex diseases. Nat. Biomed. Eng. 9, 2155–2180 (2025).

49 Sun, X. et al. MIOFlow 2.0: A unified framework for inferring cellular stochastic dynamics from single cell and spatial transcriptomics data. Preprint at https://arxiv.org/abs/2603.22564 (2026).

50 Qiu, X. et al. Spatiotemporal modeling of molecular holograms. Cell 187, 7351–7373.e61 (2024).

51 Halmos, P. et al. DeST-OT: Alignment of spatiotemporal transcriptomics data. Cell Syst.,101160 (2025).

52 Halmos, P., Gold, J., Liu, X. & Raphael, B. J. Learning latent trajectories in developmental time series with Hidden-Markov optimal transport. In Research in Computational Molecular Biology (ed. Sankararaman, S.) 367–370 (Springer, 2025).

53 Huizing, G.-J. et al. STORIES: learning cell fate landscapes from spatial transcriptomics using optimal transport. Nat. Methods 23, 522–531 (2026).

54 Peng, Q., Zhou, P. & Li, T. stVCR: spatiotemporal dynamics of single cells. Nat. Methods 23, 542–553 (2026).

55 Lázár, E. & Lundeberg, J. Spatial architecture of development and disease. Nat. Rev. Genet. 27, 118–136 (2026).

56 Longo, S. K., Guo, M. G., Ji, A. L. & Khavari, P. A. Integrating single-cell and spatial transcriptomics to elucidate intercellular tissue dynamics. Nat. Rev. Genet. 22, 627–644 (2021).

57 Browaeys, R., Saelens, W. & Saeys, Y. NicheNet: modeling intercellular communication by linking ligands to target genes. Nat. Methods 17, 159–162 (2020).

58 Jin, S. et al. Inference and analysis of cell-cell communication using CellChat. Nat. Commun. 12, 1088 (2021).

59 Cang, Z. et al. Screening cell–cell communication in spatial transcriptomics via collective optimal transport. Nat. Methods 20, 218–228 (2023).

60 Zohora, F. T. et al. CellNEST reveals cell–cell relay networks using attention mechanisms on spatial transcriptomics. Nat. Methods 22, 1505–1519 (2025).

61 Cheng, X. et al. Cell-Cell Communication Inference and Analysis: Biological Mechanisms, Computational Approaches, and Future Opportunities. CSIAM Trans. Life Sci. 2, 214–258 (2026).

62 Almet, A. A., Tsai, Y. C., Watanabe, M. & Nie, Q. Inferring pattern-driving intercellular flows from single-cell and spatial transcriptomics. Nat. Methods 21, 1806–1817 (2024).

63 Tejada-Lapuerta, A. et al. Causal machine learning for single-cell genomics. Nat. Genet. 57, 797–808 (2025).

64 Yan, L., Zhang, D. & Sun, X. Decoding cell state transitions driven by dynamic cell–cell communication in spatial transcriptomics. Nat. Comput. Sci. 6, 193–207 (2026).

65 Liu, J. et al. CytoSignal detects locations and dynamics of ligand–receptor signaling at cellular resolution from spatial transcriptomic data. Nat. Genet. 58, 1396–1408 (2026).

66 Weinreb, C., Rodriguez-Fraticelli, A., Camargo, F. D. & Klein, A. M. Lineage tracing on transcriptional landscapes links state to fate during differentiation. Science 367, eaaw3381 (2020).

67 Qiu, Q. et al. Massively parallel and time-resolved RNA sequencing in single cells with scNT-seq. Nat. Methods 17, 991–1001 (2020).

68 Mantri, M. et al. Spatiotemporal single-cell RNA sequencing of developing chicken hearts identifies interplay between cellular differentiation and morphogenesis. Nat. Commun. 12, 1771 (2021).

69 Chen, A. et al. Spatiotemporal transcriptomic atlas of mouse organogenesis using DNA nanoball-patterned arrays. Cell 185, 1777–1792.e21 (2022).

70 Wei, X. et al. Single-cell Stereo-seq reveals induced progenitor cells involved in axolotl brain regeneration. Science 377, eabp9444 (2022).

71 10x Genomics. Xenium In Situ Analysis of Alzheimer’s Disease Mouse Model Brain Coronal Sections from One Hemisphere Over a Time Course, https://www.10xgenomics.com/datasets/xenium-in-situ-analysis-of-alzheimers-disease-mouse-model-brain-coronal-sections-from-one-hemisphere-over-a-time-course-1-standard (2023).

72 Liu, C. et al. Spatiotemporal mapping of gene expression landscapes and developmentaltrajectories during zebrafish embryogenesis. Dev. Cell 57, 1284–1298.e5 (2022).

73 Zeira, R., Land, M., Strzalkowski, A. & Raphael, B. J. Alignment and integration of spatial transcriptomics data. Nat. Methods 19, 567–575 (2022).

74 Liu, X., Zeira, R. & Raphael, B. J. Partial alignment of multislice spatially resolved transcriptomics data. Genome Res. 33, 1124–1132 (2023).

75 Shen, X. et al. Inferring cell trajectories of spatial transcriptomics via optimal transport analysis. Cell Syst. 16, 101194 (2025).

76 Huguet, G. et al. Manifold interpolating optimal-transport flows for trajectory inference. Adv. Neural Inf. Process. Syst. 35, 29705–29718 (2022).

77 Raghavan, V., Zheng, Y., Li, Y. & Ding, J. Harnessing agent-based frameworks in CellAgentChat to unravel cell–cell interactions from single-cell and spatial transcriptomics. Genome Res. 35, 1646–1663 (2025).

78 Dettman, R. W., Denetclaw Jr, W., Ordahl, C. P. & Bristow, J. Common epicardial origin of coronary vascular smooth muscle, perivascular fibroblasts, and intermyocardial fibroblasts in the avian heart. Dev. Biol. 193, 169–181 (1998).

79 Lotto, J. et al. Cell diversity and plasticity during atrioventricular heart valve EMTs. Nat. Commun. 14, 5567 (2023).

80 Tian, Y. & Morrisey, E. E. Importance of myocyte-nonmyocyte interactions in cardiac development and disease. Circ. Res. 110, 1023–1034 (2012).

81 Wittig, J. G. & Münsterberg, A. The chicken as a model organism to study heart development. Cold Spring Harb. Perspect. Biol. 12, a037218 (2020).

82 Grove, E. A., Tole, S., Limon, J., Yip, L.-w. & Ragsdale, C. W. The hem of the embryonic cerebral cortex is defined by the expression of multiple Wnt genes and is compromised in Gli3-deficient mice. Development 125, 2315–2325 (1998).

83 Kelly, N. H., Huynh, N. P. T. & Guilak, F. Single cell RNA-sequencing reveals cellular heterogeneity and trajectories of lineage specification during murine embryonic limb development. Matrix Biol. 89, 1–10 (2020).

84 Di Bella, D. J. et al. Molecular logic of cellular diversification in the mouse cerebral cortex. Nature 595, 554–559 (2021).

85 Lust, K. et al. Single-cell analyses of axolotl telencephalon organization, neurogenesis, and regeneration. Science 377, eabp9262 (2022).

86 Amamoto, R. et al. Adult axolotls can regenerate original neuronal diversity in response to brain injury. eLife 5, e13998 (2016).

87 Deczkowska, A. et al. Disease-associated microglia: a universal immune sensor of neurodegeneration. Cell 173, 1073–1081 (2018).

88 Keren-Shaul, H. et al. A unique microglia type associated with restricting development of Alzheimer’s disease. Cell 169, 1276–1290.e17 (2017).

89 Heneka, M. T. et al. Neuroinflammation in Alzheimer’s disease. Lancet Neurol. 14, 388–405 (2015).

90 Price, B. R. et al. Therapeutic Trem2 activation ameliorates amyloid-beta deposition and improves cognition in the 5XFAD model of amyloid deposition. J. Neuroinflammation 17, 238 (2020).

91 De Schepper, S. et al. Perivascular cells induce microglial phagocytic states and synaptic engulfment via SPP1 in mouse models of Alzheimer’s disease. Nat. Neurosci. 26, 406–415 (2023).

92 Ganassi, M. et al. Myogenin promotes myocyte fusion to balance fibre number and size.Nat. Commun. 9, 4232 (2018).

93 Powell, G. T. & Wright, G. J. Jamb and Jamc are essential for vertebrate myocyte fusion. PLoS Biol. 9, e1001216 (2011).

94 Carvalho, L. & Heisenberg, C.-P. The yolk syncytial layer in early zebrafish development. Trends Cell Biol. 20, 586–592 (2010).

95 Bruce, A. E. E. Zebrafish epiboly: Spreading thin over the yolk. Dev. Dyn. 245, 244–258 (2016).

96 Hallou, A., He, R., Simons, B. D. & Dumitrascu, B. A computational pipeline for spatial mechano-transcriptomics. Nat. Methods 22, 737–750 (2025).

97 Ichbiah, S., Delbary, F., McDougall, A., Dumollard, R. & Turlier, H. Embryo mechanics cartography: inference of 3D force atlases from fluorescence microscopy. Nat. Methods 20, 1989–1999 (2023).

98 Bahrami, M., Richter, T., Schmacke, N. A., Egea Lavandera, A. & Theis, F. J. From modality-specific to compositional foundation models for cell biology. Cell Syst. 17, 101534 (2026).

99 Sashittal, P. et al. Inferring cell differentiation maps from lineage tracing data. Nat. Methods 23, 532–541 (2026).

100 Wagner, D. E. & Klein, A. M. Lineage tracing meets single-cell omics: opportunities and challenges. Nat. Rev. Genet. 21, 410–427 (2020).

101 Monge, G. Mémoire sur la théorie des déblais et des remblais. Mémoires de l’Académie royale des sciences, 666–704 (1781).

102 Benamou, J. D. & Brenier, Y. A computational fluid mechanics solution to the Monge-Kantorovich mass transfer problem. Numer. Math. 84, 375–393 (2000).

103 Kantorovich, L. V. On the translocation of masses. Dokl. Akad. Nauk SSSR 37, 227–229 (1942).

104 Chizat, L., Peyré, G., Schmitzer, B. & Vialard, F. X. An Interpolating Distance Between Optimal Transport and Fisher–Rao Metrics. Found. Comput. Math. 18, 1–44 (2018).

105 Chizat, L., Peyré, G., Schmitzer, B. & Vialard, F. X. Unbalanced optimal transport: Dynamic and Kantorovich formulations. J. Funct. Anal. 274, 3090–3123 (2018).

106 Léonard, C. A survey of the Schrödinger problem and some of its connections with optimal transport. Discret. Contin. Dyn. Syst. Ser. A 34, 1533–1574 (2014).

107 Schrödinger, E. Sur la théorie relativiste de l’électron et l’interprétation de la mécanique quantique. Ann. Inst. Henri Poincaré 2, 269–310 (1932).

108 Backhoff, J., Conforti, G., Gentil, I. & Léonard, C. The mean field Schrödinger problem: ergodic behavior, entropy estimates and functional inequalities. Probab. Theory Relat. Fields 178, 475–530 (2020).

109 Chen, R. T. Q., Rubanova, Y., Bettencourt, J. & Duvenaud, D. K. Neural Ordinary Differential Equations. In Advances in Neural Information Processing Systems Vol. 31 (Curran Associates, Inc., 2018).

110 Flamary, R. et al. POT: Python Optimal Transport. J. Mach. Learn. Res. 22, 1–8 (2021).

