## Supplementary Information for "Virtual dynamics of spatiotemporal single-cell interactions"

**Supplementary Table 1. Comparison of related transport formulations 47**

**Supplementary Table 2. Graph-construction and training settings used for the five applications 48**

**Supplementary Table 3. Computational cost of the full CytoBridge models** 49

**Supplementary Figure S1. CytoBridge analysis workflow** 50

**Supplementary Figure S2. Simulation benchmark of growth, attention and distributional recovery** 51

**Supplementary Figure S3. Velocity agreement across time and state space in the AGIST benchmark** 52

**Supplementary Figure S4. Synthetic spatiotemporal benchmark with finite-range attractive cell–cell interactions** 53

**Supplementary Figure S5. Interaction-informed dynamics in the Weinreb lineage-tracing dataset** 55

**Supplementary Figure S6. Interaction-informed dynamics in the scNT-seq cortical activation dataset** 57

**Supplementary Figure S7. Five-dataset leave-one-time-point-out benchmark across methods** 59

**Supplementary Figure S8. Five-dataset leave-one-time-point-out benchmark** 61

**Supplementary Figure S9. Ligand–receptor-prior and interaction ablations across five spatial datasets** 63

**Supplementary Figure S10. Stability of the interaction and intrinsic-context decomposition in zebrafish** 65

**Supplementary Figure S11. Cross-method comparison of CytoBridge interaction summaries** 67

**Supplementary Figure S12. Sensitivity of chicken-heart alignment and downstream results to small coordinate perturbations** 69

**Supplementary Figure S13. Chicken-heart alignment after small coordinate perturbations** 71

**Supplementary Figure S14. Growth estimates during chicken-heart development** 73

**Supplementary Figure S15. Velocity components in the developing chicken heart** 74

**Supplementary Figure S16. Generated spatial maps across the MOSTA trajectory** 76

**Supplementary Figure S17. Growth-rate maps across the MOSTA trajectory** 77

**Supplementary Figure S18. Cell counts and composition across the MOSTA trajectory** 78

**Supplementary Figure S19. Cell-state transitions across the MOSTA trajectory** 79

**Supplementary Figure S20. Temporal gene programs in the MOSTA dataset** **80**

**Supplementary Figure S21. Functional enrichment of temporal gene programs in the MOSTA dataset** **81**

**Supplementary Figure S22. Developmental waves in the MOSTA dataset** **82**

**Supplementary Figure S23. Ligand–receptor temporal patterns in the MOSTA dataset** **83**

**Supplementary Figure S24. Observed and generated stages across the ARTISTA trajectory** **84**

**Supplementary Figure S25. Particle flows and composition across the ARTISTA trajectory** **85**

**Supplementary Figure S26. Localized interaction domains in the ARTISTA 5-DPI field** **86**

**Supplementary Figure S27. Temporal gene programs and GO analysis in the ARTISTA dataset** **88**

**Supplementary Figure S28. Ligand–receptor communication pattern prototypes in the ARTISTA dataset** **89**

**Supplementary Figure S29. Representative ligand–receptor communication profiles in the ARTISTA dataset** **90**

**Supplementary Figure S30. Growth-rate maps across the ARTISTA trajectory** **91**

**Supplementary Figure S31. Reconstructed AD progression in TgCRND8 mice** **92**

**Supplementary Figure S32. Pattern 1 functional enrichment in the AD dataset** **93**

**Supplementary Figure S33. Pattern 4 functional enrichment in the AD dataset** **94**

**Supplementary Figure S34. Temporal ligand activity and ligand–receptor networks in the AD dataset** **95**

**Supplementary Figure S35. Functional-module changes after in silico Spp1 perturbation** **96**

**Supplementary Figure S36. Observed and generated zebrafish development** **97**

**Supplementary Figure S37. Growth-rate maps across observed zebrafish stages** **99**

**Supplementary Figure S38. External and spatial comparison of learned interaction attention in zebrafish** **100**

**Supplementary Figure S39. Development after virtual YSL and EVL removal** **102**

**Supplementary Figure S40. Spatial divergence after virtual YSL and EVL removal 104**

**Supplementary Figure S41. YSL-lineage gene dynamics 105**

**Supplementary Figure S42. Inverse-PCA reconstruction of observed YSL expression 107**

**Supplementary Figure S43. Representative six-stage training histories 108**

**Supplementary Figure S44. Sensitivity to training-loss weights in zebrafish 110**

**Supplementary Figure S45. Daughter-cell perturbation sensitivity in zebrafish 111**

**Supplementary Figure S46. Effect of spatial smoothing on cell-type assignments 112**

**Supplementary Figure S47. Sensitivity of ligand–receptor rankings to multi-subunit aggregation 113**

### Supplementary Note S1. Joint transcriptomic–spatial formulation

This note collects the formal definitions and derivations that are only summarized in the main-text Methods. Throughout the SI, we write the joint transcriptomic–spatial state as $\mathbf{z} = (\mathbf{x}, \mathbf{r}) \in \mathcal{Y}$, where $\mathbf{x} \in \mathbb{R}^{d_x}$ is the transcriptomic embedding and $\mathbf{r} \in \mathbb{R}^{d_r}$ is the spatial coordinate. The observed snapshot at experimental time $t_k$ is $\mu_k$, its continuous-time population density is $\rho_t$, $\mathbf{b}_t$ denotes the intrinsic drift in the diffusive formulation, $\mathbf{v}_t$ denotes the probability-flow drift used by the particle solver and in the main text, $\mathbf{u}_t = \mathbf{v}_t + \mathcal{I}[\rho_t]$ is the transport velocity, $g_t$ is the net growth field, and $\sigma(t)$ is the diffusion scale. Supplementary Fig. S1 summarizes the CytoBridge workflow.

#### S1.1 Overview of optimal transport and Schrödinger bridge formulations

CytoBridge uses optimal transport (OT) and Schrödinger bridge (SB) formulations to connect discrete spatial-transcriptomic snapshots through a continuous population trajectory. Supplementary Note S2 gives the particle approximation and numerical solver. Related approaches to dynamical modeling of spatiotemporal single-cell data are reviewed in [1].

We model the observed snapshots $\{\mu_k\}_{k=0}^{K}$ as time-indexed marginals of an underlying continuous population flow on the joint transcriptomic–spatial state space $\mathcal{Y}$. Dynamical optimal transport (OT) provides a deterministic view of this problem by seeking a velocity field that transports one population distribution into another with minimal kinetic cost [2–4]. In the mass-preserving setting, the density $\rho_t$ evolves under a continuity equation driven by a transport field, and the associated action measures the amount of motion required to match endpoint or intermediate distributions.

$$\partial_t \rho_t(\mathbf{z}) + \nabla\cdot_{\mathbf{z}}\big(\rho_t(\mathbf{z})\, \mathbf{v}_t(\mathbf{z})\big) = 0. \tag{S1}$$

Among all pairs $(\rho, \mathbf{v})$ satisfying (S1) and matching the observed marginals, dynamical OT selects those with minimal kinetic action

$$\mathcal{A}_{\mathrm{OT}}(\rho, \mathbf{v}) := \int_0^1 \int_{\mathcal{Y}} \tfrac{1}{2}\, \|\mathbf{v}_t(\mathbf{z})\|^2\, \rho_t(\mathbf{z})\, \mathrm{d}\mathbf{z}\, \mathrm{d}t. \tag{S2}$$

For the Euclidean joint state space used here, the Benamou–Brenier dynamic formulation is equivalent to the static Kantorovich problem with quadratic ground cost $c(\mathbf{z}_0, \mathbf{z}_1) = \frac{1}{2}\|\mathbf{z}_1 - \mathbf{z}_0\|^2$. For prescribed endpoint densities $\rho_0$ and $\rho_1$, the minimum of Eq. (S2) subject to Eq. (S1) is the corresponding static quadratic transport cost, $\frac{1}{2}W_2^2(\rho_0, \rho_1)$. The kinetic action follows from the quadratic transport cost, a modeling choice rather than a biophysical action.

For developmental and tissue data, however, strict mass conservation is usually inappropriate because cell numbers change through proliferation and death, and each stage is measured on a different section with a different number of captured cells. Unbalanced transport addresses this issue by introducing a scalar growth term $g_t$ that allows local mass creation and depletion while penalizing large deviations from conservation [5, 6]. This extension keeps the transport picture intact but permits the model to represent biologically meaningful changes in population size.

$$\partial_t \rho_t(\mathbf{z}) + \nabla\cdot_{\mathbf{z}}\big(\rho_t(\mathbf{z})\, \mathbf{v}_t(\mathbf{z})\big) = g_t(\mathbf{z})\, \rho_t(\mathbf{z}).$$

The corresponding unbalanced action is

$$\mathcal{A}_{\mathrm{UOT}}(\rho, \mathbf{v}, g) := \int_0^1 \int_{\mathcal{Y}} \Big[\tfrac{1}{2}\, \|\mathbf{v}_t(\mathbf{z})\|^2 + \alpha\, \Psi(g_t(\mathbf{z}))\Big] \rho_t(\mathbf{z})\, \mathrm{d}\mathbf{z}\, \mathrm{d}t.$$

Schrödinger bridge (SB) theory adds stochasticity by starting from a reference diffusion process and asking for the most likely stochastic evolution that matches the observed marginals [7, 8].

$$\mathrm{d}\mathbf{z}_t = \mathbf{b}_t(\mathbf{z}_t)\,\mathrm{d}t + \Sigma\,\mathrm{d}\mathbf{W}_t,$$

where $\mathbf{W}_t$ is a standard Brownian motion and $D := \Sigma\Sigma^{\mathsf{T}}$. For isotropic diffusion, $D(t) = \sigma^2(t)I$, so the diffusion matrix is diagonal with common diagonal entry $\sigma^2(t)$. The reported applications used constant $\sigma$ unless stated otherwise. The corresponding density satisfies the Fokker–Planck equation

$$\partial_t \rho_t(\mathbf{z}) = -\nabla\cdot_{\mathbf{z}}\big(\rho_t(\mathbf{z})\,\mathbf{b}_t(\mathbf{z})\big) + \tfrac{1}{2}\nabla\cdot_{\mathbf{z}}\big(D\,\nabla_{\mathbf{z}}\rho_t(\mathbf{z})\big). \tag{S3}$$

Under this formulation, the Schrödinger bridge objective can be written as the minimum kinetic control cost

$$\mathcal{A}_{\mathrm{SB}}(\rho, \mathbf{b}) := \int_0^1 \int_{\mathcal{Y}} \tfrac{1}{2}\left\|\mathbf{b}_t(\mathbf{z})\right\|^2 \rho_t(\mathbf{z})\,\mathrm{d}\mathbf{z}\,\mathrm{d}t, \tag{S4}$$

among all pairs $(\rho, \mathbf{b})$ satisfying (S3) and matching the prescribed marginals. For constant isotropic diffusion $\Sigma = \sigma I$, this action equals $\sigma^2$ times the drift-dependent path-space KL cost. For a general diffusion matrix, the KL cost uses the inverse-covariance metric given below. The reference dependence is explicit in the path-space formulation. Let $\mathsf{R}$ be the path measure on $\Omega = C([0,1], \mathcal{Y})$ induced by a selected reference diffusion

$$\mathrm{d}\mathbf{Z}_t = \mathbf{a}_t(\mathbf{Z}_t)\,\mathrm{d}t + \Sigma_t(\mathbf{Z}_t)\,\mathrm{d}\mathbf{W}_t, \qquad D_t = \Sigma_t\Sigma_t^{\mathsf{T}}.$$

For prescribed endpoint distributions $\mu_0$ and $\mu_1$, the classical balanced Schrödinger bridge is the path measure

$$\mathsf{P}^\star = \underset{\substack{\mathsf{P}\ll\mathsf{R}:\\ (e_0)_\#\mathsf{P}=\mu_0,\ (e_1)_\#\mathsf{P}=\mu_1}}{\operatorname{argmin}} \mathrm{KL}(\mathsf{P}\,\|\,\mathsf{R}),$$

where $e_t(\omega) = \omega_t$. Thus, the bridge selects a path law that satisfies the endpoint constraints while remaining closest in relative entropy to the chosen reference dynamics. If an admissible process changes the drift by a control $\mathbf{c}_t$ while retaining the same diffusion coefficient, Girsanov's theorem gives, under the usual absolute-continuity and integrability conditions,

$$\mathrm{KL}(\mathsf{P}\,\|\,\mathsf{R}) = \mathrm{KL}(\mu_0\,\|\,\mathsf{R}_0) + \frac{1}{2}\mathbb{E}_{\mathsf{P}}\left[\int_0^1 \mathbf{c}_t(\mathbf{Z}_t)^{\mathsf{T}} D_t(\mathbf{Z}_t)^{-1}\mathbf{c}_t(\mathbf{Z}_t)\,\mathrm{d}t\right].$$

Because the initial marginal is prescribed, the first term is fixed. The corresponding density-level control penalty is therefore

$$\frac{1}{2}\int_0^1 \int_{\mathcal{Y}} \rho_t(\mathbf{z})\,\mathbf{c}_t(\mathbf{z})^{\mathsf{T}} D_t(\mathbf{z})^{-1}\mathbf{c}_t(\mathbf{z})\,\mathrm{d}\mathbf{z}\,\mathrm{d}t.$$

For an isotropic reference diffusion $D(t) = \sigma^2(t)I$, this becomes $\frac{1}{2}\int_0^1\int_{\mathcal{Y}} \rho_t(\mathbf{z})\left\|\mathbf{c}_t(\mathbf{z})\right\|^2/\sigma^2(t)\,\mathrm{d}\mathbf{z}\,\mathrm{d}t$. By Girsanov's theorem, this is the relative-entropy cost of changing the reference drift. The Euclidean expression in Eq. (S4) uses a fixed reference metric. For the mean-field formulation used by CytoBridge, the reference construction is nonlinear. Conditional on a specified interaction operator $\mathcal{I}$ and a candidate path law $\mathsf{P}$ with marginals $\rho_t = (e_t)_\#\mathsf{P}$, let $\Gamma_{\mathcal{I}}(\mathsf{P})$ denote the law of the uncontrolled McKean–Vlasov diffusion

$$\mathrm{d}\mathbf{Z}_t = \mathcal{I}[\rho_t](\mathbf{Z}_t)\,\mathrm{d}t + \Sigma_t(\mathbf{Z}_t)\,\mathrm{d}\mathbf{W}_t.$$

The mass-conserving mean-field Schrödinger bridge minimizes the large-deviation rate functional

$$\mathsf{P}^\star = \operatorname*{argmin}_{(e_0)_\#\mathsf{P}=\mu_0,\,(e_1)_\#\mathsf{P}=\mu_1} \mathrm{KL}\big(\mathsf{P}\,\|\,\Gamma_{\mathcal{I}}(\mathsf{P})\big),$$

which is the mean-field counterpart of the fixed-reference path-space problem above [9]. Its controlled representation adds an intrinsic drift $\mathbf{b}_t$ to the same interacting diffusion,

$$\mathrm{d}\mathbf{Z}_t = \big[\mathcal{I}[\rho_t](\mathbf{Z}_t) + \mathbf{b}_t(\mathbf{Z}_t)\big]\,\mathrm{d}t + \Sigma_t(\mathbf{Z}_t)\,\mathrm{d}\mathbf{W}_t.$$

Under the usual Girsanov conditions, the quadratic term is the path-space relative-entropy cost of the added control $\mathbf{b}_t$. When stochasticity and non-conserved population size must be handled simultaneously, regularized unbalanced optimal transport (RUOT), also known as an unbalanced Schrödinger bridge, provides a natural extension of the above SB picture by combining diffusion-driven dynamics with explicit growth or depletion terms. In a simple isotropic setting, the density evolves as

$$\partial_t\rho_t(\mathbf{z}) = -\nabla\cdot_{\mathbf{z}}\big(\rho_t(\mathbf{z})\,\mathbf{b}_t(\mathbf{z})\big) + \frac{\sigma^2(t)}{2}\Delta_{\mathbf{z}}\rho_t(\mathbf{z}) + g_t(\mathbf{z})\,\rho_t(\mathbf{z}),$$

and the corresponding objective augments the stochastic transport cost with a growth penalty,

$$\mathcal{A}_{\mathrm{RUOT}}(\rho,\mathbf{b},g) := \int_0^1\int_{\mathcal{Y}}\Big[\tfrac{1}{2}\,\|\mathbf{b}_t(\mathbf{z})\|^2 + \alpha\,\Psi(g_t(\mathbf{z}))\Big]\rho_t(\mathbf{z})\,\mathrm{d}\mathbf{z}\,\mathrm{d}t.$$

In the CytoBridge formulation, "reference" denotes the mass-conserving mean-field diffusion defined above. This process has the interaction drift $\mathcal{I}[\rho_t]$ and stochastic diffusion. The intrinsic-control field $\mathbf{b}_t$ and reaction field $g_t$ are introduced in the controlled, unbalanced formulation. This viewpoint is especially useful for learning stochastic developmental trajectories from snapshot observations [10]. The UMFSB formulation used by CytoBridge combines these ingredients: transport in the joint state space, stochastic regularization, unbalanced growth, and an explicit interaction field that depends on the current population.

#### S1.2 Data model, state space, and snapshot measures

We consider time-series spatial transcriptomic measurements at ordered experimental times

$$t_0 < t_1 < \cdots < t_K.$$

Experimental stages are ordered by their measured times and assigned consecutive model times $0, 1, \ldots, K$. Within each adjacent experimental interval, time is mapped linearly to the corresponding unit-length model interval. The theoretical derivation below uses $[0, 1]$ as a general time domain. Unless a conversion is stated, the learned drift and growth fields are rates in normalized model time rather than in days, hours, or months. At time $t_k$, we observe $N_k$ cells/spots indexed by $i \in \{1, \ldots, N_k\}$, each with (i) a transcriptomic embedding $\mathbf{x}_i^{(k)} \in \mathbb{R}^{d_x}$ (obtained from the high-dimensional gene expression by a fixed embedding map) and (ii) a physical coordinate $\mathbf{r}_i^{(k)} \in \mathbb{R}^{d_r}$. We write the joint state vector as

$$\mathbf{z} = (\mathbf{x},\mathbf{r}) \in \mathcal{Y} := \mathbb{R}^{d_x}\times\mathbb{R}^{d_r}.$$

Here $d_x$ is the dimension of the transcriptomic embedding and $d_r$ is the dimension of the spatial coordinate system. The empirical snapshot measure at $t_k$ is

$$\mu_k := \frac{1}{N_0}\sum_{i=1}^{N_k}\delta_{(\mathbf{x}_i^{(k)},\mathbf{r}_i^{(k)})} \in \mathcal{M}_+(\mathcal{Y}),$$

where $\mathcal{M}_+(\mathcal{Y})$ denotes finite nonnegative measures. The total mass $\mu_k(\mathcal{Y}) = N_k/N_0$ gives the population size relative to the initial observation. Normalizing $\mu_k$ by this mass gives the empirical probability distribution of cell states. The measure $\mu_k$ approximates the population measure $\mu_{t_k}$ with density $\rho_{t_k}$. Unless otherwise specified, when considering a fixed snapshot at time $t_k$, we drop the index $k$ and write $N$ and $(\mathbf{x}, \mathbf{r})$ in place of $N_k$ and $(\mathbf{x}^{(k)}, \mathbf{r}^{(k)})$. Our goal is to infer a continuous-time evolution $\{\mu_t\}_{t\in[0,1]}$ (with density $\rho_t$) and the underlying single-cell dynamics that interpolate the observed snapshots.

##### S1.3 Unbalanced mean-field diffusion with interaction operator

CytoBridge models population evolution via a controlled diffusion with unbalanced growth and mean-field interactions [11]. The probability-flow representation used for numerical optimization is derived below. Throughout this subsection, $\mathbf{b}_t$ denotes the intrinsic drift in the diffusive formulation, $\mathbf{v}_t$ denotes the corresponding probability-flow drift after the diffusion-to-flow conversion, and $\mathcal{I}[\rho_t]$ denotes the interaction field induced by the current population distribution. The deterministic transport velocity is written as $\mathbf{u}_t = \mathbf{v}_t + \mathcal{I}[\rho_t]$, so the intrinsic and interaction contributions remain explicit. At the population (density) level, we consider the generalized Fokker–Planck equation on $\mathcal{Y}$:

$$\partial_t \rho_t(\mathbf{z}) = -\nabla\cdot_{\mathbf{z}}\Big(\rho_t(\mathbf{z})\,[\mathbf{b}_t(\mathbf{z}) + \mathcal{I}[\rho_t](\mathbf{z})]\Big) + \frac{\sigma^2(t)}{2}\,\Delta_{\mathbf{z}}\rho_t(\mathbf{z}) + g_t(\mathbf{z})\,\rho_t(\mathbf{z}),$$

where $\mathbf{b}_t : \mathcal{Y} \to \mathbb{R}^{d_x+d_r}$ is the intrinsic-context drift, representing intrinsic regulation together with unmeasured contextual influences, $\sigma(t) \geq 0$ controls the diffusion intensity, $g_t : \mathcal{Y} \to \mathbb{R}$ is a growth field encoding net proliferation or death (and potentially absorbing sampling bias), and $\mathcal{I}[\rho_t] : \mathcal{Y} \to \mathbb{R}^{d_x+d_r}$ is a mean-field interaction operator that depends on the current population $\rho_t$.

**Kernel form as a special case.** A classical (pairwise) interaction specification is

$$\mathcal{I}[\rho](\mathbf{z}) := \int_{\mathcal{Y}} \kappa(\mathbf{z}, \mathbf{z}')\,\mathbf{K}(\mathbf{z}, \mathbf{z}')\,\rho(\mathbf{z}')\,\mathrm{d}\mathbf{z}', \tag{S5}$$

where $\kappa(\mathbf{z}, \mathbf{z}') \geq 0$ is a scalar interaction weight and $\mathbf{K}(\mathbf{z}, \mathbf{z}') \in \mathbb{R}^{d_x+d_r}$ is a vector-valued force kernel (for example, derived from a potential). This form is pairwise and depends on $\rho$ through an integral.

**General operator form.** To avoid restricting interactions to a fixed analytic kernel, we treat

$$\mathcal{I} : \mathcal{M}_+(\mathcal{Y}) \to \mathcal{V}(\mathcal{Y}),$$

where $\mathcal{M}_+(\mathcal{Y})$ denotes finite nonnegative measures, allowing population mass to vary with time. The operator maps a population measure to a vector field on $\mathcal{Y}$. Here $\mathcal{V}(\mathcal{Y})$ denotes a suitable function space of vector fields on $\mathcal{Y}$ (e.g., $L^2(\rho; \mathbb{R}^{d_x+d_r})$). CytoBridge instantiates $\mathcal{I}[\rho_t](\cdot)$ through message passing on a time-varying cell–cell graph (Supplementary Note S3), which can be viewed as a data-driven approximation of (S5) in which the integral is replaced by neighborhood aggregation.

##### S1.4 Variational principle and Fisher-information form

We derive the Fisher-information-regularized probability-flow form used for training.

##### S1.4.1 Unbalanced mean-field Schrödinger bridge action

Fix endpoint marginals $\rho_{t_0}$ and $\rho_{t_K}$ (or, in practice, match all intermediate snapshots via distributional losses). The UMFSB action functional on $[0, 1]$ takes the form [11]

$$\mathcal{J}(\rho, \mathbf{b}, g) := \int_0^1 \int_{\mathcal{Y}} \Big( \tfrac{1}{2} \|\mathbf{b}_t(\mathbf{z})\|^2 + \alpha\, \Psi(g_t(\mathbf{z})) \Big) \rho_t(\mathbf{z})\, \mathrm{d}\mathbf{z}\, \mathrm{d}t, \tag{S6}$$

The quadratic drift term in Eq. (S6) has the stochastic-control interpretation established in Supplementary Note S1.1. In the mass-conserving case with constant isotropic diffusion, it equals $\sigma^2$ times the path-space relative-entropy cost of adding the intrinsic control $\mathbf{b}_t$ to the mean-field reference diffusion. This motivates the quadratic drift penalty in the unbalanced objective. Its form is determined by the chosen state-space metric and reference diffusion.

The action in Eq. (S6) is minimized subject to the controlled diffusion constraint

$$\partial_t \rho_t = -\nabla \cdot_{\mathbf{z}} \big( \rho_t [\mathbf{b}_t + \mathcal{I}[\rho_t]] \big) + \tfrac{\sigma^2(t)}{2} \Delta_{\mathbf{z}} \rho_t + g_t \rho_t, \tag{S7}$$

where $\mathbf{b}_t$ plays the role of the intrinsic control drift in the diffusive formulation.

**Fisher–Rao interpretation of the quadratic growth penalty.** The unbalanced extension adds the reaction term $g_t \rho_t$ to the population equation. For a pure reaction tangent $\dot{\rho}_t = g_t \rho_t$, the Fisher–Rao squared norm is

$$\|\dot{\rho}_t\|^2_{\mathrm{FR}, \rho_t} := \int_{\mathcal{Y}} \frac{\dot{\rho}_t(\mathbf{z})^2}{\rho_t(\mathbf{z})}\, \mathrm{d}\mathbf{z} = \int_{\mathcal{Y}} \rho_t(\mathbf{z}) g_t(\mathbf{z})^2\, \mathrm{d}\mathbf{z}.$$

Thus, $\Psi(g) = g^2$ makes the density-weighted growth penalty in Eq. (S6) the Fisher–Rao reaction energy. It measures the size of a local relative change in population mass within the chosen density geometry. The coefficient $\alpha$ sets its scale relative to the drift-control cost. Birth–death reference models can lead to other reaction penalties [10].

**Classical models as limiting cases.** The controlled population equation in Eq. (S7) provides a common view of six related formulations (Supplementary Table 1). Setting $\mathcal{I}[\rho_t] \equiv 0$ and $g_t \equiv 0$ gives a standard Schrödinger bridge. Retaining $g_t$ while setting $\mathcal{I}[\rho_t] \equiv 0$ gives an unbalanced Schrödinger bridge, also described as regularized unbalanced optimal transport. Retaining $\mathcal{I}[\rho_t]$ while imposing $g_t \equiv 0$ gives a balanced mean-field Schrödinger bridge. Keeping both terms gives the unbalanced mean-field Schrödinger bridge used by the full CytoBridge model. With $\mathcal{I}[\rho_t] \equiv 0$, taking the vanishing-diffusion limit removes the stochastic score correction and yields dynamical optimal transport when $g_t = 0$ or unbalanced dynamical optimal transport when $g_t$ is retained.

Each restriction removes a fitted component. Setting $\sigma = 0$ removes the score and diffusion correction and makes transport deterministic, whereas setting $g_t = 0$ removes the growth output and its mass-change objective and conserves total mass. Similarly, setting $\mathcal{I}[\rho_t] = 0$ removes the interaction network, leaving all changes to be represented by the intrinsic drift, diffusion and, in the unbalanced case, growth. Supplementary Notes S4.8, S6 and S7 compare the full model with versions that omit the interaction term and with other transport and interpolation methods.

##### S1.4.2 Probability-flow drift, transport velocity, and Fisher information

Define the probability-flow drift

$$\mathbf{v}_t(\mathbf{z}) := \mathbf{b}_t(\mathbf{z}) - \tfrac{\sigma^2(t)}{2} \nabla_{\mathbf{z}} \log \rho_t(\mathbf{z}). \tag{S8}$$

and the deterministic transport velocity

$$\mathbf{u}_t(\mathbf{z}) := \mathbf{v}_t(\mathbf{z}) + \mathcal{I}[\rho_t](\mathbf{z}) = \mathbf{b}_t(\mathbf{z}) + \mathcal{I}[\rho_t](\mathbf{z}) - \tfrac{\sigma^2(t)}{2}\,\nabla_{\mathbf{z}} \log \rho_t(\mathbf{z}). \tag{S9}$$

Then (S7) is equivalent to the unbalanced continuity equation

$$\partial_t \rho_t + \nabla\cdot_{\mathbf{z}}(\rho_t \mathbf{u}_t) = g_t \rho_t. \tag{S10}$$

The price of replacing diffusion in (S7) by the deterministic flow (S10) is an explicit regularizer involving the Fisher information

$$\mathcal{F}(\rho_t) := \int_{\mathcal{Y}} \|\nabla_{\mathbf{z}} \log \rho_t(\mathbf{z})\|^2\, \rho_t(\mathbf{z})\,\mathrm{d}\mathbf{z}. \tag{S11}$$

**Theorem S1.1** (Fisher-information reformulation)**.** *Assume $\rho_t$ is smooth and strictly positive on $\mathcal{Y}$, with sufficient decay at infinity so that integrations by parts are valid. Let $\mathbf{v}_t$ and $\mathbf{u}_t$ be defined by* (S8) *and* (S9)*. Then the controlled Fokker–Planck equation* (S7) *is equivalent to the unbalanced continuity equation* (S10)*. Moreover, the kinetic term in* (S6) *admits the decomposition*

$$\int_{\mathcal{Y}} \tfrac{1}{2}\|\mathbf{b}_t(\mathbf{z})\|^2\, \rho_t(\mathbf{z})\,\mathrm{d}\mathbf{z} = \int_{\mathcal{Y}} \tfrac{1}{2}\|\mathbf{v}_t(\mathbf{z})\|^2\, \rho_t(\mathbf{z})\,\mathrm{d}\mathbf{z} + \frac{\sigma^4(t)}{8}\mathcal{F}(\rho_t) + \frac{\sigma^2(t)}{2}\int_{\mathcal{Y}} \langle \mathbf{v}_t, \nabla_{\mathbf{z}} \log \rho_t\rangle\, \rho_t\,\mathrm{d}\mathbf{z}. \tag{S12}$$

*Minimizing* (S6) *under* (S7) *is therefore equivalent to minimizing the reformulated action under* (S10)*, retaining the kinetic, Fisher-information, and cross terms in* (S12) *together with the growth penalty.*

*Proof.* **(i)** Starting from (S9), we have

$$\begin{aligned} -\nabla\cdot(\rho_t \mathbf{u}_t) &= -\nabla\cdot\Big(\rho_t(\mathbf{b}_t + \mathcal{I}[\rho_t])\Big) + \frac{\sigma^2(t)}{2}\nabla\cdot\Big(\rho_t \nabla_{\mathbf{z}} \log \rho_t\Big) \\ &= -\nabla\cdot\Big(\rho_t(\mathbf{b}_t + \mathcal{I}[\rho_t])\Big) + \frac{\sigma^2(t)}{2}\Delta_{\mathbf{z}}\rho_t, \end{aligned}$$

where we used $\nabla\cdot(\rho\nabla\log\rho) = \Delta\rho$. Substituting into (S7) yields (S10), and the converse follows by reversing the calculation.

**(ii)** Rearranging (S8) gives

$$\mathbf{b}_t = \mathbf{v}_t + \tfrac{\sigma^2(t)}{2}\nabla_{\mathbf{z}} \log \rho_t.$$

Expanding the square and integrating against $\rho_t$ yields

$$\int_{\mathcal{Y}} \tfrac{1}{2}\|\mathbf{b}_t\|^2 \rho_t = \int_{\mathcal{Y}} \tfrac{1}{2}\|\mathbf{v}_t\|^2 \rho_t + \frac{\sigma^4(t)}{8}\int_{\mathcal{Y}} \|\nabla_{\mathbf{z}} \log \rho_t\|^2 \rho_t + \frac{\sigma^2(t)}{2}\int_{\mathcal{Y}} \langle \mathbf{v}_t, \nabla_{\mathbf{z}} \log \rho_t\rangle\, \rho_t.$$

Recognizing the Fisher information (S11) gives (S12). □

### Supplementary Note S2. Particle-based approximation and numerical solver

#### S2.1 Weighted particle system for unbalanced continuity equations

We approximate the population density by interacting particles whose weights change with the growth field.

We consider the deterministic unbalanced flow constraint

$$\partial_t \rho_t + \nabla\cdot(\rho_t \mathbf{u}_t) = g_t \rho_t, \qquad t \in [0,1], \tag{S13}$$

with given initial density $\rho_0$. A natural approximation is a weighted particle system. Let $\{\mathbf{z}_i(t)\}_{i=1}^N$ be particles and $\{w_i(t)\}_{i=1}^N$ be nonnegative weights evolving by

$$\dot{\mathbf{z}}_i(t) = \mathbf{u}_t(\mathbf{z}_i(t)), \qquad \dot{w}_i(t) = g_t(\mathbf{z}_i(t))\, w_i(t), \tag{S14}$$

with initialization $\mathbf{z}_i(0) \sim \rho_0$ and $w_i(0) = 1$. Define the weighted empirical measure

$$\rho_t^{(N)} := \frac{1}{N}\sum_{i=1}^N w_i(t)\, \delta_{\mathbf{z}_i(t)}. \tag{S15}$$

**Proposition S2.1** (Particle representation in weak form)**.** *Assume $\mathbf{u}_t$ and $g_t$ are continuous and globally Lipschitz in $\mathbf{z}$. Then for any test function $\varphi \in C_c^\infty(\mathcal{Y})$, the empirical measure* (S15) *satisfies*

$$\frac{\mathrm{d}}{\mathrm{d}t}\int_{\mathcal{Y}} \varphi(\mathbf{z})\, \rho_t^{(N)}(\mathrm{d}\mathbf{z}) = \int_{\mathcal{Y}} \Big( \langle \nabla\varphi(\mathbf{z}), \mathbf{u}_t(\mathbf{z})\rangle + g_t(\mathbf{z})\,\varphi(\mathbf{z})\Big)\, \rho_t^{(N)}(\mathrm{d}\mathbf{z}). \tag{S16}$$

*Consequently, any weak limit point of $\{\rho_t^{(N)}\}_{N\to\infty}$ solves* (S13) *in the weak sense.*

*Proof.* By definition,

$$\int \varphi\, \rho_t^{(N)} = \frac{1}{N}\sum_{i=1}^N w_i(t)\, \varphi(\mathbf{z}_i(t)).$$

Differentiating and using (S14) yields

$$\begin{aligned}
\frac{\mathrm{d}}{\mathrm{d}t}\int \varphi\, \rho_t^{(N)} &= \frac{1}{N}\sum_{i=1}^N \dot{w}_i\, \varphi(\mathbf{z}_i) + \frac{1}{N}\sum_{i=1}^N w_i\, \langle \nabla\varphi(\mathbf{z}_i), \dot{\mathbf{z}}_i\rangle \\
&= \frac{1}{N}\sum_{i=1}^N w_i\, g_t(\mathbf{z}_i)\, \varphi(\mathbf{z}_i) + \frac{1}{N}\sum_{i=1}^N w_i\, \langle \nabla\varphi(\mathbf{z}_i), \mathbf{u}_t(\mathbf{z}_i)\rangle,
\end{aligned}$$

which is exactly (S16). □

#### S2.2 Interaction-aware probability-flow and transport particle system

In CytoBridge, the probability-flow drift $\mathbf{v}_t$ is obtained from the intrinsic drift $\mathbf{b}_t$ through the diffusion-to-flow correction in Theorem S1.1:

$$\mathbf{v}_t(\mathbf{z}) = \mathbf{b}_t(\mathbf{z}) - \frac{\sigma^2(t)}{2}\, \nabla_{\mathbf{z}} \log \rho_t(\mathbf{z}).$$

The total transport velocity is then

$$\mathbf{u}_t(\mathbf{z}) := \mathbf{v}_t(\mathbf{z}) + \mathcal{I}[\rho_t](\mathbf{z}) = \mathbf{b}_t(\mathbf{z}) + \mathcal{I}[\rho_t](\mathbf{z}) - \frac{\sigma^2(t)}{2}\, \nabla_{\mathbf{z}} \log \rho_t(\mathbf{z}).$$

Here $\mathbf{b}_t$ denotes an intrinsic drift field, and $\mathcal{I}[\rho_t]$ is an interaction operator that depends on the population distribution (implemented in practice by a dynamic-graph GNN). In the particle approximation, $\rho_t$ is replaced by the weighted empirical measure $\rho_t^{(N)}$ in (S15), leading to an empirical interaction term

$$\mathcal{I}_t^{(N)}(\mathbf{z}) := \mathcal{I}[\rho_t^{(N)}](\mathbf{z}),$$

an empirical probability-flow drift

$$\mathbf{v}_t^{(N)}(\mathbf{z}) := \mathbf{b}_t(\mathbf{z}) - \frac{\sigma^2(t)}{2}\,\mathbf{s}_t(\mathbf{z}),$$

and the corresponding empirical transport velocity

$$\mathbf{u}_t^{(N)}(\mathbf{z}) := \mathbf{v}_t^{(N)}(\mathbf{z}) + \mathcal{I}[\rho_t^{(N)}](\mathbf{z}),$$

where $\mathbf{s}_t(\mathbf{z}) = \nabla_{\mathbf{z}} \log \rho_t(\mathbf{z})$ is a score estimate (learned jointly or estimated from particles, as detailed in Supplementary Note S2.5). We use the empirical interaction field $\mathcal{I}_t^{(N)}$ to approximate $\mathcal{I}[\rho_t]$. Accordingly, the weighted particle dynamics (S14) should be understood as

$$\dot{\mathbf{z}}_i(t) = \mathbf{u}_t^{(N)}(\mathbf{z}_i(t)) = \mathbf{b}_t(\mathbf{z}_i(t)) + \mathcal{I}[\rho_t^{(N)}](\mathbf{z}_i(t)) - \frac{\sigma^2(t)}{2}\,\mathbf{s}_t(\mathbf{z}_i(t)), \qquad \dot{w}_i(t) = g_t(\mathbf{z}_i(t))\,w_i(t),$$

which makes the empirical interaction contribution explicit.

##### S2.3 Mean-field limit of weighted interacting particles

We show that a limit of the weighted interacting particle system satisfies the mean-field equation. The argument accounts for changing population mass and the disappearance of particle-level Brownian fluctuations in the empirical measure.

**Assumption S2.2** (Regularity of drift, growth and interaction operator)**.** *Work on $\mathcal{Y} = \mathbb{R}^d$. Assume $\mathbf{b}_t$ and $g_t$ are bounded, continuous in $t$, and globally Lipschitz in $\mathbf{z}$, uniformly in $t \in [0, 1]$, with $|g_t| \le G$. Assume $\sigma(t)$ is bounded and continuous. For finite nonnegative measures of mass at most $e^G$, assume $\mathcal{I}[\mu]$ is uniformly bounded and Lipschitz in $\mathbf{z}$ and satisfies*

$$\sup_{\mathbf{z}} \|\mathcal{I}[\mu](\mathbf{z}) - \mathcal{I}[\nu](\mathbf{z})\| \le L_\mu\, d_{\mathrm{BL}}(\mu, \nu),$$

*where $d_{\mathrm{BL}}(\mu, \nu) = \sup_{\|f\|_\infty + \mathrm{Lip}(f) \le 1} |\int f\, \mathrm{d}(\mu - \nu)|$ is the bounded-Lipschitz distance. This distance allows unequal masses and metrizes weak convergence of finite nonnegative measures. Bounded Lipschitz convolution kernels satisfy these assumptions.*

**Microscopic interacting system.** Let $(\mathbf{B}_i)_{i=1}^N$ be i.i.d. $d$-dimensional Brownian motions. Define the (weighted) empirical measure

$$\rho_t^{(N)} := \frac{1}{N}\sum_{i=1}^N w_i(t)\,\delta_{\mathbf{z}_i(t)}.$$

Consider the interacting diffusion with unbalanced weights

$$\mathrm{d}\mathbf{z}_i(t) = \big(\mathbf{b}_t(\mathbf{z}_i(t)) + \mathcal{I}[\rho_t^{(N)}](\mathbf{z}_i(t))\big)\,\mathrm{d}t + \sigma(t)\,\mathrm{d}\mathbf{B}_i(t), \qquad \mathrm{d}w_i(t) = g_t(\mathbf{z}_i(t))\,w_i(t)\,\mathrm{d}t, \tag{S17}$$

with i.i.d. initialization $\mathbf{z}_i(0) \sim \rho_0$, where $\rho_0$ is a probability measure, and $w_i(0) = 1$.

**Theorem S2.3** (Identification of the mean-field limit)**.** *Under Assumption S2.2, any limit in law of* $\rho^{(N)}$ *in* $C([0,1], \mathcal{M}_+(\mathcal{Y}))$*, equipped with the uniform* $d_{\mathrm{BL}}$ *topology, almost surely satisfies the mean-field controlled Fokker–Planck equation with initial measure* $\rho_0$*,*

$$\partial_t \rho_t = -\nabla \cdot_{\mathbf{z}} \big(\rho_t \, (\mathbf{b}_t + \mathcal{I}[\rho_t])\big) + \tfrac{\sigma^2(t)}{2} \, \Delta_{\mathbf{z}} \rho_t + g_t \, \rho_t.$$

*If the limit has a positive, sufficiently smooth density, defining* $\mathbf{v}_t = \mathbf{b}_t - \frac{\sigma^2(t)}{2} \nabla_{\mathbf{z}} \log \rho_t$ *and* $\mathbf{u}_t = \mathbf{v}_t + \mathcal{I}[\rho_t]$ *gives the unbalanced continuity equation* (S10)*.*

*Proof.* Fix $\varphi \in C_c^\infty(\mathcal{Y})$ and write $\langle \varphi, \rho_t^{(N)} \rangle := \frac{1}{N} \sum_{i=1}^N w_i(t) \, \varphi(\mathbf{z}_i(t))$. Applying Itô's formula to $w_i(t) \, \varphi(\mathbf{z}_i(t))$ under (S17) gives

$$\begin{aligned} \mathrm{d}\big(w_i \varphi(\mathbf{z}_i)\big) = w_i \, \langle \nabla \varphi(\mathbf{z}_i), \mathbf{b}_t(\mathbf{z}_i) + \mathcal{I}[\rho_t^{(N)}](\mathbf{z}_i) \rangle \, \mathrm{d}t + \tfrac{\sigma^2(t)}{2} \, w_i \, \Delta \varphi(\mathbf{z}_i) \, \mathrm{d}t \\ + g_t(\mathbf{z}_i) \, w_i \, \varphi(\mathbf{z}_i) \, \mathrm{d}t + \sigma(t) \, w_i \, \langle \nabla \varphi(\mathbf{z}_i), \mathrm{d}\mathbf{B}_i \rangle. \end{aligned} \tag{S18}$$

Summing (S18) over $i$ and dividing by $N$ yields the (stochastic) weak evolution

$$\mathrm{d}\langle \varphi, \rho_t^{(N)} \rangle = \Big\langle \langle \nabla \varphi, \mathbf{b}_t \rangle + \tfrac{\sigma^2(t)}{2} \, \Delta \varphi + g_t \, \varphi, \, \rho_t^{(N)} \Big\rangle \, \mathrm{d}t + \int_{\mathcal{Y}} \langle \nabla \varphi(\mathbf{z}), \mathcal{I}[\rho_t^{(N)}](\mathbf{z}) \rangle \, \rho_t^{(N)}(\mathrm{d}\mathbf{z}) \, \mathrm{d}t + \mathrm{d}M_t^{(N)}, \tag{S19}$$

where $M_t^{(N)}$ is the summed Brownian martingale. Since $w_i(t) = \exp(\int_0^t g_\tau(\mathbf{z}_i(\tau)) \, \mathrm{d}\tau) \le e^G$, its quadratic variation and Doob's inequality give

$$\mathbb{E} \sup_{0 \le t \le 1} |M_t^{(N)}|^2 \le \frac{4 e^{2G} \|\sigma\|_\infty^2 \|\nabla \varphi\|_\infty^2}{N} \longrightarrow 0.$$

The empirical masses are bounded by $e^G$. On this mass-bounded set, Assumption S2.2 makes $\mu \mapsto \int \langle \nabla \varphi, \mathcal{I}[\mu] \rangle \, \mathrm{d}\mu$ continuous in $d_{\mathrm{BL}}$: the change in the field is controlled uniformly by the measure distance, and the remaining integrand is bounded and Lipschitz. The other weak-form terms are continuous for the same reason. Thus, along a convergent subsequence, the continuous mapping theorem applied to the integrated form of (S19), together with $M^{(N)} \to 0$ in probability, yields almost surely

$$\langle \varphi, \rho_t \rangle - \langle \varphi, \rho_0 \rangle = \int_0^t \Big\langle \langle \nabla \varphi, \mathbf{b}_s + \mathcal{I}[\rho_s] \rangle + \tfrac{\sigma^2(s)}{2} \Delta \varphi + g_s \varphi, \rho_s \Big\rangle \, \mathrm{d}s.$$

The initial empirical measure converges to $\rho_0$ by the law of large numbers. Using a countable dense set of smooth test functions and then continuity gives the weak formulation for all test functions. The probability-flow equivalence follows by substituting $\rho \nabla \log \rho = \nabla \rho$ when the density is positive and smooth. □

#### S2.4 Numerical integration and snapshot matching

We work in the deterministic continuity formulation induced by the probability-flow drift

$$\partial_t \rho_t + \nabla_{\mathbf{z}} \cdot (\rho_t \mathbf{u}_t) = g_t \rho_t, \qquad \mathbf{u}_t(\mathbf{z}) = \mathbf{v}_t(\mathbf{z}) + \mathcal{I}[\rho_t](\mathbf{z}), \tag{S20}$$

where the probability-flow drift is defined by

$$\mathbf{v}_t(\mathbf{z}) \; := \; \mathbf{u}_t(\mathbf{z}) - \mathcal{I}[\rho_t](\mathbf{z}) \; = \; \mathbf{b}_t(\mathbf{z}) - \frac{\sigma^2(t)}{2} \nabla_{\mathbf{z}} \log \rho_t(\mathbf{z}). \tag{S21}$$

In practice, the probability-flow drift $\mathbf{v}_t$ and growth field $g_t$ are parameterized by neural networks, the score is estimated by $\mathbf{s}_\theta \approx \nabla_{\mathbf{z}} \log \rho_t$, and $\mathcal{I}[\rho_t]$ is realized by a time-varying interaction graph and an equivariant GNN. The intrinsic diffusive drift can then be recovered as $\mathbf{b}_t = \mathbf{v}_t + \frac{\sigma^2(t)}{2} \mathbf{s}_\theta$ whenever the original SDE representation is needed.

**Weighted interacting particle solver.** Let $\{\mathbf{z}_i(t), w_i(t)\}_{i=1}^N$ be weighted particles. We evolve

$$\dot{\mathbf{z}}_i(t) = \mathbf{u}_\theta(\mathbf{z}_i(t), t) = \mathbf{v}_\theta(\mathbf{z}_i(t), t) + \mathcal{I}_\theta\big[\rho_t^{(N)}\big](\mathbf{z}_i(t), t), \qquad \dot{w}_i(t) = g_\theta(\mathbf{z}_i(t), t)\, w_i(t), \tag{S22}$$

with initialization $\mathbf{z}_i(0) \sim \mu_0$ and $w_i(0) = 1$. Here

$$\rho_t^{(N)} := \frac{1}{N}\sum_{i=1}^N w_i(t)\, \delta_{\mathbf{z}_i(t)}$$

is the weighted empirical measure, and $\mathcal{I}_\theta[\rho_t^{(N)}]$ evaluates interactions among the simulated particles.

We integrate (S22) with a standard ODE solver (e.g. RK4) on $t \in [0, 1]$, using a fixed set of solver nodes $\{t_m\}_{m=0}^M$ (or adaptive nodes). For time-series ST, we also record particle states at experimental times $\{t_k\}_{k=0}^K$ for snapshot matching. This continuous-time parameterization is closely related to neural ODE-style modeling of latent dynamics [12].

**Snapshot matching loss.** Let $\widehat{\mu}_{t_k}^{(N)} := \frac{1}{N}\sum_{i=1}^N w_i(t_k)\delta_{\mathbf{z}_i(t_k)}$ be the predicted weighted snapshot. We match $\widehat{\mu}_{t_k}^{(N)}$ to the observed empirical snapshot $\mu_k$ using a composite distributional loss (e.g. unbalanced mass matching + normalized transport in joint space). The implementation stores particle masses $m_i(t) = w_i(t)/N$, initialized at $1/N$, so that $\widehat{\mu}_t^{(N)} = \sum_i m_i(t)\delta_{\mathbf{z}_i(t)}$. For the balanced transport term, we compare $\widehat{\mu}_{t_k}^{(N)}/\widehat{\mu}_{t_k}^{(N)}(\mathcal{Y})$ with $\mu_k/\mu_k(\mathcal{Y})$, so that both have unit total mass. The local mass term compares $m_i(t_k)$ with $(N_k/N_0)\,\mathrm{card}(h_k^{-1}(\mathbf{z}_i(t_k)))/B_k$, where $B_k$ is the number of observed points in the matched batch and $h_k$ assigns each of them to its nearest simulated particle. The global term compares $\sum_i m_i(t_k)$ with $N_k/N_0$.

##### S2.5 Energy loss, weight function, and Monte Carlo estimator

**Energy integrand in Fisher form.** Under Theorem S1.1, the UMFSB action contains a kinetic term and Fisher-information regularization, plus growth penalty:

$$\begin{aligned}\mathcal{J}(\rho, \mathbf{v}, g) = \int_0^1 \int_{\mathcal{Y}} \Big(&\frac{1}{2}\|\mathbf{v}_t(\mathbf{z})\|^2 + \frac{\sigma^4(t)}{8}\,\|\nabla_{\mathbf{z}} \log \rho_t(\mathbf{z})\|^2 \\ &+ \frac{\sigma^2(t)}{2}\,\langle \mathbf{v}_t(\mathbf{z}), \nabla_{\mathbf{z}} \log \rho_t(\mathbf{z})\rangle + \alpha\, \Psi(g_t(\mathbf{z}))\Big)\rho_t(\mathbf{z})\, d\mathbf{z}\, dt.\end{aligned} \tag{S23}$$

Here $\mathbf{v}_t$ is the probability-flow drift defined in (S21), i.e. $\mathbf{v}_t = \mathbf{b}_t - \frac{\sigma^2}{2}\nabla \log \rho_t$, while the interaction component remains explicit in the transport velocity $\mathbf{u}_t = \mathbf{v}_t + \mathcal{I}[\rho_t]$ in (S20).

**Trajectory weight induced by growth.** For the weighted particle dynamics (S22), the weight admits the closed form

$$w_i(t) = \exp\Big(\int_0^t g_\theta(\mathbf{z}_i(\tau), \tau)\, d\tau\Big). \tag{S24}$$

Thus, growth exponentially reweights particles along their trajectories. We will use (S24) to build a Monte Carlo estimator of (S23).

**Upper bound for training.** Directly optimizing (S23) can be numerically unstable because the cross term $\langle \mathbf{v}_t, \nabla \log \rho_t \rangle$ couples the optimization of the drift and the score. We bound this term using the Cauchy–Schwarz inequality:

$$\langle \mathbf{v}_t, \nabla \log \rho_t \rangle \;\le\; \|\mathbf{v}_t\| \, \|\nabla \log \rho_t\|. \tag{S25}$$

Substituting (S25) into (S23) yields the training surrogate

$$\mathcal{J}(\rho, \mathbf{v}, g) \;\le\; \int_0^1 \int_{\mathcal{Y}} \Big( \underbrace{\frac{1}{2}\|\mathbf{v}_t\|^2}_{\text{drift energy}} + \underbrace{\frac{\sigma^4(t)}{8}\|\nabla \log \rho_t\|^2}_{\text{Fisher/score energy}} + \underbrace{\frac{\sigma^2(t)}{2}\|\mathbf{v}_t\| \, \|\nabla \log \rho_t\|}_{\text{coupling term}} + \alpha\, \Psi(g_t) \Big) \rho_t \, d\mathbf{z} \, dt. \tag{S26}$$

**Monte Carlo estimator along trajectories.** Let $\{\mathbf{z}_i(t_m), w_i(t_m)\}$ be the solver states along the integration grid $\{t_m\}_{m=0}^{M}$, and let $\Delta t_m := t_{m+1} - t_m$. Define the score field

$$\mathbf{s}_\theta(\mathbf{z}, t) := \frac{2}{\sigma^2(t)} \nabla_{\mathbf{z}} \phi_\theta(\mathbf{z}, t) \approx \nabla_{\mathbf{z}} \log \rho_t(\mathbf{z}),$$

for $\sigma(t) > 0$. The score network parameterizes the scalar potential $\phi_\theta$. Its gradient gives the score contribution to the drift,

$$\mathbf{c}_\theta(\mathbf{z}, t) := \nabla_{\mathbf{z}} \phi_\theta(\mathbf{z}, t) = \frac{\sigma^2(t)}{2} \mathbf{s}_\theta(\mathbf{z}, t). \tag{S27}$$

We directly parameterize the probability-flow drift $\mathbf{v}_\theta(\mathbf{z}, t)$. The corresponding intrinsic diffusive drift is recovered by

$$\mathbf{b}_\theta(\mathbf{z}, t) := \mathbf{v}_\theta(\mathbf{z}, t) + \mathbf{c}_\theta(\mathbf{z}, t) = \mathbf{v}_\theta(\mathbf{z}, t) + \frac{\sigma^2(t)}{2} \mathbf{s}_\theta(\mathbf{z}, t).$$

Then the energy upper bound (S26) admits the estimator

$$\begin{aligned} \mathcal{L}_{\text{energy}} := \frac{1}{N} \sum_{i=1}^{N} \sum_{m=0}^{M-1} \Big[ & \frac{1}{2}\|\mathbf{v}_\theta(\mathbf{z}_i(t_m), t_m)\|^2 + \frac{\sigma(t_m)^4}{8} \|\mathbf{s}_\theta(\mathbf{z}_i(t_m), t_m)\|^2 \\ & + \frac{\sigma(t_m)^2}{2} \|\mathbf{v}_\theta(\mathbf{z}_i(t_m), t_m)\| \, \|\mathbf{s}_\theta(\mathbf{z}_i(t_m), t_m)\| \\ & + \alpha\, \Psi(g_\theta(\mathbf{z}_i(t_m), t_m)) \Big] \; w_i(t_m) \; \Delta t_m. \end{aligned} \tag{S28}$$

**Combined training objective.** In the combined training objective, the energy loss (S28) is added to the snapshot matching loss and (optionally) a physics-informed residual enforcing (S20) in weak/PINN form.

$$\mathcal{L}_{\text{total}} = \mathcal{L}_{\text{energy}} + \lambda_{\text{recon}} \, \mathcal{L}_{\text{recon}} + \lambda_{\text{FP}} \, \mathcal{L}_{\text{FP}}.$$

The training procedure is summarized below.

#### Training procedure

Algorithm 1 summarizes the training loop. Each iteration alternates between forward simulation of weighted interacting particles in the joint transcriptomic–spatial state space and parameter updates based on snapshot-matching, energy, and optional continuity residual losses.

After training, trajectories and derived quantities are obtained by forward simulation as summarized in Algorithm 2.

**Algorithm 1** CytoBridge training

**Require:** Time-series snapshots $\{\mu_k\}_{k=0}^{K}$ at times $\{t_k\}_{k=0}^{K}$ in joint space $\mathcal{Y}$, hyperparameters $(N, M)$ (particles, solver nodes), weights $\lambda_{\text{recon}}, \lambda_{\text{FP}}$, noise schedule $\sigma(t)$, growth penalty $\Psi(\cdot)$ and energy weight $\alpha$.

**Require:** Parametric fields including probability-flow drift $\mathbf{v}_\theta(\mathbf{z}, t)$, growth $g_\theta(\mathbf{z}, t)$, score $\mathbf{s}_\theta(\mathbf{z}, t)$, and interaction operator $\mathcal{I}_\theta[\cdot](\mathbf{z}, t)$ realized by a time-varying graph GNN.

1: **for** each training iteration **do**

2: Sample an initial mini-batch $\{\mathbf{z}_i(0)\}_{i=1}^{N} \sim \mu_0$ and set $w_i(0) \leftarrow 1$.

3: Build a time-varying cell–cell graph (from spatial proximity and ligand–receptor priors) for the current mini-batch.

4: **Forward simulate** weighted interacting particles.

$$\dot{\mathbf{z}}_i(t) = \mathbf{v}_\theta(\mathbf{z}_i(t), t) + \mathcal{I}_\theta\big[\rho_t^{(N)}\big](\mathbf{z}_i(t), t), \qquad \dot{w}_i(t) = g_\theta(\mathbf{z}_i(t), t)\, w_i(t),$$

to obtain $\{\mathbf{z}_i(t_m), w_i(t_m)\}_{m=0}^{M}$ and predicted snapshots $\widehat{\mu}_{t_k}^{(N)} = \frac{1}{N}\sum_{i=1}^{N} w_i(t_k)\delta_{\mathbf{z}_i(t_k)}$.

5: Compute **snapshot matching loss** $\mathcal{L}_{\text{recon}}$ between $\widehat{\mu}_{t_k}^{(N)}$ and $\mu_k$ over $k = 1, \ldots, K$.

6: Compute **energy loss** $\mathcal{L}_{\text{energy}}$ by Monte Carlo quadrature along trajectories.

$$\mathcal{L}_{\text{energy}} \approx \frac{1}{N}\sum_{i=1}^{N}\sum_{m=0}^{M-1}\Big[\ell_{\text{energy}}\big(\mathbf{z}_i(t_m), t_m\big)\, w_i(t_m)\Big]\Delta t_m,$$

where $\ell_{\text{energy}}$ corresponds to the training upper bound in (S26).

7: Optionally compute **continuity/Fokker–Planck residual** $\mathcal{L}_{\text{FP}}$ (weak-form / PINN) enforcing the transport/continuity constraint.

8: Update parameters $\theta$ by minimizing

$$\mathcal{L}_{\text{total}} = \mathcal{L}_{\text{energy}} + \lambda_{\text{recon}}\mathcal{L}_{\text{recon}} + \lambda_{\text{FP}}\mathcal{L}_{\text{FP}}.$$

9: **end for**

10: **return** Trained model $(\mathbf{v}_\theta, g_\theta, \mathbf{s}_\theta, \mathcal{I}_\theta)$.

**Algorithm 2** CytoBridge inference

**Require:** Trained $(\mathbf{v}_\theta, g_\theta, \mathbf{s}_\theta, \mathcal{I}_\theta)$ and initial distribution $\mu_{t_0}$ (or sampled cells).

1: Sample particles $\{\mathbf{z}_i(t_0)\}_{i=1}^{N} \sim \mu_{t_0}$, set $w_i(t_0) \leftarrow 1$.

2: Forward simulate the weighted SDE

$$\begin{aligned} \mathrm{d}\mathbf{z}_i(t) &= \Big[\mathbf{v}_\theta(\mathbf{z}_i(t), t) + \frac{\sigma^2(t)}{2}\mathbf{s}_\theta(\mathbf{z}_i(t), t) + \mathcal{I}_\theta\big[\rho_t^{(N)}\big](\mathbf{z}_i(t), t)\Big]\mathrm{d}t + \sigma(t)\,\mathrm{d}\mathbf{B}_i(t), \\ \mathrm{d}w_i(t) &= g_\theta(\mathbf{z}_i(t), t)\, w_i(t)\,\mathrm{d}t. \end{aligned}$$

to obtain stochastic trajectories $\{\mathbf{z}_i(t), w_i(t)\}$.

3: Read out probability-flow drift $\mathbf{v}_t = \mathbf{v}_\theta$, intrinsic drift $\mathbf{b}_t = \mathbf{v}_\theta + \frac{\sigma^2(t)}{2}\mathbf{s}_\theta$, transport velocity $\mathbf{u}_t = \mathbf{v}_t + \mathcal{I}_\theta[\rho_t^{(N)}]$, growth $g_\theta$, and interaction component via $\mathcal{I}_\theta$.

4: **return** Trajectories $\{\mathbf{z}_i(t), w_i(t)\}$ and the fields evaluated along them.

### Supplementary Note S3. Interaction graphs and equivariant GNN parameterization

#### S3.1 Dynamic cell–cell graph construction with ligand–receptor priors

At each time point, CytoBridge constructs a directed graph

$$\mathcal{G}_t = (\mathcal{V}_t, \mathcal{E}_t),$$

whose nodes are observed or simulated cells and whose edges represent candidate interactions. At observed stages, cell pairs are labeled using spatial proximity and ligand–receptor compatibility. An edge predictor is trained on the concatenated states of ordered cell pairs using binary cross-entropy. Its parameters are then held fixed during dynamics training and prediction.

As cells move and their expression states change, candidate pairs are recomputed within the spatial cutoff and retained when the edge-predictor score exceeds the selected threshold. Thus, the graph changes with the simulated population while using the same learned rule for ligand–receptor compatibility. The spatial cutoff and edge-predictor threshold are determined as described in Supplementary Note S4.5.

#### S3.2 Equivariant attention GNN as an interaction operator

CytoBridge maps the current graph to interaction-induced changes in gene expression and spatial position with an equivariant attention GNN (equations in Methods, *GNN-based interaction operator*). For an edge from sender $j$ to receiver $i$, the distance $\|\mathbf{r}_j - \mathbf{r}_i\|$ is expanded in radial basis functions and, together with the expression embeddings of the two cells, determines the attention weight that each head assigns to the message from $j$. Scalar messages update the expression features, and learned scalar coefficients multiply the relative direction $(\mathbf{r}_j - \mathbf{r}_i)/\|\mathbf{r}_j - \mathbf{r}_i\|$ to give vector messages, so the spatial update is equivariant to rotations and translations. Messages are weighted by the sender particle weights and normalized by the total incoming weight, and the scalar and vector readouts give

$$\mathcal{I}_\psi(\mathbf{z}_i, t; \mathcal{G}_t) = \left(\Delta\mathbf{x}_i^{\text{int}},\, \Delta\mathbf{r}_i^{\text{int}}\right). \tag{S29}$$

We aggregate the learned attention weights by sender and receiver cell type at each time point to summarize interaction patterns.

#### S3.3 Identifiability of the dynamical components

We consider how accurately snapshots determine the population distribution over time, and when these changes can be attributed to intrinsic-context velocity, growth and interaction. The first result uses the UMFSB action to control changes between observations. The second shows that, within a specified model, the three components can be recovered if their effects on the population can be distinguished.

**Reconstructing the population between observations.** On the compact periodic state domain $X = \mathbb{T}^d$, consider

$$\partial_t \rho = -\nabla \cdot \{\rho(b + I[\rho])\} + \nu \Delta \rho + \rho g, \tag{S30}$$

$$\mathcal{A}(\rho, b, g) = \int_0^T \int_X \rho \left(\tfrac{1}{2}|b|^2 + \alpha g^2\right) \mathrm{d}x\, \mathrm{d}t, \qquad \alpha > 0,$$

where $b$ is intrinsic drift, $g$ is net growth and $\nu > 0$ is fixed. For positive smooth densities, the probability-flow velocity is $v = b - \nu\nabla \log \rho$, giving

$$\partial_t \rho = -\nabla \cdot \{\rho(v + I[\rho])\} + \rho g. \tag{S31}$$

The total transport velocity is $v + I[\rho]$. Define a distance between finite nonnegative population measures by

$$d(\lambda, \zeta) = \sup_{\|f\|_{C^2(X)} \leq 1} |\langle f, \lambda - \zeta\rangle|.$$

Here $\langle f, \lambda\rangle = \int_X f \, \mathrm{d}\lambda$, and the $C^2$ norm controls $f$ and its first two spatial derivatives. The distance includes total mass.

**Proposition S3.1** (Recovery of the population path)**.** *Let the true path $\mu$ and reconstructed paths $\rho^n$ satisfy* (S30)*, with uniformly bounded actions, initial masses and interaction fields. On observation grids containing $0, T$, let $h_n$ be the largest time gap. Suppose the observations $\widehat{\mu}_{n,k}$ and reconstructions satisfy*

$$\max_k d(\widehat{\mu}_{n,k}, \mu_{t_{n,k}}) \leq \varepsilon_n, \qquad \max_k d(\rho^n_{t_{n,k}}, \widehat{\mu}_{n,k}) \leq \tau_n.$$

*Then, with constants independent of $n$,*

$$\delta_n := \sup_t d(\rho^n_t, \mu_t) \leq C_1\sqrt{h_n} + C_2 h_n + \varepsilon_n + \tau_n. \tag{S32}$$

*Thus the population paths converge uniformly in $d$ when the gaps and errors vanish.*

*Proof.* Writing $m(t) = \rho_t(X)$, the growth equation and Cauchy–Schwarz give

$$\sqrt{m(t)} \leq \sqrt{m(0)} + \tfrac{1}{2}\sqrt{T\mathcal{A}/\alpha}.$$

Testing (S30) against $\|f\|_{C^2} \leq 1$ bounds the drift and growth contributions over $[s, t]$ by $C\sqrt{|t-s|}$, and the bounded interaction and diffusion contributions by $C'|t - s|$. Both paths therefore satisfy $d(\rho_t, \rho_s) \leq C\sqrt{|t-s|} + C'|t-s|$ with uniform constants. Choose an observed time $t_{n,k}$ within $h_n$ of $t$. The triangle inequality gives

$$\begin{aligned} d(\rho^n_t, \mu_t) \leq{}& d(\rho^n_t, \rho^n_{t_{n,k}}) + d(\rho^n_{t_{n,k}}, \widehat{\mu}_{n,k}) \\ &+ d(\widehat{\mu}_{n,k}, \mu_{t_{n,k}}) + d(\mu_{t_{n,k}}, \mu_t). \end{aligned}$$

The first and last terms are bounded by the time regularity of the two paths, and the middle terms by the fitting and observation errors $\tau_n$ and $\varepsilon_n$, which gives (S32). □

An exact UMFSB minimizer has no greater action than a feasible comparison path. Comparison paths with a common action bound therefore give the required bound for minimizers, including approximate minimizers with bounded optimization error.

**Separating intrinsic-context velocity, growth and interaction.** For a fixed state representation and measured input features, write each component as a sum of specified functions, with separate coefficient vectors $a$, $c$ and $\theta$:

$$v_a = \sum_{\ell=1}^{p_v} a_\ell V_\ell, \qquad g_c = \sum_{\ell=1}^{p_g} c_\ell H_\ell, \qquad I_\theta[\rho] = \sum_{\ell=1}^{p_I} \theta_\ell F_\ell[\rho]. \tag{S33}$$

Here $v_a$ represents the intrinsic-context probability-flow velocity. Collect the three coefficient vectors as $\beta = (a^\top, c^\top, \theta^\top)^\top \in \mathbb{R}^p$, where $p = p_v + p_g + p_I$ and $\|\beta\|_2 \le R$. The functions may depend nonlinearly on state, time and measured LR or neighborhood features, and $F_\ell[\rho]$ can depend on the population distribution. Assume that these functions represent the true dynamics with coefficients $\beta^* = (a^{*\top}, c^{*\top}, \theta^{*\top})^\top$.

To relate these coefficients to observations, choose smooth functions $\psi_j(t, x)$, $j = 1, \dots, J$, vanishing at $t = 0, T$. Each function can weight a region of state space over a chosen time interval. Multiplying (S31) by $\psi_j$ and integrating by parts gives $A(\rho)\beta = y(\rho)$, with $A = [A_v\ A_g\ A_I]$ and

$$(A_v)_{j\ell}(\rho) = \int_0^T \langle V_\ell \cdot \nabla\psi_j, \rho_t\rangle \, \mathrm{d}t,$$

$$(A_g)_{j\ell}(\rho) = \int_0^T \langle H_\ell \psi_j, \rho_t\rangle \, \mathrm{d}t,$$

$$(A_I)_{j\ell}(\rho) = \int_0^T \langle F_\ell[\rho_t] \cdot \nabla\psi_j, \rho_t\rangle \, \mathrm{d}t, \tag{S34}$$

$$y_j(\rho) = -\int_0^T \langle \partial_t \psi_j, \rho_t\rangle \, \mathrm{d}t. \tag{S35}$$

Here $y(\rho)$ records changes in the $\psi_j$-weighted cell counts, and each column of $A(\rho)$ gives the contribution of one specified function. Because all derivatives act on $\psi_j$, the relation can be evaluated without differentiating estimated densities. The true dynamics satisfy $A(\mu)\beta^* = y(\mu)$.

To determine the strengths separately, require

$$\|A(\mu)u\|_2 \ge \kappa\|u\|_2 \quad \text{for all } u \in \mathbb{R}^p, \qquad \kappa > 0. \tag{S36}$$

For the fixed matrix $A(\mu)$, this is equivalent to full column rank. Its smallest singular value gives a fixed $\kappa > 0$ for the chosen feature and test-function scales. Full column rank guarantees uniqueness, and the size of $\kappa$ controls how errors are amplified. Changing the coefficients must change at least one of the population measurements, even when all three components are adjusted together. Two coefficient vectors explaining the same complete path would satisfy $A(\mu)(\beta_1 - \beta_2) = 0$, so (S36) implies $\beta_1 = \beta_2$.

**Error in the estimated components.** We next relate errors in the distribution to errors in these strengths. Assume the integrands in (S34)–(S35) have uniform spatial $C^2$ bounds for paths with the mass bound above, and $F_\ell$ are Lipschitz from $d$ to the uniform norm. Adding and subtracting integrals with the same interaction input gives

$$\|A(\rho^n) - A(\mu)\|_{\mathrm{op}} \le L_A\delta_n, \qquad \|y(\rho^n) - y(\mu)\|_2 \le L_y\delta_n. \tag{S37}$$

Let $\widehat{A}_n, \widehat{y}_n$ be their numerical estimates, with errors bounded by $q_{A,n}, q_{y,n}$. These include errors in integration and input features. For estimated coefficients $\|\widehat{\beta}_n\|_2 \le R$, let $r_n = \|\widehat{A}_n\widehat{\beta}_n - \widehat{y}_n\|_2$ measure the error in satisfying the integrated population equation.

**Theorem S3.2** (Recovery of the true components). *Under* (S33)–(S37) *and the preceding assumptions,*

$$\|\widehat{\beta}_n - \beta^*\|_2 \le \frac{r_n + (L_y + RL_A)\delta_n + q_{y,n} + Rq_{A,n}}{\kappa}. \tag{S38}$$

*If the basis functions are bounded, define*

$$\|f\|_\mu^2 = \int_0^T \int_X |f|^2 \, \mathrm{d}\mu_t \, \mathrm{d}t, \qquad \mathcal{E}_n^2 = \|v_{\widehat{a}_n} - v_{a^*}\|_\mu^2 + \|g_{\widehat{c}_n} - g_{c^*}\|_\mu^2 + \|I_{\widehat{\theta}_n}[\rho^n] - I_{\theta^*}[\mu]\|_\mu^2.$$

*Then* $\mathcal{E}_n \le C_0\|\widehat{\beta}_n - \beta^*\|_2 + C_1\delta_n$*. With the model class and all bounds fixed, including* $\kappa > 0$*, suppose*

$$h_n,\ \varepsilon_n,\ \tau_n,\ r_n,\ q_{A,n},\ q_{y,n} \longrightarrow 0.$$

*Proposition S3.1 gives* $\delta_n \to 0$*, and* (S38) *then gives*

$$\widehat{\beta}_n \longrightarrow \beta^*, \qquad \mathcal{E}_n \longrightarrow 0.$$

*Thus the coefficients converge to the true coefficients, and the estimated intrinsic-context velocity, growth and interaction converge to the true fields in* $L^2(\mu)$*.*

*Proof.* Subtract the true and estimated population equations:

$$\begin{aligned} A(\mu)(\widehat{\beta}_n - \beta^*) = [A(\mu) - \widehat{A}_n]\widehat{\beta}_n + [\widehat{A}_n\widehat{\beta}_n - \widehat{y}_n] \\ + [\widehat{y}_n - y(\mu)]. \end{aligned}$$

The three terms have norms at most $R(L_A\delta_n + q_{A,n})$, $r_n$ and $L_y\delta_n + q_{y,n}$. Condition (S36) bounds the left-hand norm below by $\kappa\|\widehat{\beta}_n - \beta^*\|_2$, proving (S38). Velocity and growth are linear combinations of bounded functions, so their errors are bounded by the coefficient error. The interaction error also includes the change in its input distribution, which is controlled by the Lipschitz assumption. □

For example, minimizing $\|\widehat{A}_n\beta - \widehat{y}_n\|_2^2$ over $\|\beta\|_2 \le R$ gives $r_n \le (L_y + RL_A)\delta_n + q_{y,n} + Rq_{A,n}$ by comparison with $\beta^*$. Its coefficients therefore converge to $\beta^*$ as the errors decrease. The same conclusion holds for any estimator in this class for which $r_n \to 0$.

**Implications for the data.** Closer observation times and more accurate population measurements reduce the distribution error in (S32). Separating the components also requires relevant LR and neighborhood features to be measured where they act. Variation in neighborhoods among comparable receiver states can help distinguish their effects, which must satisfy (S36). This condition ensures that different combinations of the three components produce distinguishable changes in the population. The theorem applies to the model in (S33). For the nonlinear CytoBridge networks, we compare predictions with withheld stages (Supplementary Figs. S7 and S8) and with known simulation dynamics (Fig. 2 and Supplementary Figs. S2, S3 and S4), and examine repeated training and alternative settings (Supplementary Figs. S10 and S9).

### Supplementary Note S4. Additional implementation details

#### S4.1 Dataset sampling and inputs

For chicken heart [13], D4, D7, D10, and D14 denote embryonic days 4, 7, 10, and 14. We selected one tissue section per stage before alignment, retaining 147 of 747, 528 of 1,966, 908 of 1,916, and 1,967 of 1,967 input spots, respectively. Spatial clustering separated the tissue sections, and one section per stage was selected by its position on the slide before model fitting. Raw counts were recovered from the corresponding GSE149457 matrices.

For ARTISTA [14], the source study produced the injury by surgically removing a reproducible portion of the dorsal pallium in the left telencephalic hemisphere. We used one annotated section at each of 2, 5, 10, 15, and 20 days post injury; after removing isolated spatial outliers within each section, the input contained 46,199 cells.

For MOSTA [15], we used one representative section at each of E12.5, E13.5, E14.5, and E15.5, comprising 344,603 spatial observations. For the AD mouse dataset [16], we analyzed one TgCRND8 coronal section at each of 2.5, 5.7, and 17.9 months, with 172,092 cells in total. For zebrafish [17], we used one section at each of 5.25, 10, 12, 18, and 24 hours post fertilization (hpf), comprising 563, 1,036, 2,081, 3,048, and 5,271 observations, respectively, and 11,999 observations in total.

##### S4.2 Embedding, preprocessing, and scaling

Transcriptomic embeddings were represented by the first 50 principal components of log-normalized gene expression and concatenated with 2 spatial coordinates, yielding a 52-dimensional joint state vector. Time was normalized as described in Supplementary Note S1.2.

##### S4.3 Network architectures

The probability-flow drift $\mathbf{v}_\theta(\mathbf{z}, t)$ was parameterized by a decoupled residual MLP with 5 hidden layers, hidden dimension 256, and LeakyReLU activations, with output dimension equal to the full 52-dimensional state. The growth field $g_\theta(\mathbf{z}, t)$ was modeled by a scalar-output MLP on the same joint state-time input. The interaction operator $\mathcal{I}_\theta$ was implemented as an attention-based graph neural network on dynamic spatial interaction graphs with 8 attention heads and hidden dimension 256, together with a pre-trained edge predictor for graph pruning. The scalar score potential $\phi_\theta(\mathbf{z}, t)$ was parameterized by an MLP with hidden dimension 400 for score matching in the same 52-dimensional joint state space.

##### S4.4 Optimization of the real-data models

The real-data models were fitted in six consecutive stages, with interactions introduced in the third stage and the score fitted before and after joint fine-tuning:

| Stage | Updated fields | Epochs |
|---|---|---|
| Pretrain | $\mathbf{v}_\theta, g_\theta$ | 100 |
| Refine | $\mathbf{v}_\theta, g_\theta$ | 100 |
| Init interaction | $\mathbf{v}_\theta, g_\theta, \mathcal{I}_\theta$ | 50 |
| Train score | $\mathbf{s}_\theta$ | 2,001 |
| Finetune | $\mathbf{v}_\theta, g_\theta, \mathcal{I}_\theta$ | 1,000 |
| Score refine | $\mathbf{s}_\theta$ | 2,001 |

Pretrain and Refine used the snapshot-matching objective with $(\lambda_{\text{OT}}, \lambda_{\text{mass}}, \lambda_{\text{energy}}) = (1, 0.01, 0)$ and learning rate $10^{-4}$. During Init interaction, the standard settings were $(\lambda_{\text{OT}}, \lambda_{\text{mass}}, \lambda_{\text{energy}}) = (10, 10, 0.01)$. Train score and Score refine used score matching with AdamW and initial learning rate $10^{-4}$. The learning rate remained constant in the zebrafish, MOSTA, ARTISTA, and chicken-heart score stages; AD mouse used cosine learning-rate decay with $T_{\max} = 1{,}000$ and $\eta_{\min} = 10^{-5}$. Finetune used snapshot matching with the fitted score field enabled, $(\lambda_{\text{OT}}, \lambda_{\text{mass}}) = (10, 10)$ ($(1, 1)$ for ARTISTA; Supplementary Table 2), $\lambda_{\text{energy}} = 0.01$, and initial learning rate $10^{-4}$. Zebrafish, MOSTA, ARTISTA, and chicken heart used a learning rate that was halved after 50 epochs without improvement during Finetune, monitored on the forward last-interval OT loss. AD mouse used cosine learning-rate decay with $T_{\max} = 1{,}000$ and $\eta_{\min} = 10^{-7}$. The OT and mass coefficients, batch sizes, and graph thresholds are listed in Supplementary Table 2.

We recorded the objective once per epoch. Supplementary Fig. S43 shows the ARTISTA training history, with recorded values and centered moving means for each stage. The score stages optimize score matching, whereas the remaining stages optimize their respective composite losses, so each curve is shown on its own scale.

#### S4.5 Dataset-specific hyperparameters

Supplementary Table 2 lists the values used for the five applications. Let $d_t$ be the median distance from each observation to its nearest neighbor at observed time $t$. We estimated the spot diameter at that time as $1.2d_t$ and set the dataset-level spatial cutoff to four times the mean estimated diameter, $4\langle 1.2d_t \rangle_t$. We trained the edge predictor on 90% of the labelled edges and selected its probability threshold on the remaining 10%. The threshold was selected by maximizing the validation F1 score. The score-matching batch size was 128 for datasets with fewer than 100,000 observations, 256 for 100,000–249,999 observations, and 512 for datasets with at least 250,000 observations.

#### S4.6 Sensitivity to training-loss weights

We examined two training-loss choices in the zebrafish dataset, training one model per setting with the same seed on all measured stages. For the expression-loss analysis, we independently trained models with $\alpha_{\text{expr}} = 0.015$ or 0.05. For the transport-to-mass analysis, we fixed $\alpha_{\text{expr}} = 0.015$ and compared $\lambda_{\text{OT}} : \lambda_{\text{mass}} = 10 : 1$, $1 : 1$, and $1 : 10$ in the interaction stages while holding $\lambda_{\text{OT}} + \lambda_{\text{mass}} = 20$. Generated distributions were compared with the observed snapshots at $t = 1, 2, 3$, and 4 using Wasserstein-1 distance (W1) in the joint 52-dimensional state, the 50-dimensional expression state, and the two-dimensional physical space (Supplementary Fig. S44).

Increasing $\alpha_{\text{expr}}$ from 0.015 to 0.05 reduced mean joint-state and expression-state W1 by 8.7% and 8.4%, respectively, but increased mean physical-space W1 by 25.3%. The 10:1 transport-to-mass setting increased mean W1 relative to 1:1 by 1.1% in the joint state, 1.0% in expression space, and 3.7% in physical space. The 1:10 transport-to-mass setting reduced physical-space W1 by 2.0% but increased joint-state and expression-state W1 by 7.6% and 7.8%. We therefore retained the 1:1 interaction-stage ratio as the setting with the best balance between expression and spatial reconstruction.

#### S4.7 Stability of the interaction decomposition in zebrafish

We evaluated the interaction and intrinsic-context decomposition in the zebrafish dataset under independent training and changes to selected model settings (Supplementary Fig. S10). Five default models trained with different random seeds were evaluated on the same 11,999 observed cells at five model times and compared in all ten pairs. Agreement was measured using median cellwise cosine similarity for the total dynamics and intrinsic-context field in the 50-dimensional expression-state space, Spearman correlation for growth, and Spearman correlation between directed cell-type-pair rankings calculated from exact directed-edge messages. The median agreements across these ten pairs were 0.899 for total dynamics, 0.931 for the intrinsic-context field, 0.870 for growth, and 0.980 for the directed cell-type interaction ranking.

We next changed one model setting at a time. The spatial-neighborhood cutoff was set to 0.8 or 1.2 times its default value in three matched training seeds. An expression-loss weight of 0.05 and a transport-to-mass ratio of 10:1 were each evaluated using one training seed. Compared with the default model trained with the same seed, we evaluated cosine similarity of the expression-space interaction vectors, Spearman correlation of sender–receiver pair rankings (0.963–0.997 across the tested settings), and weighted overlap among the highest-ranked 20% of pairs with nonzero contributions.

Reconstruction remained similar across these comparisons. Across the five independently trained default models, the coefficients of variation of mean spatial, expression-state, and joint Wasserstein-1 distances were 4.11%, 0.90%, and 0.98%, respectively. After changing one model setting, the spatial and expression-state distances changed by -1.2% to 3.7% relative to the matched default.

#### S4.8 Interaction ablations and method comparisons

Both ablations below use models trained on all measured stages and compare reconstructions of these stages. We tested the contribution of the ligand–receptor (LR) prior by training models with and without it in five spatial datasets. The Full model used the trained edge predictor to select LR-compatible neighbors. The No-LR-prior model retained the interaction GNN and spatial cutoff but connected all cells within that distance. All other settings, including the training seed, were identical. Reconstruction from the initial population to later stages was measured in joint, gene-expression, and physical space. For each target stage and evaluation space, we averaged sliced-W2 over five projection repeats and divided the No-LR-prior error by the Full-model error. These ratios were averaged with equal weight within each dataset (Supplementary Fig. S9a,b).

Removing the LR-informed gate increased mean sliced-W2 by 13.5% in zebrafish, 26.1% in MOSTA, 177.3% in ARTISTA, 25.5% in AD mouse, and 1.8% in chicken heart. The increase occurred in all 12 ARTISTA target-by-space comparisons. The chicken-heart effect was modest and target dependent: error decreased slightly at the first two targets and increased at the third in all three spaces.

We also tested the contribution of the learned interaction module during inference (Supplementary Fig. S9c,d). For each dataset, we generated predictions from the trained Full model with and without its interaction-network output, starting from the same 5,000 particles. Growth was retained through continuous, unnormalized particle weights without resampling. The two conditions used paired random streams for three inference seeds, a time step of 0.01, and a diffusion scale of 0.03. Each population was propagated continuously from the earliest source to all later observed stages. Weighted sliced-W2 used all predicted and observed particles in the same standardized joint, spatial, and gene-state coordinates. For each inference seed, target stage, and evaluation space, errors were averaged over the same five sets of 1,024 projection directions, and the ratio of the error without interaction to the error with interaction was averaged across the three paired seeds. These target–space ratios were then averaged with equal weight within each dataset. Disabling interaction increased mean sliced-W2 by 10.4% in zebrafish, 2.5% in MOSTA, 30.1% in ARTISTA, 2.0% in AD mouse, and 97.9% in chicken heart, with higher error in 36 of the 48 target–space comparisons and lower error in 12.

For the external comparison, CytoBridge and stVCR [18] were fitted separately with each target stage omitted. Five projection repeats were averaged within target and space before calculating paired ratios (Supplementary Fig. S8a). We used the stVCR predictions with growth enabled and assigned each output particle a mass of 1/5,000. Sliced-W2 normalized these weights and used all predicted and observed particles at their original counts, whereas total-mass variation used the unnormalized output mass. CytoBridge had lower sliced-W2 in 28 of 33 comparisons across target stages and evaluation spaces, and in 11 of 15 comparisons after averaging over target stages within each dataset, including physical space in all five datasets. stVCR had slightly lower mean errors in the joint and gene-state spaces of MOSTA and ARTISTA.

We extended this leave-one-time-point-out evaluation to MOSCOT [19], PASTE2 [20, 21], Spateo [22], Linear OT displacement and random-interpolation controls, MIOFlow 2.0 [23, 24], STORIES [25], Waddington-OT [26], and PCA-based SpaTrack [27] (Supplementary Figs. S7 and S8). Each method started from the same 5,000 initial particles selected without using the held-out target, and all methods except stVCR returned 5,000 predicted particles. Because MIOFlow 2.0, STORIES, and Waddington-OT do not output spatial coordinates, they were evaluated in gene-state space only; MIOFlow 2.0 used a joint embedding of expression PCs and spatial-neighborhood features with stochastic dynamics and a growth model. PASTE2 aligned adjacent observed stages with the overlap fraction set to 1, and SpaTrack used Euclidean PCA distances in place of its default expression-based

KL cost, with both methods predicting the held-out stage by barycentric interpolation through the fitted coupling. Finally, the Linear OT control interpolated between observed populations through a fitted transport coupling, whereas the random-interpolation control paired each cell at the preceding stage with a uniformly sampled cell from the following stage with replacement before linear interpolation.

For every matched dataset, held-out target, method, and available evaluation space, Sliced-W2 used 1,024 projection directions per repeat, with the same directions used for all methods evaluated at that target stage and in that space. For each target stage and evaluation space, we averaged the five repeats and divided each comparison method's error by the CytoBridge error. These ratios contributed equally to the dataset mean. CytoBridge had lower Sliced-W2 in 28 of 33 matched comparisons for stVCR, 27 of 33 for MOSCOT, 23 of 33 for PASTE2, 32 of 33 for Spateo, 29 of 33 for random interpolation, 30 of 33 for SpaTrack, 11 of 11 for MIOFlow 2.0, and 9 of 11 for each of STORIES and Waddington-OT. Across the 33 combinations of held-out stage and evaluation space, CytoBridge had the lowest Sliced-W2 in 12, more than any other method or control, while providing interaction-resolved continuous dynamics not modeled by the other methods compared here.

##### S4.9 Computational cost

We measured training time, peak host memory, and peak GPU allocation for one full-model run per dataset on an NVIDIA GeForce RTX 4090 D (Supplementary Table 3). The six training stages took 11.6–33.3 min, excluding preprocessing and downstream evaluation.

#### Supplementary Note S5. Atlas-grounded in silico spatiotemporal benchmark

To evaluate recovery of continuous spatiotemporal dynamics beyond snapshot matching, we constructed an atlas-grounded in silico spatiotemporal (AGIST) benchmark using a CytoBridge model trained on the MOSTA mouse organogenesis atlas. We simulated the evolution of an initial population of brain cells and recorded the generator's velocity, growth and attention on the resulting snapshots.

For method-level comparison on this benchmark, we included STORIES and stVCR as representative OT-based baselines for reconstructing developmental dynamics from snapshot data [18, 25].

##### S5.1 Generative simulation from an atlas-trained model

We started from the CytoBridge model trained on the four-stage MOSTA dataset. MOSTA expression was normalized and log-transformed once from the count layer before selecting the 50-dimensional gene state. We sampled 5,000 Brain-annotated cells from E12.5 without replacement and retained their two aligned spatial coordinates and 50 gene-state coordinates.

The split-SDE framework in Supplementary Note S8.2 was used with a time step of 0.01, diffusion coefficient 0.03 and interaction groups of 1,024 cells. Growth-dependent division and extinction were applied at $t = 1, 2, 3$, with zero daughter displacement. We recorded the exact initial population at $t = 0$ and retained all descendants, yielding 5,000, 7,637, 9,733 and 10,667 cells at the four snapshots.

The atlas-trained generator was used to evaluate the ground-truth velocity components, growth field $g_t^\star(\mathbf{z})$ and attention matrix $A_t^\star$ on the simulated cells. We trained a new CytoBridge model using the four snapshots, without supplying these ground-truth fields. The new model used the same spatial cutoff and fixed ligand–receptor edge classifier as the generator. Training followed the six-stage schedule and epoch counts in Supplementary Note S4.4. We compared the recovered fields

$$\left(\widehat{\mathbf{v}}_t, \widehat{\mathcal{I}}_t, \widehat{g}_t, \widehat{A}_t\right)$$

against the corresponding generator values.

##### S5.2 Metrics

We evaluated recovery of velocity, interaction, growth, and population distributions in AGIST. For velocity recovery in Fig. 2b and Supplementary Fig. S3, we evaluated the complete field defined in Eq. (S40). Let $\widehat{\mathbf{q}}_t(\mathbf{z})$ denote a predicted vector field and $\mathbf{q}_t^\star(\mathbf{z})$ the corresponding generator field, and let $P_{\mathrm{sp}}$ and $P_{\mathrm{ge}}$ denote projection onto physical space and gene-expression space, respectively. We quantified directional agreement by

$$\mathrm{Sim}_P(\widehat{\mathbf{q}}, \mathbf{q}^\star) = \frac{1}{\sum_k n_k} \sum_k \sum_{i=1}^{n_k} \frac{\left\langle P\widehat{\mathbf{q}}_{t_k}(\mathbf{z}_i^{(k)}), P\mathbf{q}_{t_k}^\star(\mathbf{z}_i^{(k)}) \right\rangle}{\left\| P\widehat{\mathbf{q}}_{t_k}(\mathbf{z}_i^{(k)}) \right\| \left\| P\mathbf{q}_{t_k}^\star(\mathbf{z}_i^{(k)}) \right\|}, \qquad P \in \{P_{\mathrm{sp}}, P_{\mathrm{ge}}\}.$$

This score is reported separately in physical space and in gene-expression space, thereby assessing whether the inferred flows recover the generator-implied directions in both domains.

To assess recovery of interaction structure, we summarized each time-varying attention matrix by the outgoing interaction strength of each cell,

$$s_i(t_k) := \frac{1}{n_k} \sum_{j=1}^{n_k} A_{ij}(t_k), \qquad \widehat{s}_i(t_k) := \frac{1}{n_k} \sum_{j=1}^{n_k} \widehat{A}_{ij}(t_k).$$

At each benchmark time point, we computed the Spearman correlation between $\{\widehat{s}_i(t_k)\}_{i=1}^{n_k}$ and $\{s_i(t_k)\}_{i=1}^{n_k}$ over cells with nonzero outgoing strength in at least one of the two matrices. The four correlations were 0.782, 0.800, 0.798 and 0.805, with a mean of 0.796. Figure 2c shows the $t = 0$ comparison. Growth recovery was measured by the Pearson correlation between predicted and generator growth values across all benchmark cells and times,

$$\mathrm{Corr}_g = \mathrm{corr}\Big(\{\widehat{g}_{t_k}(\mathbf{z}_i^{(k)})\}_{k,i}, \{g_{t_k}^\star(\mathbf{z}_i^{(k)})\}_{k,i}\Big).$$

This measures the linear association between inferred and generator growth rates across cells and time points.

For distributional recovery, we quantified the discrepancy between the predicted particles and the benchmark snapshot by a weighted Wasserstein-2 distance on projection $P$,

$$W_{2,P}\big(\widehat{\mu}_{t_k}, \mu_{t_k}^\star\big) := \left[ \min_{\pi \in \Pi(\mathbf{w}^{(k)}, \mathbf{a}^{(k)})} \sum_{j=1}^{\widehat{n}_k} \sum_{i=1}^{n_k} \pi_{ji} \left\| P\widehat{\mathbf{z}}_j^{(k)} - P\mathbf{z}_i^{(k)} \right\|^2 \right]^{1/2}, \tag{S39}$$

where $\mathbf{w}^{(k)} = (w_1^{(k)}, \ldots, w_{\widehat{n}_k}^{(k)})$ denotes the predicted particle weights normalized to sum to one at time $t_k$ and $\mathbf{a}^{(k)} = (1/n_k, \ldots, 1/n_k)$ denotes the empirical benchmark weights. We evaluated (S39) in physical and gene-expression space. These weighted OT quantities can be solved with standard software such as POT [28]. Methods that do not reconstruct spatial coordinates were evaluated only in gene-expression space. The change in total population size was assessed separately through the mass metric below. Finally, to quantify recovery of population-size dynamics under the unbalanced formulation, we defined the relative mass error at time $t_k$ by

$$\mathrm{Err}_{\mathrm{mass}}(t_k) = \frac{|\widehat{m}_k - m_k^\star|}{m_k^\star}, \qquad \widehat{m}_k := \frac{\sum_j w_j(t_k)}{\sum_j w_j(t_0)}, \qquad m_k^\star := \frac{n_k}{n_0},$$

where $\widehat{m}_k$ and $m_k^\star$ denote the predicted and benchmark relative masses, respectively. For Fig. 2e, each method was trained once on the same four snapshots. The official stVCR implementation used 2,000 epochs, 500 sampled cells per stage, growth enabled and alignment disabled. STORIES used 100 training iterations and batches of 128 cells, with spatial coordinates and the same 50 gene-state coordinates. Predictions started from the same 5,000 initial cells and continued to $t = 3$. For CytoBridge and stVCR, we kept the trained model and initial cells fixed and repeated prediction ten times with different random seeds, using a time step of 0.01. CytoBridge simulation includes diffusion, stochastic cell division and death, and random grouping of cells for interaction calculations. The stVCR velocity integration is deterministic, but its official population simulator samples cell division and death according to the predicted growth rates and adds a small perturbation to daughter-cell states. We used all predicted cells to calculate exact empirical Wasserstein-2 distance, with uniform weights normalized within each population. STORIES supplied one deterministic gene-state prediction per target time and retained 5,000 particles. In Fig. 2e, each open circle is the Wasserstein-2 distance for one predicted population at the indicated time. Bars and error bars show the mean and standard deviation across the ten predictions. Two-sided sign-flip tests compared the ten prediction errors from CytoBridge with those from stVCR using corresponding seeds and with the fixed STORIES prediction, with Holm correction across nine comparisons.

Figure 2d displays each growth distribution after clipping at its first and 99th percentiles and scaling to $[0, 1]$. The raw and displayed growth correlations were 0.923 and 0.925, respectively. The spatial maps in Fig. 2b,c and Supplementary Fig. S2 show the main tissue region. All cells were included in the numerical evaluations.

##### S5.3 Velocity agreement across time and state-space partitions

For the stratified analysis, we evaluated the total drift of the stochastic dynamics,

$$\widehat{\mathbf{q}}_t = \widehat{\mathbf{v}}_t + \widehat{\mathcal{I}}_t + \widehat{\mathbf{c}}_t, \qquad \mathbf{q}_t^\star = \mathbf{v}_t^\star + \mathcal{I}_t^\star + \mathbf{c}_t^\star, \tag{S40}$$

where $\mathbf{c}_t$ denotes the score drift defined in Eq. (S27). Cellwise cosine similarity was calculated separately over the two physical coordinates and the 50-dimensional gene state. We summarized agreement by the median cellwise cosine similarity at each of the four simulated time points and within each expression-state cluster.

We grouped the 33,037 cells from all four benchmark snapshots by MiniBatch $k$-means in the 50-dimensional gene-state space. Three clusters were selected by the highest silhouette score among $k = 3, \ldots, 8$, evaluated on 8,000 cells sampled across time points. The same cluster centroids were used at every time point.

Across the four time points, median velocity cosine similarity was at least 0.996 in physical space and 0.972 in gene-expression space. Across the three state-space partitions, the corresponding medians ranged from 0.996 to 0.998 and from 0.976 to 0.990, respectively (Supplementary Fig. S3).

### Supplementary Note S6. Synthetic benchmark with finite-range cell–cell attraction

We constructed a controlled interacting-particle benchmark to test whether CytoBridge could recover a finite-range attractive interaction while simultaneously reconstructing motion in physical and gene-expression space and changes in population mass (Supplementary Fig. S4). The low-dimensional simulator specified the interaction law and the other dynamical components. The joint state of cell $i$ was $\mathbf{z}_i = (\mathbf{x}_i, \mathbf{r}_i) \in \mathbb{R}^2 \times \mathbb{R}^2$, where $\mathbf{r}_i$ denotes physical position and $\mathbf{x}_i$ denotes a two-dimensional expression state.

#### S6.1 Ground-truth dynamics and snapshot generation

The ground-truth particle system combined a translational spatial drift, relaxation in expression space, finite-range attraction, diffusion and uniform net population growth:

$$\begin{aligned} \mathrm{d}\mathbf{r}_i(t) &= [\mathbf{v}_r + \mathbf{F}_i(t)]\,\mathrm{d}t + \sigma_r\,\mathrm{d}\mathbf{B}_i^{(r)}(t), \\ \mathrm{d}\mathbf{x}_i(t) &= [-\kappa\mathbf{x}_i(t) + \Gamma\mathbf{F}_i(t)]\,\mathrm{d}t + \sigma_x\,\mathrm{d}\mathbf{B}_i^{(x)}(t), \\ \mathrm{d}\log m_i(t) &= g^\star\,\mathrm{d}t, \end{aligned}$$

with parameters defined in the table below. The Brownian motions $\mathbf{B}_i^{(r)}$ and $\mathbf{B}_i^{(x)}$ were independent. The interaction force was

$$\mathbf{F}_i(t) = \frac{1}{N-1}\sum_{j\neq i} \widetilde{m}_j(t)\,\phi^\star(r_{ij})\frac{\mathbf{r}_j(t)-\mathbf{r}_i(t)}{r_{ij}}, \qquad r_{ij} = \|\mathbf{r}_j(t)-\mathbf{r}_i(t)\|, \tag{S41}$$

where $N$ is the number of simulated particles, $\widetilde{m}_j = m_j/(N^{-1}\sum_k m_k)$, and

$$\phi^\star(r) = \begin{cases} A\sin(\pi r/r_c), & 0 < r < r_c, \\ 0, & \text{otherwise.} \end{cases} \tag{S42}$$

Because the unit vector in Eq. (S41) points from cell $i$ to cell $j$, $\phi^\star(r) > 0$ denotes attraction. Initial spatial positions were drawn from an equal mixture of isotropic Gaussians centered on $\mathcal{C}_r$, and initial expression states were drawn from an equal mixture centered on $\mathcal{C}_x$. We denote the corresponding within-component standard deviations by $\tau_r$ and $\tau_x$. At each observation time, cells were sampled in proportion to their particle masses and independent Gaussian measurement errors with standard deviations $\eta_r$ and $\eta_x$ were added to the two coordinate blocks.

The numerical values used to generate the benchmark were as follows:

| Symbol | Value | Definition |
|---|---|---|
| $t_k$ | $0, 1, 2, 3, 4$ | observation times |
| $N_0$ | 400 | initial number of particles |
| $\mathbf{v}_r$ | $(0.02, 0.01)^\mathsf{T}$ | background spatial velocity |
| $\kappa$ | 0.5 | expression-relaxation rate |
| $A$ | 0.5 | attraction amplitude |
| $r_c$ | 0.30 | interaction cutoff |
| $\Gamma$ | $3I_2$ | spatial-to-expression interaction map |
| $\sigma_r, \sigma_x$ | $0.015, 0.015$ | diffusion amplitudes |
| $g^\star$ | $\log(1.25)/4$ | log-mass growth rate |
| $\Delta t$ | 0.02 | Euler–Maruyama step size |
| $\mathcal{C}_r$ | $\{0.2, 0.4, 0.6, 0.8\}^2$ | spatial mixture centers |
| $\tau_r$ | 0.03 | initial spatial spread |
| $\mathcal{C}_x$ | $\{(\pm 0.25, \pm 0.25)\}$ | expression mixture centers |
| $\tau_x$ | 0.05 | initial expression spread |
| $\eta_r, \eta_x$ | $0.005, 0.01$ | measurement-noise scales |

The resulting snapshot sizes were 400, 423, 447, 473 and 500 cells.

#### S6.2 Benchmark-specific model and training configuration

The benchmark retained the CytoBridge dynamical decomposition. The low-dimensional network sizes, interaction head and stage-specific objectives were set for this controlled system as described below.

**Interaction head.** Within the interaction operator, the general attention readout in Eq. (S29) was replaced by a radial interaction function that depends only on the distance between cells to match the rotational symmetry of the data-generating interaction:

$$\mathbf{F}_i^{\psi} = \frac{1}{N-1} \sum_{j \neq i} \widetilde{m}_j \, c_\psi(r_{ij}) \frac{\mathbf{r}_j - \mathbf{r}_i}{r_{ij}}, \qquad \mathcal{I}_\psi(\mathbf{z}_i) = \big(\Gamma \mathbf{F}_i^{\psi}, \mathbf{F}_i^{\psi}\big).$$

Pair distance was encoded by eight fixed Gaussian radial basis functions with centers spanning $[0, r_c]$. A neural network with two hidden layers of 400 units and LeakyReLU activations mapped these distance features to the signed coefficient $c_\psi(r)$. The radial coefficient was fitted from the observed snapshots and could take either sign. Messages were restricted to $r < r_c$, weighted by source-cell mass and averaged over the training group. The fixed map $\Gamma = 3I_2$ transferred the inferred radial force to expression space, as in the simulator.

The probability-flow and growth networks were two-layer multilayer perceptrons with hidden dimension 400 and ReLU activations. The score network contained four hidden layers with dimension 128 and LeakyReLU activations. The interaction graph and training mini-batches both used groups of 192 cells.

**Training settings.** Training followed Supplementary Note S4.4, with the same stage sequence, epoch counts, and Pretrain and Refine settings. Snapshot matching used a detached Sinkhorn loss. For the remaining stages, the settings differed as follows. Interaction initialization used $\lambda_{\mathrm{OT}} = 1$ and learning rate $10^{-5}$. The two score stages used flow matching with penalty weight 1, learning rate $10^{-3}$ and cosine decay to $10^{-5}$. Joint fine-tuning used $(\lambda_{\mathrm{OT}}, \lambda_{\mathrm{mass}}, \lambda_{\mathrm{energy}}) = (10, 10, 0.01)$ and learning rate $10^{-5}$. Because the two spatial and two expression coordinates were generated on comparable numerical scales, their loss weights were set to $\alpha_{\mathrm{spatial}} = 1$ and $\alpha_{\mathrm{expr}} = 1$. The diffusion scale was $\sigma = 0.015$. The model with the lowest stage-specific objective was retained.

##### S6.3 Evaluation of dynamics, growth and interaction recovery

Coordinate dynamics were evaluated by following the same 400 particles from the initial ground-truth population. Particle masses were held uniform and growth-dependent resampling was disabled, separating coordinate reconstruction from population growth. Ground-truth and inferred particles were propagated with $\Delta t = 0.02$ and $\sigma = 0.015$. We computed Wasserstein-1 and Wasserstein-2 distances separately in physical space, expression space and the joint four-dimensional state space at $t = 1, 2, 3, 4$. Stochastic inference was repeated five times with different random seeds.

Growth was evaluated separately by comparing the inferred relative total mass with the ground-truth mass ratio $M(t)/M(0) = \exp(g^\star t)$. For time $t_k$, total-mass variation was defined as

$$\mathrm{TMV}(t_k) = \left| \frac{\widehat{M}(t_k)}{\widehat{M}(0)} - \frac{M^\star(t_k)}{M^\star(0)} \right| .$$

The learned radial coefficient was evaluated directly over $0 < r < 0.30$ and compared with Eq. (S42) using Pearson correlation, Spearman correlation and sign agreement. For visualization only, the corresponding interaction potential was obtained by post-hoc integration,

$$U(r) = -\int_r^{r_c} c(s)\,\mathrm{d}s, \qquad r_c = 0.30,$$

which fixes the additive gauge by setting $U(r_c) = 0$. Finally, the contribution of the inferred interaction was assessed using the same trained model and initial particles, either retaining $\mathcal{I}_\theta$ or setting $\mathcal{I}_\theta \equiv 0$ during forward simulation. All other model components and stochastic settings were unchanged.

### Supplementary Note S7. Non-spatial evaluation with lineage tracing and metabolic RNA labeling

We evaluated the non-spatial adaptation of CytoBridge using lineage tracing in the Weinreb/LARRY hematopoiesis dataset and newly transcribed RNA in the scNT-seq cortical KCl time course. These measurements were reserved for evaluation. In both datasets, the interaction graph and GNN operate in expression-state space because physical coordinates are unavailable.

#### S7.1 Lineage-traced hematopoiesis

##### S7.1.1 Weinreb/LARRY lineage-tracing data and preprocessing

In the Weinreb/LARRY mouse hematopoietic lineage-tracing dataset [29], heritable lentiviral barcodes link transcriptionally profiled cells to their clonal descendants, providing measurements of endpoint fate. The dataset contained 49,302 cells and 25,289 genes, comprising 4,638 cells at day 2, 14,985 cells at day 4, and 29,679 cells at day 6. A lineage identifier was defined by concatenating the starting population and clone barcode so that identical numerical clone labels from different starting populations were not merged.

The input expression matrix contained library-normalized linear values. We renormalized each cell to a total of 10,000, applied the $\log(1+x)$ transformation, selected 2,000 highly variable genes, and retained 50 principal components (PCs). Days 2, 4, and 6 were mapped to model times 0, 1, and 2; model fitting used only the expression states and observation times, whereas SPRING coordinates and lineage, cell-type, and fate annotations were used for visualization and evaluation.

##### S7.1.2 LR-informed non-spatial interaction graph

The model state was the complete 50-PC expression vector $z_i(t) \in \mathbb{R}^{50}$, with no physical-coordinate block. Candidate ordered cell pairs satisfied

$$0 < \|z_j - z_i\|_2 < c, \qquad c \approx 25.8,$$

where $c$ was the maximum, across the three observed times, of the within-time 99th percentile of pairwise PC distances.

For the directed ligand–receptor (LR) edge prior, we used the mouse CellChatDB distributed with COMMOT [30, 31]. We retained secreted-signaling entries for which every ligand or receptor complex subunit was detected in at least 50 cells. Complex activity was defined by the least-expressed required subunit. This yielded 193 LR pairs in 52 pathways. For a candidate sender $j$ and receiver $i$, the expression-derived compatibility score was

$$Q_{j\to i} = \mathrm{mean}_p \left[\mathrm{mean}_{(L,R)\in p}\, \widetilde{a}_{j,L}\widetilde{a}_{i,R}\right],$$

where activities were scaled by their training-cell 95th percentiles. The within-pathway mean followed by the between-pathway mean prevented pathways with more database entries from dominating the target.

Within each time point, the top 20% of the training-cell $Q$ distribution was labeled LR-compatible. We trained a multilayer perceptron to predict LR compatibility from paired expression states, with 70% of cells for training, 15% for validation, and 15% for testing. The predictor was then held fixed during dynamics training. Held-out AUROC and average precision for LR edge-label prediction were 0.79 and 0.50.

The interaction operator used the CytoBridge GNN to predict changes in the 50-dimensional expression state instead of the two-dimensional spatial coordinates. Cells were processed in groups of 16. Messages depended on the distance between cell states and were weighted by the predicted mass of the sender cell. The network used eight radial basis functions applied to distance divided by $c$, a 256-dimensional hidden state, eight attention heads, one message-passing layer, and a bias-free 50-dimensional readout. Reapplying the trained edge predictor at each step updates the graph as expression states evolve.

##### S7.1.3 Dynamical model, training schedule, and ablations

For particle $i$, the fitted stochastic dynamics were

$$\begin{aligned} \mathrm{d}z_i &= [v_\theta(t, z_i) + I_\eta(z_i, \{z_j, w_j\}) + \mathbf{c}_\theta(t, z_i)]\,\mathrm{d}t + \sigma\,\mathrm{d}W_i(t), \\ \mathrm{d}\log w_i &= g_\psi(t, z_i)\,\mathrm{d}t, \end{aligned}$$

where $v_\theta$ is the cell-autonomous state drift, $I_\eta$ is the GNN interaction contribution, $\mathbf{c}_\theta = \nabla_z \phi_\theta$ is the score drift defined in Eq. (S27), and $g_\psi$ controls particle mass. The velocity and growth networks contained five and three 256-unit hidden layers, respectively. The scalar score network contained three 400-unit hidden layers.

To evaluate the contribution of the interaction network, we trained three Full models and three No-interaction models using the six-stage procedure described in Supplementary Note S4.4. Across the four dynamical stages, the OT-loss weights were 1, 1, 10, and 1 and the mass-loss weights were 0.01, 0.01, 10, and 1. In both conditions, the energy penalty had a weight of 0.01 during the third training stage and joint fine-tuning. The learning rate was $10^{-4}$, neural-ODE and score batch sizes were 512 and 128, respectively, and $\sigma = 0.1$.

##### S7.1.4 Distribution and clone-fate evaluation

Each trained model, fitted to all three measured stages, was used to predict the day-4 and day-6 populations in 50 simulations with different random seeds. Within each simulation repeat, the two conditions used the same 2,048 sampled day-2 cells and Brownian noise, with Euler–Maruyama step size 0.05. Generated and observed day-4 and day-6 cells were compared in the complete, unwhitened 50-PC state using growth-weighted W1 and W2, with weights normalized for transport and at most 1,024 points per side. We averaged the errors first over simulations and then over the three models in each condition (Supplementary Figure S5c).

Clone-fate evaluation propagated all 4,638 day-2 cells together to day 6 with step size 0.1 in ten simulations per trained model. Scoring used the 1,714 source cells in 1,237 lineages with observed day-6 descendants. A uniform 20-nearest-neighbor classifier fitted to all 29,679 observed day-6 PC states assigned each endpoint to one of 11 fates. For each lineage, we summed the predicted endpoint weights for each fate over source cells and repeats, then averaged the resulting fate proportions across the three models in each condition before comparing them with those of its observed day-6 descendants.

We report total-variation agreement,

$$\mathrm{TV}_{\mathrm{agree}}(\ell) = 1 - \frac{1}{2}\sum_k \left|\widehat{p}_{\ell k} - p_{\ell k}^{\mathrm{obs}}\right|,$$

and one minus the normalized Jensen–Shannon divergence. Metrics were first averaged equally over evaluable lineages. Full versus No-interaction agreement was 0.480 versus 0.468 for total variation, 0.569 versus 0.560 for Jensen–Shannon similarity, and 0.476 versus 0.473 for dominant-fate matching (Supplementary Figure S5d).

##### S7.1.5 Model-derived interactions and cross-method consistency

For sender type $A$ and receiver type $B$, the day-6 network summarized the GNN message vectors as

$$D_{A\to B} = \frac{1}{N_B} \sum_{i:\text{type}(i)=B} \left\| \sum_{j:\text{type}(j)=A} \mathbf{c}_{j\to i} \right\|_2 .$$

Here $\mathbf{c}_{j\to i} \in \mathbb{R}^{50}$ denotes the instantaneous contribution of sender cell $j$ to the interaction drift of receiver cell $i$. Let $M_\eta$ denote the attention block, which takes the sender and receiver states and their distance as inputs. The contribution is

$$\mathbf{c}_{j\to i} = R\left[ \frac{m_j}{\sum_{\ell\in\mathcal{N}(i)} m_\ell} M_\eta(\mathbf{z}_j, \mathbf{z}_i, \|\mathbf{z}_j - \mathbf{z}_i\|_2) \right],$$

where $\mathcal{N}(i)$ is the set of senders connected to $i$ by the trained LR edge predictor, $m_j$ is the predicted mass of sender $j$, and $R$ maps the hidden message back to 50-PC state space. Thus $\mathbf{c}_{j\to i}$ describes the direction and magnitude of the contribution from sender $j$ to receiver $i$ in the fitted model. Self-edges were excluded, but same-type edges between distinct cells were retained. For the interaction analysis, we evaluated each of the three Full models using five random cell groupings and averaged the scores across groupings and models.

CellChat was run on the same expression data and 193 LR pairs, and rankings were compared by Spearman correlation across all 121 directed cell-type pairs (top 30 positive inter-type edges shown in Supplementary Figure S5e).

The pathway score was

$$S_{A\to B,p} = D_{A\to B} Q_{A\to B,p},$$

with the LR-expression term

$$Q_{A\to B,p} = \frac{1}{P n_p N_{AB}} \sum_{(L,R)\in p} \sum_{j\in A,\, i\in B} \widetilde{a}_{j,L} \widetilde{a}_{i,R}.$$

Here $P = 52$, $n_p$ counts LR rows in pathway $p$, and $N_{AB}$ counts directed non-self cell pairs. Activities were scaled by training-cell 95th percentiles and clipped to $[0, 1]$. Complexes used the least-expressed required subunit. Dividing by $Pn_p$ gives each pathway equal weight regardless of the number of LR pairs it contains.

At day 6, positive $S_{A\to B,p}$ values were summed across all ordered pairs of different cell types. Supplementary Figure S5f shows the eight largest pathway shares of the total positive $S$ and uses only CytoBridge scores with the fixed LR annotation.

#### S7.2 Metabolic-RNA-labeled cortical activation

##### S7.2.1 scNT-seq cortical KCl data and preprocessing

We next analyzed the scNT-seq primary mouse cortical culture time course (GEO GSE141851) from Qiu et al. [32]. The experiment used 4sU metabolic labeling to distinguish newly transcribed from pre-existing RNA after KCl stimulation. The complete annotated cortical dataset contained 20,547 cells and 24,078 genes at 0, 15, 30, 60, and 120 min, with 2,232, 4,031, 4,470, 6,814, and 3,000 cells, respectively. We retained the nine cell-type annotations (Ex, EX-NP1, EX-NP2, RG, Inh-NP, and Inh1–Inh4) for visualization and cell-type summaries.

For every cell, the raw total count was reconstructed as the sum of the GEO new- and old-RNA count matrices. Training used total RNA only, preprocessed into 50 PCs in the same way as in Supplementary Note S7.1.1. Model times were 0, 0.25, 0.5, 1, and 2 h. Newly transcribed RNA was reserved for evaluating the direction predicted by the trained model.

###### S7.2.2 State-space LR graph and model fitting

The 50-PC vector was the full model state. Candidate ordered pairs at each observed time satisfied $0 < \|z_j - z_i\|_2 < c$, where $c \approx 23.7$ was obtained by estimating the within-time 99th percentile of pairwise PC distances and taking the maximum across the five times.

We applied the CellChatDB filtering and LR-compatibility calculation described for Weinreb, retaining 314 LR pairs in 50 pathways. The edge predictor used the same architecture and data split as in the Weinreb analysis and was applied to the evolving cell states during simulation. The GNN architecture and grouping procedure were the same as in the Weinreb analysis.

We compared three independently trained Full models with three No-interaction models, following the ablation procedure described for Weinreb. For scNT-seq, both conditions used 128-cell batches, a learning rate of $10^{-4}$ and diffusion $\sigma = 0.1$.

###### S7.2.3 Distributional and metabolic-direction evaluation

Each trained model was simulated once from the same 2,048 sampled 0-min cells, using the same Brownian noise and an Euler–Maruyama step size of 0.05 h. Generated and observed cells were compared at 15, 30, 60 and 120 min using the growth-weighted W1 and W2 procedure described for Weinreb. Errors were averaged across the three models in each condition. The Full models had lower mean W1 and W2 at all four time points (Supplementary Figure S6c).

We evaluated transcriptional direction using newly transcribed RNA. A one-shot labeling model

$$n_g = \frac{\alpha_g}{\gamma_g}\left(1 - \exp[-\gamma_g \tau]\right),$$

with labeling time $\tau = 2$ h and gene-specific degradation estimated from baseline cells was used to obtain a new-RNA transcriptional direction. Total and new RNA were normalized with the same total-library size factor, and the resulting log-expression direction was projected through the exact 2,000-HVG PCA loadings. At each observed cell, we compared this reference with the model drift, comprising velocity and score plus the GNN contribution in the Full model. The GNN contribution was averaged over five random cell groupings. For each time point and trained model, we calculated the percentage of cells whose predicted drift formed an angle of less than 90° with the new-RNA direction. We then took the mean and median of the four time-point percentages within each model and averaged each summary across the three models. The mean percentages were 51.19% for Full and 51.12% for No-interaction, and the corresponding temporal medians were 53.73% and 53.34% (Supplementary Figure S6d).

###### S7.2.4 Interaction messages and associated LR pathways

Using one of the three Full models (Supplementary Figure S6b,e,f), we summarized the learned sender-specific GNN messages at 120 min. For sender type $A$ and receiver type $B$, we summed the 50-dimensional messages from type $A$ to each receiver cell, calculated the norm of this sum, and averaged over cells of type $B$ to obtain $D_{A \to B}$, as in the Weinreb analysis. Across all 81 directed cell-type pairs, these rankings correlated at Spearman $\rho = 0.752$ with CellChat run on the same total-RNA expression and LR database (Supplementary Figure S6e, which displays the top 24 positive inter-type edges per method with edge widths indicating within-method rank).

Finally, we annotated the learned message field with the LR pathway score $S_{A \to B,p}$ defined in the Weinreb analysis. PTN contributed the largest share of the pathway score at 120 min (Supplementary Figure S6f). The largest PTN-supported contributions involved excitatory neurons or radial glia as senders and EX-NP or radial-glia states as receivers.

### Supplementary Note S8. Downstream analysis details

#### S8.1 Non-spatial interaction ablation

Supplementary Figures S5c,d and S6c,d compare Full and separately trained No-interaction models using the distributional, clone-fate and transcriptional-direction evaluations described in Supplementary Note S7.

#### S8.2 Forward simulation of developmental trajectories

Given the trained probability-flow network $\mathbf{v}_\theta$, growth network $g_\theta$, scalar score potential $\phi_\theta$, and interaction operator $\mathcal{I}_\theta$, we simulated developmental trajectories from an observed initial population using stochastic dynamics consistent with the UMFSB formulation. Here $\mathbf{s}_\theta(\mathbf{x}, t)$ approximates the score $\nabla_\mathbf{x} \log \rho_t(\mathbf{x})$ (with $\mathbf{x}$ denoting the $d$-dimensional joint state in this subsection), and the intrinsic diffusive drift is recovered as

$$\mathbf{b}_\theta(\mathbf{x}, t) = \mathbf{v}_\theta(\mathbf{x}, t) + \frac{\sigma^2}{2}\mathbf{s}_\theta(\mathbf{x}, t).$$

We considered two complementary simulation modes.

**Weighted SDE (fixed particle number).** In the weighted formulation, each particle $i$ carries a state $\mathbf{x}_i(t) \in \mathbb{R}^d$ and a log-weight $\ell_i(t) = \log w_i(t)$. The coupled system evolves as

$$\mathrm{d}\mathbf{x}_i = \left[\mathbf{v}_\theta(\mathbf{x}_i, t) + \frac{\sigma^2}{2}\mathbf{s}_\theta(\mathbf{x}_i, t) + \mathcal{I}_\theta(\mathbf{x}_i, t; \{\mathbf{x}_j, \ell_j\})\right] \mathrm{d}t + \sigma\, \mathrm{d}B_i,$$

$$\mathrm{d}\ell_i = g_\theta(\mathbf{x}_i, t)\, \mathrm{d}t.$$

Here $B_i$ is a standard $d$-dimensional Brownian motion and $\sigma \geq 0$ is the diffusion coefficient. The interaction term is evaluated by the learned interaction operator on random sub-groups of $m$ particles, where $\mathcal{G}_i$ denotes the sub-group containing particle $i$. Within each sub-group, candidate cell pairs are filtered by the spatial cutoff and the fixed edge predictor. The GNN then computes attention and interaction contributions on the retained edges. The system is integrated with Euler–Maruyama,

$$\mathbf{x}_i^{(n+1)} = \mathbf{x}_i^{(n)} + f\left(\mathbf{x}_i^{(n)}, t^{(n)}\right) \Delta t + \sigma\, \varepsilon_i \sqrt{\Delta t}, \qquad \varepsilon_i \sim \mathcal{N}(0, I_d),$$

$$\ell_i^{(n+1)} = \ell_i^{(n)} + g_\theta\left(\mathbf{x}_i^{(n)}, t^{(n)}\right) \Delta t,$$

where

$$f(\mathbf{x}_i, t) = \mathbf{v}_\theta(\mathbf{x}_i, t) + \frac{\sigma^2}{2}\mathbf{s}_\theta(\mathbf{x}_i, t) + \mathcal{I}_\theta(\mathbf{x}_i, t).$$

**Split SDE (variable particle number with stochastic division).** In the split formulation, the particle count varies dynamically over time to directly represent cellular proliferation and death. The state SDE is the same as in the weighted formulation, while the key distinction occurs at each output time point $t_k$. For each particle $i$, let

$$q_i = \frac{w_i(t_k)}{w_i(t_{k-1})} = \exp\big(\ell_i(t_k) - \ell_i(t_{k-1})\big)$$

denote the weight ratio relative to the previous output time. When $q_i \geq 1$, particle $i$ is replaced by

$$m_i = \lfloor q_i \rfloor + \text{Bernoulli}\big(q_i - \lfloor q_i \rfloor\big)$$

daughter particles. Each daughter inherits the parent state with additive Gaussian noise,

$$\mathbf{x}_j^{\text{daughter}} = \mathbf{x}_i + \eta_j, \qquad \eta_j \sim \mathcal{N}(0, \sigma_{\text{noise}}^2 I_d),$$

where $\sigma_{\text{noise}}$ is the perturbation scale applied when daughter particles are created. For the standard simulations, we set $\sigma_{\text{noise}} = 0$, so daughter particles initially inherited the parent state without an additional displacement. This parameter is distinct from the continuous SDE diffusion coefficient, which was $\sigma = 0.03$. When $q_i < 1$, particle $i$ survives with probability $q_i$ and is removed otherwise. After division, all surviving particles are re-assigned uniform log-weights,

$$\ell_j = -\log N_0,$$

where $N_0$ is the initial particle count. Unless otherwise noted, we used step size $\Delta t = 0.05$, diffusion coefficient $\sigma = 0.03$, and interaction sub-group size $m = 1{,}024$, with the initial particle count capped at $N_0 = 12{,}000$.

##### S8.3 Zebrafish developmental trajectory and observed growth

For Supplementary Figure S36, the 563 cells observed at $t = 0$ were propagated as one fixed population through $t = 4$ using the learned drift, score, interaction and diffusion fields. The five observed stages are shown separately from the generated trajectory. Cell-state labels were assigned with the same classifier at every generated time point. Supplementary Figure S37 shows the learned growth field evaluated directly on cells from the five observed stages. Growth values were scaled by their within-stage percentiles for display.

##### S8.4 Daughter-cell perturbation sensitivity

For the zebrafish analysis in Supplementary Figure S45, we used the same trained model and all 563 cells at the initial stage. Five paired simulations compared $\sigma_{\text{noise}} = 0$, 0.01, 0.03, and 0.06 while keeping the continuous diffusion coefficient fixed at $\sigma = 0.03$. Each initial cell received a lineage identifier inherited by its daughters, and generated cells were assigned cell-type labels with the same classifier.

At each stage, cell-type composition was calculated as the fraction of living particles assigned to each cell type. For each initial cell type, lineage transition fractions were calculated from the labels of its living descendants. Differences from the $\sigma_{\text{noise}} = 0$ setting were summarized by total-variation distance, $\text{TV}(p, q) = \frac{1}{2} \sum_c |p_c - q_c|$. The lineage score was averaged across initial cell types using their initial abundances as weights. Curves and sensitivity scores report the mean and standard error across the five paired simulations.

##### S8.5 Interpolation of intermediate states

Intermediate states were generated by forward simulation of the learned split-SDE at requested times between observed stages. Spatial maps use the simulated coordinates directly.

**Split-SDE interpolation.** Given observed stages $\{t_0 < t_1 < \cdots < t_S\}$ and interpolation times $\mathcal{T}_{\text{interp}} = \{t_0, \tau_1, \tau_2, \ldots, t_S\}$, the split-SDE was integrated forward to generate simulated cell populations $\{X(\tau)\}_{\tau \in \mathcal{T}_{\text{interp}}}$. In one mode, a single forward integration from $t_0$ to $t_S$ was performed from the observed population at $t_0$. In the interval-by-interval mode, the simulation was reinitialized at each observed stage so that, for each interval $[t_s, t_{s+1}]$, the split-SDE was integrated from the observed population at $t_s$ to the intermediate and endpoint times within that interval.

##### S8.6 Chicken-heart alignment sensitivity

We tested the sensitivity of the chicken-heart analysis to small changes in the initial section coordinates by applying two translations, two rotations, and two combined translation–rotation perturbations to D4, D7, D10, and D14 before alignment. Translation magnitudes were 1 and 2 times the within-section median nearest-neighbor distance and absolute rotation angles were 1° and 3° (combined as 1× with 1° and 2× with 3°); translation directions and rotation signs varied across stages before centering and scaling. For each condition, we repeated alignment, model training, and analysis with the same random seed. Each perturbed model was then compared with an unperturbed repeat (which was also compared with the original model), matching cells by identifier and directed edges by source and target cell.

Across the six perturbations, aligned coordinates differed from those of the unperturbed repeat by at most 1.55% of the reference section radius after a proper rigid transformation without reflection. After the corresponding frame rotation, the median cellwise cosine was at least 0.975 for the full velocity and 0.877 for the interaction velocity, and the normalized weighted Jaccard overlap of directed interaction edges was at least 0.838 (Supplementary Figs. S12 and S13).

##### S8.7 Cell-type label inference at simulated time points

To assign cell types at unobserved time points, we trained a multi-layer perceptron (MLP) classifier on the observed data. For cell-type maps and proportions, we applied spatial $k$-nearest neighbors (k-NN) smoothing as described below. Lineage transitions were calculated from the classifier predictions.

Let $\{(\tilde{\mathbf{z}}_i, y_i)\}_{i=1}^{N_{\text{obs}}}$ denote the observed dataset, where

$$\tilde{\mathbf{z}}_i = \left(t_i, z_i^{(1)}, \ldots, z_i^{(d_f)}\right) \in \mathbb{R}^{d_f+1}$$

is the concatenation of the normalized time stamp and the $d_f$ joint-state coordinates, and $y_i \in \{1, \ldots, C\}$ is the cell-type label. For all five datasets, we used the complete model state, $d_f = 52$, comprising two aligned spatial coordinates and 50 principal-component scores. In chicken heart, the classifier distinguished 13 cell types. Anatomical annotations were used for tissue maps and alignment checks. The classifier

$$h_\phi : \mathbb{R}^{d_f+1} \to \Delta^{C-1}$$

is a residual MLP with hidden size 128 and LeakyReLU activations, trained by minimizing cross-entropy loss. The same classifier was used across analyses of each dataset.

We combined the observed cells from all measured stages and used 90% of each cell type for training and 10% for validation. Singleton classes were retained in the training set. We fitted a separate classifier for each dataset with the same architecture and training schedule, using Adam for 500 epochs and cosine learning-rate decay from $10^{-3}$ to $10^{-5}$. The model selected by validation balanced accuracy (macro-averaged recall) was used directly to assign cell-type labels to generated cells.

For spatial smoothing, the MLP predictions were refined by a $k$-nearest neighbors classifier in the 2D spatial coordinate space using the relation:

$$\widetilde{y}_j = \arg \max_{c \in \{1,\ldots,C\}} \sum_{i \in \mathcal{N}_k(j)} \mathbf{1}[\widehat{y}_i = c]\,,$$

where $\widehat{y}_i$ is the MLP-predicted label and $\mathcal{N}_k(j)$ denotes the spatial $k$-nearest neighbors of cell $j$. For cell-type maps and proportions, we applied spatial smoothing with $k = 10$ in zebrafish, MOSTA and ARTISTA. We retained the unsmoothed labels ($k = 1$) in AD mouse and chicken heart, where the decrease in held-out balanced accuracy was much larger. At $k = 1$, the procedure returns the unsmoothed MLP label.

We quantified the effect of this choice using an evaluation on held-out cells at $k = 1, 5, 10, 20,$ and 50 in all five datasets and a separate generated-trajectory analysis in zebrafish (Supplementary Fig. S46). In the held-out analysis, $k = 1$ gave the highest balanced accuracy in each dataset. ARTISTA balanced accuracy decreased from 0.688 at $k = 1$ to 0.605 at $k = 10$, and chicken-heart balanced accuracy decreased from 0.707 to 0.347 when evaluated against the 13-class cell-type annotation. Across nine generated zebrafish time points, the mean total-variation distance between the $k = 10$ and $k = 1$ cell-type compositions was 7.6%. We followed the same 563 particles at model times $0, 1, 2, 3,$ and 4 and calculated the fraction whose predicted cell type differed between each pair of consecutive time points. Averaged across the four intervals, this fraction was 87.48% at $k = 1$ and 87.52% at $k = 10$.

##### S8.8 Estimation of spatiotemporal cell–cell interactions

To summarize the learned interaction network, we extracted and aggregated attention weights from the GNN-based interaction operator.

At each time point $t$, the interaction operator $\mathcal{I}_\theta$ constructs a spatiotemporal cell–cell graph with edge set $\mathcal{E}_t$. Each edge $(i, j)$ carries attention weights $\alpha_{ij}^{(h)}$ from $H$ attention heads. We extracted the absolute multi-head mean according to

$$\bar{\alpha}_{ij} = \frac{1}{H} \sum_{h=1}^{H} \left| \alpha_{ij}^{(h)} \right| .$$

Self-loops ($i = j$) were removed for downstream analysis.

Given $C$ cell types with $n_c$ cells per type, we then summarized attention between sender and receiver cell types in matrices. The raw summed matrix is

$$M_{\text{sum}}(A, B) = \sum_{\substack{(i,j) \in \mathcal{E}_t \\ y_i = A,\, y_j = B}} \bar{\alpha}_{ij}.$$

To account for differences in cell-type abundance, we divided the summed attention by the number of sender cells:

$$M_{\text{src}}(A, B) = \frac{M_{\text{sum}}(A, B)}{n_A}.$$

We also computed the row-normalized matrix

$$M_{\text{row}}(A, B) = \frac{M_{\text{sum}}(A, B)}{\sum_{B'} M_{\text{sum}}(A, B')},$$

and the per-edge mean

$$M_{\text{mean}}(A, B) = \frac{M_{\text{sum}}(A, B)}{|\{(i, j) \in \mathcal{E}_t : y_i = A,\ y_j = B\}|}.$$

Directional asymmetry was measured as

$$M_{\text{asym}}(A, B) = M_{\text{src}}(A, B) - M_{\text{src}}(B, A).$$

To mitigate the influence of extreme outlier attention values, weights were winsorized at quantile $q$ (default $q = 0.995$).

$$\bar{\alpha}_{ij} \leftarrow \min(\bar{\alpha}_{ij},\ Q_q(\{\bar{\alpha}_e\}_{e \in \mathcal{E}_t}))\,.$$

For analyses of a selected tissue, edges were retained only if either the sender or receiver matched the selected label. Spatially representative centroids for that label were computed using only the fraction $\rho$ (default $\rho = 0.2$) of cells nearest to the global centroid, so that centroid-to-centroid connections in the 3D view reflected the spatial core of the selected tissue.

Cell-type interaction networks were displayed in three dimensions, with physical coordinates on the horizontal axes and time on the vertical axis. Edges were drawn as lines connecting cell-type centroids, with width proportional to $M_{\text{src}}(A, B)$. Cell fate transitions were additionally shown as source-to-target connections between centroids at $t_s$ and $t_e$, with width proportional to the transition fraction $\pi_{A \to B}$.

#### S8.9 Cross-method comparison of spatial interaction summaries

We compared CytoBridge with COMMOT [31] and CellAgentChat [33] on the chicken-heart, MOSTA, ARTISTA, and zebrafish datasets. We ranked all combinations of sender and receiver cell types, assigning zero to pairs absent from a method's output. Agreement was summarized by Spearman correlation over all pairs and by Jaccard overlap between the highest-ranked 20% of directed pairs (Supplementary Fig. S11). The CellAgentChat comparison used the same ligand–receptor database as the CytoBridge analysis and three fixed sampling seeds.

For the molecular comparison, we combined the CytoBridge interaction contribution, normalized for cell-type abundance, with ligand expression in sender cells and receptor expression in receiver cells. Candidate axes required at least 10 active edges in the model. The representative axis in each dataset was selected from CytoBridge before the external results were examined. COMMOT was then used to report the rank of the same ligand–receptor axis within that directed pair and the percentile of the directed pair. For the NicheNet comparison, we report the combination of an LR pair and sender–receiver cell types ranked highest by CytoBridge for which NicheNet reported a positive ligand–target link and list the predicted receiver targets. For zebrafish, genes were mapped to one-to-one mouse orthologs in Ensembl and analyzed using the mouse NicheNet prior.

#### S8.10 ARTISTA local interaction-domain analysis

The local analysis used the 5-DPI region and trained ARTISTA model shown in Fig. 5c. For each cell, we compared the spatial component of the full velocity with the corresponding velocity obtained from the interaction operator alone by cosine similarity. Cells in the upper quartile of cosine similarity were connected using the spatial radius used during training. Connected components containing at least 20 cells were defined before attention or ligand–receptor enrichment was evaluated. This yielded the two reported domains.

Within each domain, we compared attention per cell and LR scores with random cell sets drawn from the same 5-DPI region and containing the same number of cells of each type. We used 9,999 random sets for attention and 1,999 for LR scores. For either test, the upper-tail $P$ value was $[1 + \sum_{b=1}^{B} \mathbf{1}\{T_b \geq T_{\text{obs}}\}]/(B+1)$, where $B$ is the number of random sets, $T_{\text{obs}}$ is the observed score, and $T_b$ is the score for random set $b$. Sender–receiver patterns were summarized from attention on edges within each domain.

For LR enrichment, Benjamini–Hochberg correction was applied separately to pathway scores and to the 531 pair-level tests within each domain. One pair was displayed per significant pathway, ranked by adjusted $P$ value, followed by score and fold enrichment.

##### S8.11 Learned interaction attention in zebrafish

For external method comparison, cell-level learned attention was aggregated over all 361 directed cell-type pairs in the zebrafish model. Rank comparisons with COMMOT and a CellAgentChat calculation using the same LR database were adjusted for sender abundance, receiver abundance, mean spatial distance, and whether the sender and receiver had the same cell type. For the permutation comparison, CytoBridge scores were shuffled within 22 groups of pairs with similar abundance and distance and the same self-pair status, while COMMOT scores were held fixed. Adjusted correlations were recalculated after each of 1,000 permutations (Supplementary Figure S38a shows the COMMOT comparison at $t = 4$).

The JAM analysis used the 375 Somite cells at 18 hpf and used the same 677 model edges for the trained, pre-interaction, and randomized-weight controls. A JAM-compatible edge required *jam2a* detection at one endpoint and *jam3b* detection at the other. The pre-interaction control used the same model immediately before the interaction-learning stage, whereas the randomized control permuted the learned interaction weights on the same cells and graph edges. Top- and bottom-attention quartiles, Somite-to-Somite rankings, and corresponding odds ratios were calculated using the same cells and edges.

For the spatial co-detection test, two distinct Somite cells were considered neighbors when their Euclidean distance in the aligned spatial coordinates was at most 0.0961 (the model's spatial-neighborhood radius), giving 4,426 undirected neighbor pairs among the 375 cells. A complementary pair had *jam2a* detected at one endpoint and *jam3b* at the other, in either orientation, and was counted once. We compared the observed count against 10,000 permutations that independently shuffled the two genes' detection labels among the Somite cells while preserving cell positions, neighbor pairs, and the positive-cell count of each gene. The anatomical map displays all 3,048 cells, with the 375 Somite cells highlighted. Associations of *jam3b* or *jam2a* detection with *myog* detection were summarized by odds ratios and Fisher's exact tests. Prior studies establish Jam2a/Jam3b as heterophilic myocyte-fusion factors and report reduced *jam3b*, but not *jam2a*, in *myog*-null embryos [34, 35].

##### S8.12 Cell lineage transition analysis

Cell lineage transition summaries were estimated from simulated trajectories after assignment of cell-type identities.

For lineage transition analysis, a fixed set of particles was propagated with the weighted SDE formulation described in Supplementary Note S8.2 from a source time $t_s$ to later output times, using the same learned velocity, growth, and interaction fields as in the main simulations but without stochastic division. At each output time, cell types were assigned by the classifier $h_\phi$. Transitions were counted by following the same particle across time. Let

$$\mathcal{I}_A(t_s) = \{i : \widehat{y}_i(t_s) = A\}$$

denote the set of simulated particles belonging to source label $A$ at time $t_s$. The transition fraction from source label $A$ at $t_s$ to target label $B$ at time $t_e$ was estimated as

$$\pi_{A\to B}(t_s, t_e) = \frac{1}{|\mathcal{I}_A(t_s)|} \sum_{i\in\mathcal{I}_A(t_s)} \mathbf{1}[\widehat{y}_i(t_e) = B]\,.$$

For Fig. 5a and Supplementary Fig. S25, transitions involving fewer than 10 particles were omitted. The largest remaining transitions were retained until their cumulative fraction reached 85% for each source cell type.

For the cartilage-primordium analysis in Fig. 4c, particles classified as cartilage primordium at E15.0 were selected from the continuous MOSTA trajectory and followed to E15.5. Their target cell types defined the transition fractions. Source and target centroids were connected by lines whose widths increased with the transition fraction. Supplementary Fig. S25 summarizes ARTISTA transitions and cell-type composition using the same classified particles.

#### S8.13 Velocity vector decomposition and spatiotemporal embedding

The trained neural networks define a continuous velocity field over the joint spatial-expression state space. We decomposed and visualized these velocity fields to characterize intrinsic differentiation dynamics and multicellular interaction forces.

At any cellular state $\mathbf{x} \in \mathbb{R}^d$ and normalized time $t$, the total instantaneous velocity of the split-SDE is

$$\mathbf{v}_{\text{full}}(\mathbf{x}, t) = \underbrace{\mathbf{v}_\theta(\mathbf{x}, t)}_{\text{drift}} + \underbrace{\frac{\sigma^2}{2}\nabla_{\mathbf{x}} \log \rho_t(\mathbf{x}, t)}_{\text{score contribution}} + \underbrace{\mathbf{F}_{\text{int}}(\mathbf{x}, t)}_{\text{interaction}}.$$

Here $\mathbf{v}_\theta$ is the probability-flow velocity, $(\sigma^2/2)\nabla_{\mathbf{x}} \log \rho_t(\mathbf{x}, t)$ is the score contribution, and $\mathbf{F}_{\text{int}}(\mathbf{x}, t)$ is the interaction force induced by the learned interaction operator on the spatiotemporal cell–cell graph. In the particle implementation, $\mathbf{F}_{\text{int}}$ is evaluated from local graph neighborhoods of the current particle system, with interaction neighborhood size $m$ set to 1,024 by default.

In physical space, the spatial velocity components were used directly as 2D vectors overlaid on anatomical coordinates. For gene-expression space visualization, we adapted the scVelo velocity embedding algorithm by constructing a $k$-nearest-neighbor graph on latent gene-expression features, computing cosine similarity between each cell's velocity vector and neighbor displacements, converting these similarities into transition probabilities via softmax normalization, and projecting the resulting transition matrix into a 2D embedding. In ARTISTA, Fig. 5c shows the full spatial velocity and its cosine similarity with the interaction component. Figure 5d instead shows the intrinsic-context gene-expression velocity in transcriptomic principal-component space.

To examine regionally resolved dynamics within the developing brain, velocity analysis was restricted to cells annotated as brain and further subclassified into telencephalon cell types. Stream plots were then generated within a spatial zoom region to inspect migration and differentiation flow patterns at higher local resolution.

#### S8.14 Chicken-heart velocity components and spatial comparison

For the chicken-heart analysis, the full, drift, and interaction components of the model-implied gene-expression velocity were evaluated at D10 and displayed over the aligned tissue coordinates. We summarized the projected full-velocity magnitude, the interaction-to-drift magnitude ratio, and the cosine similarity between the interaction and drift components. At D14, CytoBridge and veloAgent [36] streamlines were plotted on the same anatomical map to compare their spatial fields (Supplementary Fig. S15).

#### S8.15 Developmental gene wave map, temporal phase segmentation, and pattern clustering

The split-SDE operates in a latent coordinate space $x = [x_{\mathrm{sp}}, x_{\mathrm{ge}}]$, where $x_{\mathrm{sp}} = (z_1, z_2) \in \mathbb{R}^2$ are spatial coordinates and $x_{\mathrm{ge}} = (z_3, \ldots, z_{52}) \in \mathbb{R}^{50}$ correspond to the first 50 principal components of log-normalized gene expression. To reconstruct expression values in the original gene space, we inverted the PCA transformation via the inverse relation

$$\widehat{g}_i = W^\top z_i^{[3:52]} + \mu.$$

Here $W$ is the PCA loading matrix and $\mu$ is the gene-wise mean vector estimated from the training data. Pseudo-count values were obtained by inverse log1p transformation with negative values clipped to zero.

For a given cell type of interest, simulated cells were filtered by inferred labels and the mean reconstructed expression was computed at each time point using the formula

$$\bar{g}^{(c)}(t) = \frac{1}{|\{i : \widetilde{y}_i = c,\ t_i = t\}|} \sum_{i:\widetilde{y}_i = c,\ t_i = t} \widehat{g}_i.$$

Stacking these values across time yielded a $G \times T$ expression matrix.

Genes were ranked by temporal variance, standardized to Z-score profiles, and clustered using agglomerative hierarchical clustering with Ward linkage. For each cluster, a prototype temporal pattern was defined as the mean Z-score trajectory. Representative genes were selected by combining temporal variance and correlation with the prototype curve.

Each variable gene was assigned a peak activation time and ordered accordingly to construct a developmental wave map. The ordered peak times were further partitioned into contiguous temporal phases by minimizing intra-segment variance with dynamic programming. For the MOSTA temporal programs and developmental waves, enrichment was performed with clusterProfiler 4.10.0 and org.Mm.eg.db 3.18.0. Biological-process, molecular-function, and cellular-component terms were combined for enrichment analysis, using the eligible highly variable genes detected in Brain cells as the background; Benjamini–Hochberg correction was applied once over the merged term collection for each query. For the ARTISTA temporal programs, annotated axolotl ortholog symbols were mapped through org.Mm.eg.db, and the enrichGO function in clusterProfiler was run for biological-process terms with the 1,736 mapped genes detected in the dataset as the background. Benjamini–Hochberg-adjusted $P < 0.05$ defined significance. The 20 displayed terms per ARTISTA program were ranked by raw $P$ value for visualization.

#### S8.16 Ligand–receptor interaction scoring

To examine ligand–receptor signaling between cell types, we combined the aggregated attention weights with ligand expression in sender cells and receptor expression in receiver cells, using pairs from CellChatDB.

We used CellChatDB ligand–receptor annotations, including multi-subunit complexes. MOSTA and zebrafish used the mouse database. ARTISTA and chicken heart used human entries matched through conserved gene symbols. Gene expression values were obtained from the reconstructed gene-space representation for interpolated time points or from the measured expression data for observed time points.

For each cell type $c$, mean gene expression was computed as

$$\bar{e}_c(g) = \frac{1}{n_c} \sum_{i:y_i=c} e_i(g).$$

For multi-subunit ligands or receptors, the effective expression was taken as the element-wise minimum across subunits, providing an AND-gate approximation limited by the least abundant component.

For each ligand–receptor pair $(L, R)$ and sender–receiver cell-type pair $(A, B)$, the interaction score was computed as

$$\mathrm{LR}(A \to B; L, R) = \bar{e}_A(L) \cdot \bar{e}_B(R) \cdot M_{\mathrm{src}}(A, B).$$

#### S8.17 Continuous ligand–receptor dynamics

Continuous LR dynamics were evaluated along a single split-SDE trajectory through the observed and intermediate times. Observed time points retained their original cell-type annotations, whereas interpolated time points were assigned labels using the trained MLP classifier described above.

At each time point, we calculated the cell-type interaction matrix $M_{\mathrm{src}}(A, B; t)$, normalized by the number of sender cells, using the attention aggregation, self-loop exclusion, and winsorization described in Supplementary Note S8.8.

For the MOSTA ligand–receptor temporal-pattern panel and complex-aggregation sensitivity analysis, we used the total cell-type interaction matrix

$$M_{\mathrm{sum}}(A, B; t) = \sum_{i:y_i(t)=A} \sum_{j:y_j(t)=B} \bar{\alpha}_{ij}(t).$$

The MOSTA analysis used 12,000 initial cells, propagated continuously from $t = 0$ to seven half-step states. In the following equations, $M_{\mathrm{sum}}$ replaces $M_{\mathrm{src}}$ for this panel and its aggregation-sensitivity analysis.

Expression at interpolated times was reconstructed by inverse PCA and inverse log transformation as described in Supplementary Note S8.15. Observed times used measured expression in the same count-space convention. LR pairs and multi-subunit complexes were scored as described in Supplementary Note S8.16, retaining one entry per distinct LR pair.

Let $S_\ell(A, B; t)$ denote the LR score for pair $\ell$ and sender–receiver types $(A, B)$ at time $t$, calculated as in Supplementary Note S8.16. To obtain a pair-level continuous trajectory, this matrix was reduced to a single global score

$$G_\ell(t) = \sum_{A,B} S_\ell(A, B; t).$$

Raw pair-level trajectories $G_\ell(t)$ were used for ranking and reporting, and row-wise normalized trajectories were used in heatmaps and small-multiple plots to show their changes over time. LR pairs were then grouped by trajectory shape using hierarchical clustering on the normalized time-course profiles.

#### S8.18 NicheNet analysis of the reconstructed AD trajectory

For each adjacent interval of the reconstructed AD trajectory, the 50 microglial genes with the largest positive change were used as the target set for NicheNet. Candidate ligands were ranked by corrected area under the precision–recall curve (AUPR) and summarized by their frequency among the five highest-ranked ligands and their mean activity across analyses. At six inferred ages, prioritized ligands were connected to receptors expressed in microglia using the NicheNet prior ligand–receptor weights (Supplementary Fig. S34).

##### S8.19 In silico Trem2 and Spp1 perturbations

Trem2 and Spp1 activation and knockdown were applied along the corresponding gene's PCA loading direction without changing the initial spatial coordinates. The shift was applied to all cells of the initial 2.5-month population and equaled the gene's normalized PCA loading multiplied by the standard deviation of each principal component, scaled by 1.0 for Trem2 and 2.5 for Spp1. Perturbed and unperturbed TgCRND8 states were propagated with the same split-SDE settings, assigned cell types with the trained classifier, and mapped back to gene space. Cell-type distributions and proportions were compared with the unperturbed reference for Trem2 (Fig. 6f). Functional-module scores were calculated at model time 2.5, beyond the last observed age of 17.9 months, as the mean Z-scored expression of the detected genes in each module and reported relative to the unperturbed reference (Fig. 6g and Supplementary Fig. S35).

##### S8.20 Sensitivity to multi-subunit ligand–receptor aggregation

The primary LR analysis represented a multi-subunit ligand or receptor complex by the minimum expression among all required subunits. To test whether the reported LR dynamics depended on this aggregation choice, we repeated only the complex-expression aggregation with a zero-preserving geometric mean. Both rules assign zero activity when any required subunit has zero expression and were evaluated on the same model outputs and LR pairs at each time point.

Across zebrafish, MOSTA, ARTISTA, and chicken heart, Spearman correlations between the two aggregation rules ranged from 0.960 to 0.999 when all scored LR pair–time observations were combined. Overall LR rankings were similar, with larger differences among the top-ranked complexes in ARTISTA and chicken heart (Supplementary Figure S47).

##### S8.21 Virtual tissue ablation

To assess the impact of specific cell types in developmental morphogenesis, we performed *in silico* virtual tissue ablation and quantified the resulting morphological divergence. Given an observed cell population at time $t_0$, we constructed a baseline simulation arm by integrating the full population forward with the split-SDE and a counterfactual arm by removing a target cell type at $t_0$ and integrating the remaining population forward under the same dynamics.

For the zebrafish analyses in Supplementary Figures S39 and S40, YSL or EVL cells were removed from the 563-cell initial population, leaving 534 or 291 cells. Baseline and virtual-removal populations were propagated to $t = 4$ with the same trained model and paired simulation seeds. Spatial Wasserstein-1 distance and centroid shift quantified the deviation from the paired baseline. Curves report mean and standard error across five simulations, and endpoint centroid shifts are shown with 95% confidence intervals.

##### S8.22 YSL-lineage gene dynamics and inverse-PCA validation

The YSL-lineage analysis in Supplementary Figure S41 begins with the 29 YSL cells observed at $t = 0$. Their descendants were represented with the continuous-weight formulation through $t = 4$, and gene expression was reconstructed from the evolving PCA state. The two adjacent heat maps show row-wise z scores for 250 temporally variable genes.

We compared inverse-PCA expression with observed expression at the five measured stages in Supplementary Figure S42. For each stage, clipped inverse-PCA mean expression was compared with observed mean log-transformed expression over the analyzed genes. Across the five stages, Pearson $r$ ranged from 0.890 to 0.996 and root-mean-square error from 0.01 to 0.06.

#### Supplementary Table 1. Comparison of related transport formulations

| Formulation | Diffusion | Growth or loss | Cell–cell interaction | What the model describes |
|---|---|---|---|---|
| Dynamical optimal transport | No | No | No | Deterministic transport. Total mass is fixed. |
| Unbalanced dynamical optimal transport | No | Yes | No | Deterministic transport with population growth or loss. |
| Schrödinger bridge | Yes | No | No | Stochastic transport around a reference diffusion. Total mass is fixed. |
| Unbalanced Schrödinger bridge (regularized unbalanced optimal transport) | Yes | Yes | No | Stochastic transport with population growth or loss. |
| Mean-field Schrödinger bridge | Yes | No | Yes | Stochastic transport with population-dependent interactions. Total mass is fixed. |
| Unbalanced mean-field Schrödinger bridge (CytoBridge) | Yes | Yes | Yes | Stochastic transport with population growth or loss and population-dependent interactions. |

**Supplementary Table 2. Graph-construction and training settings used for the five applications**

| Dataset | ODE batch | Spatial cutoff | Edge cutoff | Score batch | $(\lambda_{\mathrm{OT}}, \lambda_{\mathrm{mass}})$[a] |
|---|---|---|---|---|---|
| AD mouse | 512 | 0.0121 | 0.9957 | 256 | $(1, 0.01)$; $(10, 10)$; $(10, 10)$ |
| ARTISTA | 512 | 0.0315 | 0.6524 | 128 | $(1, 0.01)$; $(10, 10)$; $(1, 1)$ |
| Chicken heart | 128 | 0.2168 | 0.1501 | 128 | $(1, 0.01)$; $(10, 10)$; $(10, 10)$ |
| MOSTA | 1024 | 0.0240 | 0.1192 | 512 | $(1, 0.01)$; $(10, 10)$; $(10, 10)$ |
| Zebrafish | 512 | 0.0961 | 0.6064 | 128 | $(1, 0.01)$; $(10, 10)$; $(10, 10)$ |

[a]Entries give the values for Pretrain and Refine, Init interaction, and Finetune, respectively.

#### Supplementary Table 3. Computational cost of the full CytoBridge models

| Dataset | Time points used for training | Observed cells/spots | Training time (min) | Peak host memory (GiB) | Peak GPU allocation (GiB) |
|---|---|---|---|---|---|
| AD mouse | 3: 2.5, 5.7, 17.9 months | 172,092 | 11.6 | 2.39 | 0.69 |
| ARTISTA | 5: 2, 5, 10, 15, 20 DPI | 46,199 | 22.1 | 9.05 | 0.85 |
| Chicken heart | 4: D4, D7, D10, D14 | 3,550 | 12.7 | 2.77 | 1.88 |
| MOSTA | 4: E12.5, E13.5, E14.5, E15.5 | 344,603 | 33.3 | 16.62 | 1.03 |
| Zebrafish | 5: 5.25, 10, 12, 18, 24 hpf | 11,999 | 20.7 | 2.12 | 0.56 |

#### Supplementary Figure S1. CytoBridge analysis workflow

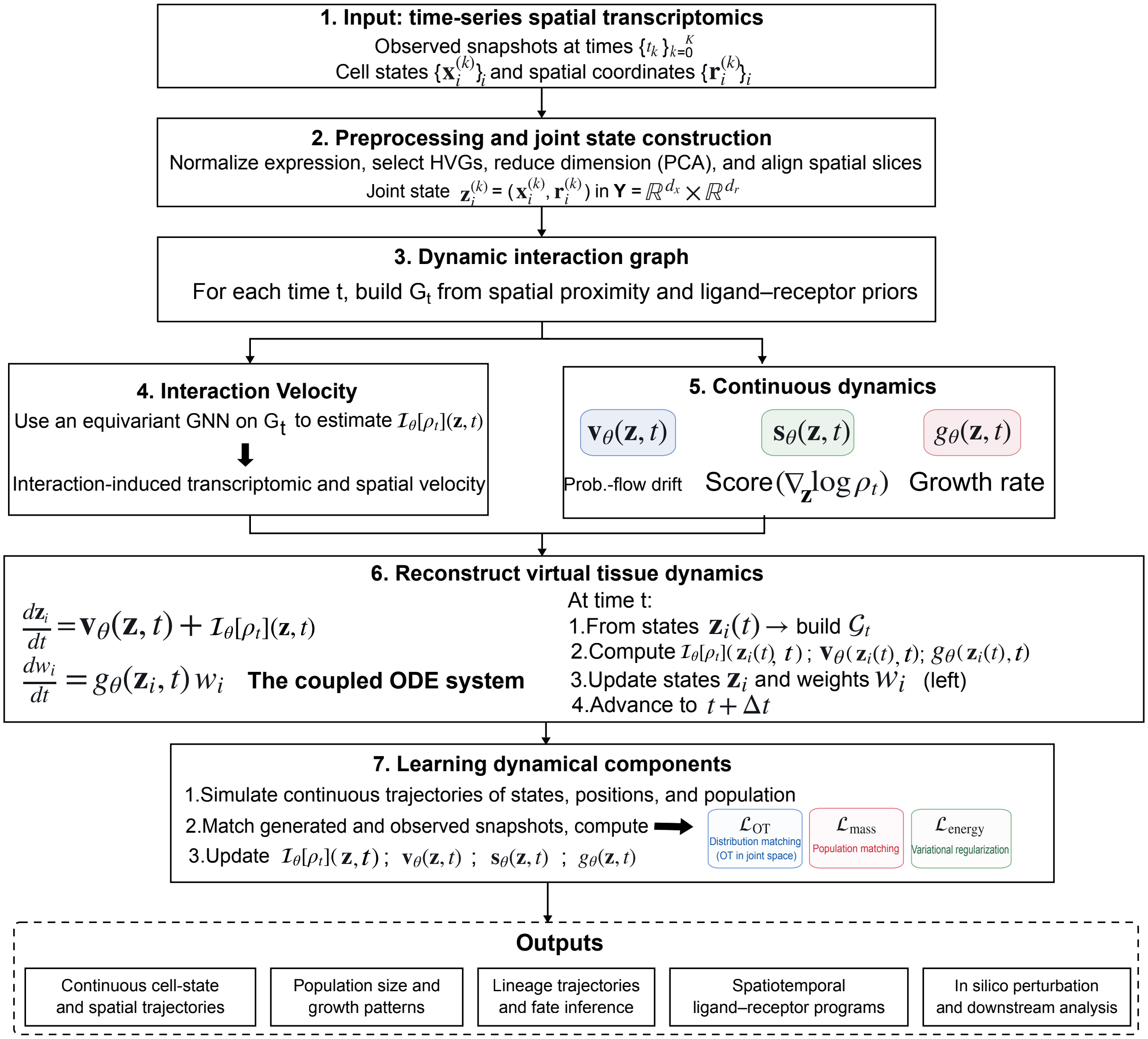

**Supplementary Figure S1.** CytoBridge analysis workflow. Spatial transcriptomic snapshots are preprocessed into joint transcriptomic–spatial cell states, and a dynamic interaction graph is built from spatial proximity and ligand–receptor priors. An equivariant graph neural network estimates the interaction velocity on this graph, and separate networks estimate the probability-flow drift, the score and the growth rate. Simulated weighted particles are matched to the observed snapshots during training, and the trained model is used for the analyses shown at the bottom.

### Supplementary Figure S2. Simulation benchmark of growth, attention and distributional recovery

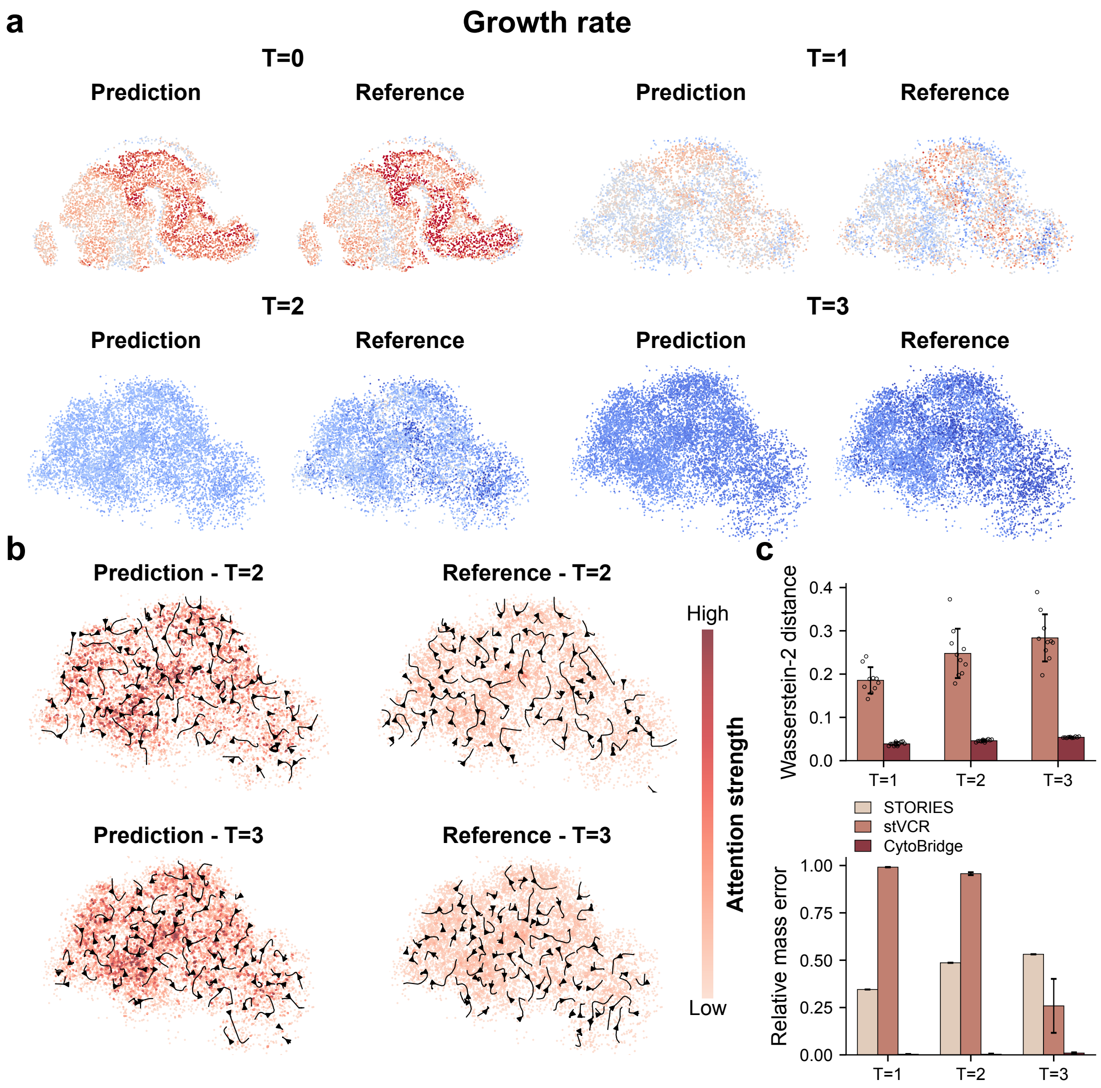

**Supplementary Figure S2.** Simulation benchmark of growth, attention and distributional recovery. (a) Inferred and generator growth rates evaluated at the same simulated cell states, with a shared color scale. (b) Attention strengths and flows at $t = 2$ and $t = 3$. (c) Spatial Wasserstein-2 distance and relative population-mass error for the predictions in Fig. 2e. For CytoBridge and stVCR [18], bars and error bars show the mean and standard deviation across ten predictions with different random seeds from one trained model per method, and open circles in the spatial-distance plot show individual predictions. Population mass changes through stochastic cell division and death in stVCR and stays at the initial particle count in STORIES [25].

#### Supplementary Figure S3. Velocity agreement across time and state space in the AGIST benchmark

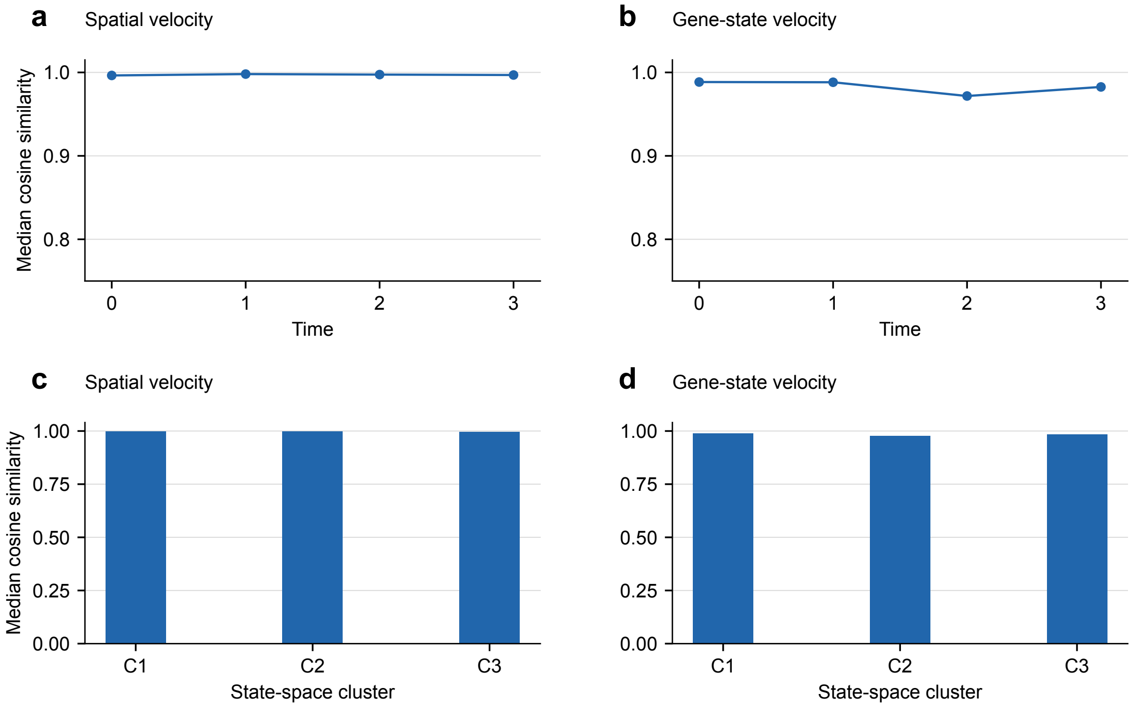

**Supplementary Figure S3.** Velocity agreement across time and state space in the AGIST benchmark. (a,b) Median cellwise cosine similarity between inferred and generator velocity vectors in physical and gene-expression space, respectively, at each simulated time point. The velocity vectors comprise the probability-flow, interaction and score contributions defined in Supplementary Note S5.3. (c,d) Median velocity agreement in three expression-state clusters obtained by MiniBatch $k$-means in the 50-dimensional gene-state space. The same cluster centroids were used at every time point.

#### Supplementary Figure S4. Synthetic spatiotemporal benchmark with finite-range attractive cell–cell interactions

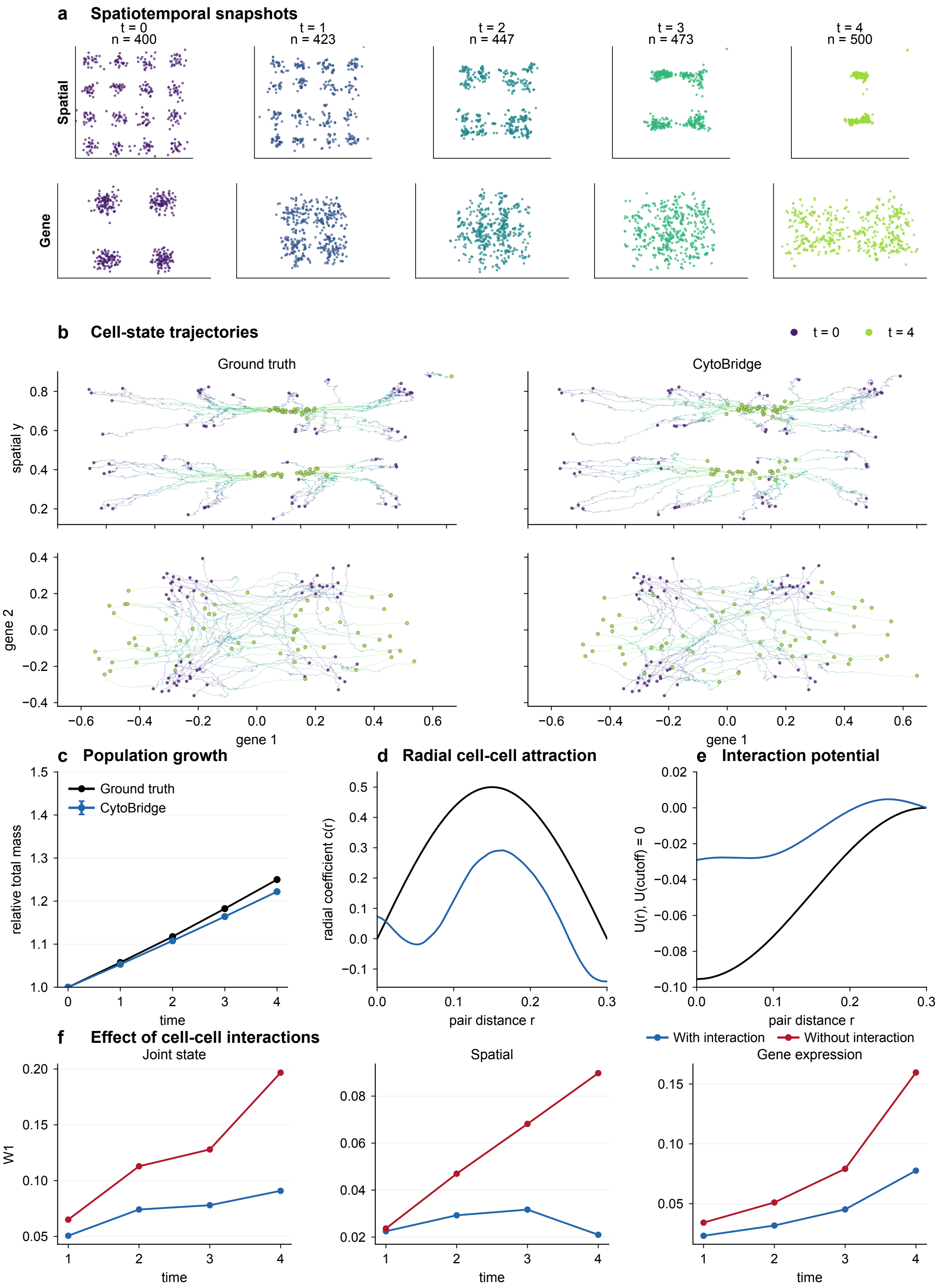

**Supplementary Figure S4.** Synthetic spatiotemporal benchmark with finite-range attractive cell–cell interactions. (a) Simulated spatial and gene-expression snapshots. Cell number increased from 400 to 500. (b) Ground-truth and inferred trajectories from the same 400 initial cells, propagated without growth-dependent resampling. (c) Ground-truth and inferred relative population mass (mean absolute TMV over $t = 1$–$4$, excluding $t = 0$: 0.0151). (d) Ground-truth and inferred radial interaction coefficients. Positive values denote attraction (Pearson $r = 0.876$, Spearman $\rho = 0.899$, sign agreement 73.1%). (e) Interaction potentials obtained by integrating the radial force profiles and setting $U(r_c) = 0$ (Pearson $r = 0.980$). (f) W1 after propagation with or without the inferred interaction term. At the final time point, interaction reduced W1 by 0.1059, 0.0688 and 0.0820 in joint, physical and gene-expression space, respectively. In (c), points show the mean and error bars show s.d. across five inference seeds. In (f), curves show the mean and error bars show s.e.m. across the same five seeds.

### Supplementary Figure S5. Interaction-informed dynamics in the Weinreb lineage-tracing dataset

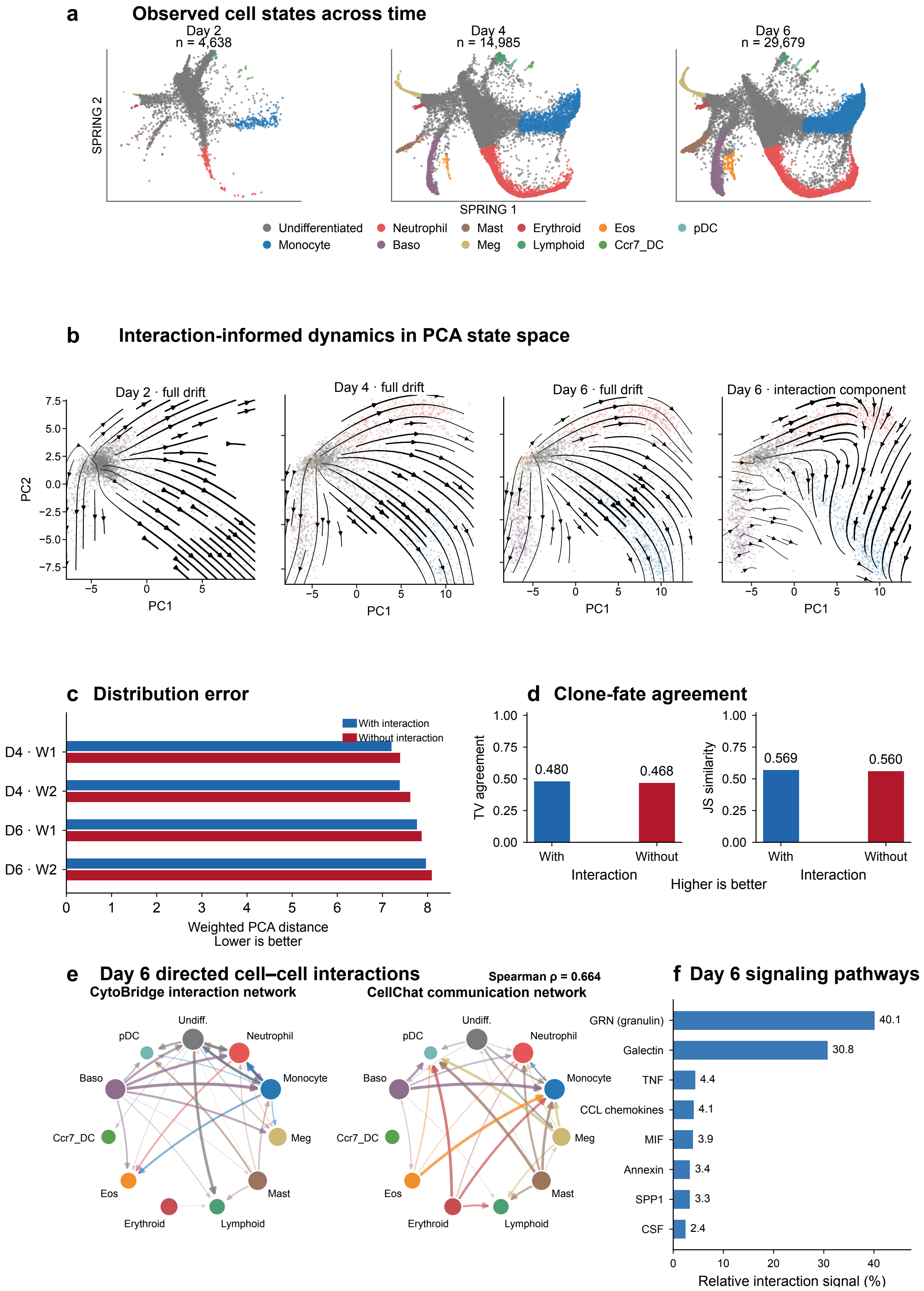

**Supplementary Figure S5.** Interaction-informed dynamics in the Weinreb lineage-tracing dataset. (a) SPRING visualization colored by cell type. (b) The Full-model state-change field and its interaction component at day 6 in PC1–PC2 space. (c) Distributional reconstruction errors at days 4 and 6. Bars show the mean error across three independently trained models per condition, each evaluated in 50 simulations. (d) Agreement with the observed fate proportions of 1,237 lineages, calculated after averaging predicted proportions across the three models, each evaluated in ten simulations (Supplementary Note S7.1). (e) The top 30 day-6 directed cell-type edges from CytoBridge and CellChat. Spearman $\rho = 0.664$ was calculated over all 121 edges. (f) Day-6 pathway annotation of the CytoBridge interactions. GRN and galectin dominated, with the largest contributions involving Monocyte→Neutrophil and Baso→Monocyte/Neutrophil. Smaller TNF, CCL, MIF, annexin, SPP1, and CSF signals support a late myeloid signaling interpretation.

### Supplementary Figure S6. Interaction-informed dynamics in the scNT-seq cortical activation dataset

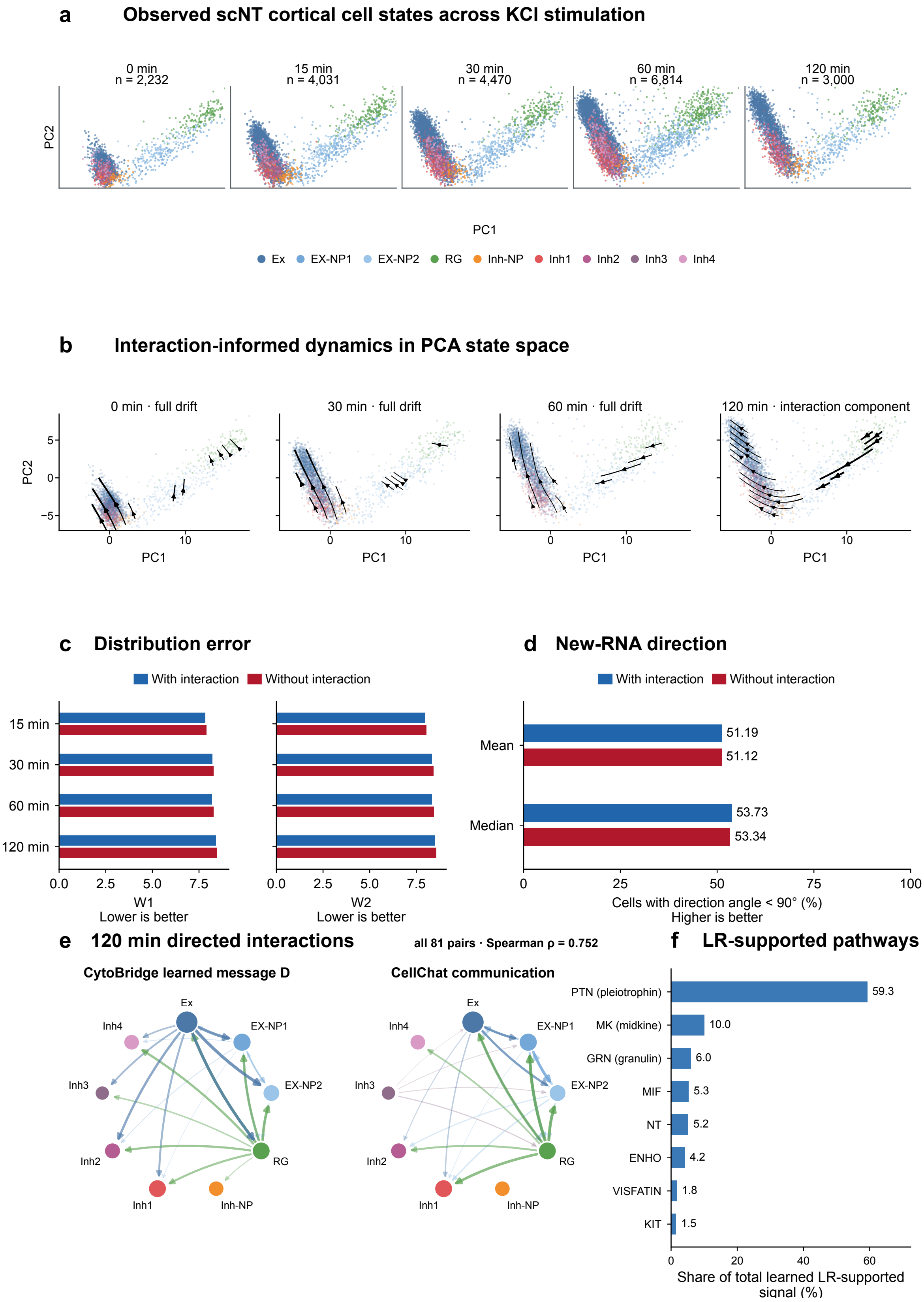

**Supplementary Figure S6.** Interaction-informed dynamics in the scNT-seq cortical activation dataset. (a) Primary mouse cortical cells at 0, 15, 30, 60, and 120 min after KCl stimulation, shown in PC1–PC2 and colored by the nine cell-type annotations. (b) Full-model state-change fields at 0, 30, and 60 min and the learned interaction component at 120 min in PC1–PC2 space. (c) Distributional reconstruction errors at four observed times, averaged across three independently trained models per condition. (d) Percentage of cells whose predicted drift forms an angle of less than 90° with the direction estimated from newly transcribed RNA. Mean and Median summarize the four time-point percentages within each model, followed by averaging across the three models. Training and evaluation are described in Supplementary Note S7.2. (e) The top 24 positive edges between different cell types at 120 min, shown for CytoBridge and CellChat. Spearman $\rho = 0.752$ was calculated over all 81 directed pairs. (f) The 120-min LR annotation of the CytoBridge interaction field. PTN dominated, with prominent contributions from excitatory neurons or radial glia to excitatory-neuron progenitors or radial glia, followed by MK, GRN, MIF, NT, ENHO, VISFATIN, and KIT.

### Supplementary Figure S7. Five-dataset leave-one-time-point-out benchmark across methods

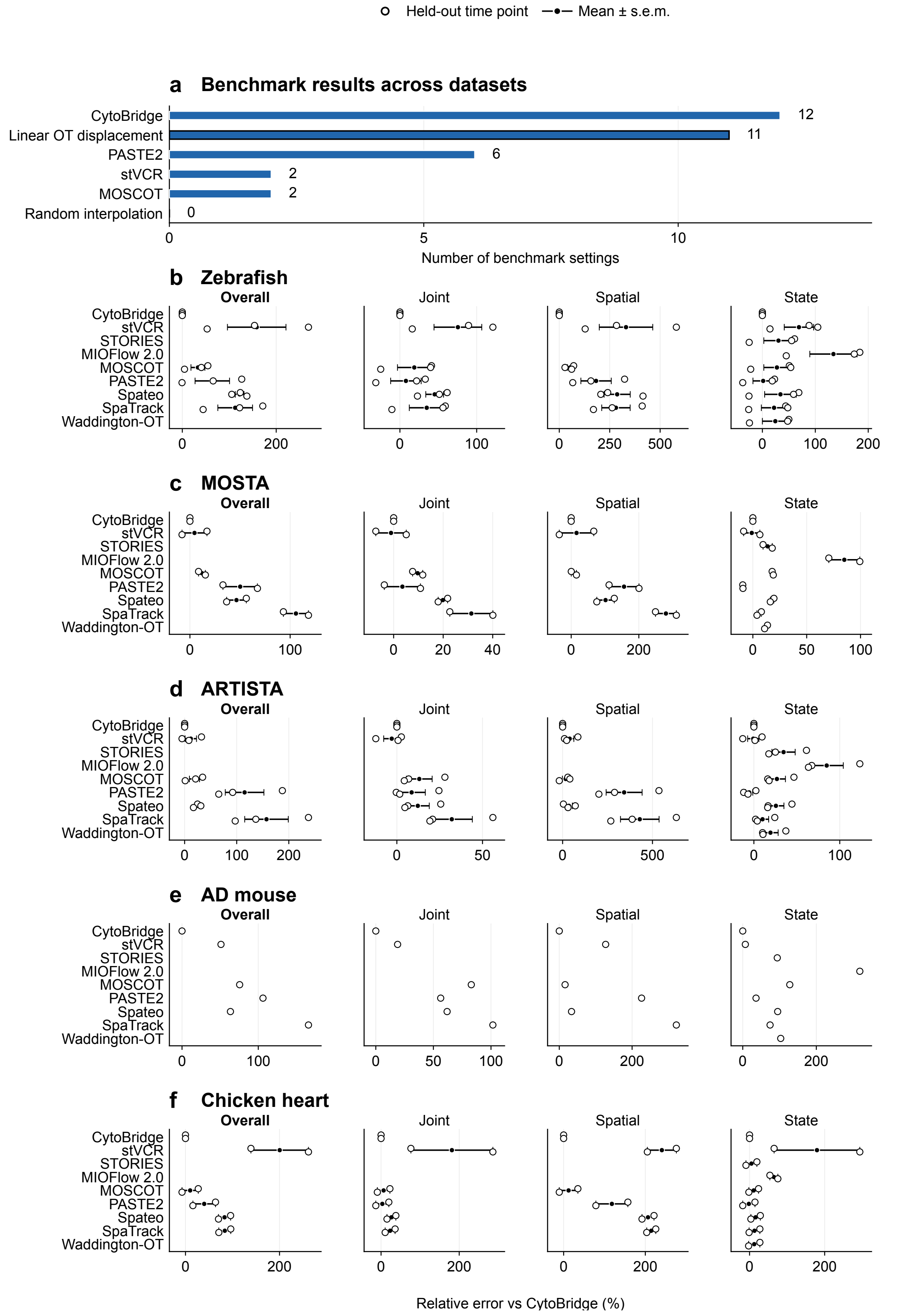

**Supplementary Figure S7.** Five-dataset leave-one-time-point-out benchmark across methods. (a) Number of benchmark settings in which each method had the lowest mean Sliced-W2 among the applicable methods. The 33 settings span matched datasets, held-out time points, and evaluation spaces. CytoBridge had the lowest value in 12 settings, Linear OT displacement in 11, PASTE2 in 6, stVCR in 2, and MOSCOT in 2. Random interpolation did not have the lowest value in any setting. (b–f) Relative Sliced-W2 compared with CytoBridge for zebrafish, MOSTA, ARTISTA, AD mouse, and chicken heart. Positive values indicate lower error for CytoBridge. Open circles represent held-out time points, filled circles represent means, and error bars show s.e.m. across held-out time points. Overall values are calculated for each held-out time point by averaging the joint, spatial, and gene-state relative errors. MIOFlow 2.0, STORIES, and Waddington-OT are shown only for gene state because they do not provide benchmark spatial coordinates. Each target–space Sliced-W2 value was averaged over five shared random-projection repeats before ranking or comparison. The input data, interpolation procedures and shared evaluation settings are described in Supplementary Note S4.8.

#### Supplementary Figure S8. Five-dataset leave-one-time-point-out benchmark

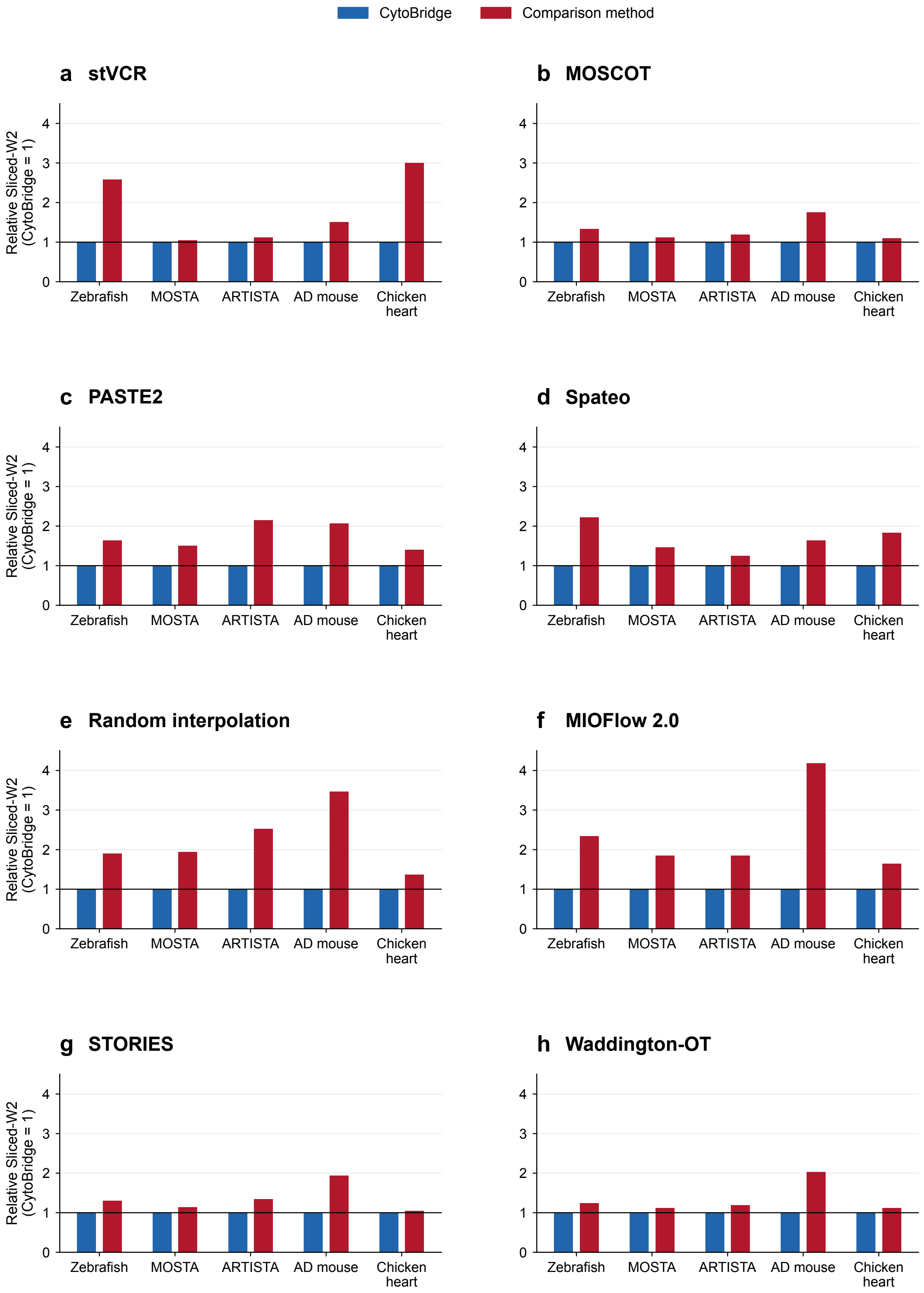

**Supplementary Figure S8.** Five-dataset leave-one-time-point-out benchmark. (a–h) Dataset-level Sliced-W2 relative to CytoBridge for stVCR, MOSCOT, PASTE2, Spateo, random interpolation, MIOFlow 2.0, STORIES, and Waddington-OT, respectively. CytoBridge is fixed at 1 and values above 1 indicate higher reconstruction error. Ratios were calculated within each matched dataset, held-out target, and evaluation space, and every matched target–space ratio contributed equally to its dataset mean. Panels a–e include joint, gene-state, and physical-space comparisons. Panels f–h include gene-state comparisons only because MIOFlow 2.0, STORIES, and Waddington-OT do not output benchmark spatial coordinates. CytoBridge had lower Sliced-W2 in 28 of 33 matched comparisons with stVCR, 27 of 33 with MOSCOT, 23 of 33 with PASTE2, 32 of 33 with Spateo, and 29 of 33 with random interpolation. The corresponding count was 11 of 11 for MIOFlow 2.0 and 9 of 11 for each of STORIES and Waddington-OT. All methods used the same 5,000 initial particles. The prediction and evaluation procedures are described in Supplementary Note S4.8. Each target–space value is the mean of five shared random-projection repeats, which measure numerical variation in Sliced-W2.

#### Supplementary Figure S9. Ligand–receptor-prior and interaction ablations across five spatial datasets

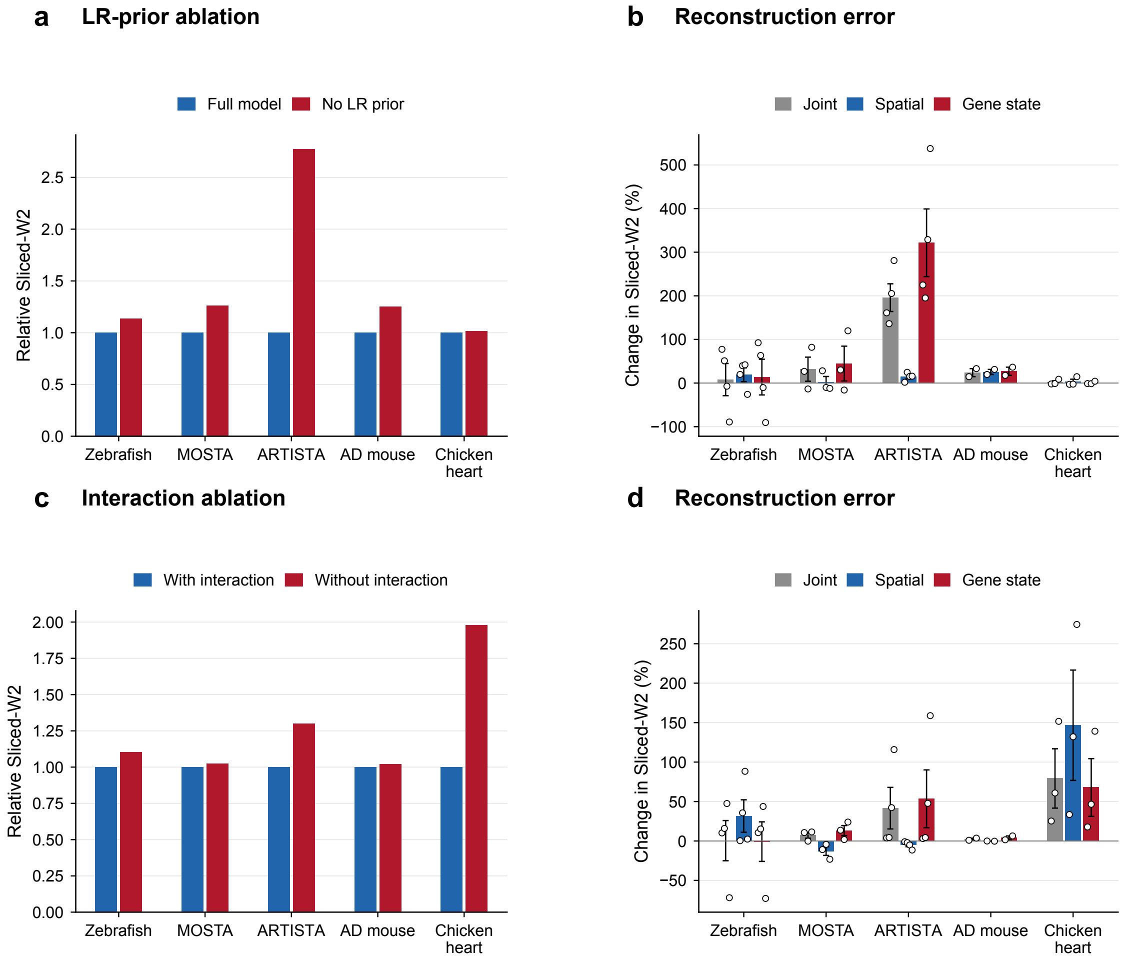

**Supplementary Figure S9.** Ligand–receptor-prior and interaction ablations across five spatial datasets. (a) Dataset-level reconstruction error for independently trained Full and No-LR-prior CytoBridge models, normalized to Full. The interaction network and spatial cutoff are retained in both models. (b) The same LR-prior comparison shown separately for each evaluation space. (c) Reconstruction error with and without the interaction-network output during inference, normalized to the model with interaction. Both conditions use the same trained model and 5,000 initial particles. (d) The inference-time comparison shown separately for each evaluation space. Growth is retained through continuous particle weights without resampling. In panels a and c, paired error ratios are averaged equally across target stages and evaluation spaces within each dataset. In panels b and d, bars show means across target stages, error bars show s.e.m., and white points show individual target stages. Positive changes indicate higher error in the ablated condition. Panels a and b use the same training seed for both models. Panels c and d average the paired ratios from three inference seeds within each target and space. All models were trained on all measured stages. Simulation and distance calculations are described in Supplementary Note S4.8.

### Supplementary Figure S10. Stability of the interaction and intrinsic-context decomposition in zebrafish

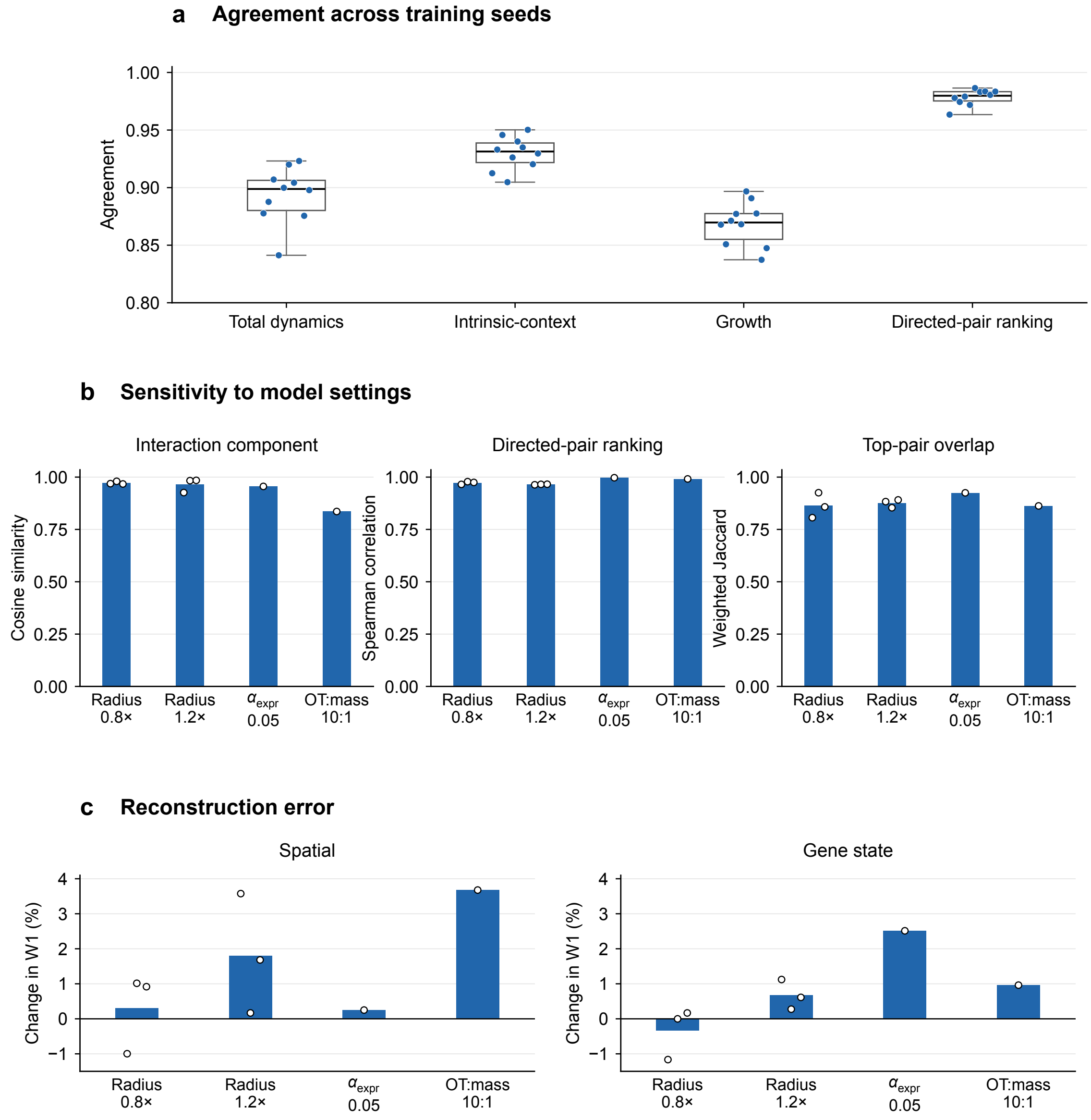

**Supplementary Figure S10.** Stability of the interaction and intrinsic-context decomposition in zebrafish. (a) Agreement among five independently trained default models with different random seeds. Boxes span the 25th–75th percentiles, central lines mark medians, and whiskers extend to the most extreme values within 1.5 interquartile ranges. Points show all ten pairwise comparisons, each evaluated on the same 11,999 observed cells at five model times. Agreement in the total dynamics and intrinsic-context component is measured by the median cosine similarity across cells in the 50-dimensional expression-state space. Growth agreement is the Spearman correlation across cells. Interaction agreement is the Spearman correlation between rankings of sender–receiver cell-type pairs calculated from the interaction messages on graph edges. (b) Stability after changing one model setting at a time. The left panel shows cosine similarity between the interaction vectors in expression-state space. The middle panel compares the rankings of sender–receiver cell-type pairs by Spearman correlation. The right panel shows weighted overlap among the highest-ranked 20% of pairs with nonzero contributions. The neighborhood cutoff was set to 0.8 or 1.2 times its default value in three matched training seeds. The expression-loss weight of 0.05 and transport-to-mass ratio of 10:1 each used one training seed. Each model is compared with the default model trained with the same seed. (c) Percentage change in mean Wasserstein-1 distance from the initial population to observed target times 1–4 after changing one model setting. Lower values indicate better reconstruction. In panels b and c, bars show means and open circles show individual runs. Spatial and expression-state distances are shown separately.

### Supplementary Figure S11. Cross-method comparison of CytoBridge interaction summaries

#### a Directed-pair consistency

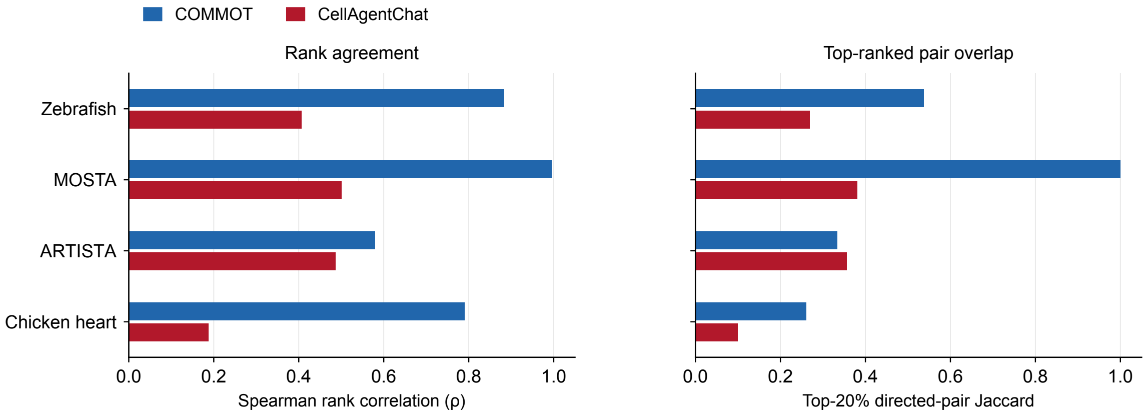

#### b CytoBridge-selected LR axes

| | CytoBridge selection | | | | COMMOT comparison | |
|---|---|---|---|---|---|---|
| **Dataset** | **Sender** | **Receiver** | **Selected LR axis** | **Database pathway** | **Same-pair LR rank** | **Pair percentile** |
| **Zebrafish** | Pronephros | Musculature / YSL | **COL1A2–SDC4** | COLLAGEN | 7 / 1,991 | 75.9 |
| **MOSTA** | Muscle | Connective tissue | **COL6A3–CD44** | COLLAGEN | 44 / 1,757 | 93.3 |
| **ARTISTA** | sfrpEGC | tINBL | **PSAP–GPR37L1** | PSAP | 2 / 480 | 94.2 |
| **Chicken heart** | Fibroblast cells | Valve cells | **CD99–CD99** | CD99 | 2 / 1,105 | 96.5 |

#### c NicheNet receiver targets for CytoBridge-selected axes

| | CytoBridge selection | | COMMOT and NicheNet results | |
|---|---|---|---|---|
| **Dataset** | **Directed pair (sender; receiver)** | **LR axis Database pathway** | **COMMOT same-axis percentile** | **NicheNet-predicted receiver targets** |
| **Zebrafish** | Sender: Spinal Cord Anterior Region<br>Receiver: Nervous System | **CDH2–CDH2**<br>CDH | 99.2 | Ccnd1, Cbx5, Cdk1 |
| **MOSTA** | Sender: Muscle<br>Receiver: Connective tissue | **COL6A3–CD44**<br>COLLAGEN | 99.3 | Cd44 |
| **ARTISTA** | Sender: sstIN<br>Receiver: mpEX | **L1CAM–L1CAM**<br>L1CAM | 99.1 | EGR1, DDIT4, JUNB |
| **Chicken heart** | Sender: Valve cells<br>Receiver: Fibroblast cells | **LAMB1–DAG1**<br>LAMININ | 99.9 | ID1, FN1, HNRNPR |

**Supplementary Figure S11.** Cross-method comparison of CytoBridge interaction summaries. (a) Spearman rank correlations compare the complete rankings of sender–receiver cell-type pairs in each dataset, using CytoBridge, COMMOT and a CellAgentChat calculation with the same LR database. Jaccard indices compare the corresponding top 20% of directed pairs. (b) Representative ligand–receptor axes selected from the CytoBridge interaction contribution before external comparison. The COMMOT columns report the rank of the same ligand–receptor axis within the selected directed pair and the percentile of that directed pair. (c) For each dataset, we show the combination of an LR pair and sender–receiver cell types ranked highest by CytoBridge for which NicheNet reports a positive ligand–target link. The table gives the COMMOT percentile for that combination and the predicted target genes in receiver cells. For zebrafish, genes were mapped to one-to-one mouse orthologs in Ensembl and analyzed using the mouse NicheNet prior.

### Supplementary Figure S12. Sensitivity of chicken-heart alignment and downstream results to small coordinate perturbations

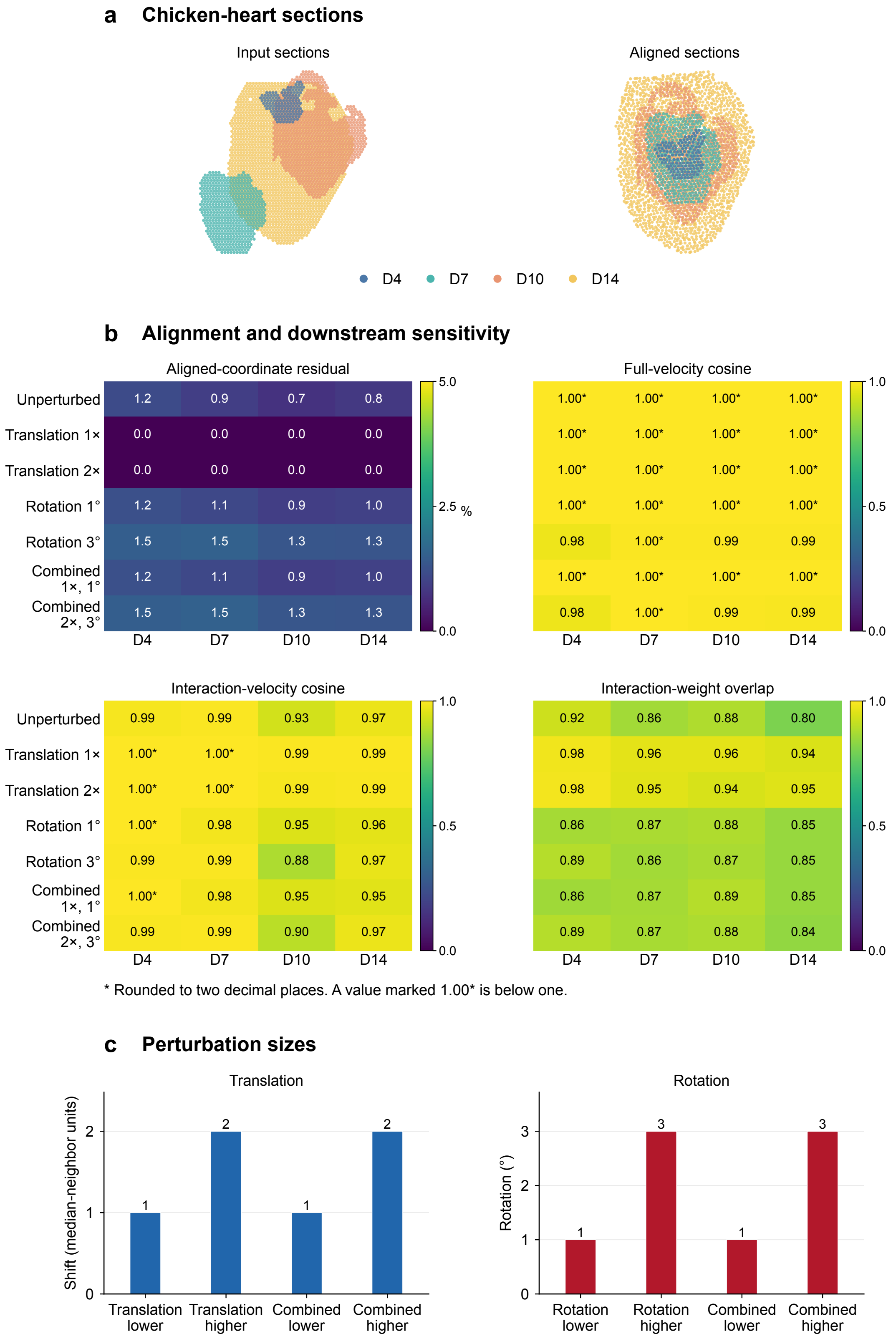

Aligned-coordinate residual (%)

| | D4 | D7 | D10 | D14 |
|---|---|---|---|---|
| Unperturbed | 1.2 | 0.9 | 0.7 | 0.8 |
| Translation 1× | 0.0 | 0.0 | 0.0 | 0.0 |
| Translation 2× | 0.0 | 0.0 | 0.0 | 0.0 |
| Rotation 1° | 1.2 | 1.1 | 0.9 | 1.0 |
| Rotation 3° | 1.5 | 1.5 | 1.3 | 1.3 |
| Combined 1×, 1° | 1.2 | 1.1 | 0.9 | 1.0 |
| Combined 2×, 3° | 1.5 | 1.5 | 1.3 | 1.3 |

Full-velocity cosine

| | D4 | D7 | D10 | D14 |
|---|---|---|---|---|
| Unperturbed | 1.00* | 1.00* | 1.00* | 1.00* |
| Translation 1× | 1.00* | 1.00* | 1.00* | 1.00* |
| Translation 2× | 1.00* | 1.00* | 1.00* | 1.00* |
| Rotation 1° | 1.00* | 1.00* | 1.00* | 1.00* |
| Rotation 3° | 0.98 | 1.00* | 0.99 | 0.99 |
| Combined 1×, 1° | 1.00* | 1.00* | 1.00* | 1.00* |
| Combined 2×, 3° | 0.98 | 1.00* | 0.99 | 0.99 |

Interaction-velocity cosine

| | D4 | D7 | D10 | D14 |
|---|---|---|---|---|
| Unperturbed | 0.99 | 0.99 | 0.93 | 0.97 |
| Translation 1× | 1.00* | 1.00* | 0.99 | 0.99 |
| Translation 2× | 1.00* | 1.00* | 0.99 | 0.99 |
| Rotation 1° | 1.00* | 0.98 | 0.95 | 0.96 |
| Rotation 3° | 0.99 | 0.99 | 0.88 | 0.97 |
| Combined 1×, 1° | 1.00* | 0.98 | 0.95 | 0.95 |
| Combined 2×, 3° | 0.99 | 0.99 | 0.90 | 0.97 |

Interaction-weight overlap

| | D4 | D7 | D10 | D14 |
|---|---|---|---|---|
| Unperturbed | 0.92 | 0.86 | 0.88 | 0.80 |
| Translation 1× | 0.98 | 0.96 | 0.96 | 0.94 |
| Translation 2× | 0.98 | 0.95 | 0.94 | 0.95 |
| Rotation 1° | 0.86 | 0.87 | 0.88 | 0.85 |
| Rotation 3° | 0.89 | 0.86 | 0.87 | 0.85 |
| Combined 1×, 1° | 0.86 | 0.87 | 0.89 | 0.85 |
| Combined 2×, 3° | 0.89 | 0.87 | 0.88 | 0.84 |

**Supplementary Figure S12.** Sensitivity of chicken-heart alignment and downstream results to small coordinate perturbations. (a) Input and aligned spatial coordinates for D4, D7, D10, and D14. (b) Coordinate residuals, full- and interaction-velocity agreement, and interaction-weight overlap after translation, rotation, or combined perturbations. Each perturbed run is compared with the unperturbed repeat. The unperturbed repeat is compared with the original model. Coordinate residuals are normalized by the reference section radius after a proper rigid-frame adjustment. Velocity agreement is the median cellwise cosine after the same frame adjustment. Interaction-weight overlap is the normalized weighted Jaccard index over the union of directed edges. Values are rounded for display. An asterisk marks a value below one displayed as 1.00. (c) Applied translations of 1 and 2 median nearest-neighbor distances and rotations of 1° and 3°.

### Supplementary Figure S13. Chicken-heart alignment after small coordinate perturbations

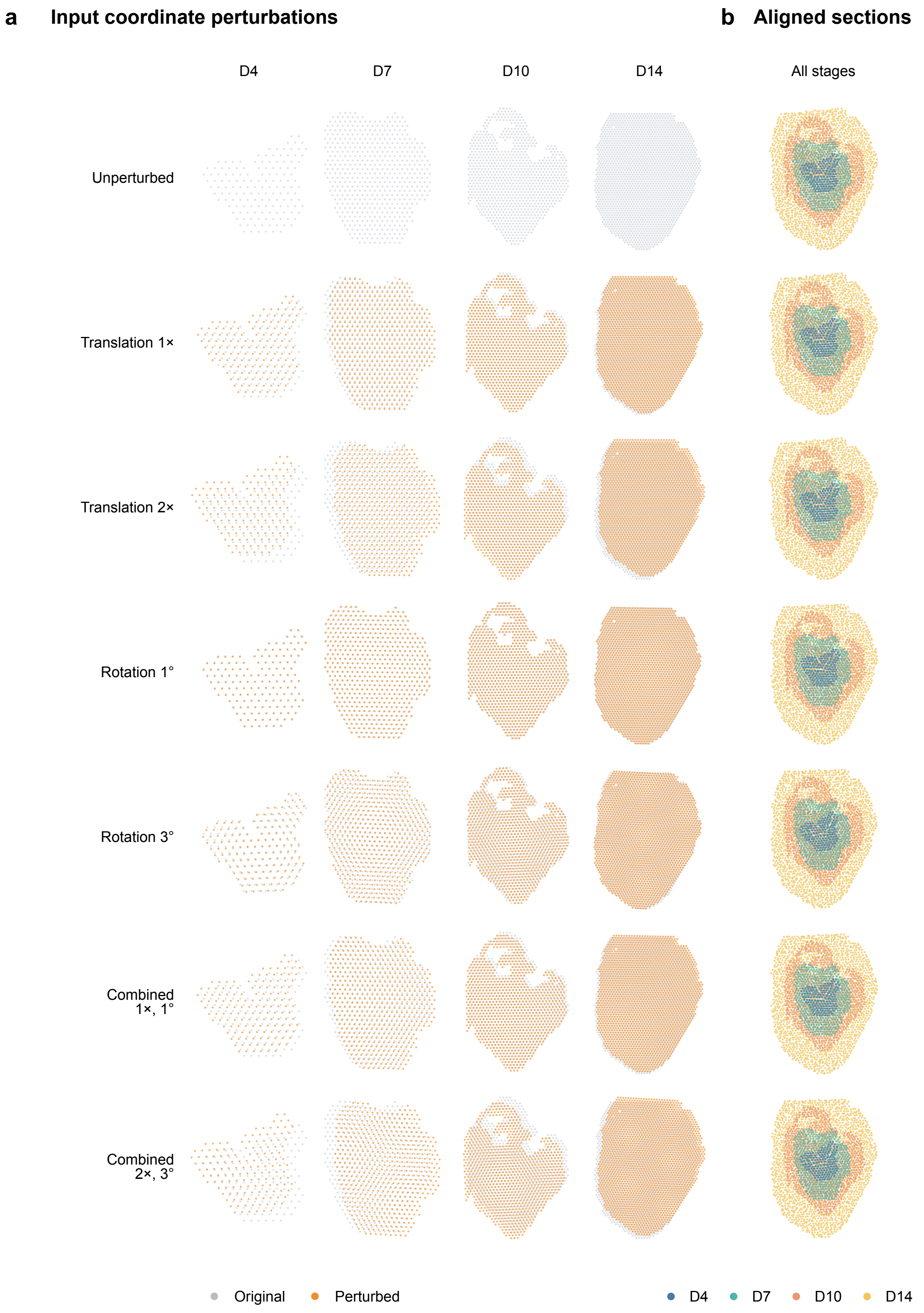

**Supplementary Figure S13.** Chicken-heart alignment after small coordinate perturbations. (a) Original input coordinates in gray and perturbed coordinates in orange at D4, D7, D10, and D14. Rows show the unperturbed input, translations of 1 and 2 times the within-section median nearest-neighbor distance, rotations of 1° and 3°, and combinations of 1× with 1° and 2× with 3°. (b) Aligned sections for each condition. Colors denote developmental time.

### Supplementary Figure S14. Growth estimates during chicken-heart development

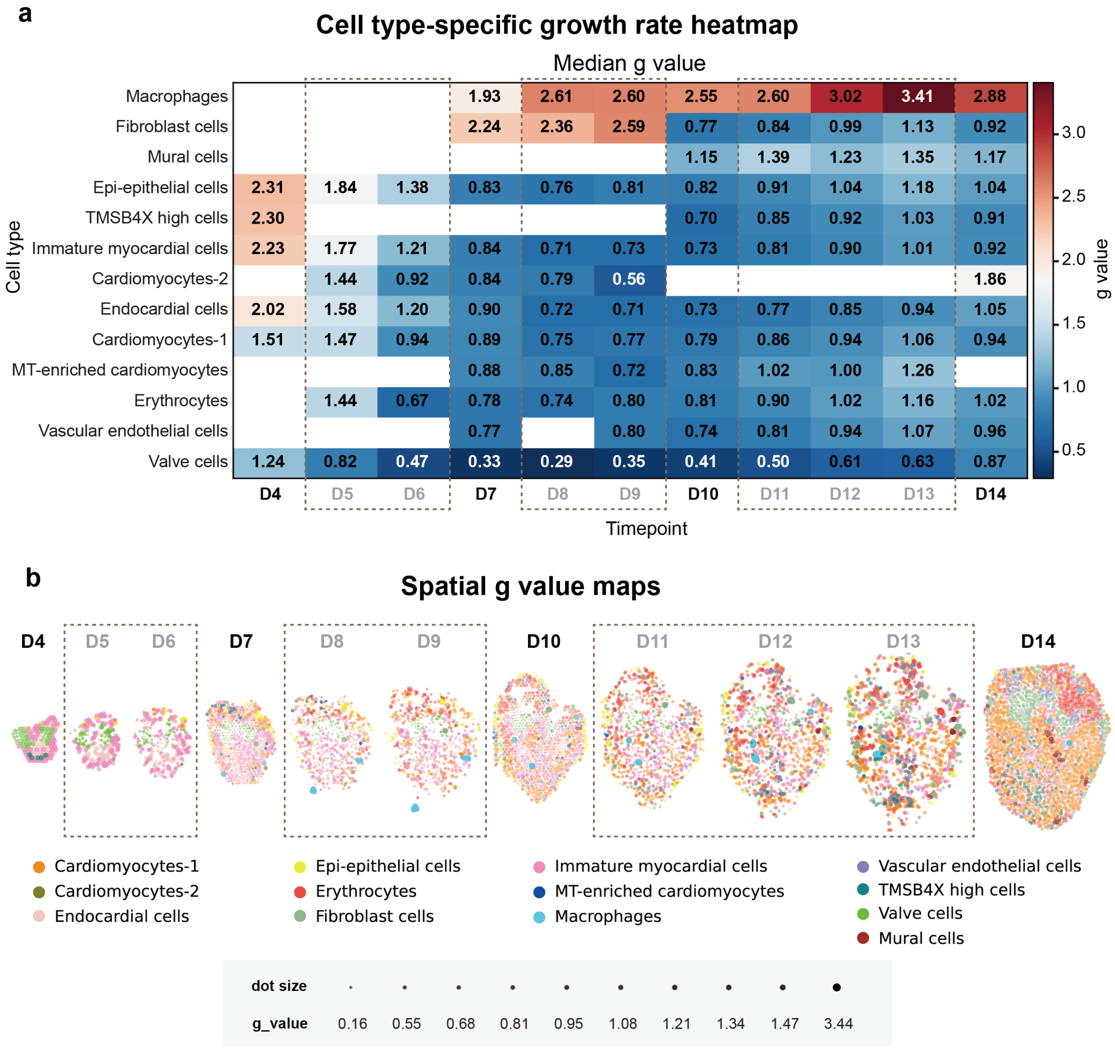

**Supplementary Figure S14.** Growth estimates during chicken-heart development. (a) Median model growth value for each cell type at observed and generated stages. (b) Spatial maps of observed and generated stages; color denotes cell type and point size denotes the model growth value.

### Supplementary Figure S15. Velocity components in the developing chicken heart

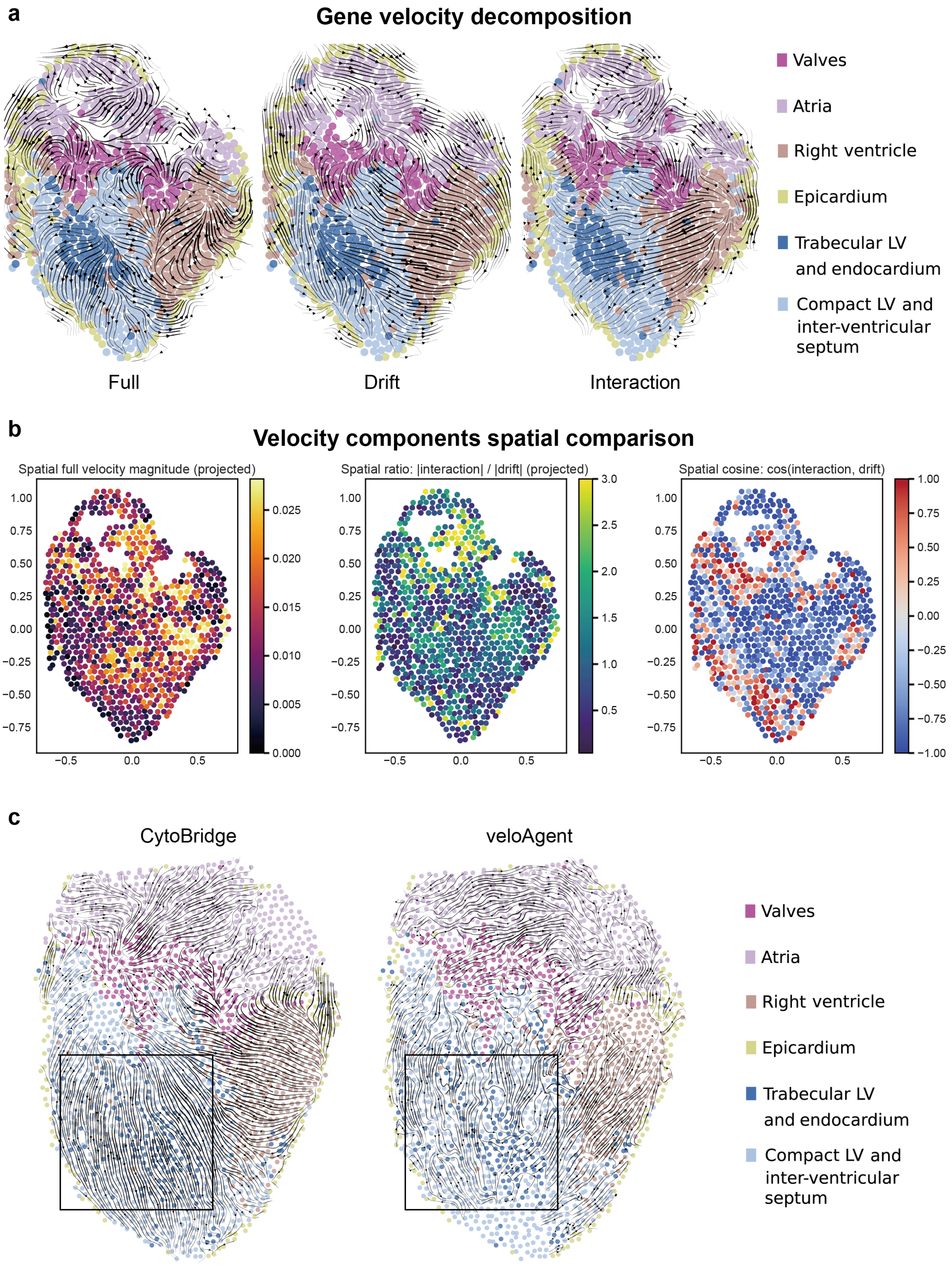

**Supplementary Figure S15.** Velocity components in the developing chicken heart. (a) Full, drift, and interaction components of the gene-expression velocity at D10, projected onto tissue coordinates. Full denotes the sum of the probability-flow, interaction and score contributions. Cells are colored by anatomical region. (b) Spatial distributions of projected full-velocity magnitude, the interaction-to-drift magnitude ratio, and cosine similarity between the interaction and drift components. (c) Spatial velocity streamlines from CytoBridge and veloAgent at D14. Black boxes mark the corresponding lower-left ventricular region, where both fields show flow from the trabecular LV toward the compact LV. LV, left ventricle.

### Supplementary Figure S16. Generated spatial maps across the MOSTA trajectory

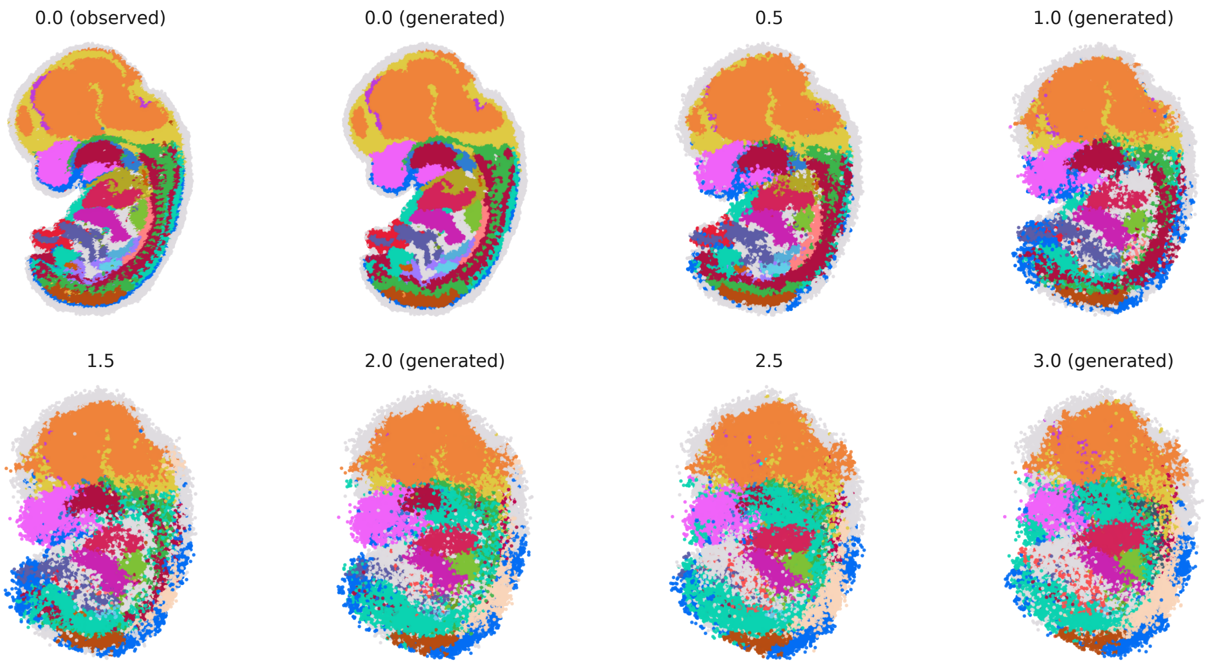

**Supplementary Figure S16.** Generated spatial maps across the MOSTA trajectory. The leftmost map is the observed $t = 0$ tissue. It is followed by generated states at $t = 0, 0.5, 1, 1.5, 2, 2.5$, and 3 from the same continuous simulation from $t = 0$. Colors indicate annotated cell populations.

#### Supplementary Figure S17. Growth-rate maps across the MOSTA trajectory

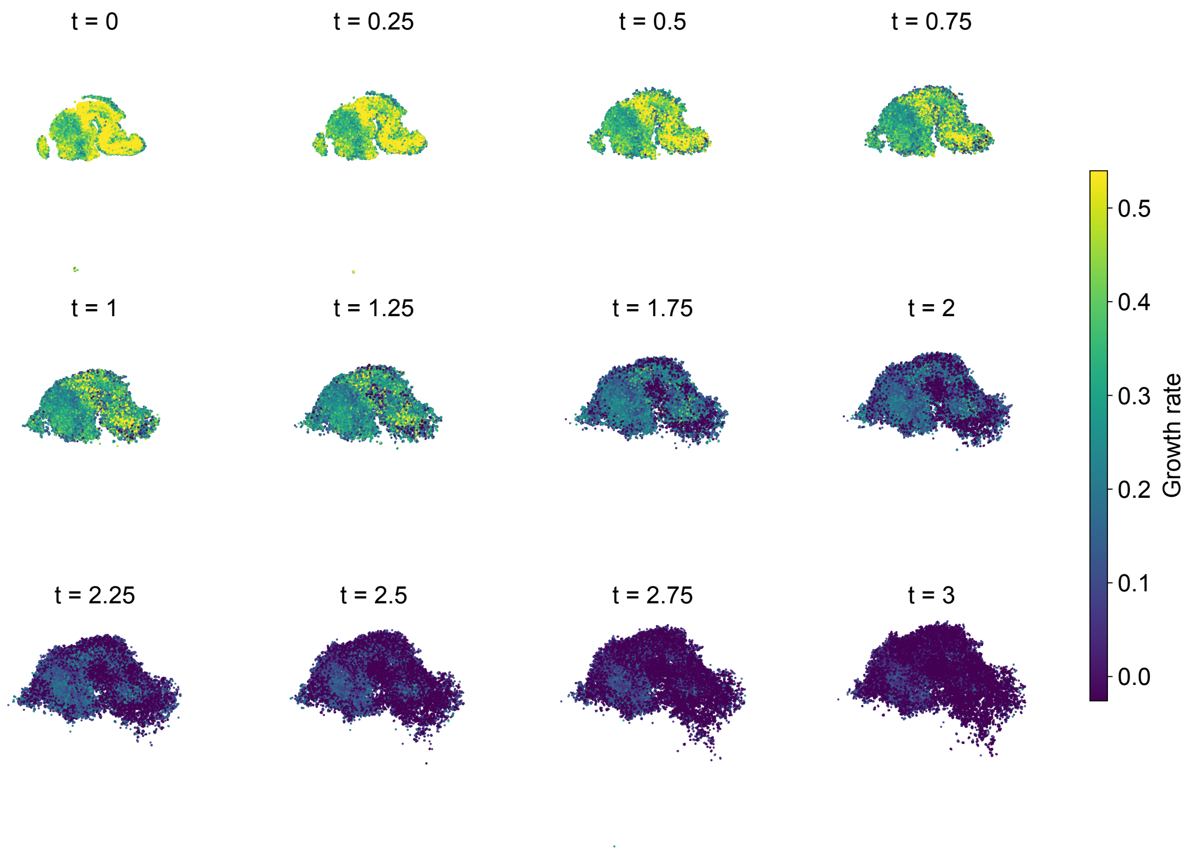

**Supplementary Figure S17.** Growth-rate maps across the MOSTA trajectory. The model growth field was evaluated on classifier-assigned Brain cells at 13 quarter-step states from $t = 0$ to $t = 3$ generated along the same trajectory initialized at $t = 0$. Twelve maps are displayed; $t = 1.50$ was omitted from the display. Colors report direct per-cell growth predictions on a common scale calculated from all 13 Brain slices.

### Supplementary Figure S18. Cell counts and composition across the MOSTA trajectory

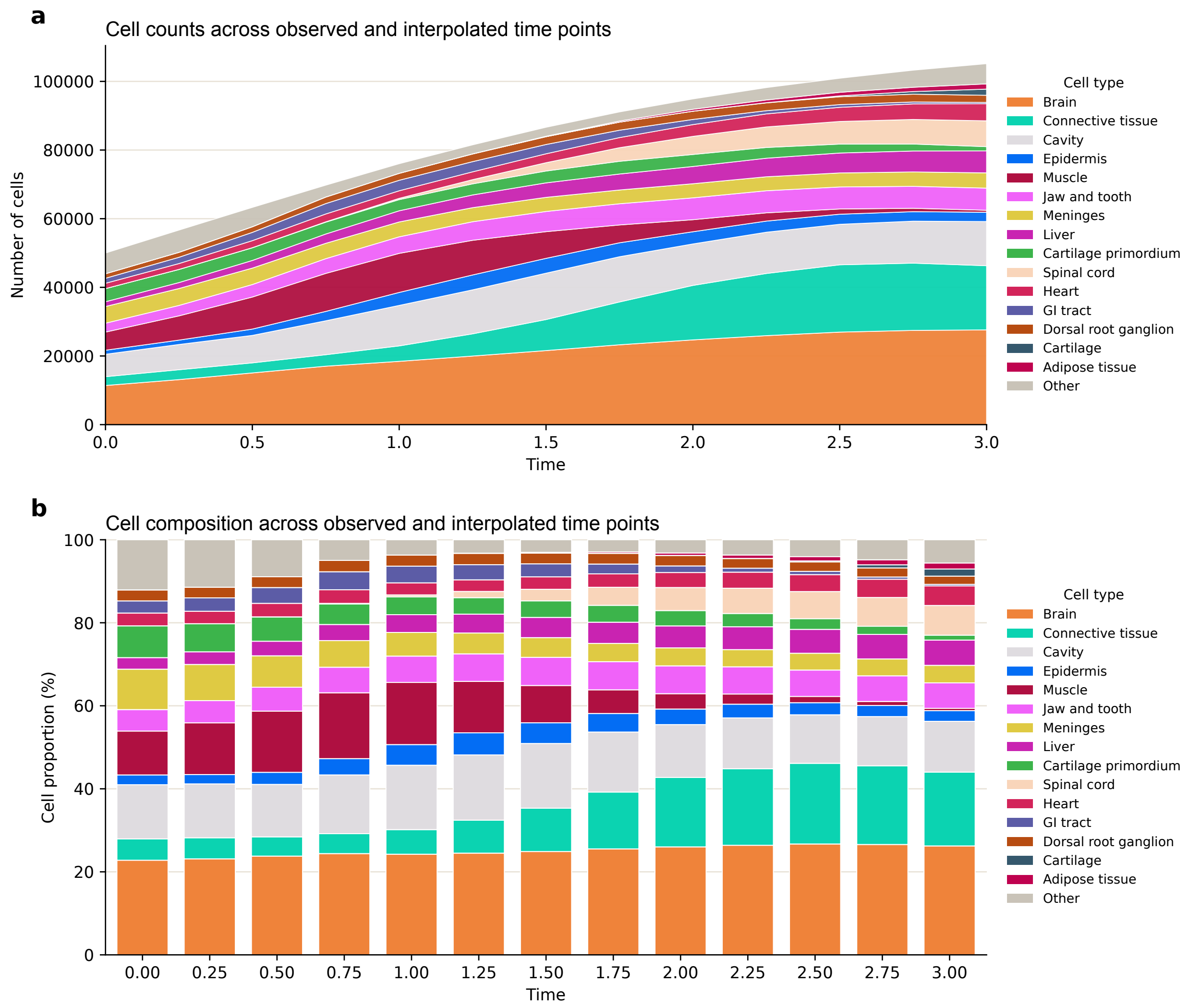

**Supplementary Figure S18.** Cell counts and composition across the MOSTA trajectory. (a) Absolute counts of annotated cell populations across 13 quarter-step states from $t = 0$ to $t = 3$. (b) The corresponding cell-type proportions.

### Supplementary Figure S19. Cell-state transitions across the MOSTA trajectory

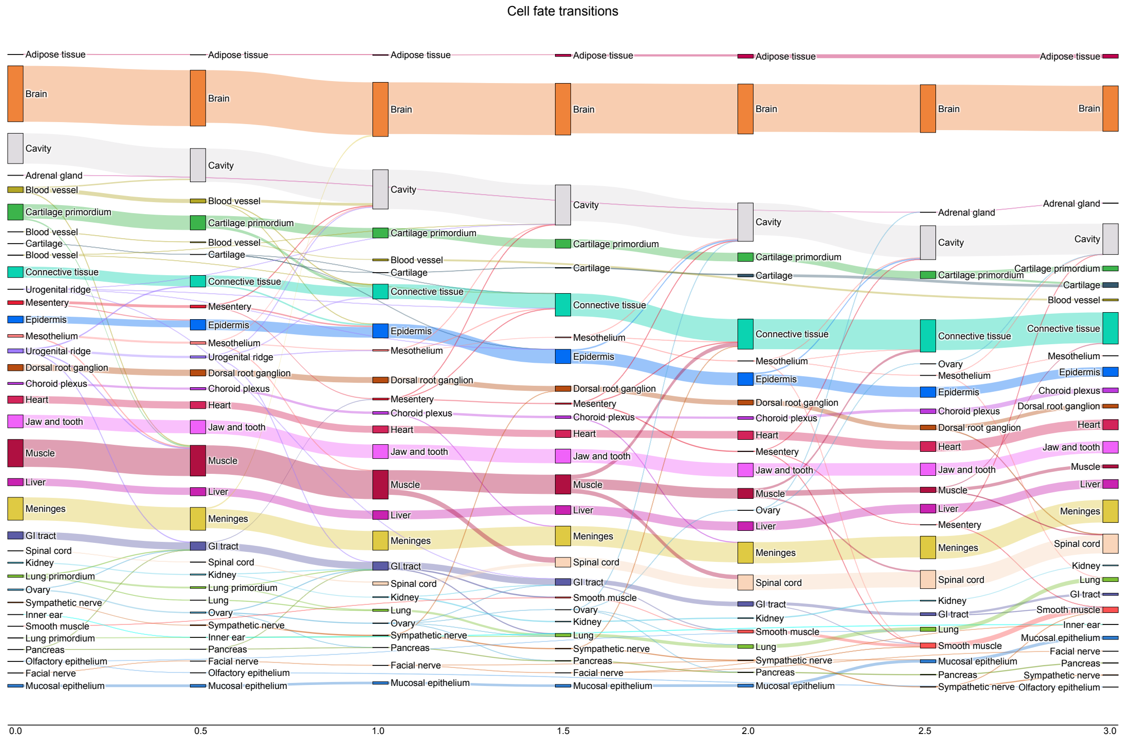

**Supplementary Figure S19.** Cell-state transitions across the MOSTA trajectory. Sankey diagram for 50,000 particles propagated from the initial stage without division. Cell types were assigned by the classifier. Persistent particle identities define transitions between adjacent half-step times. Ribbon widths report particle counts.

#### Supplementary Figure S20. Temporal gene programs in the MOSTA dataset

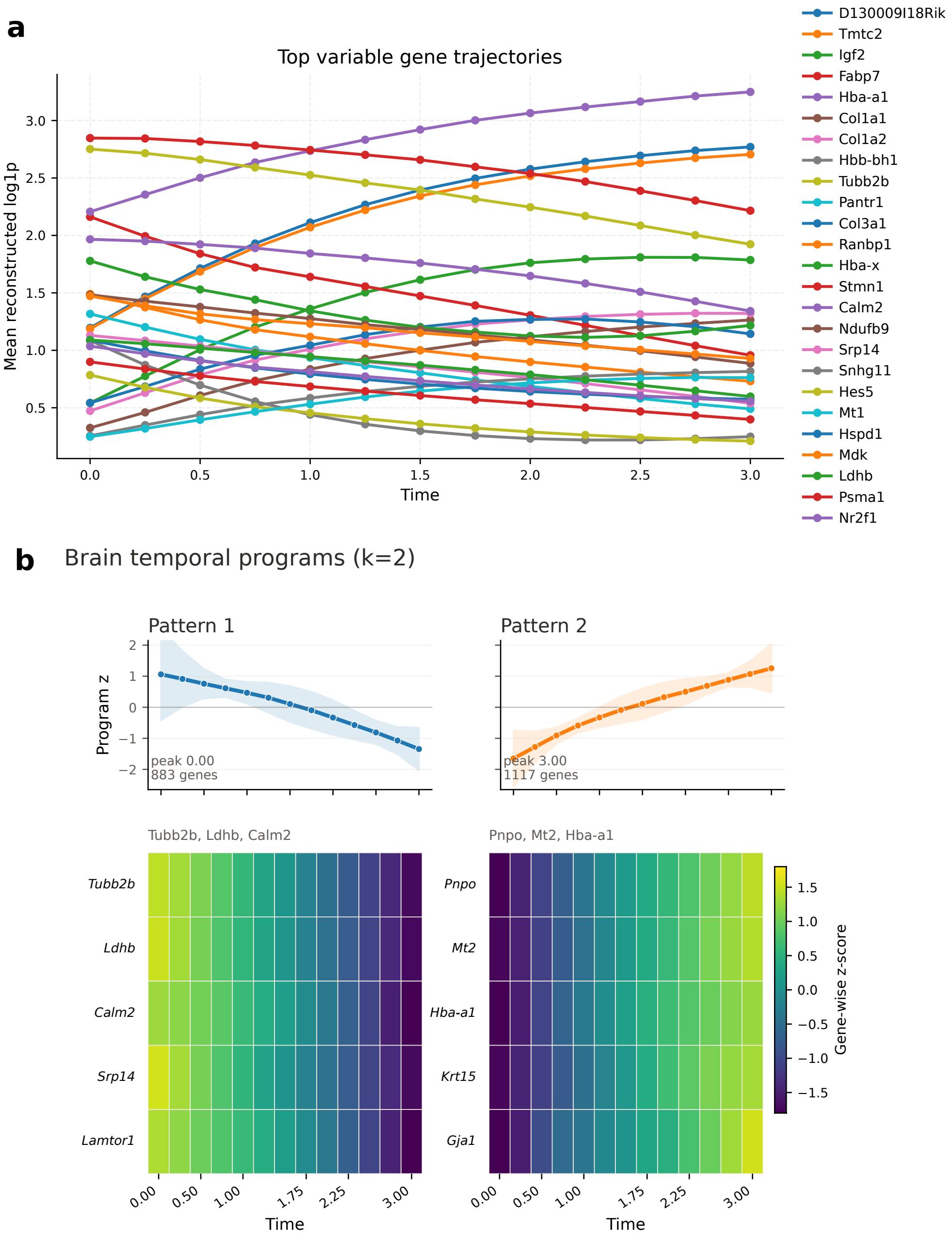

**Supplementary Figure S20.** Temporal gene programs in the MOSTA dataset. (a) Reconstructed trajectories of representative variable genes across the trajectory. (b) Two temporal programs identified from the row-normalized gene trajectories. Pattern 1 contains 883 genes and peaks at $t = 0$; Pattern 2 contains 1,117 genes and peaks at $t = 3$. Heat maps show representative gene-wise standardized profiles.

### Supplementary Figure S21. Functional enrichment of temporal gene programs in the MOSTA dataset

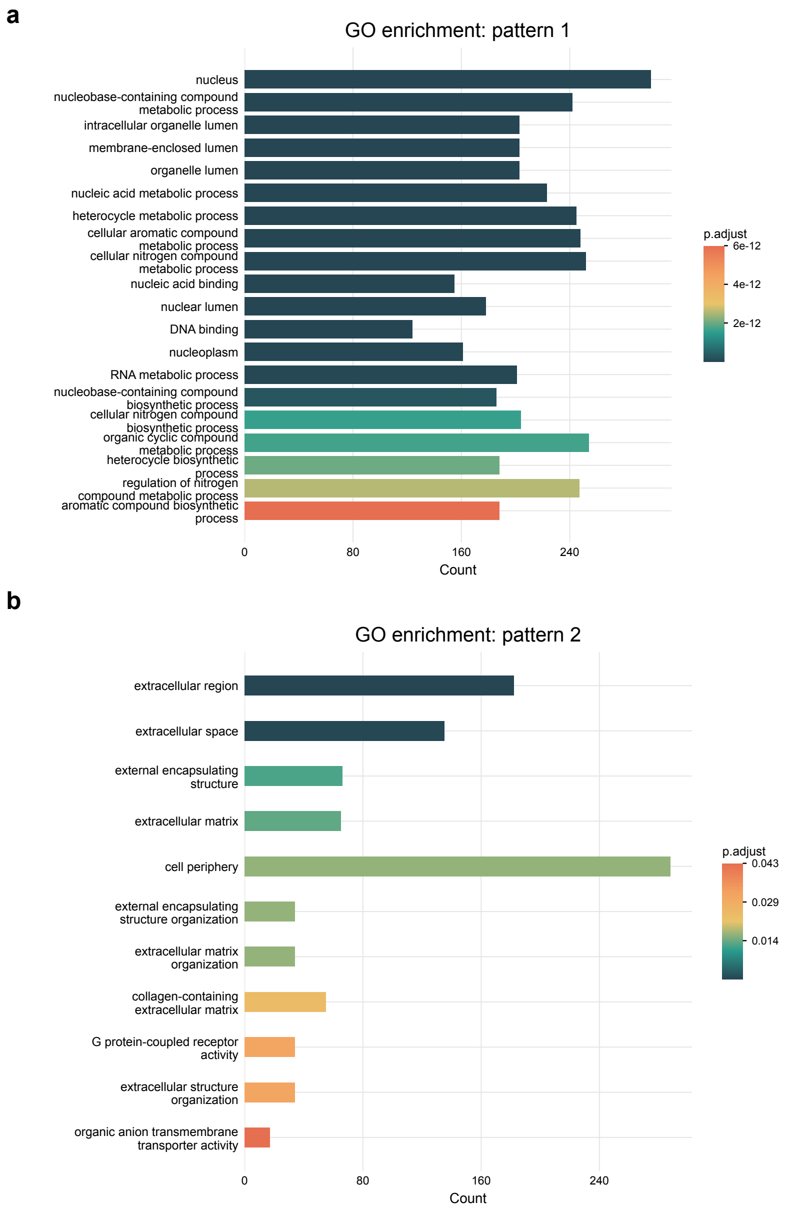

**Supplementary Figure S21.** Functional enrichment of temporal gene programs in the MOSTA dataset. Gene Ontology enrichment for the two temporal programs. Pattern 1 is enriched for nuclear and nucleic-acid metabolic terms. Pattern 2 has 11 terms passing the Benjamini–Hochberg-adjusted $P < 0.05$ threshold, mostly extracellular-region and extracellular-matrix terms.

### Supplementary Figure S22. Developmental waves in the MOSTA dataset

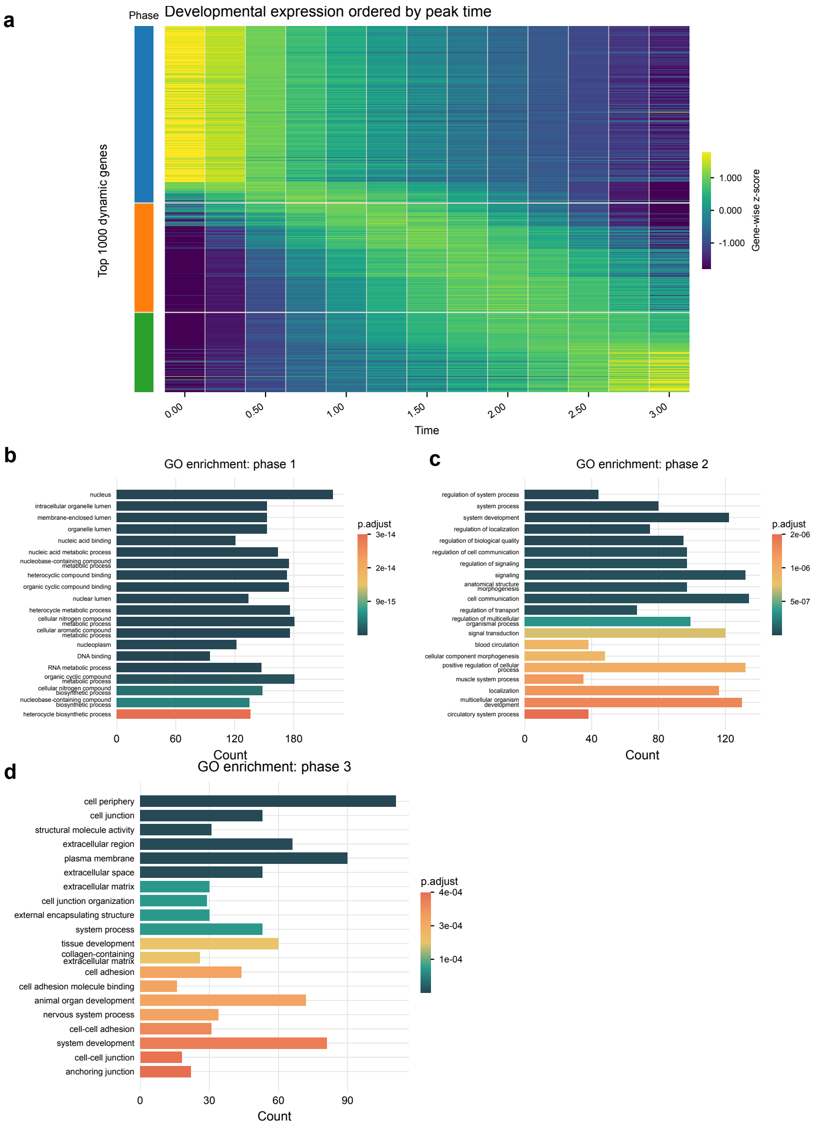

**Supplementary Figure S22.** Developmental waves in the MOSTA dataset. (a) Dynamic genes ordered by peak time and assigned to three temporal phases. (b) Phase 1 is enriched for nuclear and nucleic-acid metabolic terms. (c) Phase 2 is enriched for signaling, system-development and circulatory-system terms. (d) Phase 3 is enriched for cell-junction, extracellular-matrix and cell-adhesion terms.

#### Supplementary Figure S23. Ligand–receptor temporal patterns in the MOSTA dataset

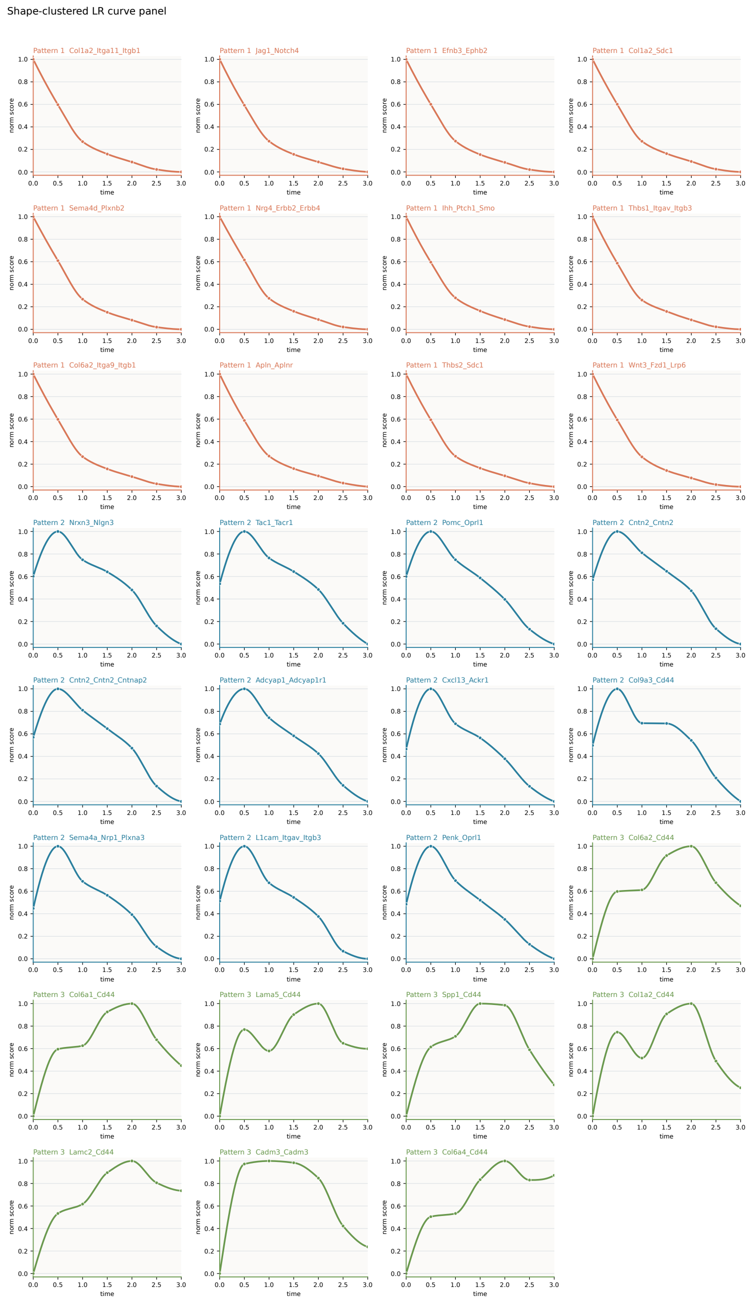

**Supplementary Figure S23.** Ligand–receptor temporal patterns in the MOSTA dataset. Ligand–receptor communication profiles were grouped by their changes over time using total cell-type communication ($M_{\text{sum}}$). Pattern 1 (orange) is strongest at the earliest stage and declines over time. Pattern 2 (blue) shows a broad early peak followed by a gradual decline. Pattern 3 (green) increases during intermediate-to-late stages and then plateaus or partially declines. Markers are the seven computed half-step values; lines are display-only interpolations.

#### Supplementary Figure S24. Observed and generated stages across the ARTISTA trajectory

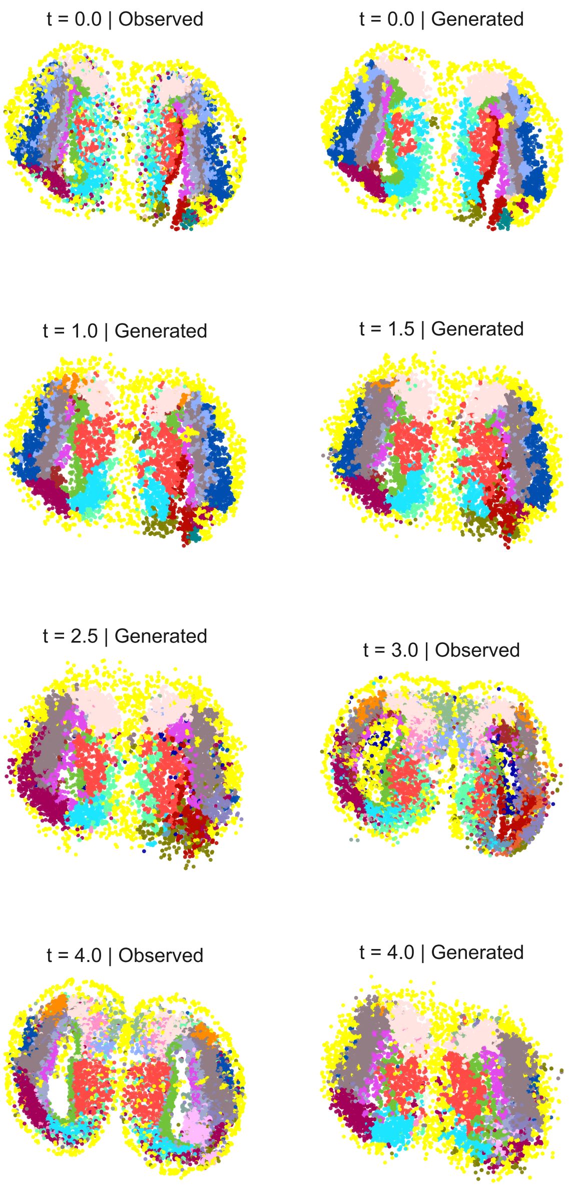

**Supplementary Figure S24.** Observed and generated stages across the ARTISTA trajectory. Observed maps at the five integer stages are shown alongside generated maps at the integer anchors and at $t = 0.5, 1.5, 2.5$, and 3.5. Generated stages are shown at their simulated spatial coordinates, colored by cell type.

### Supplementary Figure S25. Particle flows and composition across the ARTISTA trajectory

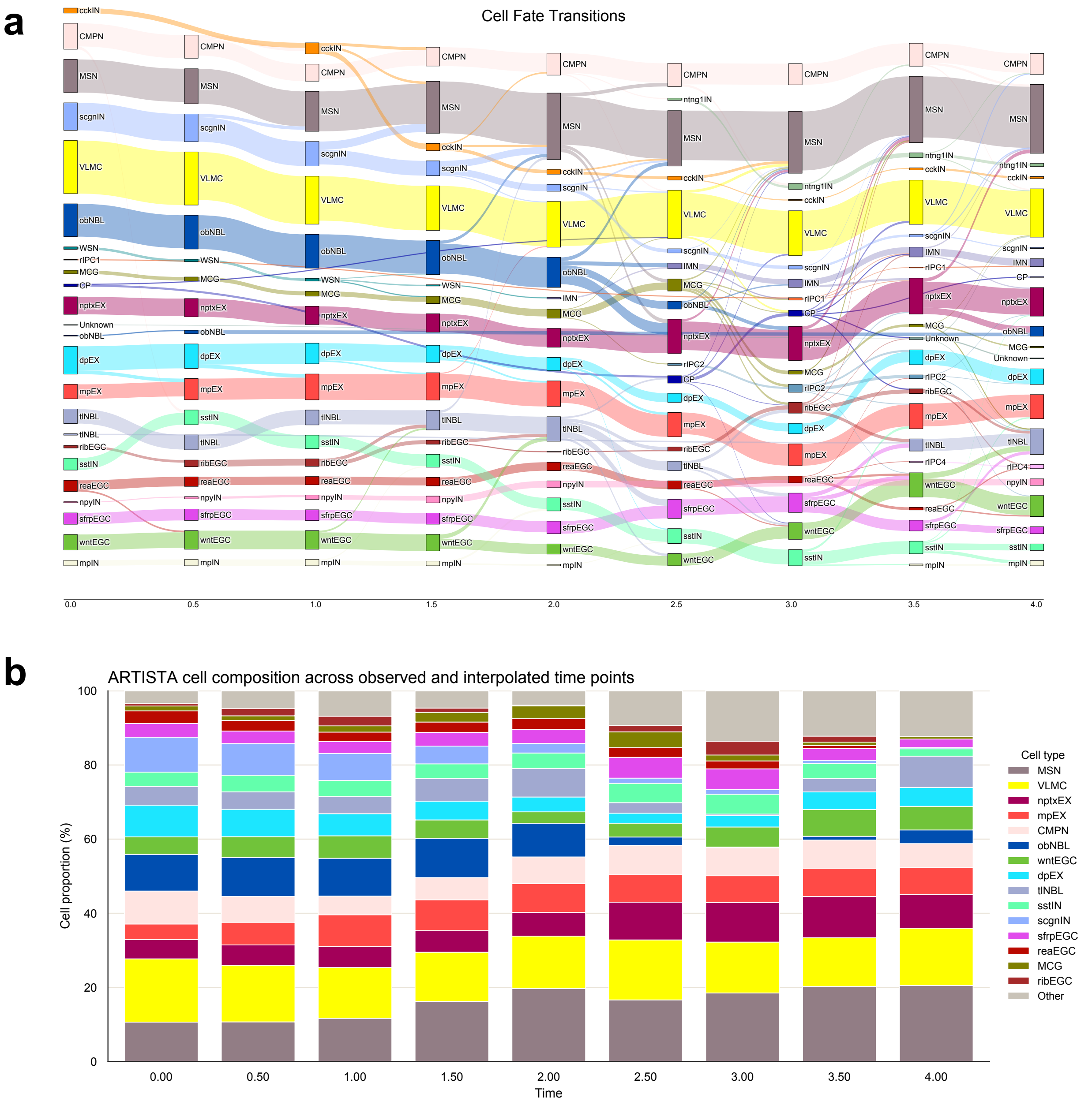

**Supplementary Figure S25.** Particle flows and composition across the ARTISTA trajectory. (a) Adjacent-time flows for 7,668 particles propagated without division across nine time points. Cell types were assigned by the classifier. Ribbon widths report particle counts. (b) Cell-type composition of the same particles, displayed as the 15 most abundant annotations plus Other.

### Supplementary Figure S26. Localized interaction domains in the ARTISTA 5-DPI field

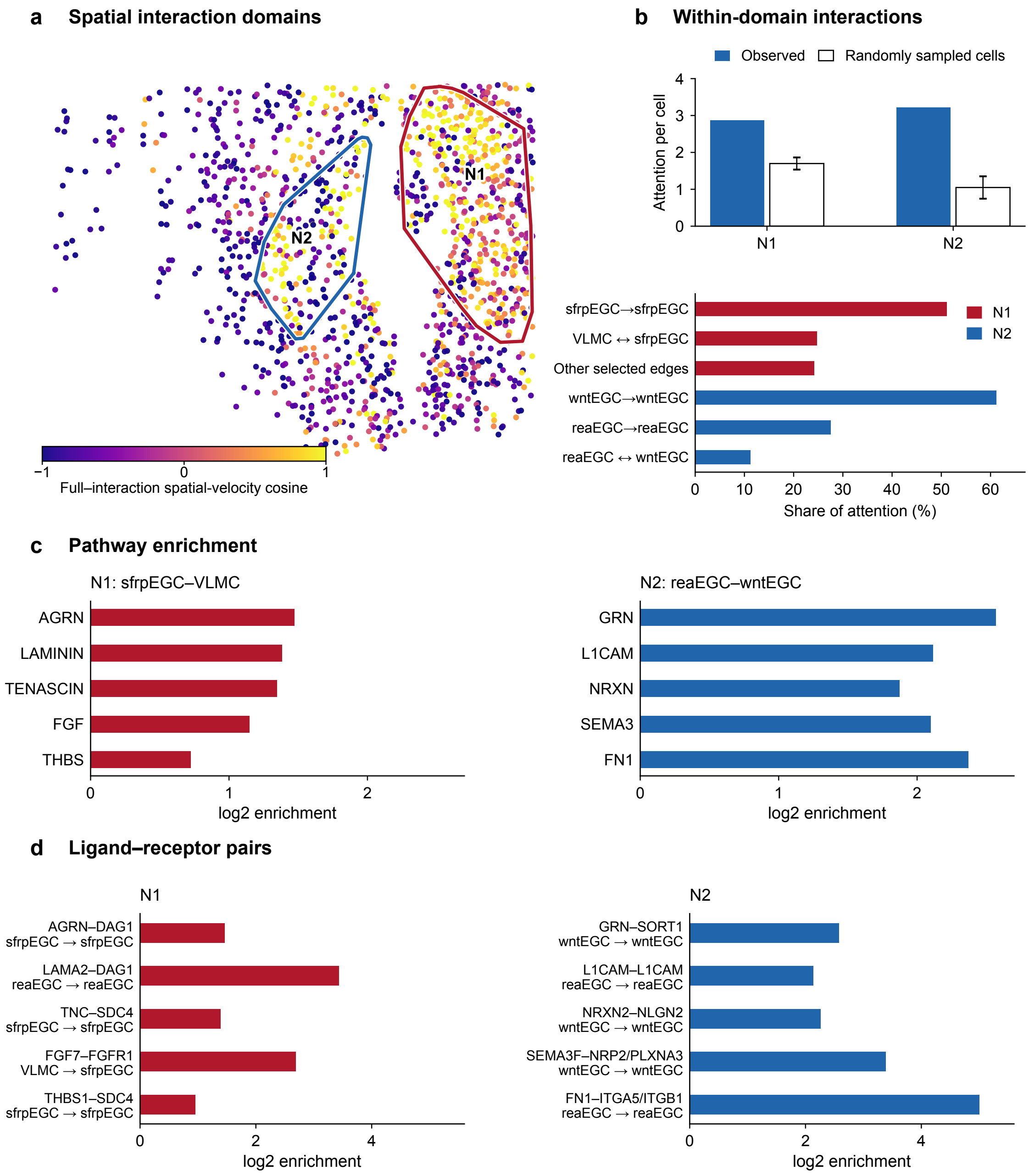

**Supplementary Figure S26.** Localized interaction domains in the ARTISTA 5-DPI field. (a) Cosine similarity between the full spatial velocity and its interaction component in the 5-DPI region shown in Fig. 5c. Two domains were defined by connecting cells in the upper quartile of cosine similarity as described in Supplementary Note S8.10. (b) Attention per domain cell compared with random cell sets from the same region and with the same cell-type composition. Filled bars show the observed values, and open bars and error bars show the random-set means and s.d. The sfrpEGC–VLMC- and reaEGC–wntEGC-associated domains showed 1.69-fold and 3.06-fold enrichment, respectively (both empirical $P = 0.0001$). Dominant sender–receiver structures summarize selected attention within each domain. (c) Representative ligand–receptor pathway scores at 5 DPI relative to the random cell sets. All displayed pathways passed Benjamini–Hochberg correction at $q < 0.05$. (d) Candidate ligand–receptor pairs within each domain. The highest-scoring directed pair is shown per pathway in (c) (for the five displayed N2 pathways, the top-scoring pairs are homotypic wntEGC or reaEGC pairs). All displayed pairs passed BH correction across 531 tested pairs within their domain. Bars show enrichment relative to the random cell sets.

### Supplementary Figure S27. Temporal gene programs and GO analysis in the ARTISTA dataset

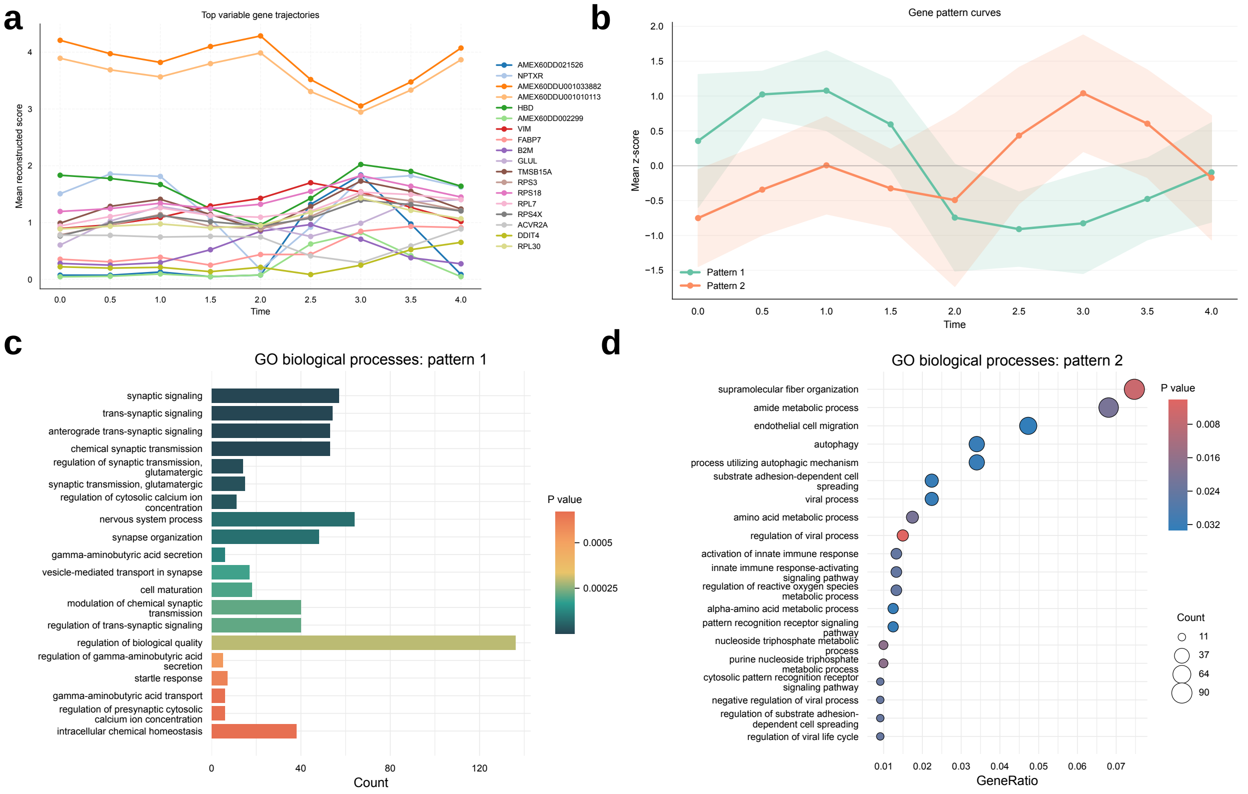

**Supplementary Figure S27.** Temporal gene programs and GO analysis in the ARTISTA dataset. Pattern 1 contains 421 raw genes (388 Entrez-mapped) and has 10 Gene Ontology terms significant after Benjamini–Hochberg correction, including synaptic signaling, glutamatergic synaptic transmission and regulation of cytosolic calcium ion concentration. Pattern 2 contains 1,362 raw genes (1,235 Entrez-mapped). Its 20 displayed terms are ranked by raw $P$ value.

#### Supplementary Figure S28. Ligand–receptor communication pattern prototypes in the ARTISTA dataset

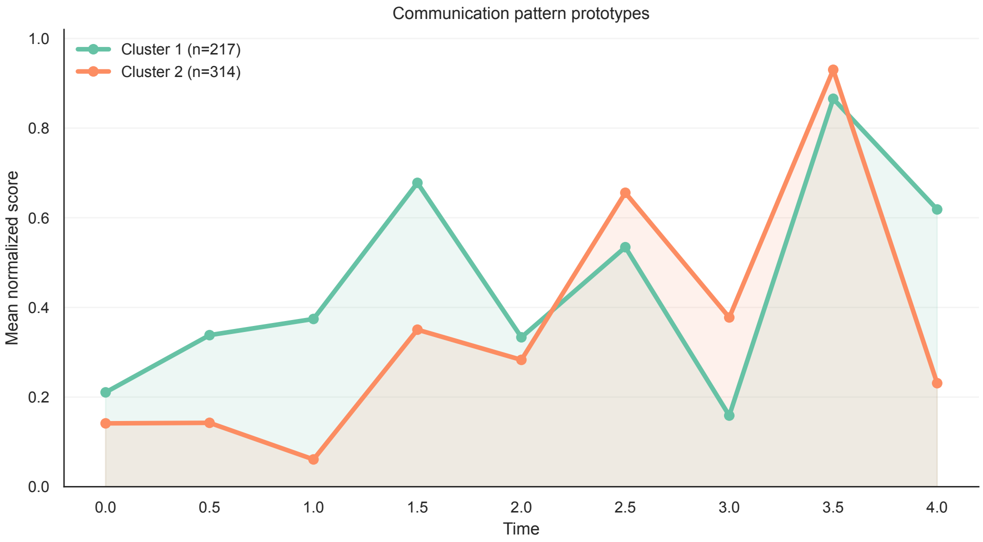

**Supplementary Figure S28.** Ligand–receptor communication pattern prototypes in the ARTISTA dataset. Row-wise min–max normalized profiles for 531 ligand–receptor pairs with all required subunits detected across nine time points were grouped by deterministic $k$-means. Among $k = 2$–$8$, $k = 2$ had the highest silhouette score; the two clusters contain 217 and 314 pairs. Curves show the corresponding cluster prototypes.

### Supplementary Figure S29. Representative ligand–receptor communication profiles in the ARTISTA dataset

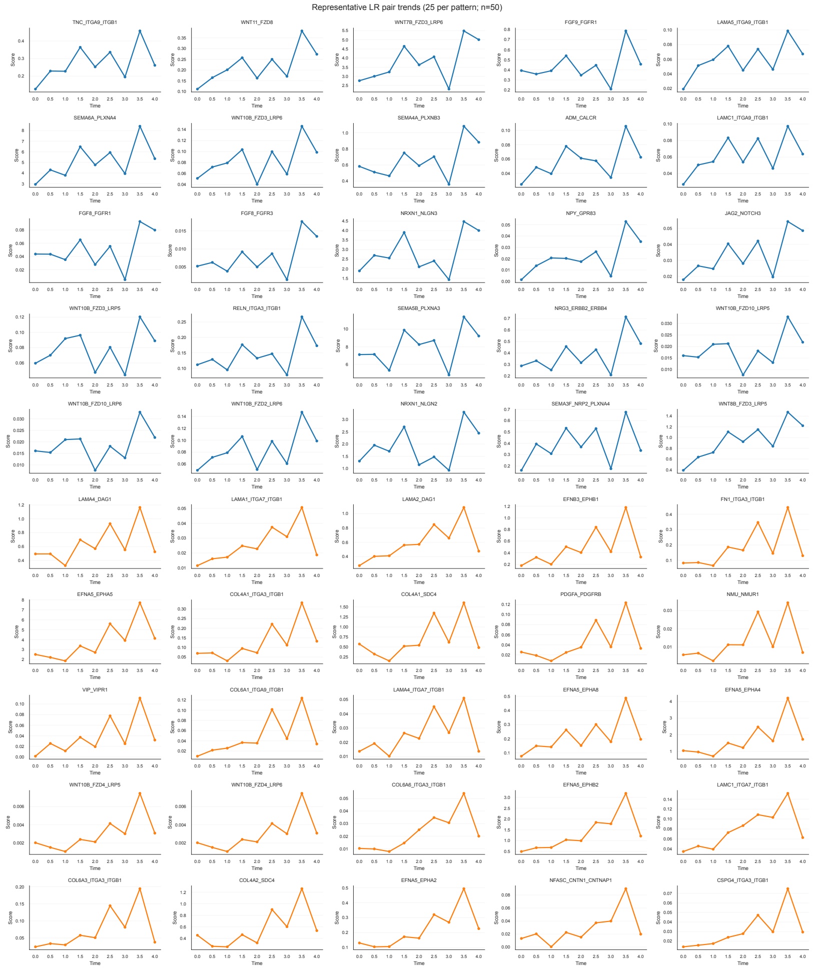

**Supplementary Figure S29.** Representative ligand–receptor communication profiles in the ARTISTA dataset. Fifty representative profiles are shown: the 25 pairs nearest to each of the two cluster prototypes after row-wise normalization. Scores require detection of all complex subunits.

### Supplementary Figure S30. Growth-rate maps across the ARTISTA trajectory

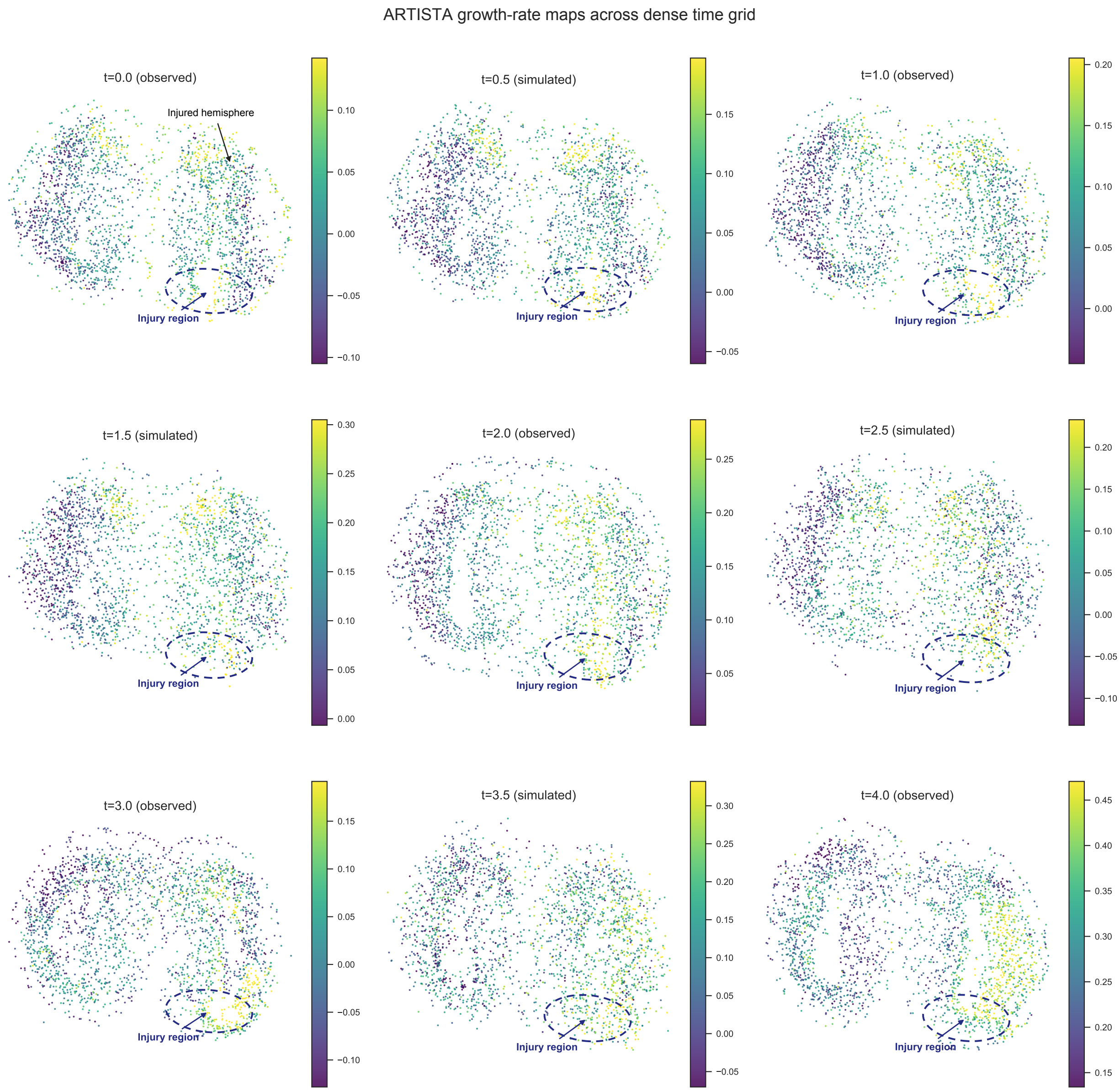

**Supplementary Figure S30.** Growth-rate maps across the ARTISTA trajectory. The learned growth field is shown at the five observed integer stages and four generated half steps. Each panel uses its own 5th–95th percentile color range. The dashed outline indicates the approximate injury region.

#### Supplementary Figure S31. Reconstructed AD progression in TgCRND8 mice

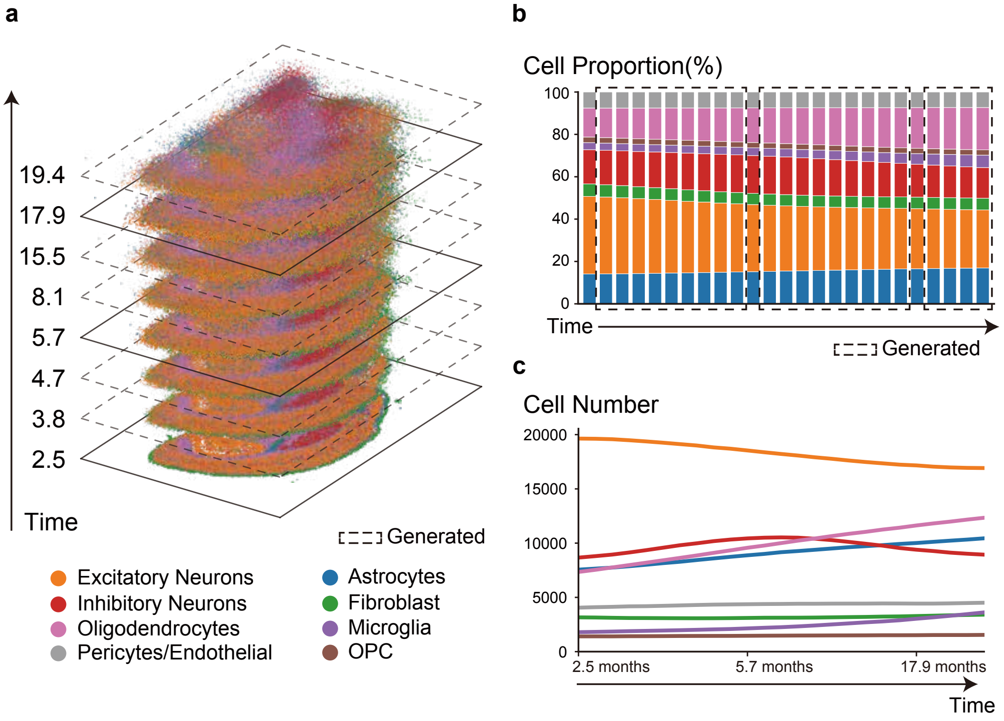

**Supplementary Figure S31.** Reconstructed AD progression in TgCRND8 mice. (a) Three-dimensional stacked visualization of observed and generated brain cell states. Solid planes mark the measured ages of 2.5, 5.7, and 17.9 months. Dashed planes mark generated states, including interpolation between observations and the displayed age of 19.4 months beyond the last observation. The latter is an extrapolation. (b) Observed and generated cell-type proportions. Dashed boxes mark generated time points. (c) Predicted cell numbers for each annotated population over the simulated trajectory.

### Supplementary Figure S32. Pattern 1 functional enrichment in the AD dataset

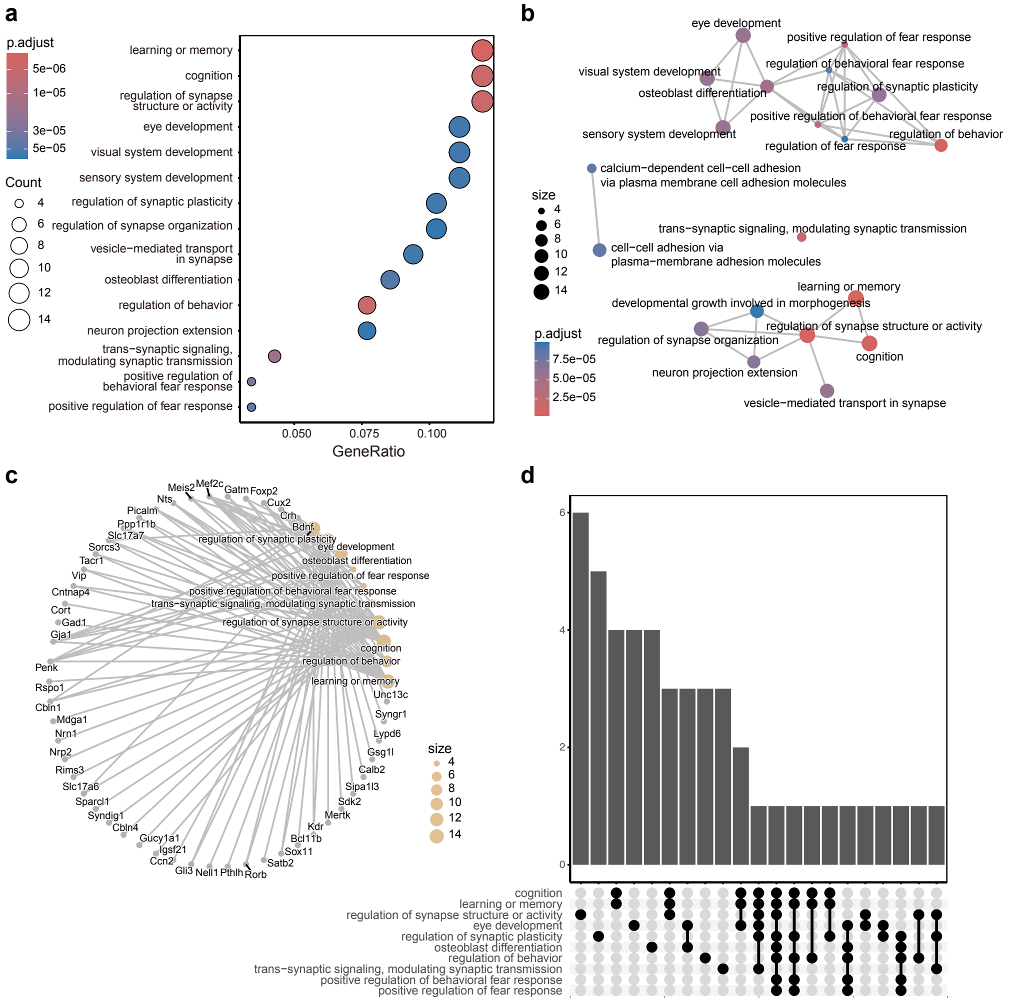

**Supplementary Figure S32.** Pattern 1 functional enrichment in the AD dataset. (a) Dot plot of enriched Gene Ontology terms for Pattern 1 genes. The x-axis represents the gene ratio, with dot size indicating gene counts and color representing the adjusted p-value. Enriched terms include cognition, learning or memory, and regulation of synapse structure or activity. (b) Enrichment map of the top Gene Ontology terms. Nodes represent terms and edges connect terms with shared gene members. Many connected terms fall under synaptic regulation, cognition, and behavioral regulation. (c) Gene-concept network showing the linkage between top enriched terms and their associated genes. (d) UpSet plot showing the overlap of genes across the top enriched Gene Ontology terms.

### Supplementary Figure S33. Pattern 4 functional enrichment in the AD dataset

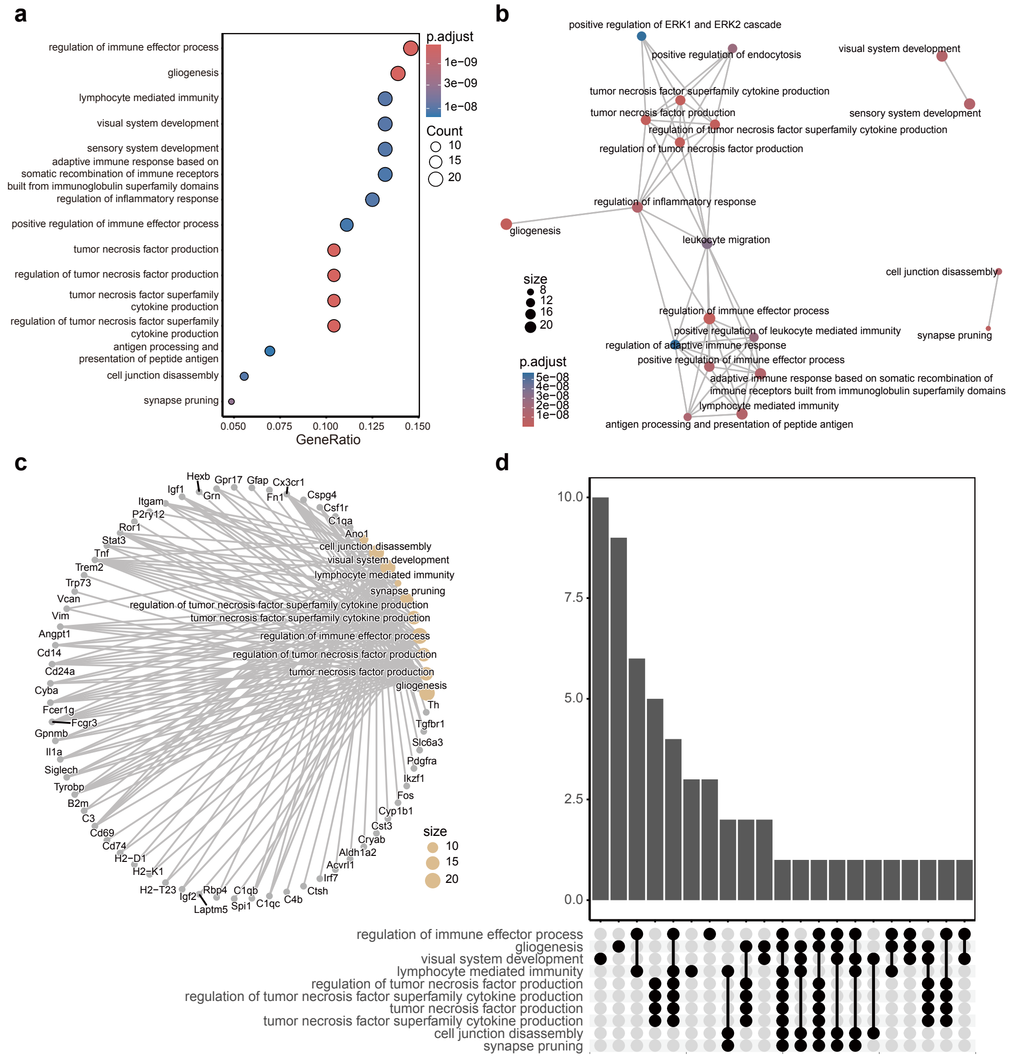

**Supplementary Figure S33.** Pattern 4 functional enrichment in the AD dataset. (a) Dot plot of enriched Gene Ontology terms for Pattern 4 genes. The x-axis represents the gene ratio, with dot size indicating gene counts and color representing the adjusted p-value. Enriched terms include immune effector processes, TNF-associated cytokine production, and inflammatory responses. (b) Enrichment map of the top Gene Ontology terms, with edges connecting terms that share gene members. The map is dominated by immune-related term clusters. (c) Gene-concept network showing the linkage between top enriched terms and their associated genes. (d) UpSet plot showing the overlap of genes across the top enriched Gene Ontology terms.

### Supplementary Figure S34. Temporal ligand activity and ligand–receptor networks in the AD dataset

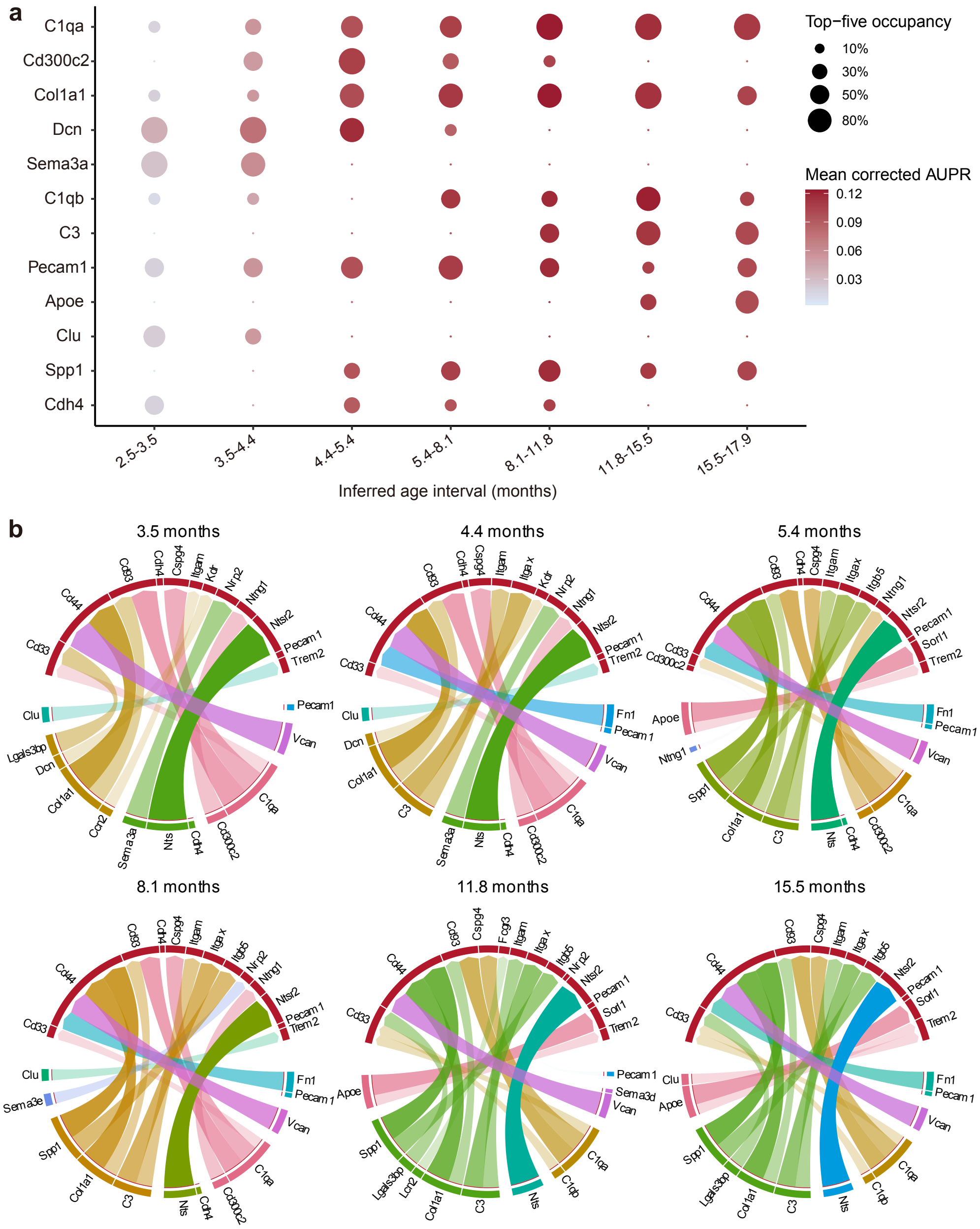

**Supplementary Figure S34.** Temporal ligand activity and ligand–receptor networks in the AD dataset. (a) NicheNet ligand activity across seven adjacent intervals of the reconstructed AD trajectory. For each interval, the 50 microglial genes with the largest positive reconstructed change were used as the target set. Dot size shows the fraction of analyses in which a ligand ranked among the top five, and color shows mean AUPR. (b) Ligand–receptor networks based on the NicheNet prior at six inferred ages. Prioritized ligands were linked to receptors expressed in microglia, with ribbon width proportional to the NicheNet prior weight.

#### Supplementary Figure S35. Functional-module changes after in silico Spp1 perturbation

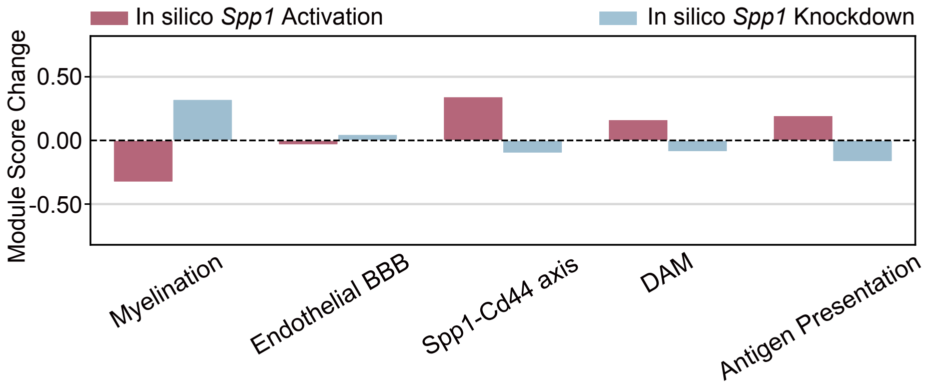

**Supplementary Figure S35.** Functional-module changes after in silico Spp1 perturbation. Changes in mean module scores at model time 2.5, beyond the last observed age of 17.9 months, relative to the unperturbed TgCRND8 reference. In silico Spp1 knockdown reduced the Spp1–Cd44, disease-associated microglia, and antigen-presentation modules and increased the myelination and endothelial–blood–brain barrier modules. Activation produced the opposite pattern. Module scores are means of Z-scored expression across detected module genes.

### Supplementary Figure S36. Observed and generated zebrafish development

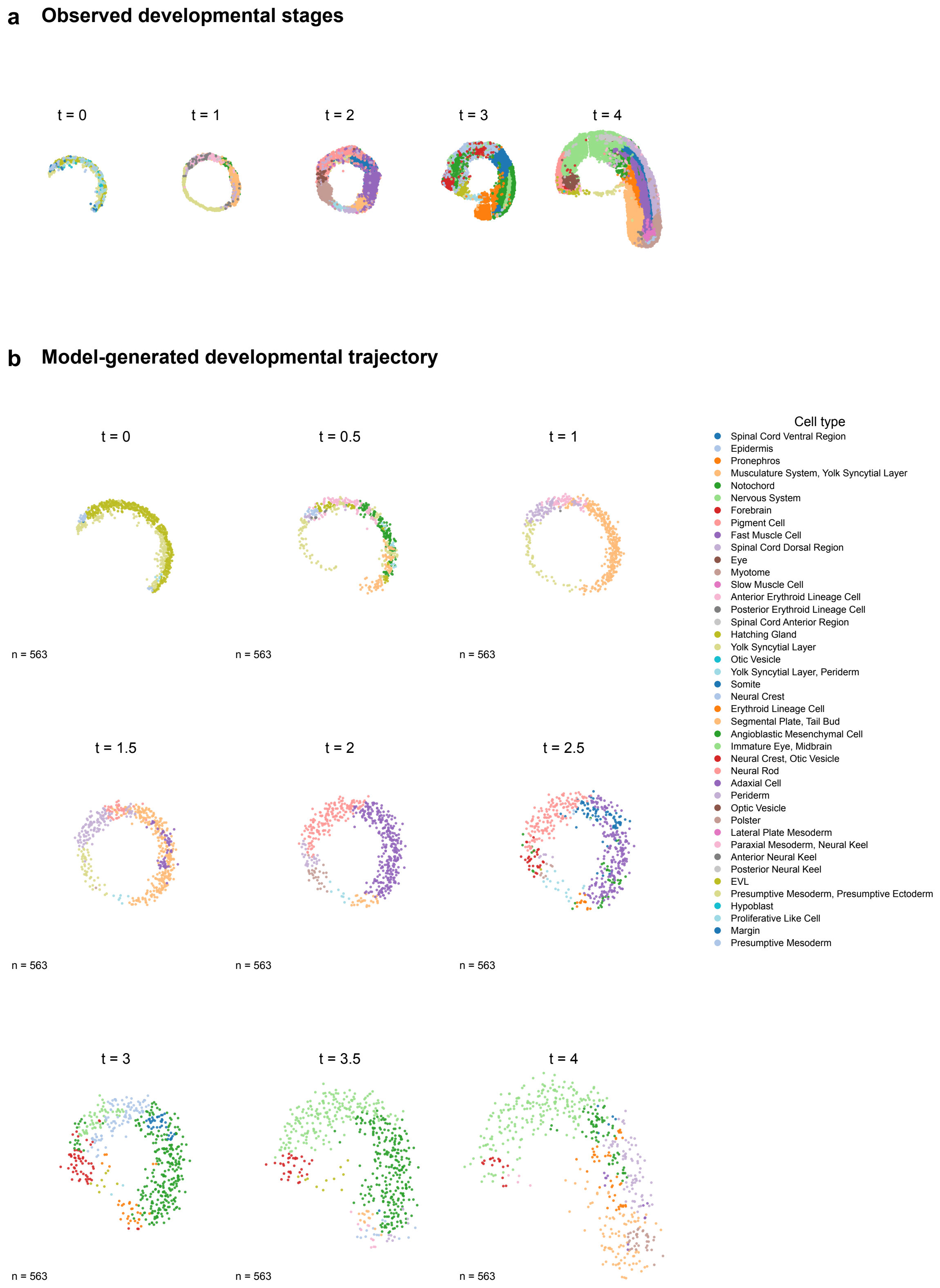

**Supplementary Figure S36.** Observed and generated zebrafish development. Observed spatial stages are shown above the generated trajectory. The generated population starts from the observed $t = 0$ cells and is propagated continuously through $t = 4$. Colors indicate predicted cell types.

#### Supplementary Figure S37. Growth-rate maps across observed zebrafish stages

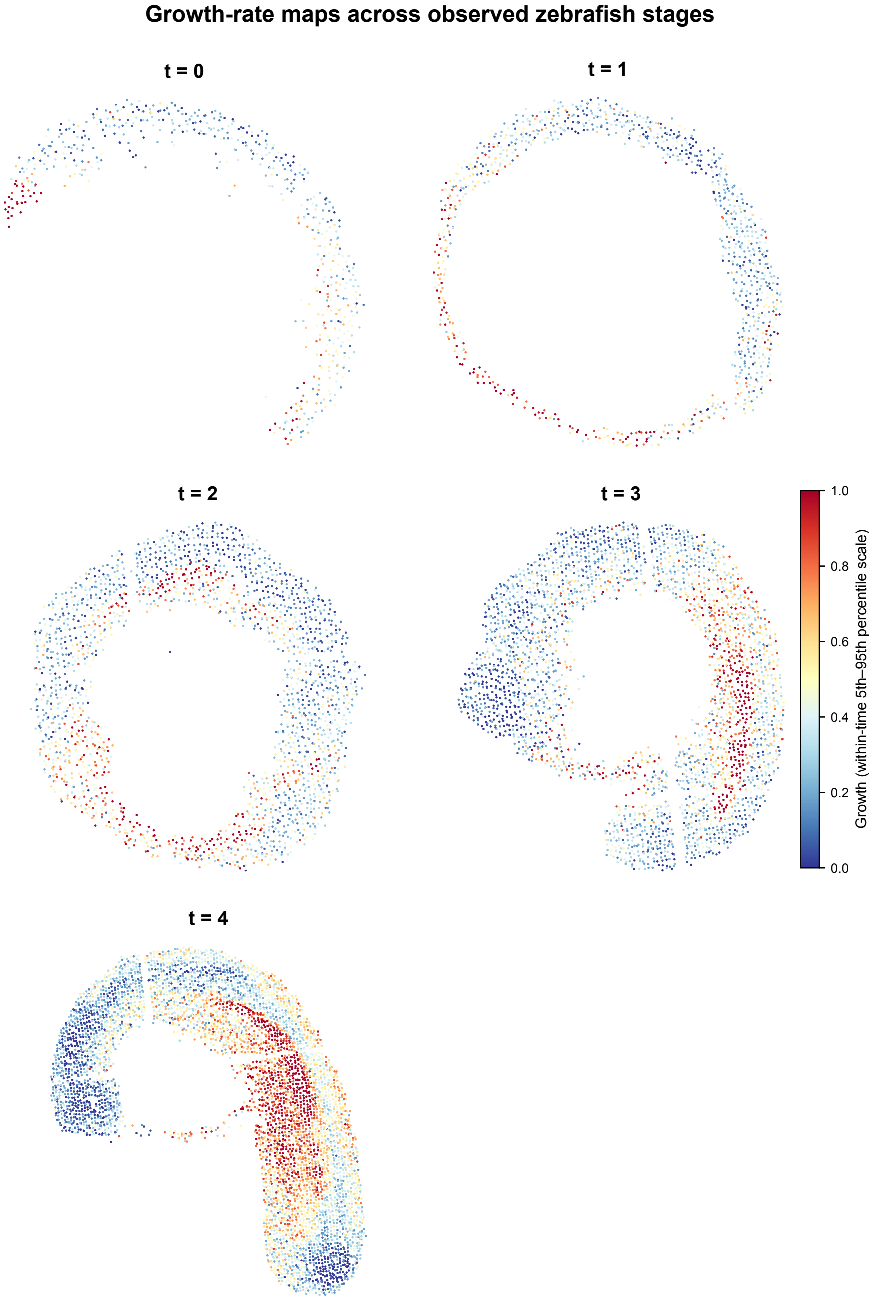

**Supplementary Figure S37.** Growth-rate maps across observed zebrafish stages. The learned growth field was evaluated on the observed cells at $t = 0, 1, 2, 3,$ and 4. Color values are scaled within each stage to show the spatial organization of relative growth.

#### Supplementary Figure S38. External and spatial comparison of learned interaction attention in zebrafish

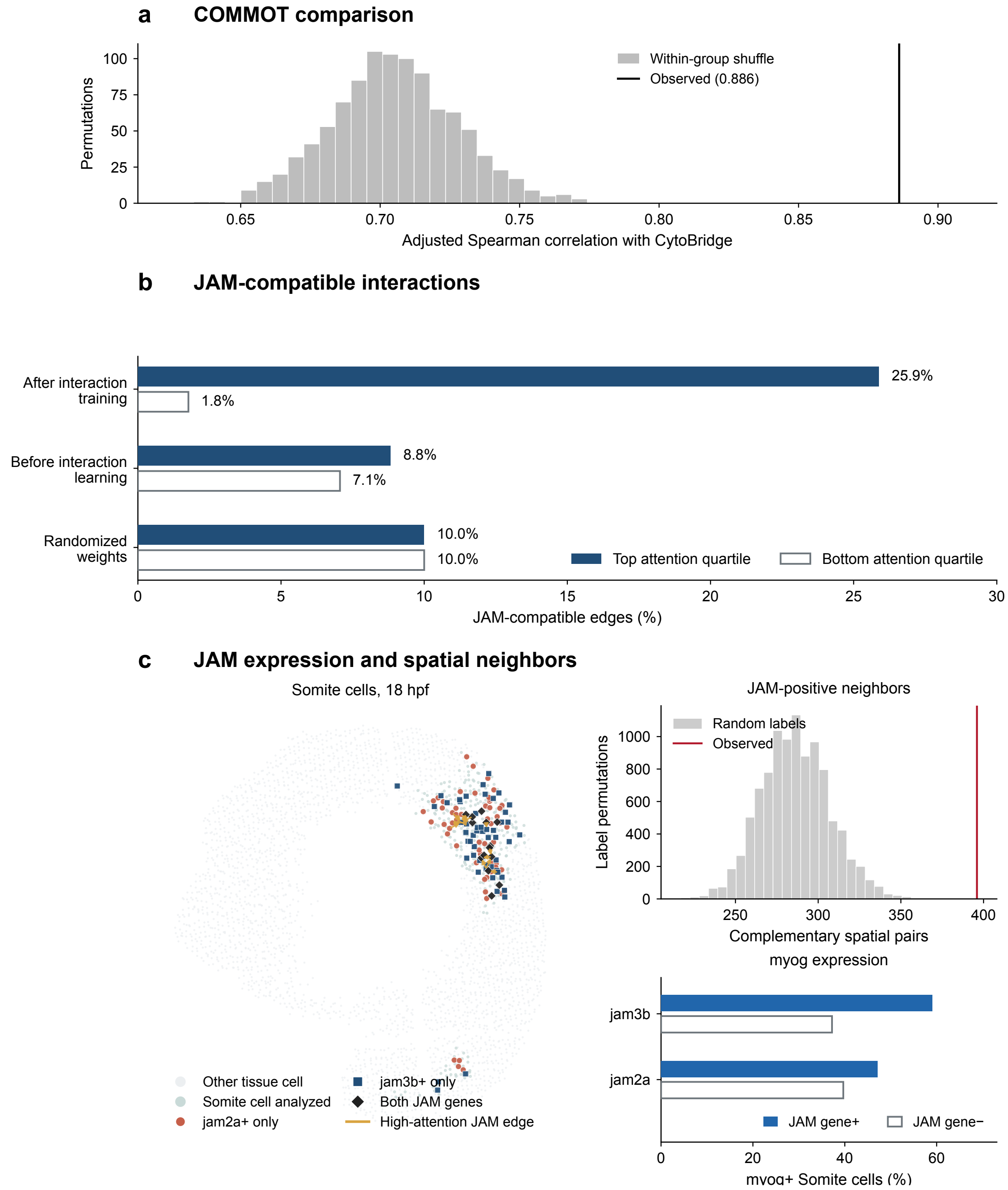

**Supplementary Figure S38.** External and spatial comparison of learned interaction attention in zebrafish. (a) Agreement between CytoBridge attention and COMMOT across all 361 directed cell-type pairs at the final observed model time ($t = 4$). The black line marks the observed Spearman correlation after adjustment for sender and receiver abundance, mean spatial distance, and whether the sender and receiver had the same cell type ($\rho = 0.886$). The gray histogram shows 1,000 permutations of the CytoBridge pair scores within 22 groups of pairs with similar abundance and distance and the same self-pair status, with COMMOT scores held fixed (one-sided permutation $P = 0.001$). (b) Comparison of attention on the same 677 model edges among 375 Somite cells at 18 hpf. A JAM-compatible edge had *jam2a* detected at one endpoint and *jam3b* at the other. In the trained model, 25.9% of edges in the highest attention quartile were JAM-compatible versus 1.8% in the bottom quartile. Controls use the model before interaction learning or shuffle its learned weights on the same cells and graph edges. (c) Spatial map highlighting the Somite cells, with neighboring cells shown for anatomical context. Gold arrows mark the 15 high-attention JAM-compatible model edges. The 396 spatially neighboring complementary $jam2a^{+}/jam3b^{+}$ Somite pairs were enriched 1.38-fold over 10,000 permutations that independently shuffled the *jam2a* and *jam3b* detection labels among the same Somite cells while retaining their positions and each gene's positive-cell count (Monte Carlo $P = 0.0001$). *jam3b* and *myog* co-detection had odds ratio 2.44 (Fisher $P = 0.0014$). The spatial and gene co-detection tests used the same 375 Somite cells (Supplementary Note S8.11).

### Supplementary Figure S39. Development after virtual YSL and EVL removal

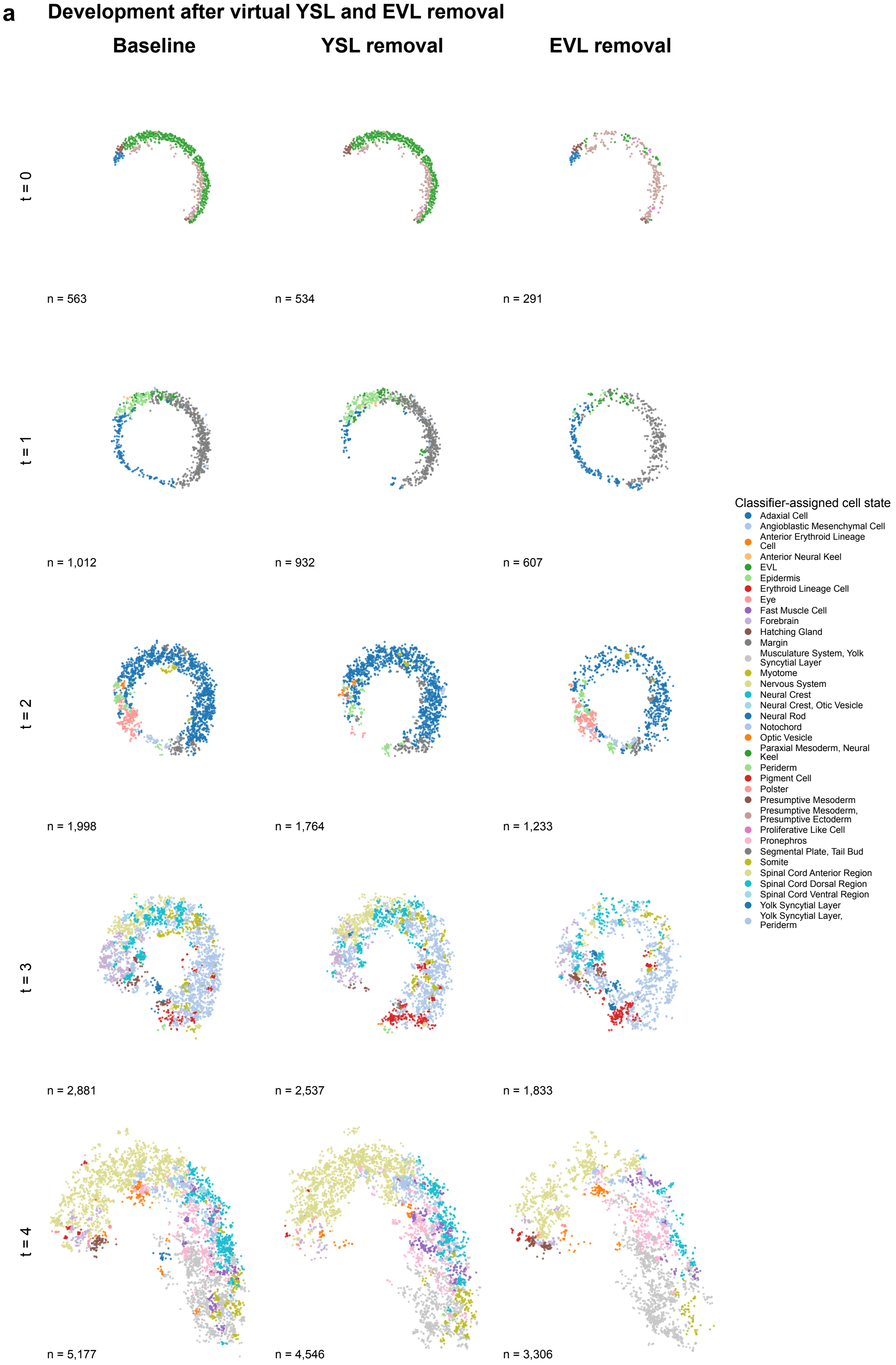

**Supplementary Figure S39.** Development after virtual YSL and EVL removal. Baseline, YSL-removal, and EVL-removal populations were propagated from the initial stage with the split SDE with growth. Colors indicate predicted cell types. YSL removal preserves more of the baseline morphology, whereas EVL removal produces a larger spatial change by the final stage.

### Supplementary Figure S40. Spatial divergence after virtual YSL and EVL removal

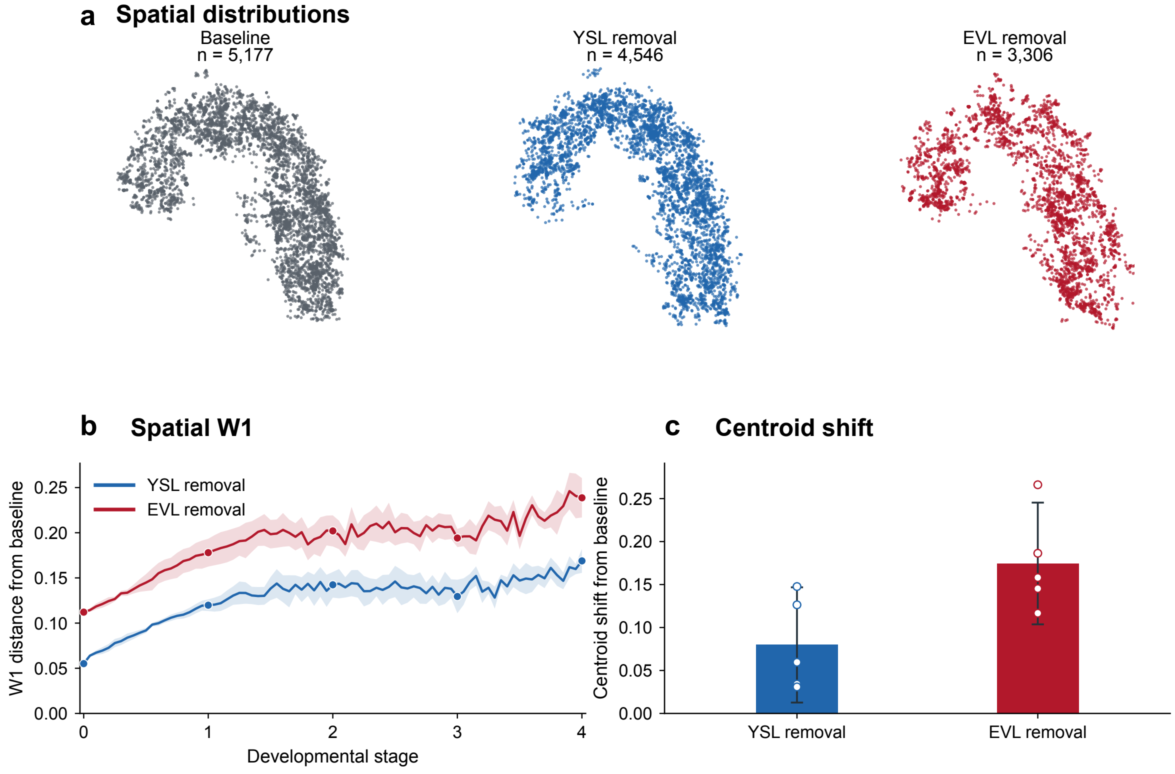

**Supplementary Figure S40.** Spatial divergence after virtual YSL and EVL removal. (a) Endpoint spatial distributions from one representative simulation. (b) Spatial Wasserstein-1 distance from the paired baseline across development. Lines and bands show mean ± s.e.m. across five paired simulations. (c) Endpoint centroid shift. Bars show the mean, error bars show the 95% confidence interval, and symbols show individual simulations. EVL removal produces the larger spatial divergence.

### Supplementary Figure S41. YSL-lineage gene dynamics

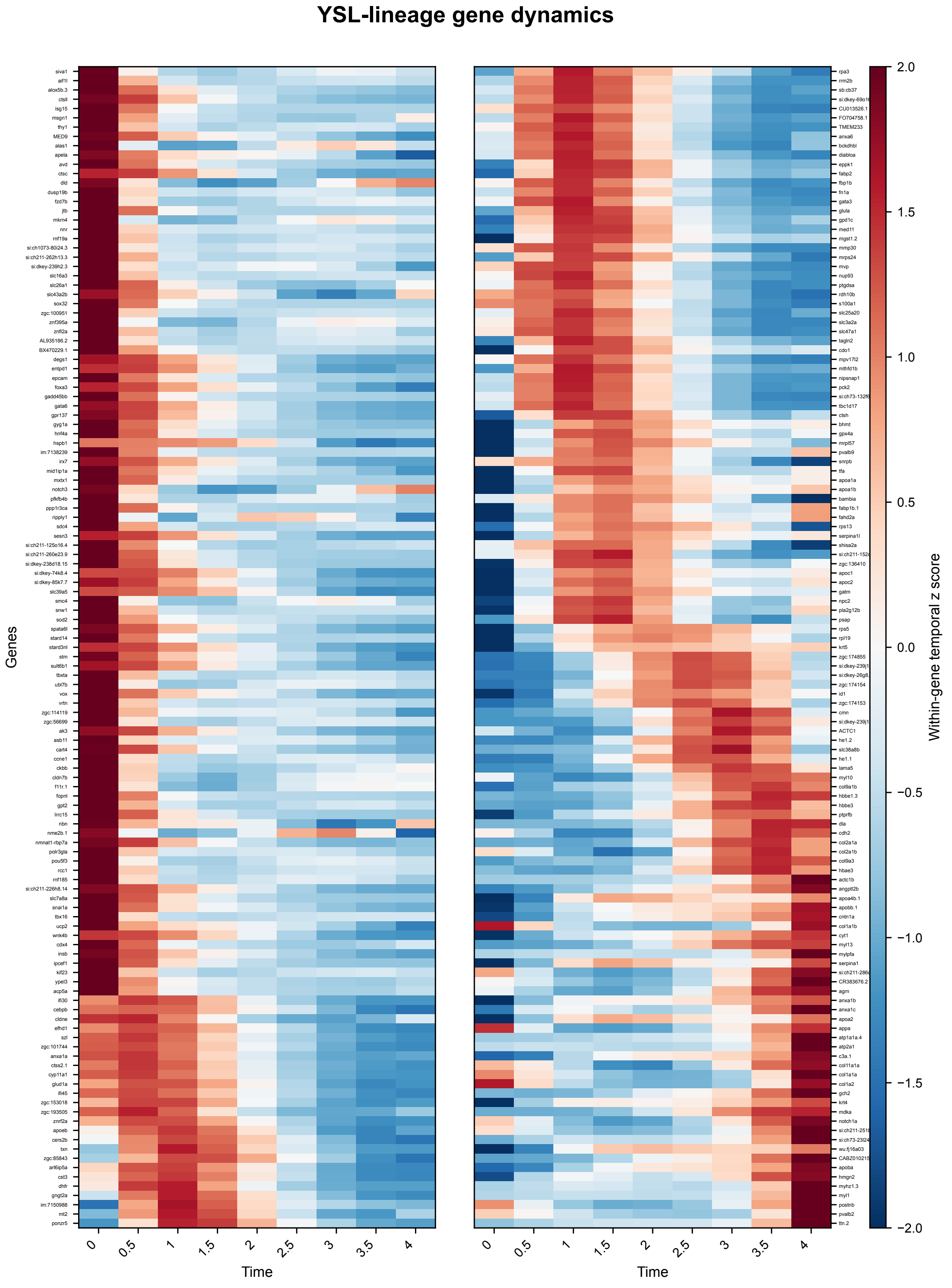

**Supplementary Figure S41.** YSL-lineage gene dynamics. Row-wise z-scored expression of 250 temporally variable genes in the descendants of the observed $t = 0$ YSL population. The initial lineage was propagated continuously to $t = 4$ with the continuous-weight formulation. Genes are ordered by their temporal expression pattern and displayed in two adjacent heat maps.

### Supplementary Figure S42. Inverse-PCA reconstruction of observed YSL expression

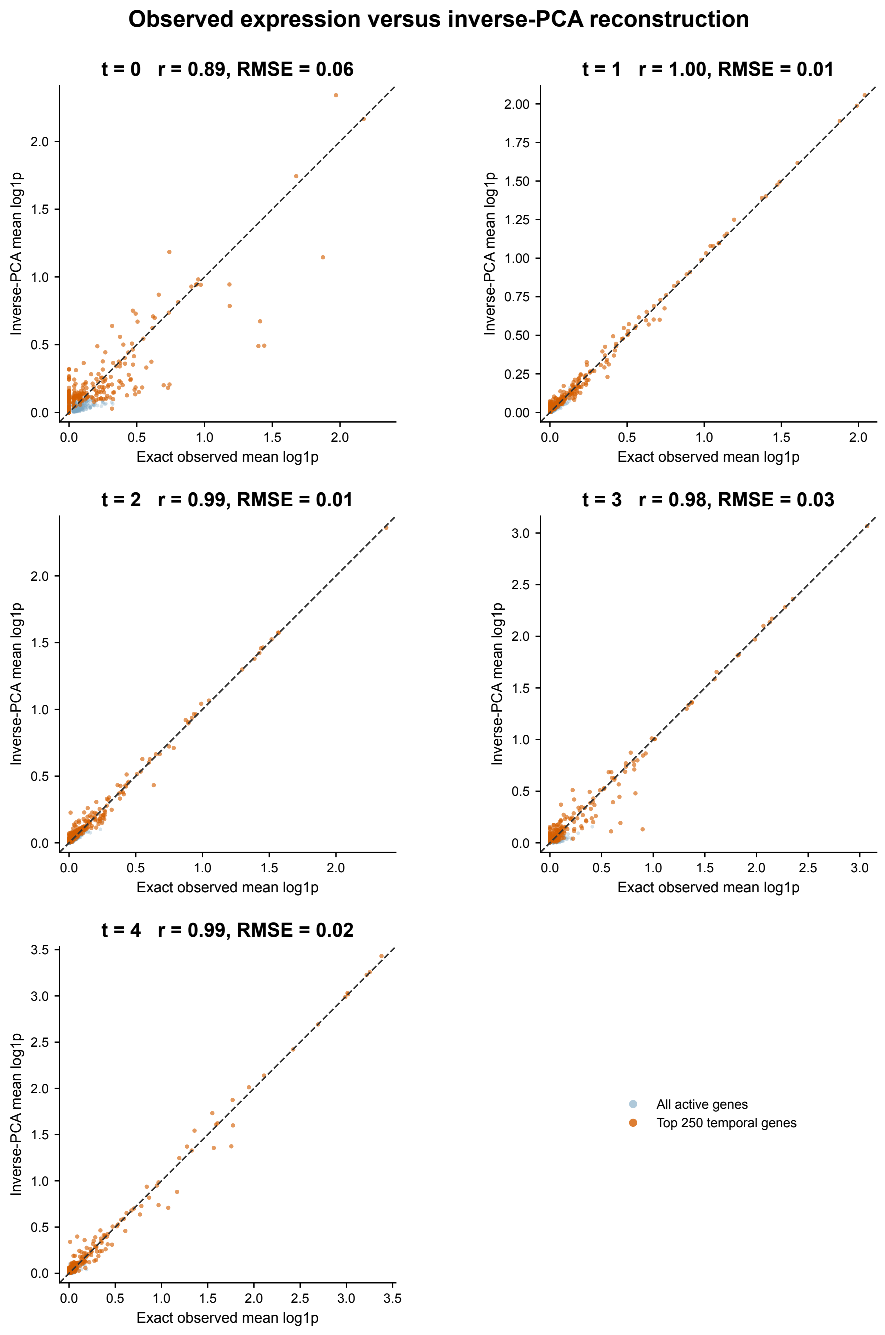

**Supplementary Figure S42.** Inverse-PCA reconstruction of observed YSL expression. Observed mean expression is compared with clipped inverse-PCA reconstruction at the five observed stages. Blue points show all analyzed genes and orange points show the 250 genes with greatest temporal variation.

#### Supplementary Figure S43. Representative six-stage training histories

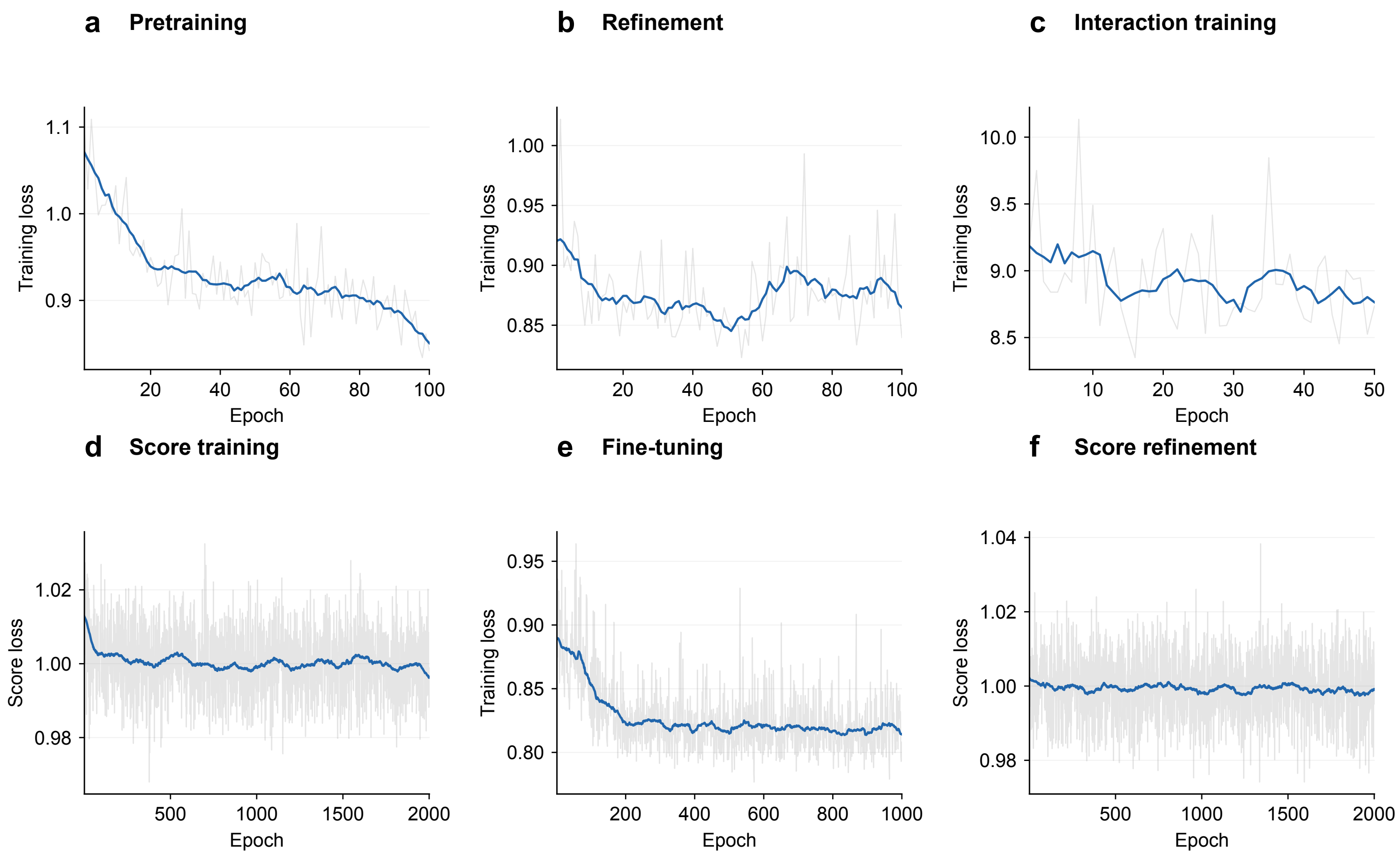

**Supplementary Figure S43.** Representative six-stage training histories. (a–f) Per-epoch objectives for Pretrain, Refine, Init interaction, Train score, Finetune, and Score refine in the ARTISTA full model. Thin lines show the recorded values and thick lines show centered moving means. The score-training stages optimize a score-matching loss, whereas the other stages report their corresponding composite training losses. Each stage is shown on its own scale. Training histories for all five datasets are included with the code.

### Supplementary Figure S44. Sensitivity to training-loss weights in zebrafish

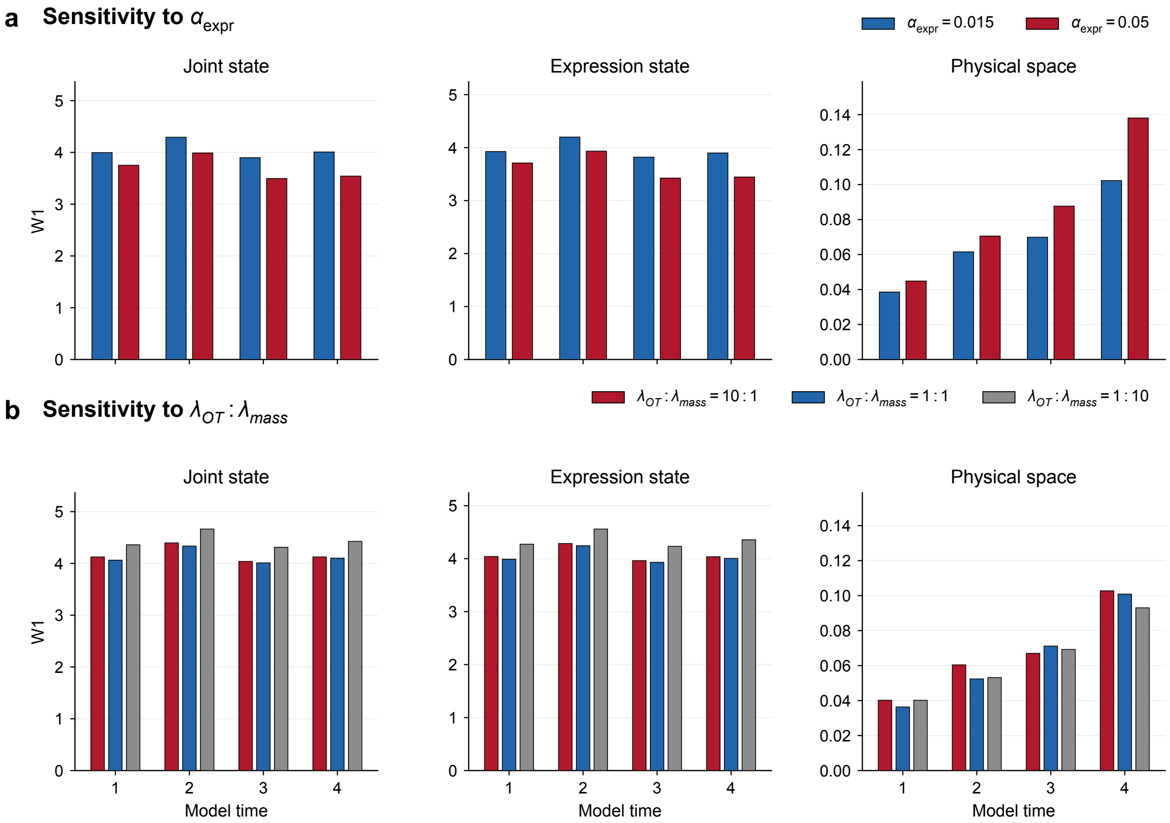

**Supplementary Figure S44.** Sensitivity to training-loss weights in zebrafish. Bars show Wasserstein-1 reconstruction distance from $t = 0$ (W1), averaged over observed model times 1–4; lower values indicate closer agreement with the observed distributions. (a) Expression-loss sensitivity comparing $\alpha_{\text{expr}} = 0.015$ and 0.05 with $\lambda_{\text{OT}} : \lambda_{\text{mass}} = 1 : 1$. (b) Transport-to-mass sensitivity comparing ratios of 10:1, 1:1, and 1:10 while holding $\lambda_{\text{OT}} + \lambda_{\text{mass}} = 20$ in the interaction stages. Training and evaluation are described in Supplementary Note S4.6.

### Supplementary Figure S45. Daughter-cell perturbation sensitivity in zebrafish

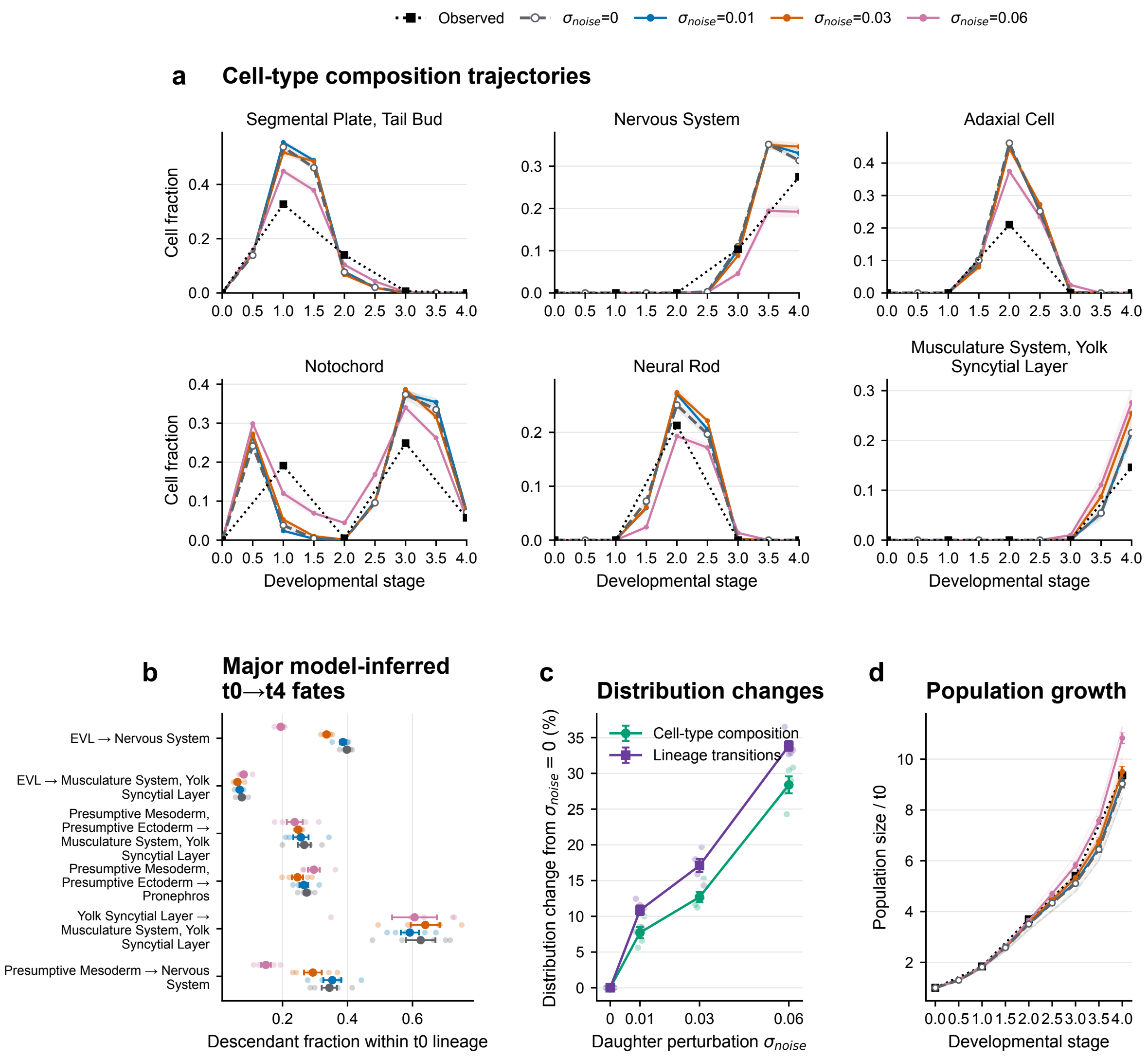

**Supplementary Figure S45.** Daughter-cell perturbation sensitivity in zebrafish. (a) Observed cell-type fractions at $t = 0, 1, 2, 3$, and 4 and simulated composition trajectories across daughter-cell perturbation scales. Observed values are connected across the five measured stages. (b) Major model-inferred lineage transitions from $t = 0$ to $t = 4$. (c) Changes in composition and lineage transitions at the final stage. (d) Simulated population growth. Increasing daughter-cell perturbation progressively changes composition and lineage outcomes. Curves and error bars show mean ± s.e.m. across five paired simulations.

### Supplementary Figure S46. Effect of spatial smoothing on cell-type assignments

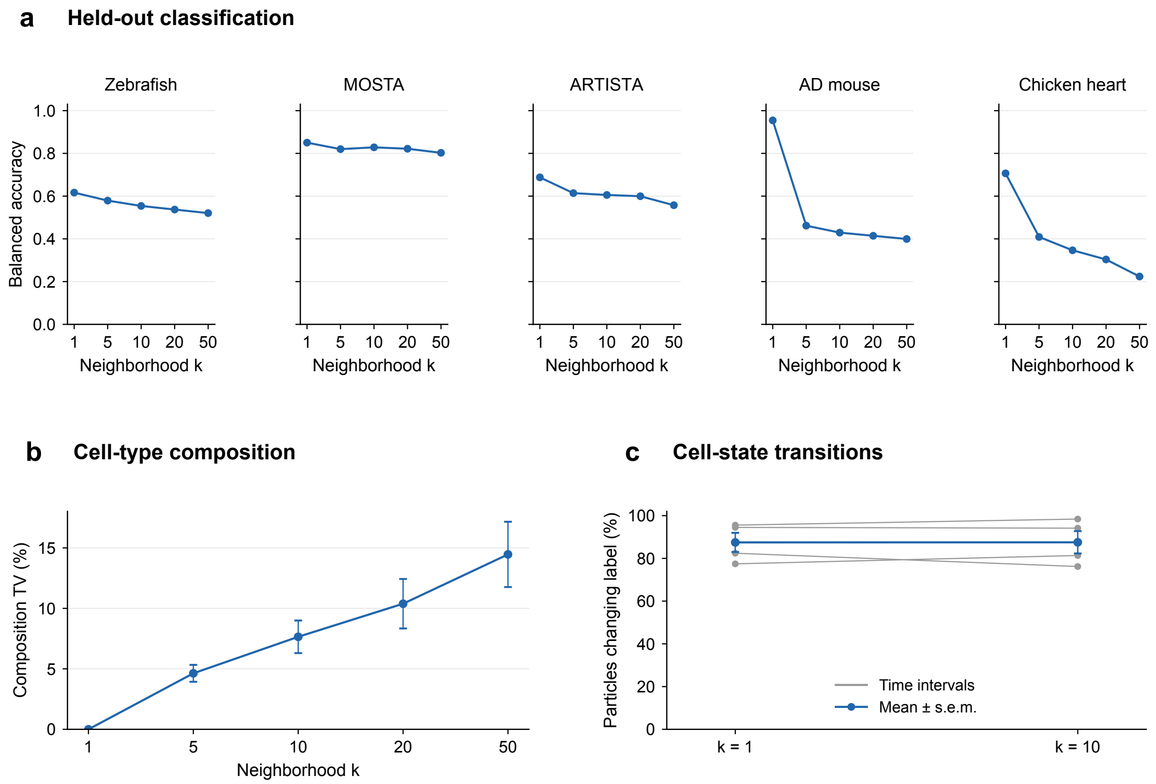

**Supplementary Figure S46.** Effect of spatial smoothing on cell-type assignments. (a) Held-out balanced accuracy after spatial voting with $k = 1, 5, 10, 20$, and 50 for classifiers trained independently in the five datasets. ARTISTA balanced accuracy was 0.688 at $k = 1$ and 0.605 at $k = 10$. Chicken-heart evaluation used the 13 cell-type annotations and gave balanced accuracy 0.707 at $k = 1$ and 0.347 at $k = 10$. (b) Total-variation distance between zebrafish cell-type compositions obtained at each $k$ and the unsmoothed $k = 1$ composition across nine generated time points. Points and error bars show the mean and s.e.m. across those time points. (c) Fraction of 563 particles that changed predicted type over each of four consecutive zebrafish intervals at $k = 1$ and $k = 10$; gray lines connect the same interval, blue circles show the mean, and error bars show s.e.m. At $k = 10$, mean zebrafish composition TV was 7.6%, whereas the mean transition fraction was 87.48% at $k = 1$ and 87.52% at $k = 10$, a difference of 0.04 percentage points. For cell-type maps and proportions, we used $k = 10$ for zebrafish, MOSTA, and ARTISTA and $k = 1$ for AD mouse and chicken heart, where spatial voting reduced held-out classification accuracy.

### Supplementary Figure S47. Sensitivity of ligand–receptor rankings to multi-subunit aggregation

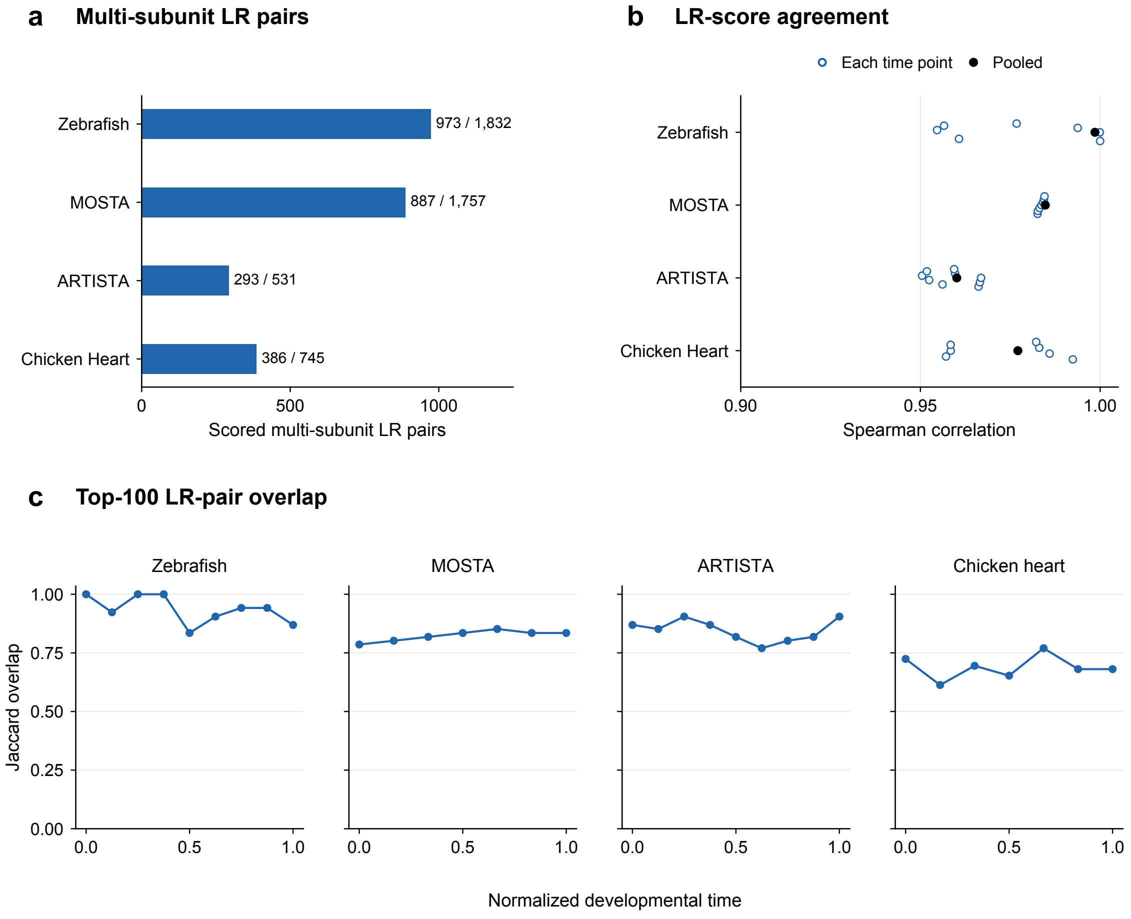

**Supplementary Figure S47.** Sensitivity of ligand–receptor rankings to multi-subunit aggregation. (a) Numbers of multi-subunit LR pairs. Annotations give the multi-subunit and total scored pair counts. (b) Spearman rank agreement between the minimum-subunit rule and the geometric mean. Open circles show individual computed model times, and filled black circles combine all LR pair–time observations. (c) Jaccard overlap of the top 100 LR pairs across normalized developmental time. Each circle is one computed model time. Within each dataset, the two calculations used the same model outputs and scored LR pairs. Both rules assign zero activity if any required subunit has zero expression. The LR scores and aggregation comparison are described in Supplementary Notes S8.17 and S8.20.

### References

[1] Zhang, Z. *et al.* Deciphering cell-fate trajectories using spatiotemporal single-cell transcriptomic data. *npj Systems Biology and Applications* **12**, 2 (2026).

[2] Monge, G. Mémoire sur la théorie des déblais et des remblais. *Mémoires de l'Académie royale des sciences* 666–704 (1781).

[3] Kantorovich, L. V. On the translocation of masses. *Doklady Akademii Nauk SSSR* **37**, 227–229 (1942).

[4] Benamou, J.-D. & Brenier, Y. A computational fluid mechanics solution to the Monge-Kantorovich mass transfer problem. *Numerische Mathematik* **84**, 375–393 (2000).

[5] Chizat, L., Peyré, G., Schmitzer, B. & Vialard, F.-X. An interpolating distance between optimal transport and Fisher–Rao metrics. *Foundations of Computational Mathematics* **18**, 1–44 (2018).

[6] Chizat, L., Peyré, G., Schmitzer, B. & Vialard, F.-X. Unbalanced optimal transport: Dynamic and Kantorovich formulations. *Journal of Functional Analysis* **274**, 3090–3123 (2018).

[7] Schrödinger, E. Sur la théorie relativiste de l'électron et l'interprétation de la mécanique quantique. *Annales de l'Institut Henri Poincaré* **2**, 269–310 (1932).

[8] Léonard, C. A survey of the Schrödinger problem and some of its connections with optimal transport. *Discrete and Continuous Dynamical Systems - Series A* **34**, 1533–1574 (2014).

[9] Backhoff, J., Conforti, G., Gentil, I. & Léonard, C. The mean field Schrödinger problem: Ergodic behavior, entropy estimates and functional inequalities. *Probability Theory and Related Fields* **178**, 475–530 (2020).

[10] Zhang, Z., Li, T. & Zhou, P. Learning stochastic dynamics from snapshots through regularized unbalanced optimal transport. In *The Thirteenth International Conference on Learning Representations* (2025).

[11] Zhang, Z., Wang, Z., Sun, Y., Li, T. & Zhou, P. Modeling cell dynamics and interactions with unbalanced mean field Schrödinger bridge. In *Advances in Neural Information Processing Systems*, vol. 38, 154733–154773 (2025).

[12] Chen, R. T. Q., Rubanova, Y., Bettencourt, J. & Duvenaud, D. K. Neural ordinary differential equations. In *Advances in Neural Information Processing Systems*, vol. 31 (2018).

[13] Mantri, M. *et al.* Spatiotemporal single-cell RNA sequencing of developing chicken hearts identifies interplay between cellular differentiation and morphogenesis. *Nature Communications* **12**, 1771 (2021).

[14] Wei, X. *et al.* Single-cell Stereo-seq reveals induced progenitor cells involved in axolotl brain regeneration. *Science* **377**, eabp9444 (2022).

[15] Chen, A. *et al.* Spatiotemporal transcriptomic atlas of mouse organogenesis using DNA nanoball-patterned arrays. *Cell* **185**, 1777–1792.e21 (2022).

[16] 10x Genomics. Xenium In Situ analysis of Alzheimer's disease mouse model brain coronal sections from one hemisphere over a time course (2023). URL https://www.10xgenomics.com/datasets/xenium-in-situ-analysis-of-alzheimers-disease-mouse-model-brain-coronal-sections-from-one-hemisphere-over-a-time-course-1-standard.

[17] Liu, C. *et al.* Spatiotemporal mapping of gene expression landscapes and developmental trajectories during zebrafish embryogenesis. *Developmental Cell* **57**, 1284–1298.e5 (2022).

[18] Peng, Q., Zhou, P. & Li, T. stVCR: spatiotemporal dynamics of single cells. *Nature Methods* **23**, 542–553 (2026). URL https://doi.org/10.1038/s41592-026-03010-3.

[19] Klein, D. *et al.* Mapping cells through time and space with moscot. *Nature* **638**, 1065–1075 (2025). URL https://doi.org/10.1038/s41586-024-08453-2.

[20] Zeira, R., Land, M., Strzalkowski, A. & Raphael, B. J. Alignment and integration of spatial transcriptomics data. *Nature Methods* **19**, 567–575 (2022). URL https://doi.org/10.1038/s41592-022-01459-6.

[21] Liu, X., Zeira, R. & Raphael, B. J. Partial alignment of multislice spatially resolved transcriptomics data. *Genome Research* **33**, 1124–1132 (2023).

[22] Qiu, X. *et al.* Spatiotemporal modeling of molecular holograms. *Cell* **187**, 7351–7373.e61 (2024). URL https://doi.org/10.1016/j.cell.2024.10.011.

[23] Huguet, G. *et al.* Manifold interpolating optimal-transport flows for trajectory inference. *Advances in Neural Information Processing Systems* **35**, 29705–29718 (2022).

[24] Sun, X. *et al.* MIOFlow 2.0: A unified framework for inferring cellular stochastic dynamics from single cell and spatial transcriptomics data. arXiv preprint arXiv:2603.22564 (2026).

[25] Huizing, G.-J. *et al.* STORIES: learning cell fate landscapes from spatial transcriptomics using optimal transport. *Nature Methods* **23**, 522–531 (2026). URL https://doi.org/10.1038/s41592-025-02855-4.

[26] Schiebinger, G. *et al.* Optimal-Transport Analysis of Single-Cell Gene Expression Identifies Developmental Trajectories in Reprogramming. *Cell* **176**, 928–943.e22 (2019). URL https://doi.org/10.1016/j.cell.2019.01.006.

[27] Shen, X. *et al.* Inferring cell trajectories of spatial transcriptomics via optimal transport analysis. *Cell Systems* **16**, 101194 (2025). URL https://doi.org/10.1016/j.cels.2025.101194.

[28] Flamary, R. *et al.* POT: Python optimal transport. *Journal of Machine Learning Research* **22**, 1–8 (2021).

[29] Weinreb, C., Rodriguez-Fraticelli, A., Camargo, F. D. & Klein, A. M. Lineage tracing on transcriptional landscapes links state to fate during differentiation. *Science* **367**, eaaw3381 (2020). URL https://doi.org/10.1126/science.aaw3381.

[30] Jin, S. *et al.* Inference and analysis of cell-cell communication using CellChat. *Nature Communications* **12**, 1088 (2021). URL https://doi.org/10.1038/s41467-021-21246-9.

[31] Cang, Z. *et al.* Screening cell–cell communication in spatial transcriptomics via collective optimal transport. *Nature Methods* **20**, 218–228 (2023). URL https://doi.org/10.1038/s41592-022-01728-4.

[32] Qiu, Q. *et al.* Massively parallel and time-resolved RNA sequencing in single cells with scNT-seq. *Nature Methods* **17**, 991–1001 (2020). URL https://doi.org/10.1038/s41592-020-0935-4.

[33] Raghavan, V., Zheng, Y., Li, Y. & Ding, J. Harnessing agent-based frameworks in CellAgentChat to unravel cell–cell interactions from single-cell and spatial transcriptomics. *Genome Research* **35**, 1646–1663 (2025).

[34] Powell, G. T. & Wright, G. J. Jamb and Jamc are essential for vertebrate myocyte fusion. *PLOS Biology* **9**, e1001216 (2011).

[35] Ganassi, M. *et al.* Myogenin promotes myocyte fusion to balance fibre number and size. *Nature Communications* **9**, 4232 (2018).

[36] Raghavan, V., Yoon, B., Fonseca, G. J., Li, Y. & Ding, J. Dissecting and steering cell dynamics using spatially-informed RNA velocity with veloAgent. *Molecular Systems Biology* **22**, 1180–1200 (2026).
